# Chromatin structure influences rate and spectrum of spontaneous mutations in *Neurospora crassa*

**DOI:** 10.1101/2022.03.13.484164

**Authors:** Mariana Villalba de la Peña, Pauliina A. M. Summanen, Martta Liukkonen, Ilkka Kronholm

## Abstract

While mutation rates have been extensively studied, variation in mutation rates throughout the genome is poorly understood. To understand patterns of genetic variation, it is important to understand how mutation rates vary. Chromatin modifications may be an important factor in determining variation in mutation rates in eukaryotic genomes. To study variation in mutation rates, we performed a mutation accumulation experiment in the filamentous fungus *Neurospora crassa*, and sequenced the genomes of the 40 MA lines that had been propagated asexually for approximately 1015 [1003, 1026] mitoses. We detected 1322 mutations in total, and observed that the mutation rate was higher in regions of low GC, in domains of H3K9 trimethylation, in centromeric regions, and in domains of H3K27 trimethylation. The rate of single nucleotide mutations in euchromatin was 2.46 [2.19, 2.77] × 10^−10^. In contrast, the mutation rate in H3K9me3 domains was tenfold higher: 2.43 [2.25, 2.62] × 10^−9^. We also observed that the spectrum of single nucleotide mutations was different between H3K9me3 and euchromatic domains. Our statistical model of mutation rate variation predicted a moderate amount of extant genetic variation, suggesting that the mutation rate is an important factor in determining levels of natural genetic variation. Furthermore, we characterized mutation rates of structural variants, complex mutations, and the effect of local sequence context on the mutation rate. Our study highlights that chromatin modifications are associated with mutation rates, and accurate evolutionary inferences should take variation in mutation rates across the genome into account.

## Introduction

New mutations are the source of all genetic diversity, and evolutionary change ultimately depends on the input of new mutations into the population. However, organisms also pay a substantial cost for their ability to evolve, as deleterious mutations are more common than beneficial mutations (Eyre-Walker and Keightley, 2007), and mutations can lead to adverse outcomes, such as a decline in fitness, hereditary diseases, and cancer. Therefore, knowledge of the rates and spectrum of spontaneous mutations is fundamental to our understanding of evolution and certain aspects of medicine.

Spontaneous mutations are rare events that were previously difficult to study. However, new sequencing technologies have made it possible to capture a large number of spontaneous mutations for analysis (Katju and Bergthorsson, 2018). Mutation rates can now be estimated through direct observations by sequencing mutation accumulation lines or parent-offspring trios (Ossowski et al., 2010; Ness et al., 2012; Keightley et al., 2014; Zhu et al., 2014; Sung et al., 2015; Ness et al., 2015; Keightley et al., 2015; Keith et al., 2016; Wang et al., 2020). These studies have produced highly precise estimates of the rate and spectrum of spontaneous mutations.

The process of mutation is stochastic, but not all mutations are equally likely. While this has been appreciated for a long time for certain classes of mutations, such as transitions and transversions, there is also variation in mutation rates that seems to depend on the structural features of the genome, such as the organization of chromatin (Makova and Hardison, 2015). In particular, positioning of nucleosomes (Tolstorukov et al., 2011; Chen et al., 2012; Li and Luscombe, 2020) and chromatin structure have a strong influence on mutation rates (Schuster-Böckler and Lehner, 2012; Polak et al., 2015; Weng et al., 2019; Monroe et al., 2022). Chromatin structure is associated with chemical modifications of histone H3. In particular, the methylation status of certain lysine residues, such as H3K9 and H3K27 methylation, is associated with closed and silenced chromatin, called heterochromatin, while the methylation of H3K36 is associated with open actively transcribed chromatin, known as euchromatin (Kouzarides, 2007). Heterochromatin appears to have higher mutation rates than euchromatin (Makova and Hardison, 2015). In addition, the local sequence context, such as GC-content, also has a strong effect on mutation rates (Makova and Hardison, 2015; Ness et al., 2015; Sung et al., 2015). Although we know that the chromatin structure can shape mutation rates, most data come from humans and a few model species.

Furthermore, to what extent variation in mutation rates determines patterns of observed genetic diversity, along with other evolutionary mechanisms, is understood mostly from population genetic data, rather than from direct observations of mutation, with a few exceptions, e.g., Monroe et al. (2022). There are some tests for selection, such as *d_N_/d_S_* ratios, which are not affected by the mutation rate. However, tests based on the site frequency spectrum to infer mutational effects or demography can benefit from data about the mutation rate if it is available (Keightley and Eyre-Walker, 2007), especially if the goal is to examine different categories of genes or regions of the genome. Furthermore, in order to understand how different evolutionary forces, such as background selection, gene conversion, demographic processes, and adaptive evolution, jointly shape patterns of diversity across the genome, obtaining mutation rate estimates will allow us to parameterize population genetic models (Campos et al., 2017; Castellano et al., 2020; Johri et al., 2020). Ultimately, taking all possible effects into account will allow us to make better estimates of how natural selection shapes genetic diversity.

To examine patterns of variation in mutation rates, we performed a mutation accumulation (MA) experiment (Halligan and Keightley, 2009) in the filamentous fungus *Neurospora crassa*. In MA experiments, an ancestor is split into multiple lines, and these lines are bottlenecked every generation, which minimizes the efficiency of natural selection. (Figure 1A). This way, even deleterious mutations can accumulate in these lines (Halligan and Keightley, 2009). We sequenced the genomes of these mutation accumulation lines using short-read sequencing. *Neurospora crassa* is a filamentous fungus with a facultative sexual cycle, producing both asexual and sexual spores. It has a small genome of 42 Mb, and the vegetative mycelium is haploid. Furthermore, *N. crassa* has a genome defence mechanism called repeat-induced point mutation (RIP), which detects duplicated regions of the genome in premeiotic cells, and induces C → T transitions in the duplicated sequences (Selker, 1990). Because RIP does not induce the same specific mutations in both copies, large repeated arrays are seldom perfect in *N. crassa* as the duplicated sequences diverge from each other due to RIP. As the sequences diverge, the efficiency of RIP decreases (Cambareri et al., 1991). The existence of imperfect repeats makes it possible to use short-read sequencing to sequence repetitive regions, e.g., centromeric regions (Smith et al., 2011), where read mapping is normally difficult in plants and animals.

**Figure 1:**
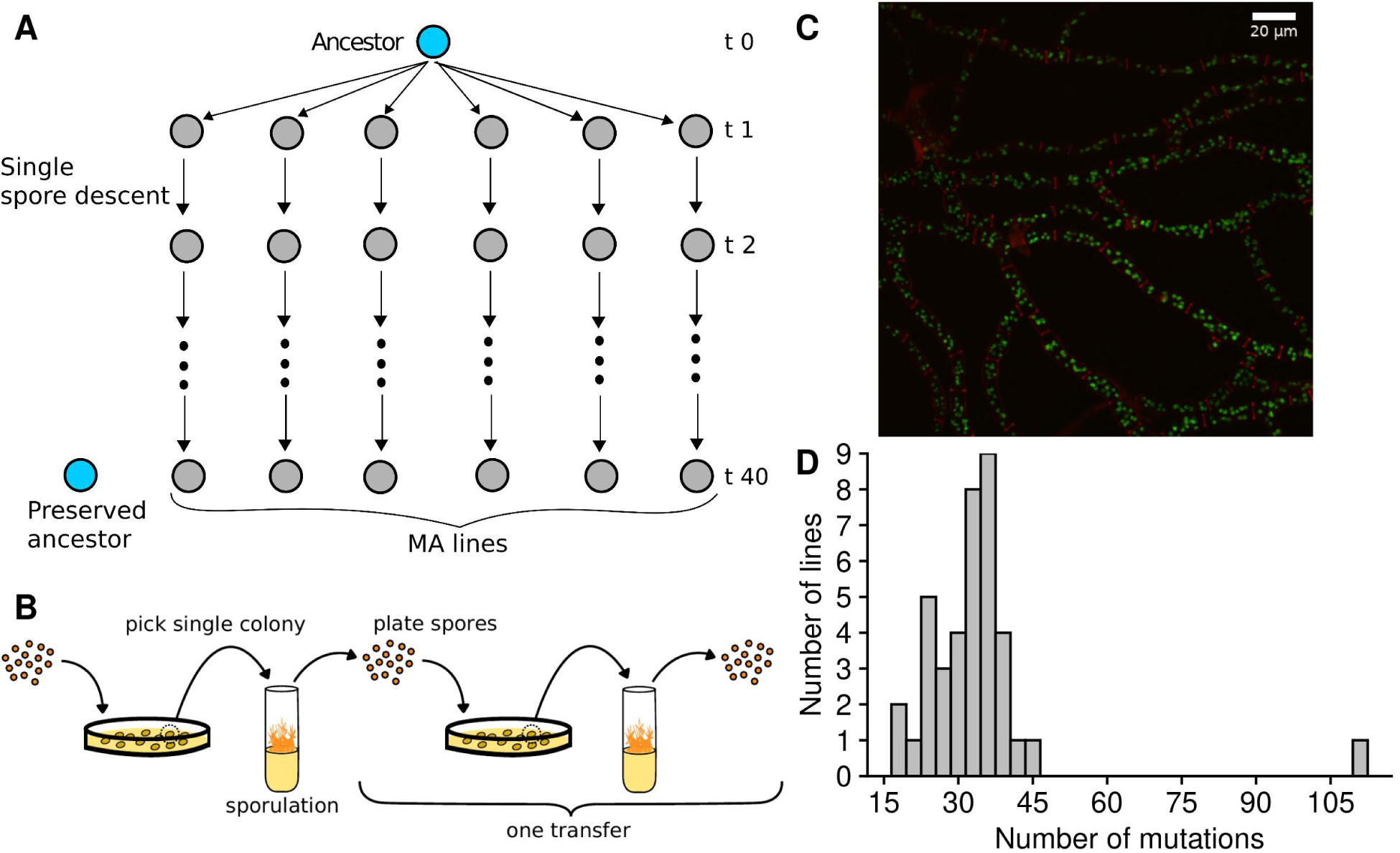
A) In the mutation accumulation experiment, ancestors were split into multiple lines, which were propagated via single spore descent. B) Lines in the MA experiment were transferred by always picking a colony originating from a single spore from a plate, moving this to a test tube with Vogel’s Medium to allow sporulation, then diluting spores and spreading them to a sorbose plate. C) Micrograph of *N. crassa* mycelium, showing nuclei in fluorescent green and cell walls in red. D) Distribution of mutations in the MA lines.

A previous study investigated the mutation rate during sexual reproduction in *N. crassa*, and revealed that the mutation rate is especially high in regions of the genome targeted by RIP (Wang et al., 2020). However, only a small number of mutations was collected during asexual reproduction; not enough to infer variation in the mutation rate across the genome. Our study complements Wang et al. (2020) by allowing us to characterize the determinants of the mutation rate and spectrum during asexual reproduction, when RIP is not active. We used information on the chromatin structure of *N. crassa* to model variation in the mutation rate across the genome. We also resequenced strains obtained from natural populations, and compared the predictions of our mutation model to patterns of natural genetic variation in order to assess whether the natural genetic variation reflects the observed mutation rate variation. Furthermore, we examined the effect of local sequence context on the mutation rate, and characterized patterns of structural variants and complex mutations.

## Results

We initiated the MA experiment with two ancestors that were isogenic, except for the mating type locus. We split both ancestors into 20 lines, giving 40 MA lines in total. We then plated asexual spores (conidia) on plates, picked a single colony, transferred this colony into a test tube, and let the mycelium grow and make asexual spores. We subsequently plated these spores again to isolate a single spore, and this process was repeated for 40 transfers (Figure 1B).

### Number of mitoses in the experiment

*Neurospora crassa* is a filamentous organism, and it does not have a defined germline. All parts of the mycelium are capable of producing structures that produce asexual spores. Thus, the number of transfers during the MA experiment does not correspond to a generation in a natural way. Therefore, a reasonable unit for measuring the mutation rate is the number of mutations per mitosis.

We estimated the number of mitoses the MA lines went through based on the counts of nuclei in different phases of a transfer (Figure 1B, C). Based on our estimate, the MA lines went through 25 [25, 26] (median, [95% HPDI]) mitoses in a single transfer. For the whole experiment of 40 transfers, this means that the MA lines went through 1015 [1003, 1026] mitoses.

### Mutations in the MA lines

To detect mutations in the MA lines, we sequenced the genomes of the MA ancestors and the MA lines using short-read sequencing with 150 bp paired-end reads. The sequencing depth was more than 30x on average, and approximately 98% of reads were mapped to the reference genome (Table S1). The reference genome of *N. crassa* contains 41 108 926 bp and we called 98.7% of those bases on average. We used a pipeline based on the GATK best practices to call the mutations, followed by a manual inspection of alignments for each mutation.

After sequencing the MA lines, it became apparent that one of the lines had many of the same mutations as another line, likely due to a mislabeling or contamination at some point of the MA experiment. This line was excluded from the analysis, leaving 39 MA lines in the data.

Accurate mutation calls are crucial for estimating mutation rates. The majority of our mutation calls had a maximum genotype quality score of 99 and were unambiguous (Figure S1). However, to ensure the accuracy of our mutation calls, we verified a sample of mutations by Sanger-sequencing. We had two verification sets: a set of mutation calls that were of lower quality based on a visual inspection of alignments, and a second set of randomly selected mutations across different genomic domains. We chose the mutations from the first set because if those calls were correct, then mutations with higher-quality scores are likely to be correct as well. We selected 29 point mutations to be confirmed. PCR or sequencing failed in 6 out 29 point mutations, and the remaining 23 point mutations were all confirmed. For the 37 small indels, PCR or sequencing failed in 10 of them, while 20 were confirmed, and 7 were false positives. Of the 16 SVs tested, PCR or sequencing failed in 9, 5 were confirmed, and 2 were false positives. The randomly selected mutations of the second set were selected to understand if heterochromatin (H3K9 or centromeric domains) had higher rates of false positive mutations. We randomly selected 15 point mutations in each of the three genomic domains (H3K9me3, centromeric, and euchromatin), 45 in total. One mutation located in a centromeric region failed to amplify by PCR, and all the rest of the 44 mutations were confirmed.

For point mutations, we never observed a false positive mutation out of the total 67 mutations checked by Sanger-sequencing, these included 23 mutations in euchromatic, 25 in H3K9me3, and 19 in centromeric regions. Since mutations in the first verification set represented mutations with the worst genotype qualities, our genotyping for point mutations was very accurate (see supplementary results), although there was some uncertainty for small indels and SVs. All mutations that were false positives were excluded from the data.

To ensure that our pipeline calls mutations in an unbiased way, we simulated mutations to the *N. crassa* genome, and then simulated short reads from this genome. We used two different scenarios to explore if the repetitive nature of heterochromatic regions created any bias. In the first scenario, the mutation rate was higher in H3K9me3 domains than in the rest of the genome, and in the second scenario, the mutation rate was uniform across the entire genome. There was no difference between the ratios of H3K9 / euchromatin mutation rates whether we used mutations called from the simulated read data, or used the true number of simulated mutations (see supplementary results).

In total, we observed 1322 mutations, with a median of 33 mutations per MA line. One of the MA lines had an excessive amount of mutations (Figure 1D), and it is possible that a mutation happened in this line that increased the mutation rate. There was a trend of increased C:G → A:T transversions in this line, but the rate was not statistically different from the rest of the MA lines (Figure S2). We did not observe any obvious candidate mutation that affected a DNA repair gene.

The breakdown of different mutation types among the MA lines was 1077 single nucleotide mutations, 134 insertions, 97 deletions, 9 complex mutations where a single mutational event created multiple adjacent nucleotide changes, and 5 translocations. The total mutation rate during asexual propagation was 0.03 [0.03, 0.04] mutations / genome / mitosis.

### Mutation rate variation across the genome

To examine whether the chromatin structure influenced mutation rates, we gathered publicly available data for H3K9 trimethylation (H3K9me3), H3K27 trimethylation (H3K27me3), H3K36 dimethylation, and locations of *N. crassa* centromeres (Jamieson et al., 2013; Bicocca et al., 2018; Smith et al., 2011). We also examined regions of the genome containing ancestral duplicates defined by Wang et al. (2020), but the duplicated regions were almost perfectly correlated with H3K9me3 domains (Figure 2A), so we did not use them in further analyses. Furthermore, H3K9me3 and H3K36me2 domains were nearly perfect mirror images of each other (Figure 2A). H3K36me2 domains were thus excluded due to containing the same information as the absence of H3K9me3.

**Figure 2:**
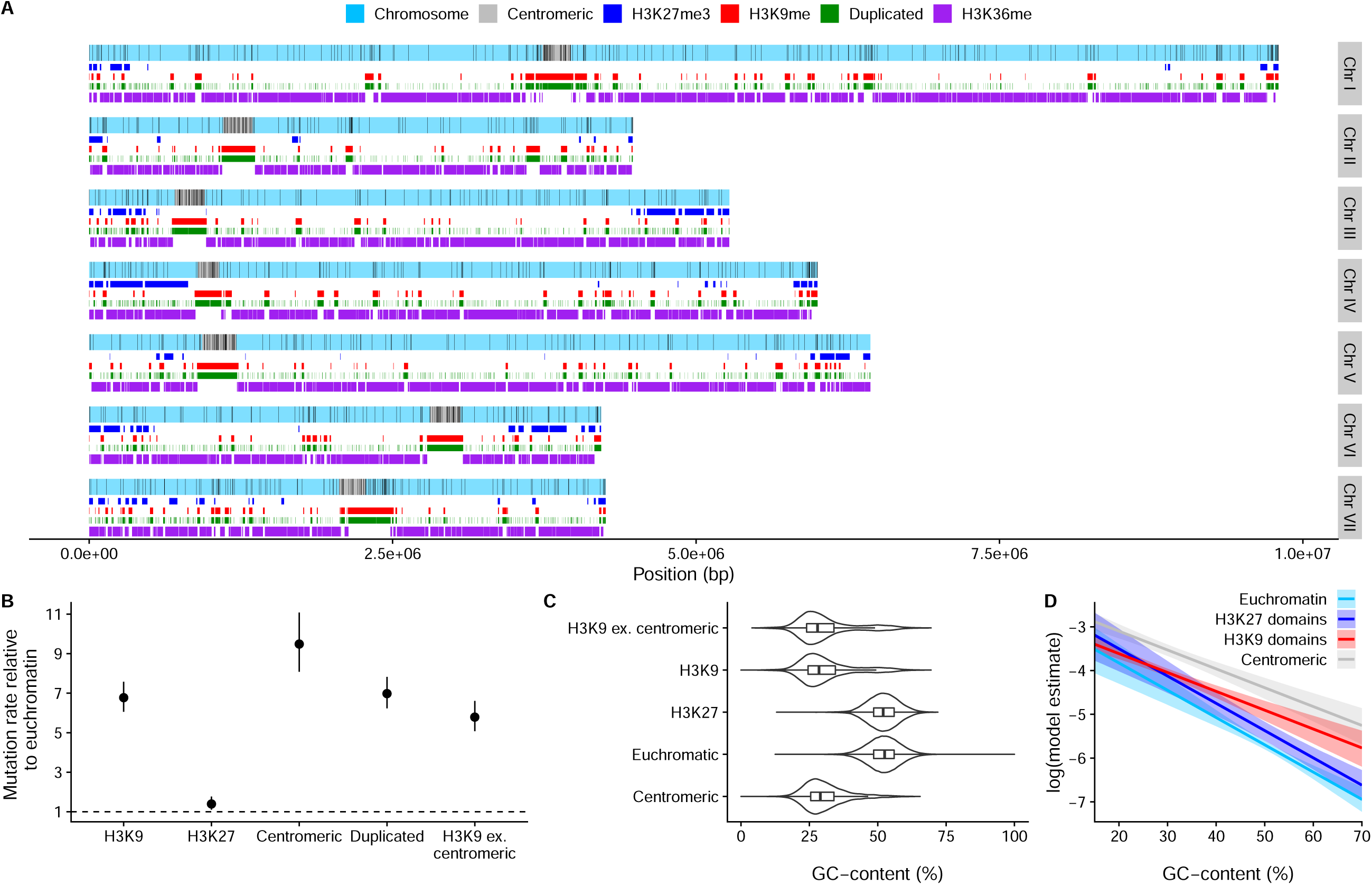
A) Distribution of mutations along the seven chromosomes. Black vertical lines indicate mutations. Centromeric regions, H3K27 trimethylation, H3K36 dimethylation, H3K9 trimethylation domains, and duplicated regions are shown. B) Relative mutation rates for different genomic domains. H3K9 ex. centromeric are H3K9me3 domains where overlaps with centromeric regions have been excluded. Posterior medians and 95% HPD intervals are shown. C) Violin plots overlaid with boxplots for GC-content in different domains. D) Model estimates (on a log-scale) for the mutation rate from a model with GC-content, H3K9me3, H3K27me3, and centromeric domains as predictors. Lines are posterior medians and envelopes are 95% HPD intervals.

Next, we examined the distribution of mutations across the seven *N. crassa* chromosomes. We observed that mutations were not uniformly distributed along the chromosomes, but were concentrated in centromeric regions and regions of the genome marked by H3K9me3 (Figure 2A). Examining relative mutation rates confirmed that the mutation rate was over sixfold higher in centromeric and H3K9me3 domains (Figure 2B), while the effect of H3K27me3 domains was much smaller: the mutation rate in H3K27me3 domains relative to euchromatin was only 1.4 [1.1, 1.78] times higher.

We also observed that GC-content displayed a bimodal distribution and was lower in H3K9me3 and centromeric domains, barely overlapping with the distribution of GC-content in euchromatin (Figure 2C). To clarify does the higher mutation rate in H3K9me3 domains arise from the GC-content itself, some other factor related to the chromatin modifications, or a combination of both, we examined the effect of GC-content on the mutation rate within the different domains. We observed that lower GC-content was associated with higher mutation rates within each domain, at different ranges of GC-content (Figure S3). However, the pattern was unclear in H3K27me3 domains, as few mutations were observed in H3K27me3 regions with low GC-content.

To investigate the joint effects of GC-content and chromatin modifications, we fitted models with different predictors, including GC-content, H3K9me3, H3K27me3, and centromeric domains. Based on model comparisons, the model with the best predictions included the effect of GC-content, the effects of H3K9me3, H3K27me3, and centromeric domains, and the interaction between the GC-content and H3K9me3 domain (Table S2). Based on model weights, the next best model that included additional interaction between the H3K27 domain and GC-content was also plausible (Table S2). However, the overall predictions for these two models were similar. There were only a few mutations in low GC areas of H3K27me domains, creating uncertainty in estimating a different slope for H3K27me3 domains; therefore, we prefer the first model with the highest weight. GC-content had a strong effect on the mutation rate, with areas of low GC having higher mutation rates (Figure 2D). Within H3K9me3 domains, GC-content had a smaller effect on the mutation rate, and centromeric regions had a statistically detectable increase in the mutation rate in addition to the effect of the H3K9me3 domain (Figure 2D, Table S3), even if centromeric regions always have H3K9me3. H3K27me3 also increased the mutation rate on top of the GC-effect (Table S3).

### Genetic variation in natural populations and mutation rate

To investigate the amount of genetic variation across the genome, we calculated nucleotide polymorphism, *θ_W_*, which measures how many polymorphic bases are found in a given length of sequence corrected for the sample size, across the genomes of natural strains in 200 bp windows. We observed that mean *θ_W_* was higher in the other domains compared to euchromatin (Figure 3A). The median estimate of *θ_W_* was 0.0150 [0.0149, 0.0151] in euchromatin, the difference to centromeric regions was 0.0213 [0.0210, 0.0216] units, the difference to H3K9me3 domains was 0.0159 [0.0156, 0.0161] units, and the difference to H3K27me3 domains was 0.0117 [0.0115, 0.0119] units.

**Figure 3:**
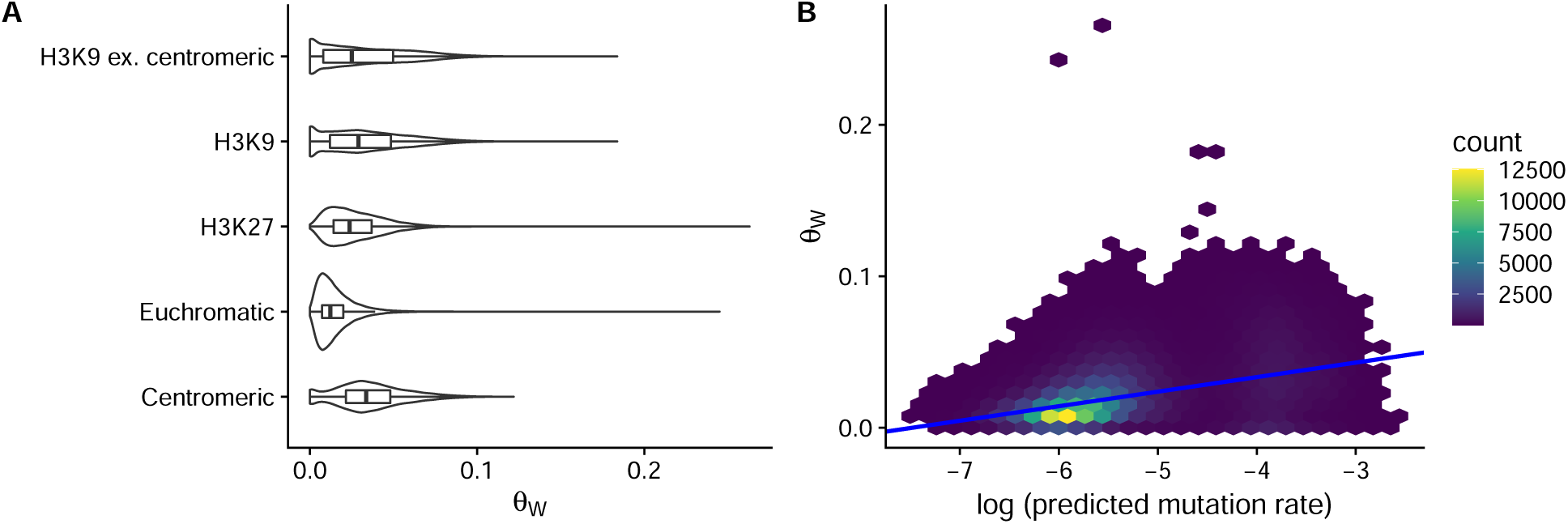
Nucleotide polymorphism, *θ_W_*, was calculated across the genome in 200 bp windows, *n* = 202310. A) Violin plots overlaid with boxplots for the distribution of *θ_W_* in different domains. B) Relationship with *θ_W_* and predicted mutation rate. For plotting, data was binned into hexes because of the high number of overlapping points, and the number of windows falling into each hex is shown by the legend.

To cross-validate our mutation model results and to investigate the role of mutation in the maintenance of genetic variation across the genome, we used our mutation model (Table S3) to predict variation in *θ_W_* across the genome. We calculated the predicted mutation rate for each 200 bp window across the genome, and observed that a simple linear model predicted a moderate amount of variation in *θ_W_* (Figure 3B). The slope of the regression line was 0.0096 [0.0095, 0.0097], so a tenfold increase in the predicted mutation rate meant an increase of 0.0096 in *θ_W_*. A measure of the model fit, the Bayesian *R*^2^ value was 0.22 [0.21, 0.22]. Although this may seem like a low *R*^2^, one should take into account that this is after our mutation model has been challenged with completely new data, and other evolutionary mechanisms besides mutation also influence *θ_W_*. The choice of the window size was not important: we tested different window sizes and found the same relationship between the predicted mutation rate and *θ_W_* (Figure S4). Larger windows even improved the fit, as there were less windows where *θ_W_* = 0; thus, our choice of a 200 bp window was conservative. We also checked that our results were robust to the windows where *θ_W_* = 0 by fitting different models that specifically modeled observations with 0 (Figure S5). We obtained the same relationship between *θ_W_* and the predicted mutation rate with these models. We further checked that the action of RIP was not solely responsible for this relationship by looking within the different domains (Figure S6 and supplementary results). The predicted mutation rate had a positive relationship with *θ* within the different domains; therefore, the action of RIP cannot solely explain these results. Consequently, the mutation rate has a substantial influence on the amount of genetic variation that is present across the genome in *N. crassa*.

### Rate and spectrum of single nucleotide mutations

Next, we examined the rate and spectrum of different types of mutations. The rate of single nucleotide mutations (SNM) across the whole genome was 6.7 [6.32, 7.11] ×10^−10^ mutations / bp / mitosis. The SNM rate in euchromatic regions was 2.46 [2.19, 2.77] ×10^−10^ mutations / bp / mitosis, and the SNM rate in H3K9me3 domains was 2.43 [2.25, 2.62] ×10^−9^ mutations / bp / mitosis. The ratio of transition to transversion (Ts/Tv) rates over the whole genome was 1.08 [0.96, 1.21], which is on the low end of reported Ts/Tv ratios. The Ts/Tv ratio of euchromatic regions was 1.49 [1.17, 1.91], which was higher than the Ts/Tv ratio in H3K9me domains 0.93 [0.8, 1.08].

As seen from the different transition to transversion ratios, the spectra of single nucleotide mutations was different for H3K9me3 domains versus the rest of the genome (Figure 4A). We calculated ratios of the relative mutation rates in different domains by taking different nucleotide or trinucleotide frequencies (see below) into account in H3K9me3 domains and euchromatin. C:G → G:C transversions were more common in H3K9 domains (Figure 4B). There was also weak evidence that A:T → C:G and A:T → G:C mutations could have different rates in the different domains. Their ratios were barely different from one when nucleotide frequencies were taken into account, but the ratios barely overlapped with one when mutation rates were corrected for trinucleotide frequencies (Figure 4B). However, A:T → T:A transversions had a lower rate in H3K9me3 domains after correcting for trinucleotide frequencies. Over the whole genome, the four different transversions occur at similar rates (Figure 4A), and the 2 transitions at higher rates. The A:T → G:C transition was the most common SNM, and more common than the C:G → T:A transition. The ratio of A:T → G:C to C:G → T:A transitions was 1.23 [1.04, 1.44].

**Figure 4:**
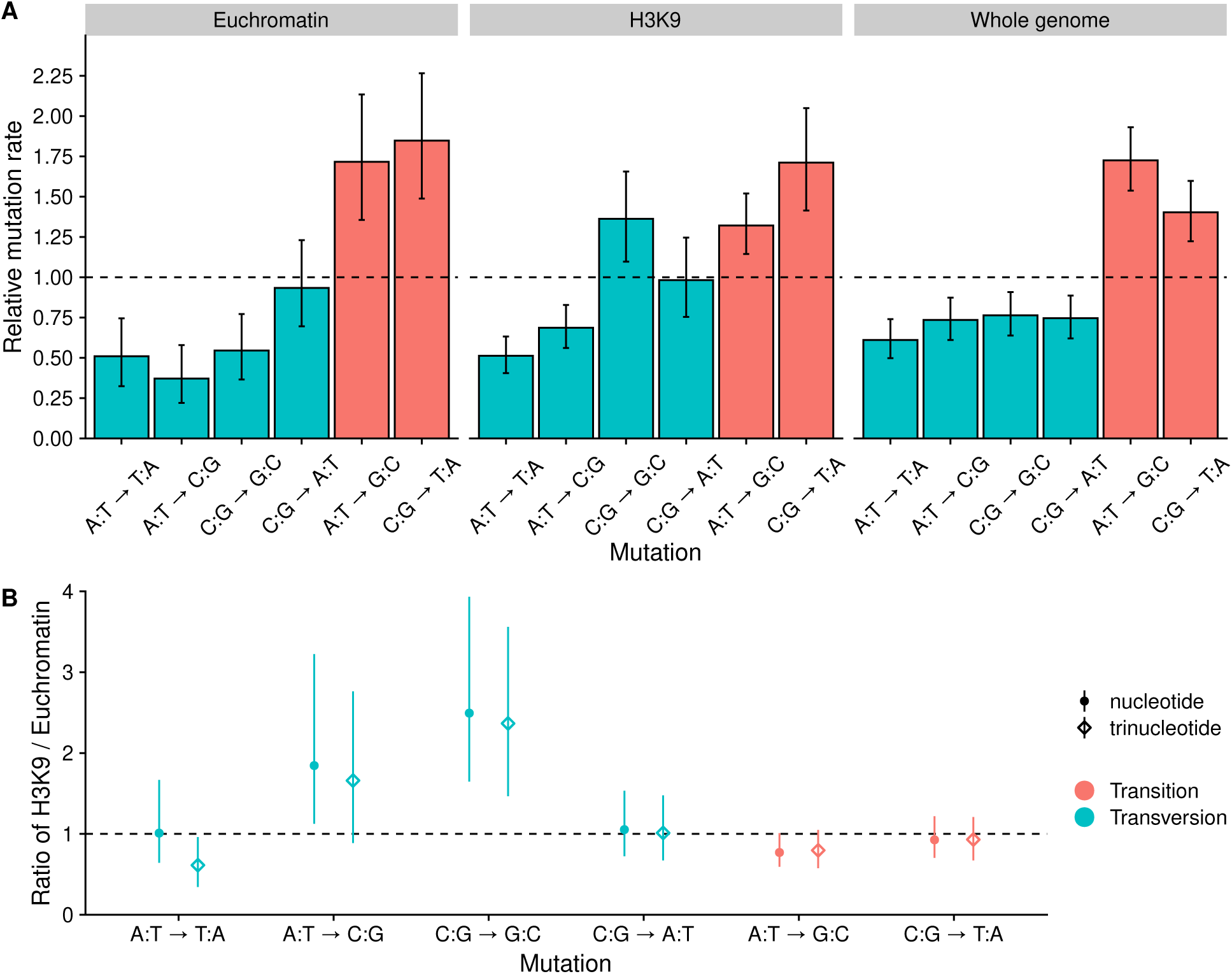
A) Spectrum of relative SNM rates, the dashed line shows the expected rate if all mutations occurred at equal frequencies. Nucleotide frequencies were taken into account in calculating the relative rates. Error bars are 95 % HPD intervals. B) Ratios of relative mutation rates in H3K9me3 over euchromatin. Points show ratios corrected for nucleotide frequencies, and diamonds show ratios corrected for trinucleotide frequenices. Estimates are medians, and the range shows 95% HPD interval of the ratio. If the interval estimate is different from one, the mutation rate is different in H3K9me3 domains and euchromatin.

### Effects of local base composition

To better understand factors influencing SNM rates, we looked at the effect of local base pair context. For each SNM, we extracted the two adjacent base pairs to get the trinucleotide context. We combined trinucleotides with respect to sequence complementarity as the strand in which the mutation occurred is unknown, this leaves 32 trinucleotide classes. We calculated trinucleotide frequencies and observed that over the whole genome, the observed frequencies were approximately at expected frequencies based on GC-content (Figure S7). However, in regions marked by H3K9 trimethylation, there were strong departures from the expected trinucleotide frequencies (Figure S7). Prompted by this observation, we investigated whether differences in the trinucleotide mutation rates could explain the observed differences in mutation rates across the genome. We compared different models with the trinucleotide classes and the effects of epigenetic domains. The model that gave the best predictions included an effect of the trinucleotide classes, effect of H3K9me3, centromeric regions, H3K27me3 regions, but no interactions between the trinucleotide class and any of the epigenetic domains (Table S4). We observed the same results regarding the epigenetic domains as before: the mutation rate was 8.1 [6.8, 9.5] times higher in H3K9me3 domains, centromeric regions had an additional increase on top of H3K9me3, and there was a small, 1.5 [1.1, 2.0] fold, increase in the mutation rate in H3K27me3 domains (Figure S8). Thus, differences in trinucleotide composition were not driving the mutation rate differences in the different domains.

After taking trinucleotide frequencies, and the effects of the epigenetic domains into account, mutations were not equally distributed across the different trinucleotide classes (Figure 5A). The trinucleotide class GAT:ATC had the lowest relative mutation rate, while TCT:AGA had the highest. Trinucleotides with adjacent C:G pairs tend to have high relative mutation rates.

**Figure 5:**
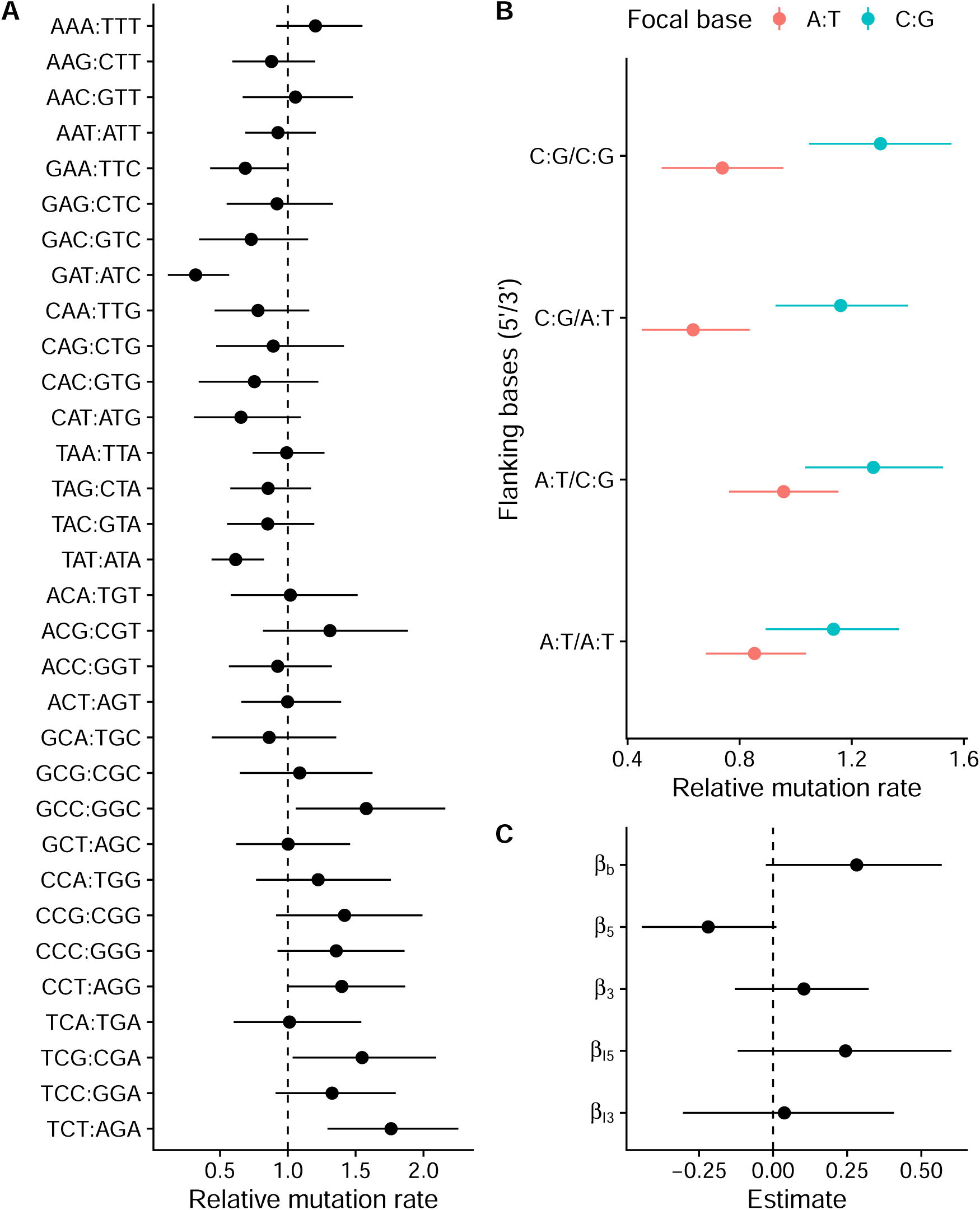
A) Relative mutation rates for the 32 different classes of trinucleotides. B) Model predictions for effects of flanking bases. C) Estimates of model coefficients for effects of flanking bases. *β_b_* is the effect of focal C:G relative to A:T, *β*_5_ is the effect of flanking 5’ C:G relative to A:T, *β*_3_ is the effect of flanking 3’ C:G relative to A:T, *B_I_*_5_ is the interaction effect of 5’ C:G when the focal base is C:G, and *B_I_*_3_ is the interaction effect of 3’ C:G when the focal base is C:G. Range shows a 95% HPD interval of the relative mutation rate.

To investigate the effects of C:G and A:T base pairs in either 5’ or 3’ flanking positions, we fitted a linear model that included the effects of the flanking bases and the mutating base pair, and we also incorporated the uncertainty in the relative mutation rate for each trinucleotide class. The model predictions are shown in Figure 5B, and there was a tendency for trinucleotides with C:G as the mutating base to have a higher relative mutation rate compared to A:T trinucleotides. However, this effect was not significant (Figure 5C). Similarly, for A:T trinucleotides, there was a tendency for 5’ C:G to protect against mutation, but this effect was not significant as the 95% HPD interval barely includes zero (Figure 5C). When the mutating base pair was C:G, neither 5’ or 3’ base had any detectable effect (Figure 5C).

### Deletions, insertions, and translocations

We also examined the structural variants that occurred in the MA lines. We observed 97 deletions and 134 insertions. Frequent lengths for deletions and insertions were changes of one base pair, 88 out of 96 1 bp indels occurred in homopolymer stretches. The second most common length was 3 bp, which were predominantly changes in microsatellite repeats. Some large deletions were observed: the largest deletion was 17.8 kb, there were 3 deletions around 8.8 kb, and one around 2.5 kb. Otherwise, most deletions were under 100 bp (Figure 6A). The largest observed insertion was 130 bp, with most insertions under 20 bp (Figure 6B).

**Figure 6:**
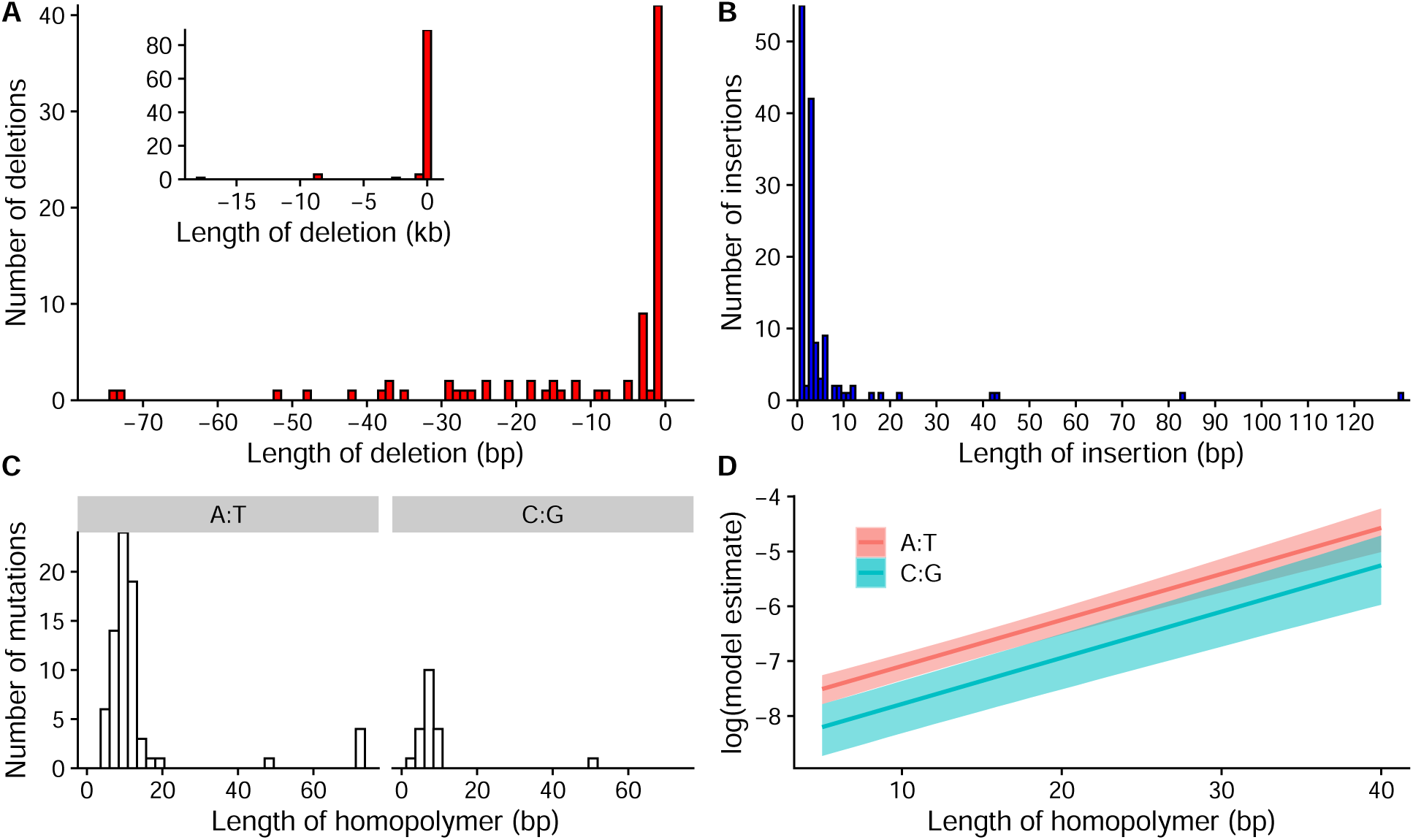
A) Distribution of deletion lengths in the range of 0 to 75 bp, inset shows the full distribution. B) Distribution of insertion lengths. C) Distribution of homopolymer lengths for those mutations that occurred in either A:T or C:G homopolymer stretches. D) Model estimates for mutation rate in homopolymers, A:T homopolymers had an overall higher mutation rate, and longer homopolymers had higher mutation rates.

Mutation rates for insertions and deletions are shown in Table 1. When all deletions and insertions were included in the analysis, insertions had slightly higher rates than deletions (Table 1). However, when we excluded homopolymers, microsatellites, and other repeats from the analysis, we observed that the rate of deletions was three times higher than insertions (Table 1). Given that the mean length of deletions excluding repeats was 1160 bp, which was longer than the 27 bp mean length for insertions, there was mutation pressure to lose DNA. Some of the large deletions observed in our data were outliers, but deletions tended to be longer at all scales (Figure 6). For mutations that occurred in repeats, the mutation rate of insertions was over twofold higher than deletions.

**Table 1:** Mutation rates (mutations / genome / mitosis) for deletions and insertions and their ratio. Deletions Insertions Insertion / deletion ratio

|  | Deletions | Insertions | Insertion / deletion ratio |
| --- | --- | --- | --- |
| All indels | 0.0024 [0.0020, 0.0029] | 0.0034 [0.0028, 0.0039] | 1.38 [1.06, 1.77] |
| Repeats excluded | 0.0011 [0.0008, 0.0014] | 0.0003 [0.0002, 0.0005] | 0.3 [0.14, 0.51] |
| Repeats only | 0.0014 [0.0010, 0.0017] | 0.0030 [0.0025, 0.0036] | 2.25 [1.6, 3.05] |

Homopolymer stretches had particularly high rates of indel mutations; we observed 92 mutations in homopolymers, and most mutations in homopolymers were indels of 1 bp. More mutations occurred in A:T than in C:G homopolymers (Figure 6C). A:T homopolymer loci are approximately 1.7 times more common in the genome, but even when adjusting for frequencies, the mutation rate in A:T homopolymers was 1.79 [1.4, 2.22] ×10^−8^ mutations / locus / mitosis compared to the rate in C:G homopolymers of 8.15 [4.89, 11.86] ×10^−9^ mutations / locus / mitosis. Thus, mutations in A:T homopolymers were 2.2 [1.27, 3.49] times more common than in C:G polymers. We also observed that longer homopolymers had higher mutation rates. In a model with a polymer length and polymer type, the length had the same effect for both A:T and C:G homopolymers (Figure 6D). This suggests that replication slippage, which is the mechanism suggested to be involved in indel mutations in repeats, tends to occur more often in longer repeats as expected.

There were differences in indel rates in the different genomic regions. We observed that deletions had a higher mutation rate in centromeric regions and in regions marked by H3K9me3 (Figure S9) compared to euchromatin, even when only deletions in repeats were considered. For deletions excluding repeats, H3K9me3 and H3K27me3 domains had a higher mutation rate than euchromatin (Figure S9). For insertions, we did not observe any differences in the mutation rate in different domains (Figure S9).

We observed 5 translocations in the MA lines, two of which were among the SVs confirmed by PCR and Sanger sequencing. Three translocations were from one chromosome to another, and two occurred among unmapped contigs. The mean translocation length was 316 bp. The translocation rate was 1.19 [0.33, 2.41] ×10^−4^ translocations / genome / mitosis. Due to their rarity, we don’t have enough translocations to further analyze their properties.

### Complex mutations

We observed 9 cases where two SNMs or 1 bp indels occurred within few base pairs of each other in the same MA line. While it is possible that two independent mutations occurred next to each other, it is unlikely. These changes were more likely caused by a single mutational event. The observed complex mutational events are listed in Table 2. To confirm whether these changes were caused by a single mutational event or two independent events, we used Sanger sequencing to check the genotypes of MA lines from intermediate transfers stored during the experiment. We always observed that the two changes appeared together (Table 2). The most parsimonious explanation is that these changes appeared as a result of single mutational events, likely caused by a DNA repair error via an error-prone DNA-polymerase.

**Table 2:**
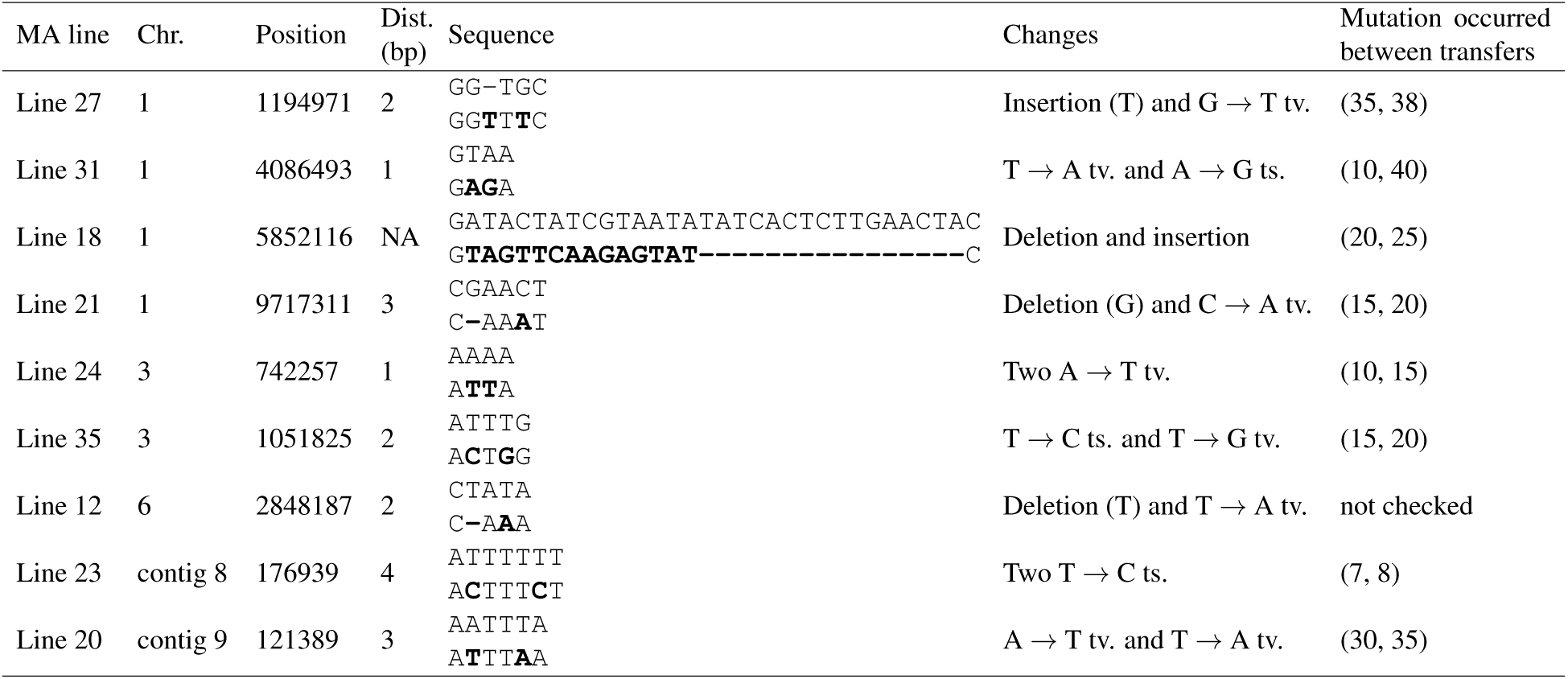
Complex mutations. Chr. is the chromosome or contig, position is the coordinate of the most 5’ end change, and dist. is the distance between nucleotide changes. Sequence column shows the ancestor sequence on the first row and the MA line sequence on the second row, and mutated bases are in bold. Final column contains the MA experiment transfer numbers (mutations absent, mutations present) of those transfers when the last transfer occurred, where none of the changes could be observed in the MA line, and the transfer when all changes were observed in the MA line. We always observed all changes in complex mutations to be present together. tv. = transversion, ts. = transition

We treated complex mutations as single events in calculations where all mutations were used to calculate overall mutation rates. The rate of complex mutations was 5.42 [2.22, 9.3] ×10^−12^ mutations / bp / mitosis. The rate of SNMs over the rate of complex mutations was 123.7 [57.48, 231.49], making point mutations over 100-fold more common than complex events.

### Comparison of mutation rate and spectra during meiosis and mitosis

Wang et al. (2020) observed an extremely high mutation rate during meiosis, as a result of C:G → T:A transitions induced by RIP. We re-analyzed their data in combination with the chromatin modification data (see supplementary results for details). We observed heterogeneity in the activation of RIP that was not taken into account in the original analysis (see Figure S10 and supplementary results). Based on our analysis, the mutation rate during meiosis in euchromatin was 1.07 [0.60, 1.67] ×10^−8^ mutations / meiosis / bp, and 2.54 [0.11, 7.55] ×10^−7^ in H3K9me3 domains. The mutation rate was substantially higher during meiosis than mitosis, as observed by Wang et al. (2020), but not as high as their analysis suggested. Comparing mutation spectra in mitosis and meiosis shows that in euchromatic regions, A:T → C:G, and C:G → G:C transversions were more common in meiosis, while C:G → A:T transversions were less common (Figure S11). In heterochromatin, C:G → T:A transitions overwhelmed all other mutations in meiosis (Figure S10, S11).

## Discussion

We have generated a highly precise estimate of spontaneous mutation rate during asexual growth in *Neurospora crassa*. Our estimate of the point mutation rate across the whole genome of 6.7 [6.32, 7.11] ×10^−10^ mutations / bp / mitosis is higher, although it is close to an estimate of asexual mutation rate of 6.03 × 10^−10^ obtained by Wang et al. (2020) with only 64 observed mutations. A previous estimate from marker gene studies suggested that the mutation rate is 4 10–4.66 × 10^−9^ (Lynch et al., 2016), but neither our, nor the results of Wang et al. (2020) agree with this. The point mutation rate in euchromatic regions was 0.007 [0.006, 0.008] mutations / genome / mitosis, which is in line with results obtained by Drake (1991), who observed that the mutation rate per genome for microbes seems to be around 0.003 mutations per genome per generation with approximately twofold variation around this mean. Thus, the asexual mutation rate in *N. crassa* in euchromatic regions seems to be rather typical for a microbe.

*N. crassa* has a striking difference in the rate and spectrum of mutation during sexual and asexual reproduction (Wang et al., 2020). During sexual reproduction, a genome defence mechanism called RIP is activated, which recognizes duplicated regions in premeiotic cells and induces C:G → T:A transitions in those regions (Selker, 1990). RIP presumably protects the genome against transposons and other selfish genetic elements. Mutations induced during sexual reproduction happen mainly in ancestral duplications (Wang et al., 2020), and these regions nearly completely overlap with regions of H3K9 trimethylation. In contrast, during asexual reproduction, while H3K9me3 domains have a higher mutation rate than euchromatin, the difference is much smaller and the spectrum of mutations is different compared to meiosis. For euchromatic regions, our analysis supports a higher mutation rate during meiosis than in mitosis, although not as high as suggested by Wang et al. (2020). We also observed that the spectrum of mutations was different in meiosis, but notably, there was no difference in C:G → T:A transitions in euchromatin. Moreover, since gene density is much higher in euchromatic regions, the action of RIP likely does not result in as high genetic load as suggested by Wang et al. (2020).

While many studies have reported effects of chromatin structure on mutation rates, these studies have often been based on indirect inference from species divergence and polymorphism (Sasaki et al., 2009; Ying et al., 2010; Washietl et al., 2008). We observed directly extensive variation across the genome in the mutation rate and mutation spectra due to chromatin modifications. In *N. crassa* H3K9 trimethylation determines heterochromatic regions, and H3K27 trimethylation is a mark for facultative heterochromatin (Jamieson et al., 2013). Centromeric regions determined by the presence of centromeric histone variant CenH3 always overlap with H3K9 trimethylation (Smith et al., 2011). Methylation in H3K9 and H3K36 are almost completely mutually exclusive. However, H3K36me2 is not a straightforward mark of euchromatin as it can be deposited by two enzymes: SET-2 and ASH1. Genes marked with H3K36 methylation by SET-2 are actively transcribed, while genes marked with H3K36 methylation by ASH1 are silenced and can be further marked by H3K27 trimethylation (Bicocca et al., 2018). SET-2 is responsible for most of H3K36 methylation, and we considered all regions that lacked H3K9me3 and H3K27me3 to be euchromatin. The mutation rate for SNMs in domains marked by H3K9me3 was tenfold higher than in euchromatic regions. We also observed that the mutation rate was slightly elevated in H3K27me3 domains, although this effect was much smaller than for H3K9me3. In centromeric regions, there was an additional effect of an increased mutation rate on top of the H3K9me3 effect.

An increased mutation rate in heterochromatic and centromeric regions has also been found by Weng et al. (2019) in the plant *Arabidopsis thaliana*. These results are also in line with observations from human cancer cells that have higher rates of mutation in heterochromatic regions (Schuster-Böckler and Lehner, 2012; Polak et al., 2015). Monroe et al. (2022) observed that in *Arabidopsis*, the mutation rate was affected by several different epigenetic marks; they suggested that the mutation rate was lower in genes that were actively transcribed, and perhaps even fine-tuned for highly expressed genes, possibly due to the presence of H3K36 methylation or other epigenetic marks of active transcription. However, our dataset does not contain enough mutations to address if gene expression levels quantitatively influence the mutation rate in *N. crassa*.

What is the mechanism causing mutation rate variation across the genome? Chromatin structure is involved in DNA repair, and this may explain why heterochromatic regions have a higher mutation rate. In yeast, actively transcribed regions contain acetylation at H3K56, which suppresses spontaneous mutations (Kadyrova et al., 2013). Moreover, analyses based on human tumors suggest that DNA mismatch repair works more efficiently in euchromatin than in heterochromatin, and it is DNA repair that is variable, not the supply of mutations themselves (Supek and Lehner, 2015). Further evidence supporting the DNA accessibility hypothesis was gathered by a study performed by Yazdi et al. (2015), the authors observed that mutation rate did not increase in high nucleosome occupancy regions in strains where DNA repair machinery was knocked out, in contrast to wild-type strains. Recently, Habig et al. (2021) investigated how epigenetic modifications affect the mutation rate in the fungus *Zymoseptoria tritici*. They observed that H3K27 trimethylation increased the mutation rate as it did in this study. Furthermore, Habig et al. (2021) observed that this effect disappeared from those regions in a mutant lacking H3K27me3. This suggested that H3K27me3 is causal, and DNA accessibility may be the reason. Habig et al. (2021) did not detect an effect for H3K9me3 like we observed, which may indicate that there are species specific differences in heterochromatin.

Differential exposure of heterochromatin to natural mutagens, such as oxidative damage, does not seem to explain our results. As mutations typically associated with oxidative damage: G → C transversions, C → T transitions, and G → T transversions (McBride et al., 1991; Cheng et al., 1992), were not systematically overrepresented in regions with H3K9 trimethylation, only the relative amount of G → C transversions differed between euchromatic and H3K9 trimethylated domains.

Among the observed mutations, sequence context had an effect on the mutation rate. It is known that local sequence context can influence the probability of mutation, and this phenomenon has been frequently observed (Ness et al., 2015). However, the mechanisms of why some sequence contexts are prone to mutation are less clear. We observed a tendency of 5’ C:G base pair to protect against mutation when the mutating base pair is A:T. A similar phenomenon has been observed before, as formation of thymine-thymine cyclobutane dimers due to DNA damage caused by sunlight is dependent on flanking sequence, even if the biophysical basis is unclear (Law et al., 2013). We also observed a tendency for trinucleotides with C:G as the mutating base pair to have higher relative mutation rates, possibly due to trinucleotides with GG dimers being susceptible to oxidation (Hanrahan et al., 1997).

We did not find evidence that trinucleotide mutation rates were different in regions of H3K9me3. One hypothesis for different trinucleotide mutation rates in heterochromatin would be that the presence of DNA methylation increases the mutation rate in certain contexts. In *N. crassa*, DNA methylation occurs only in H3K9 trimethylated domains (Tamaru and Selker, 2001), and deamination of 5-methylcytosine is a known cause of mutations (Cooper et al., 2010). However, deamination should cause mainly C:G → T:A transitions, and we did not observe an excess of C:G → A:T transitions in H3K9me3 domains. Instead, we observed an excess of C:G → G:C transversions in H3K9me3 domains. Therefore, DNA methylation does not contribute substantially to spontaneous genetic mutations in *N. crassa*, possibly because DNA methylation is so rare in this species (Hosseini et al., 2020).

For spontaneous insertions and deletions, we observed that deletions were more common than insertions when repeated sequences were excluded. Since deletions also tended to be longer, mutations have the tendency to reduce genome size. Similar patterns in mutational bias have been observed for *Drosophila melanogaster* (Leushkin et al., 2013). For repeated sequences, we analyzed homopolymer sequences in more detail. We found that while A:T homopolymers are more common in *N. crassa*, they also have a higher mutation rate. This is in contrast to observations in nematodes, where C:G homopolymers had much higher mutation rates than A:T homopolymers (Denver et al., 2004), suggesting that there are species-specific differences in mutation rates of these sequences.

We identified mutations with multiple base changes due to a single mutational event. Such complex mutations are thought to arise from the action of error-prone translesion DNA polymerases, such as Pol *ζ* (Stone et al., 2012). A recent large-scale survey of human trios identified many multinucleotide mutations, and observed that mutations that were 2–10 bp apart showed an over-representation of A:T → T:A, and A:T → G:C mutations (Besenbacher et al., 2016). Our results are compatible with this pattern as most nucleotide changes in complex mutations were either A:T → T:A or A:T → G:C. We did not observe GC → AA or GA → TT tandem mutations, which are the most common tandem mutations in humans (Harris and Nielsen, 2014; Besenbacher et al., 2016). In *N. crassa*, complex mutations represented only 0.7% of observed mutations, which is smaller than approximately 3% observed in humans. Thus, the impact of multinucleotide mutations on population genetic inference may be small in *N. crassa*.

Finally, the implication for evolutionary genetics is that variation in mutation rates must be taken into account in order to quantify which evolutionary forces act on the genome. Genes residing in different regions of the genome can differ substantially in their mutation loads, and it seems that variation in mutation rates also has a large part in determining how much genetic variation is segregating in a given region of the genome. To accurately estimate imprints of selection on natural genetic variation, we must account for variation in mutation rates in evolutionary models. Therefore, detailed models of mutation rates are needed for different species.

## Methods

### Mutation accumulation experiment

We used two ancestors for the MA experiment: B 26708 and B 26709. These strains have been generated by backcrossing mating type *mat a* from strain 4200 into 2489 background nine times (Kronholm et. al. 2020). There were 20 MA lines for both ancestors, giving 40 lines in total. Common protocols for culturing *N. crassa* were followed (see supplementary methods). We propagated the MA lines for 40 transfers.

To estimate the number of mitoses that that happened in the MA experiment during a transfer, we used a strain that had been tagged with fluorescent protein at histone H1 (Freitag et al. 2004). Using this strain, we counted the numbers of nuclei that were present in each phase of a single transfer of the MA experiment with confocal microscopy (Figure 1C), and used this data to estimate the number of mitoses the lines went through (see supplementary methods).

### Strains from natural populations

We obtained 33 strains from the Fungal Genetics Stock Center (McCluskey et al., 2010), and re-sequenced these strains. In addition, we obtained genome sequencing data for another 23 strains from Zhao et al. (2015). In total, including the laboratory strain 2489, we had 57 strains with sequencing data (Table S5).

### DNA extraction

We extracted DNA from the strains using Phenol-Chloroform extraction, and further purified it via polyethyleneglycol precipitation. DNA quality was checked on agarose gels. For a detailed protocol, see supplementary methods.

### Genome sequencing

DNA was sequenced at Novogene (Cambridge, UK), using Illumina platform with paired-end 150 bp libraries. Libraries were prepared by fragmenting the DNA by sonication, adapter ligation, and PCR-amplification. Libraries were sequenced to 30x target coverage. Reads with adapter sequences, > 10% N’s, or > 50% bases of low-quality Q score ≤ 5 were removed.

### Read mapping and genotyping

We mapped reads against the *N. crassa* reference genome (assembly version NC12) using BWA-MEM (Li, 2013). The strain used for the original genome project was 2489, so the MA experiment ancestors should be nearly identical to the reference.

We used the GATK version 4.2.0.0 (McKenna et al., 2010) pipeline to call SNMs and indels. First, we ran Haplotypecaller for each sample, and then genotyped all samples together. *Neurospora crassa* is haploid, but we ran Haplotypecaller in diploid mode because mapping errors manifest as heterozygous sites in haploids (Kronholm et al., 2017), see also Li (2014) and Ness et al. (2012). We used wormtable version 0.1.5 (Kelleher et al., 2013) and custom Python scripts to filter for high-quality sites. To produce the final set of mutations, we checked all candidate mutations manually by inspecting the alignment in IGV (Thorvaldsdóttir et al., 2013). SNPs were genotyped using the same pipeline (see supplementary methods for details).

To genotype structural variants, we first compared the performance of different SV genotyping algorithms using simulated data. We simulated SVs by modifying the *N. crassa* genome using SURVIVOR 1.0.7 (Jeffares et al., 2017). We then simulated short reads from these genomes using DWGSIM 0.1.11 (Homer, 2021). We simulated different scenarios with different amounts, types, and lengths of SVs. Then we proceeded to call the simulated SVs with different SV callers. See supplementary methods for details. Delly 0.8.7 (Rausch et al., 2012) and Lumpy 0.2.13 (Layer et al., 2014) were the callers that performed best, and they were selected for genotyping. SVs with a genotype quality score below 30 and read depth below 10 were discarded. Alignments for each called SV were manually inspected in IGV.

### Validation of mutations with Sanger sequencing

To verify a sample of the observed mutations, we performed Sanger sequencing using standard methods. First, we selected a set of mutations that passed our threshold, but had the lowest quality scores: 30 point mutations, 37 small indels and 16 larger structural variants, 83 in total. Second, we selected another set of mutations at random: 15 point mutations from three genomic regions (H3K9, centromeric and eurchromatin), 45 in total. In total, we attempted to confirm 128 mutations, list of primers and confirmed mutations are given in the supplementary file S1. Additionally, complex mutations, where a suspected single event created multiple changes, were selected for verification, and we sequenced time points from the middle of the MA experiment to see if the multiple changed sites appeared together in the MA line.

### Chromatin modifications

We used publicly available data to determine where chromatin modifications occurred. ChIP-seq reads for H3K9 trimethylation and H3K27 trimethylation were obtained from Jamieson et al. (2013), accession numbers SRX248101 and SRX248097. Jamieson et al. (2013) used the same strain (2489) that we used as the ancestor of the MA experiment. They also tested whether different growth media had an effect on H3K27me3 regions, and observed that they were stable. Jamieson et al. (2013) extracted chromatin from mycelium, which gives rise to the asexual spores used for transfers in our experiment, so the chromatin states they observed should be relevant for our experiment. Taken together, this data provides a very good approximation of the chromatin states that the ancestors of the MA lines had in our experiment. Data for H3K36 methylation was obtained from Bicocca et al. (2018), accession number SRX4549854. Data for centromeric regions were obtained from Smith et al. (2011). See supplementary methods for details.

### Statistical analysis

We used Bayesian statistics in all statistical modeling. As a rule of thumb for comparison to frequentist methods: if a 95% interval estimate of a parameter does not contain zero, the parameter is statistically different from zero, i.e., significant. We tested differences in mutation rates by computing ratios from posterior distributions. If the interval of the ratio did not include one, the mutation rates were statistically different. Estimates were reported as medians and 95% highest posterior density intervals in square brackets.

### Mutation rate estimates

Mutation rates were estimated using Bayesian Poisson models implemented with the Stan language (Carpenter et al., 2017) interfaced from R 3.6.0 (R Core Team, 2019) with the “brms” package (Bürkner, 2017). The basic model for estimating the mutation rate was

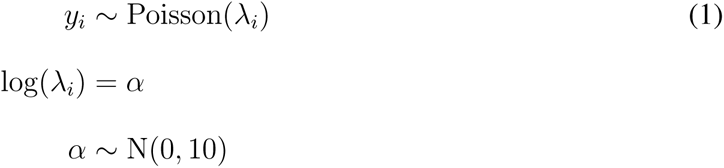

where *y_i_* is the number of mutations in *i*th MA line, *λ_i_* is the poisson rate parameter, and *α* is the intercept. The linear model part was modified accordingly if other predictors were used. We can then calculate the mutation rate, *µ*, from posterior distributions as

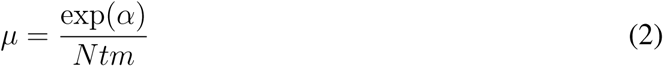

where *N* is the number of called nucleotides, *t* is the number of transfers the MA lines went through, and *m* is the number of mitoses per transfer. We used the posterior distribution for *m*, so any uncertainty in number of mitoses is incorporated into our estimated mutation rate. To get the mutation rate per genome, *N* is removed from the denominator. We used a weakly regularizing prior for *α*. Settings for MCMC estimation were: 1000 iterations of warm-up followed by 3000 iterations of sampling with four independent chains. MCMC convergence was monitored by traceplots and *R*^^^ values. No convergence problems were observed. For details of the other models, see supplementary methods.

### Mutation rate in repeats

Homopolymer sequences in the genome were detected using MISA (Beier et al., 2017). We used 5 bp as the minimum homopolymer length, and extracted repeat counts for all homopolymer loci in the genome. Counts of homopolymers of different lengths were used as an offset term in a model used to estimate mutation rates in homopolymers.

### Population genetics

We estimated nucleotide polymorphism, *θ_W_*, from the SNP data for the population sample following Ferretti et al. (2012), which allows us to deal with missing data.

### Data access

The genome sequencing data generated in this study have been submitted to the NCBI BioProject database (https://www.ncbi.nlm.nih.gov/bioproject/) under accession number PRJNA839531. Other data and scripts are available at https://github.com/ikron/mutation_ms and as supplemental material.

## Supporting information

Supplementary Datafile S1

Supplementary Datafile S2

## Acknowledgments

This study was supported by a grant from the Academy of Finland (no. 321584) to IK. We thank Dr. Visa Ruokolainen for help with confocal microscopy, the Finnish CSC-IT Center for Science Ltd. for providing computational resources, and Dr. Matthieu Bruneaux for comments on the manuscript.

## Author contributions

I.K. conceived the study. P.A.M.S., M.V., M.L., and I.K performed experiments. I.K., M.V., and P.A.M.S. analyzed the data. I.K. and M.V wrote the manuscript. All authors edited the final manuscript.

## Supplementary Information

### Supplementary methods

#### Mutation accumulation experiment

We started the MA experiment with two different strains: 2489 *mat A* and 2489 *mat a*. We have previously generated these strains by backcrossing mating type *mat a* from strain 4200 into 2489 nine times (Kronholm et al., 2020). The strains differ in their mating types, but should share over 98% of the rest of their genetic background. Their Fungal Genetics Stock Center ID’s are: B 26708 and B 26709. We used 20 lines for both these strains, giving 40 MA lines in total. We used two different mating types to later have the possibility to perform crosses between the MA lines. However, for this study the mating types of the lines do not matter as all propagation was asexual.

Common protocols for culturing *N. crassa* were followed, and sorbose plates were used to induce colonial morphology on plates (Davis and de Serres, 1970). The experiment was started by picking a single colony from a sorbose plate for both ancestors and transferring that colony into a 75×12 mm test tube with flat surface of 1 mL of Vogel’s Medium (VM) with 1.5% agar and 1.5% sucrose (Metzenberg, 2003). Tubes were incubated at 25 °C for 3 days to allow conidia (asexual spores) to develop. Then we picked small amount of conidia with a loop into a tube with 1.4 mL of 0.01% Tween-80, we then pipetted 1 µL of this conidial suspension into a 50 µL water droplet on a sorbose plate and spread it. We incubated the plates at room temperature for 2 days and picked single colonies to establish the MA lines. The MA lines were transferred the same way, so that a single colony was always picked randomly from a sorbose plate to propagate the MA line (Figure 1B). We tested that 2 days of incubation was enough time for all colonies to appear on plates. Combining the time of 2 days on plates and 3 days in a tube, a single transfer took 5 days. We propagated the MA lines for 40 transfers, the ancestors and the MA lines were stored frozen in suspended animation until sequencing.

#### Estimating the number of mitoses in the MA experiment

To estimate the mutation rate per mitosis, we needed to estimate how many mitoses happened in the MA lines during the experiment. To estimate the number of mitoses that happened during one transfer, we needed to obtain data about the number of nuclei present in each phase of a transfer: in a colony on a sorbose plate, in the mycelium in a test tube, and the conidia produced in the test tube. To estimate the density of nuclei per µm^2^ of hyphae we used the strain *mat A his-3*^+^*::Pccg-1-hH1*^+^*-sgfp*^+^ (FGSC# 9518) which expressed a green fluorescent protein that had been fused into histone H1 (Freitag et al., 2004). We grew the strain on plates with either normal VM medium or sorbose medium, cut out a piece of the agar, and mounted it on a glass coverslip using the inverted agar block method (Lichius and Zeilinger, 2019). We used Congo Red to stain cell walls: a 20 µL droplet with 2 µM Congo Red was pipetted to a glass coverslip and an agar block with the side carrying the mycelium was placed face down in the droplet.

Samples were imaged with a Nikon A1R confocal microscope, GFP was excited with a 488 nm laser and detected with a 515/30 emission filter, Congo Red was excited with a 561 nm laser and detected with a 595/50 emission filter. Plan apochromat air objectives 20x (numerical aperture 0.75) and 40x (numerical aperture 0.95) were used. Laser power was set as low as possible to avoid saturated pixels. We imaged vertical stacks of the mycelium, and used imageJ2 (Rueden et al., 2017) to measure the area covered by hyphae in sections of the image, and counted the number of nuclei in these areas (Figure 1C).

We then estimated the number of nuclei in the different phases of a transfer, and calculated the number of mitoses that the MA lines went through. The number of nuclei can only increase when the old nuclei divide. If we know the number of initial nuclei and the number of nuclei at time *t*, we can calculate the number of mitoses, *m*, that separate these time points from the equation

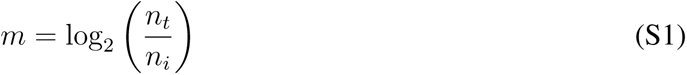

where *n_i_* is the initial number of nuclei, and *n_t_* the number of nuclei at time *t*. Thus, in order to estimate number of mitoses that happened in the MA lines during one transfer, we need to count how many nuclei were present in the colonies on sorbose plates that were picked and transferred to slants, and how many nuclei were in the mycelium that formed in the test tube, and finally how many nuclei were in the conidia that formed in the test tube (Figure 1B). This allows us to calculate how many mitoses happened during one transfer of the MA experiment, from one spore to a spore. There are multiple sources of uncertainty in these calculations, so we used a Bayesian framework to do the calculations using posterior distributions of the estimates to incorporate all sources of uncertainty in the final estimate.

Nuclei were counted from the microscope images using Fiji2 version 2.0.0-rc-54/1.51g (Schindelin et al., 2012). Short sections of mycelium were surrounded with the rectangular selection tool and the area inside was measured. All nuclei with more than 50% of their diameter inside the selection were counted manually. Multiple sections were counted from each image, with no overlap. In some fainter images, the contrast was enhanced with the enhance contrast tool, with the default value 0.3% saturated pixels and no histogram equalization. To estimate the number of nuclei in a given area of hyphae, we used the counts of nuclei and the hyphal areas measured from the microscope images to obtain the number of nuclei per µm^2^. We had images for both VM and sorbose plates, in total we collected 519 measurements. To estimate average density of nuclei for VM and sorbose we used a model where we allowed standard deviations to differ for VM and sorbose media:

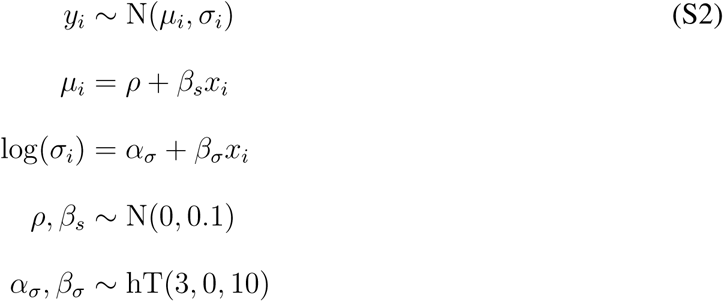

where *y_i_* is the *i*th density measurement, *ρ* is the intercept, *β_s_* is the effect of sorbose medium, *x_i_* an indicator variable for sorbose, *α_σ_* is the intercept for standard deviation, and *β_σ_* is the effect of sorbose medium on standard deviation. The average density of nuclei in VM medium is *ρ* and the density of nuclei in sorbose is obtained as *ρ_s_* = *ρ* + *β_s_*.

To estimate the average size of colonies on sorbose plates, we plated conidia on sorbose plates as in the MA experiment and photographed the plates. Millimeter paper was used as a scale. Colony area was measured from these images with ImageJ2 version 2.0.0-rc-43/1.50e. The pixels per millimeter calibration value was set by measuring the number of pixels per 1 mm of millimeter paper. The images were enhanced with the sharpen tool to make the colony outlines more distinct. The colony area was measured using the elliptical selection tool. We used 10 different genotypes from different MA lines and timepoints in this experiment, including the 2 ancestors. We collected a dataset with 482 area measurements. To estimate the average colony size, we fitted a multilevel model

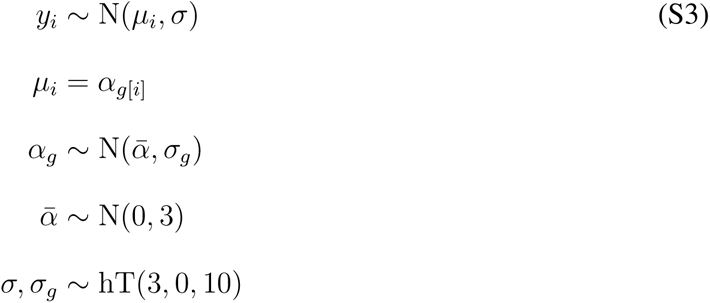

where *y_i_* is the *i*th area measurement, *ᾱ* is the overall mean, *α_j_* is the mean for *j*th genotype, *σ_g_* is the genotype standard deviation, and *σ* is the error standard deviation. Standard deviations had a weakly informative prior, which was the half-location scale version of Student’s t-distribution, where 3 is the degrees of freedom, 0 is the location, and 10 is the scale parameter. We estimated the number of nuclei in a sorbose plate colony, *n_s_*, as

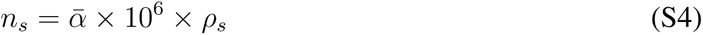

the average colony size is multiplied by 10^6^ to transform the unit from mm^2^ to µm^2^.

Once the sorbose colony is transferred to the test tube, the mycelium will cover the surface of the growth media. We estimated the number of nuclei present in the mycelium, *n_v_* by multiplying the surface of the media in the test tube with the density of nuclei in the hyphae in VM medium:

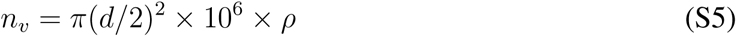

where *d* is the diameter of the test tubes (in mm) used in the experiment, area is multiplied by 10^6^ to transform the unit to µm^2^.

To estimate the number of conidia produced by the mycelium in the test tube, we counted conidia by suspending them in 1 mL of 0.01% Tween-80, making a 10000-fold dilution of the suspension, and plating 10 µL of the dilution on sorbose plates. We counted the colonies that were formed, and estimated the original number of conidia produced. We used 10 different genotypes, including the ancestors from the MA experiment to estimate produced conidia. We collected 71 measurements, the model was

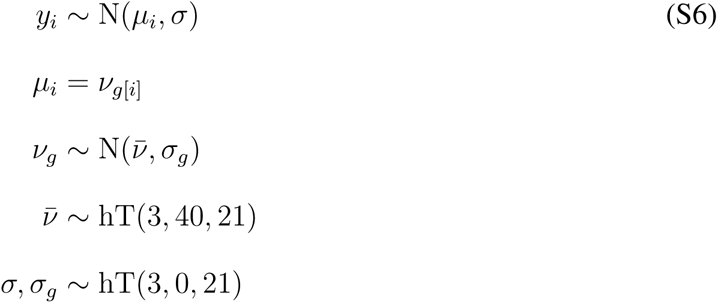

where *y_i_* is the *i*th conidial number measurement, *ν̄* is the overall mean, *α_j_* is the mean for *j*th genotype, *σ_g_* is the genotype standard deviation, and *σ* is the error standard deviation. Priors followed Student’s t-distribution. The number of nuclei contained by the conidia, *n_c_* was estimated as

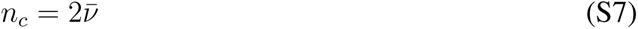

since the mode of nuclei in conidia of *N. crassa* is two.

The number of mitotic divisions separating two time points can be calculated from equation S1. First, we need to calculate the number of divisions that happened when a single spore grows to a colony on sorbose plate, then the number of divisions when the colony grows to a lawn of mycelium in the test tube, and finally the number of divisions it takes to form the final number of conidia. Thus, using the posterior distributions of numbers of nuclei in the different phases of the transfer and equation S1, we can calculate the number of mitoses that happen during a transfer, *m*, as:

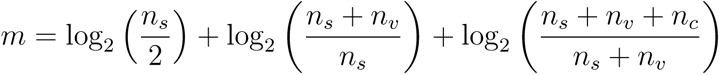

which simplifies to

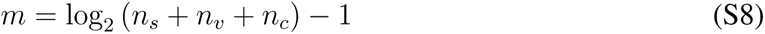

this estimate of the number of mitoses incorporates all sources of measurement error since posterior distributions are used in every step of the calculations.

#### DNA extraction

To get high quality DNA for sequencing, the natural strains, MA lines, and the ancestors were grown in 5 mL of liquid VM for two days at 25 °C with shaking. We harvested the mycelium and freeze dried it over night in a lyophilizer. Dried mycelium was then ground with a glass bead in Qiagen Tissue Lyzer for two times 20 s with frequency of 25 s^−1^. Then 500 µL of extraction buffer was added (10 mM Tris pH 8, 0.1 M EDTA, 150 mM NaCl, and 2% SDS), and the powdered tissue dissolved by shaking. Then samples were extracted with 750 µL of 25:24:1 Phenol:Chloroform:Isoamylalcohol and keeping the aqueous phase. We added 2 µL of RNAse A (10 mg/mL) and 50 U of RNAse I to each sample and incubated them for 1 h at 37 °C. Samples were then extracted with 750 µL of chloroform, 1 mL of 100% ethanol was added, and DNA was precipitated for 1 h at −20 °C. Then DNA was pelleted with centrifugation at 4 °C, ethanol aspirated, pellet washed with 70% ethanol, and air dried. We then added 77.5 µL of TE-buffer to elute the samples and incubated at 37 °C to help dissolve the pellets. We observed that occasional small DNA fragments would remain in the samples and to remove these we did a polyethyleneglycol precipitation: we added 12.5 µL of 4 M NaCl, mixed and added 12 µL 50% PEG (P3350), mixed and precipitated DNA over night at 4 °C. DNA was then pelleted with centrifugation and the supernatant aspirated, the pellet was washed twice with 70% ethanol, and aspirated. Pellets were eluted to 55 µL of TE-buffer as above. DNA concentrations were measured with the Qubit Broad Range Kit, and DNA quality was checked by running 2 µL of the sample on an 0.8% agarose gel.

#### Read mapping and genotyping

To be able to map reads to the mating type locus in the *mat a* strains, we included the mating type *a* region, as well as the mitochondrial genome, as additional contigs. Reads were mapped using BWA-MEM version 0.7.12-r1039 with default parameters (Li, 2013). Alignment files were sorted and indexed with samtools and read groups were added with picardtools. See table S1 for alignment metrics.

We used the GATK version 4.2.0.0 (McKenna et al., 2010) pipeline to call single nucleotide mutations (SNMs) and small indels. First, we ran Haplotypecaller for each sample individually to make a g.vcf file. Haplotypecaller was run with otherwise default parameters, emitting all sites, and in diploid mode. We then consolidated all of the samples together into a database using the GenomicsBDImport function in GATK. Samples were then jointly genotyped with the Genotype-GVCFs function to produce a vcf file with all samples.

We used wormtable version 0.1.5 (Kelleher et al., 2013) to convert the vcf file into an indexed database and then a custom Python script to filter for high quality sites. For a site to be included as a candidate mutation, first we required the genotypes of the ancestor and the MA line to differ for that site. Second, the site had to have five or more reads from both the ancestor and the sample. Third, the site had to have genotype quality greater or equal to 30 for both the ancestor and the sample, and finally sites that were called heterozygous in either the ancestor or the sample were excluded. There was also a filter that a site could not be called as a mutation if all of the MA lines had the same genotype. Sites were considered as invariant if their reference genotype quality was greater or equal to 30.

To produce the final dataset of curated mutations, we checked all candidate mutations manually by inspecting the alignments from BWA and or Haplotypecaller in IGV (Thorvaldsdóttir et al., 2013). Based on our manual inspection our filtering criteria were stringent enough, for our high coverage haploid genomes, to remove mapping errors and leave only real mutations, as only very few candidate mutations had to be rejected based on manual inspection and most mutations were unambiguous.

For genotyping SNPs in the strains sampled from natural populations, the above pipeline was used to call the genotypes. Other variants than SNPs were excluded. For a site to be included, it had to be polymorphic in the sample, with a mean read depth five or greater, genotype quality 30 or greater, and mapping quality 40 or greater across all samples. Then these same criteria were applied for each individual sample, and if a sample failed to meet the quality filters, its genotype was recorded as missing data. Heterozygous sites were excluded. Sites were also excluded if > 90% of samples had missing data. Sites were called as monomorphic if the mean reference genotype quality was 30 or greater and read depth 5 of greater across all samples. Then these same criteria were applied to individual samples, genotypes were recorded as missing data if a sample did not pass the filters.

#### Genotyping structural variants

There are several algorithms available to detect structural variants (SVs) from short-read sequencing data. However, because this kind of data is prone to base calling and alignment errors, none of the available computational algorithms can accurately and sensitively detect all types and sizes of SVs (Kosugi et al., 2019). To overcome this limitation it is common to use several algorithms and merge their outputs to increase sensitivity and precision. First, we assessed the performance of four different SVs algorithms (DELLY, Lumpy, PINDEL and SVaba) using simulated data.

We evaluated the performance of different SV callers on simulated data created using SURVIVOR version 1.0.7 (Jeffares et al., 2017). SURVIVOR simulates SVs by first modifying a fasta reference file by randomly altering locations according to given parameters of length and number of different SVs types (insertions, deletions, duplications, inversions and translocations). Reads are simulated based on the modified fasta and SVs are detected using the preferred SV caller. Finally SURVIVOR compares the SVs detected against the known simulated SVs, based on this FDR and sensitivity can be calculated.

We simulated four sets with 18, 40, 50 and 120 structural variants with a mutation rate of 0.001 on the reference genome (assembly NC12). The number of each type of SV simulated in each set is presented in the table S6. In set number four we simulated 20 complex SVs in which inversions and deletions occur in the same location. For duplications the min and max length parameter was set to 100-1000 bp, for INDELs 20-500 bp, for translocations 1000-3000 bp and for inversions 600-800 bp.

The SV length distribution across our four simulated sets were very similar (Figure S12), and the distribution coincides with ones reported in the literature, which indicates that short SVs are more common than large ones (Jeffares et al., 2017). The number of reads, the error rate and the coverage of the simulated data represent our sequenced reads. The inflated number of SV per genome is for testing purposes.

Based on the modified fasta we created 150 bp pair end reads with an error rate of 0.003% and a mean coverage of 30X using DWGSIM version 0.1.11 (Homer, 2021). Simulated reads were then aligned to the reference genome using BWA-MEM (Li, 2013) with default parameters, and SVs were called using DELLY version 0.8.7, LUMPY version 0.2.13, PINDEL version 0.2.5b9 and SVaba version 1.1.0 (Rausch et al., 2012; Layer et al., 2014; Ye et al., 2009; Wala et al., 2018). Finally, we used SURVIVOR to evaluate the performance of each SV caller. The SV calls were considered correct if the simulated and detected SVs were 1) of the same type 2) on same chromosome and 3) both start and stop locations were within 50 bp. The callers that performed the best were DELLY and LUMPY as they showed high sensitivity score and low false discovery rate (FDR) score (Table S6), and they were selected to call SVs on the MA lines.

For calling SVs in the MA lines we first aligned the reads to the reference genome using BWAMEM, excluded duplicated reads with SAMBLASTER version 0.1.26 (Faust and Hall, 2014), and extracted the discordant paired-end and split-read alignments using SAMTOOLS version 1.9 (Danecek et al., 2021). DELLY was used as indicated in the recommended workflow (Rausch et al., 2012). For LUMPY the read and insert lengths were extracted from alignment files using SAMTOOLS and the SVs were genotyped using SVTyper version 0.7.1 (Chiang et al., 2015). To filter out SVs that were present in the ancestor we used SnpSift version 5.0e (Cingolani et al., 2012). We removed those calls with a genotype quality score lower than 30 and read depth below 10. The analysis with both callers were carried out in somatic-germline mode, considering MA line as somatic and the ancestor as the germline. The signature of a translocation are reads with discordant mate pairs, where both mates are consistantly mapped to other chromosomes for example. Translocation length was determined from the break points of these discordant reads. All of the SVs detected by each caller were manually verified by inspecting the alignment files in IGV.

#### Genotyping copy number variants

To evaluate the performance of copy number variant (CNV) detection algorithms, we simulated 32 CNVs using SECNVs version 2.7.1 (Xing et al., 2020), then simulated 150 bp paired end reads with an error rate of 0.03% and a mean coverage of 30X using DWGSIM. We scanned for copy number variants (CNVs) using two detection programs, CNVnator version 0.4.1 (Abyzov et al., 2011) and CNV-seq version 0.2-7 (Xie and Tammi, 2009). CNV-seq was used with default parameters while CNVnator was used with two different bin sizes, 75 and 1670. Bins of 75 bp allowed the detection of small events, while bins of 1670 bp, which is the average gene length of *N. crassa* (Galagan et al., 2003), allowed the detection larger-scale events. Both callers togetherperformed better that any of the callers individually by showing the lowest FDR rate score of 0.482, and good sensitivity score of 0.906 (Table S7).

For genotyping CNVs in the MA lines we excluded MA line sites if the start or stop location of these where within 500 bp of any site detected in the ancestor. Also, we only retained the sites that were detected by both callers CNVnator and CNVseq (if 1000 bp or less overlapped at the start or end location). The remaining sites were manually verified by inspecting the alignment file in IGV. However, we did not find any evidence of copy number changes in the MA lines.

#### Chromatin modifications

To determine regions of the genome where chromatin modifications occur, ChIP-seq reads for H3K9me3, H3K27me3, and H3K36me2 were aligned to the reference genome using BWA-MEM, and duplicate reads were removed by Picard tools. Domains of chromatin modifications were identified using RSEG 0.4.9 (Song and Smith, 2011). Data for centromeric regions were obtained from Smith et al. (2011) and coordinate corrections for NC12 from Wang et al. (2020). The centromeric regions were defined based on the presence of centromeric histone 3 variant: CENPA. Smith et al. (2011) collected ChIP-seq data against CENPA and other centromeric proteins. Centromeric sequences in *N. crassa* are composed of AT-rich sequences of degraded transposable elements. However, the repeat arrays are heterogenous due to action of RIP, making almost all sequence sufficiently unique to be able to map short reads to the genome (Smith et al., 2011).

Furthermore, we used the data of the duplicated regions that were defined by Wang et al. (2020). Wang et al. (2020) identified duplicated regions using BLAST, with the criteria of at least 100 bp alignment length and at least 65% sequence identity.

#### Analysis of relative mutation rate for different classes

For cases where the relative mutation rates were computed for different classes of mutations the model was:

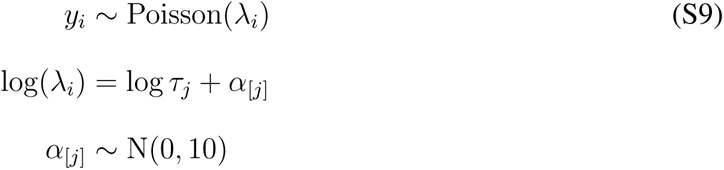

where *τ_j_* is an offset term for class *j* that allows taking into account differences in the abundance of certain classes (McElreath, 2015), such as higher frequency of A’s and T’s than G’s and C’s in the genome. Priors for different predictors remained the same as in equation 1. Furthermore, if we calculate the expected number of mutations for different classes under the assumption that all mutations in all classes are equally likely, as *τ_j_* = *f_j_n*, where *f_j_* is the frequency of class *j* and *n* is the total number of observed mutations, and use *τ_j_*, the expected number of mutations, as the offset parameter, then exp(*α*_[*j*]_) yields the relative mutation rate of class *j*. Since all estimates for different classes come from the same model, they are simultaneous comparisons in the statistical sense.

#### Mutation rate variation across the genome

To model the effects of epigenetic domains and GC-content on mutation rate we used the following model:

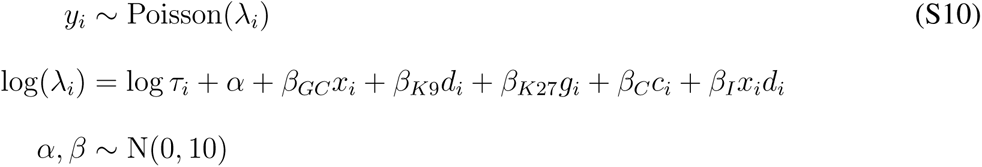

where *y_i_* is the number of mutations a class of *i* intervals contained, *τ_i_* is the number of class *i* intervals in total, *x_i_* the GC-content of those intervals, *d_i_* indicates presence or absence of H3K9me3, *g_i_* indicates presence or absence of H3K27me3, and *c_i_* indicates presence or absence of centromeric region. *β* coefficients are the corresponding effects and *α* is the intercept.

Model selection was a combination of biological and statistical reasoning, and we tested models representing plausible biological hypotheses. For instance, we had a clear biological reason to expect that GC-content influences mutation rate, and we saw a large improvement in model predictions when GC-content was included in the model. Therefore we did not further test models without GC-content and with different combinations of other terms. Furthermore, the only biologically realistic interactions are those involving GC-content and one of the domains. There are no regions where H3K27me3 and centromeric regions overlap, or regions where H3K9me3 and centromeric regions do not overlap, hence statistical interactions between domains are not possible in our data. Tested models are shown in Table S2, model comparisons were done using the widely applicable information criterion (WAIC) (McElreath, 2015; Vehtari et al., 2017).

When we assessed how well did the mutation model predict the natural genetic variation we used the predicted mutation rates from model S10 as a response and *θ_W_* calculated from a population sample of strains as a predictor in a simple regression model. Bayesian version of *R*^2^ (Gelman et al., 2019) was used to assess the model fit.

We could not asses the effect of duplicated regions defined by Wang et al. (2020) independently of H3K9me3 regions. Nearly all duplicated regions overlapped with H3K9me3 regions (Figure 2). Those regions that were marked as duplicates, but which did not overlap with H3K9me3 or H3K27me3, contained mainly mutations in microsatellite repeats. Only 10 point mutations were observed in these regions, which was not enough to obtain reasonable estimates of independent effect of duplicated regions on mutation rate. Of those 10 point mutations, 3 were C:G → T:A transitions. As C → T transitions were not over-represented, action of RIP is unlikely to be responsible for these mutations, which is expected as RIP is active only during meiosis.

#### Effects of local sequence context

To analyze effects of local base composition on the mutation rate, we estimated the effects of the trinucleotides from a model that included the effects of the epigenetic domains. First, we extracted the adjacent basepairs for every point mutation. There are 64 different trinucleotides, but as we cannot know in which strand the mutation originally occurred we grouped the trinucleotides into 32 different classes based on sequence complementarity. For example, trinucleotides ATA and TAT are complementary and were grouped. Then we counted how many times a given trinucleotide occurs in the genome in all three reading frames. Relative mutation rate was analyzed using the following model:

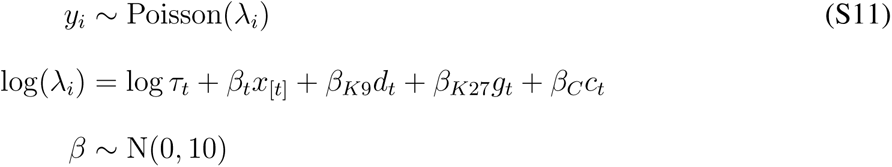

We compared different linear models (Table S4) with the same reasoning as above. We did not include an intercept in this model, as we wanted to obtain estimates for all trinucleotide classes, and not set one class as the intercept against which the others are compared. This does not alter any biological conclusions.

We further investigated how the flanking base pairs influenced the relative mutation rates of the trinucleotides. We extracted estimates of the relative mutation rates for the trinucleotides from model S11, and used these as a response in a model where we predicted relative mutation rates with the identities of the flanking base pairs and the mutating base. Since our estimates of the relative mutation rates contain uncertainty, we included the estimated error of the relative mutation rates in the model. The model was:

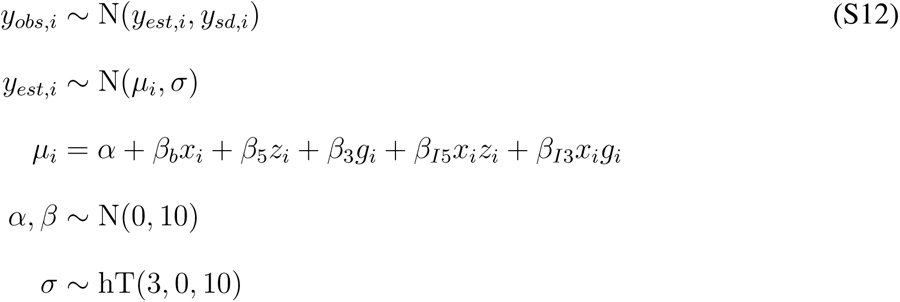

where *y_obs,i_* is the median of *i*th observed relative mutation rate, *y_sd,i_* is the observed standard deviation of the *i*th relative mutation rate, *y_est,i_* is the *i*th estimated relative mutation rate, *α* is the intercept, *β_b_* is the effect of C:G relative to A:T for the mutating base, *β*_5_ is the effect of C:G relative to A:T for the 5’ flanking base pair, *β*_3_ is the effect of C:G relative to A:T for the 3’ flanking base pair, *β_I_*_5_ is the interaction effect of 5’ CG when the mutating base is C:G, and *β_I_*_3_ is the interaction effect of 3’ C:G when the mutating base pair is C:G. *x_i_, z_i_*, and *g_i_* are indicators whether the basepair is C:G. We used the half location-scale version of Student’s t-distribution as a prior for the standard deviation with 3 degree’s of freedom, location 0, and scale 10.

### Supplementary results

#### Accuracy of mutation calling

Estimating mutation rates and particularly estimating differences in the mutation rate in different parts of the genome requires accurate mutation calls. As some regions of the genome, such as centromeric regions, may contain repetitive sequences it is important to verify that the mutations are called accurately in all regions of the genome, and that no region has an excess of false positive mutations. First, we examined sequencing coverage throughout the genome, GC-content does have an effect on sequencing coverage as regions of low GC can be preferentially amplified during library construction, and we observed slight elevation on normalized coverage around 35% GC (Figure S13). However, overall we observed that sequencing coverage was rather uniform across regions of different GC-content (Figure S13). Centromeric regions and regions marked by H3K9me3 have low GC-content, and while we did observe that coverage went down in regions of < 15% GC, those regions constitute a very small fraction of the genome. Next, we explored the accuracy of our mutation calls after the mutations had been called by our pipeline and manually inspected in IGV. We observed that overwhelming majority of mutations had the highest possible genotype quality score determined by the GATK pipeline (Figure S1). Median genotype quality for mutations was the highest possible value of 99, and only 8.6% of mutations had genotype quality less than 80 and only 1.9% less than 50. Distribution of quality scores was similar in different regions of the genome (Figure S1). While there was slightly more mutations that had lower quality scores than 99 in regions marked by H3K9me3 and in centromeric regions than in euchromatic regions (Figure S1), overwhelming majority of mutations in those regions have the highest genotype quality score of 99.

If most of the mutations had genotype quality scores of 99, then what kind of confidence we have in those mutation calls? We illustrate genotype quality scores with alignments viewed in IGV that show mutations in different regions of the genome and different genotype quality scores (Figure S14, S15, S16, S17, S18, S19, S20, S21, S22). When mutations had genotype quality score of 99 they were unambiguous (Figure S14, S17, S20). When genotype qualities were around 70 mutations could still be distinguished from unambiguously, even if few reads did not support the mutation or the mutations were in repetitive regions. When mutation genotype qualities were around 45 this was usually a sign that the region had lower mapping quality due to repeats or duplications (Figure S16, S19, S22). Despite of this, even in these regions, real mutations could be distinguished from mapping errors by looking at which reads supported the mutation and which did not (Figure S19, S22).

We have also provided screenshots of the alignments showing mutations viewed in IGV for a random sample of mutations. We selected mutations randomly, by first splitting the mutations into three genomic domains: H3K9me3, centromeric, and euchromatic, then drew a random sample of 30 from each pool, for a total of 90 mutations (see supplementary file S2). Information about the sampled mutations can be found in supplementary file S1.

The reason we chose first to do Sanger-verification for the mutations with the lowest genotype qualities was because for mutations with genotype quality of 99, there was no doubt that these were real mutations. We verified 23 base pair changes, of which 12 were in complex mutations and 11 as single nucleotide mutations. Of the 11 SNMs 5 were in regions marked by H3K9 (excluding centromes), 3 in centromeric regions, and 3 in euchromatin. Of the 12 base pair changes in complex mutations, 3 mutations were in H3K9 regions (5 base changes in total), 1 mutation in centromeric region (2 base pair changes), and 3 mutations in euchromatin (5 base pair changes). In the second verification set we sequenced 15 randomly sampled mutations from each genomic region (euchromatin, H3K9me3, and centromeric). One mutation located in centromeric region failed to amplify by PCR, the remaining 44 mutations were all confirmed. In summary, we confirmed point mutations by Sanger sequencing in centromeric, H3K9me3, and euchromatic regions. We confirmed all point mutations where PCR-amplification and Sanger sequencing were successful, so we never detected a false positive point mutation.

Why were the genotype qualities of the mutations so good in our experiment? There are several factors in this study that contributed excellent genotype calls. First, the ancestors for the MA lines were derived from line 2489 (synonym OR74a), which was the strain used for the original genome project (Galagan et al., 2003). Therefore, the reference genome used for read mapping corresponds to the genome of the MA line ancestors. This is seen in alignment metrics as 98% reads are mapped to the genome in the ancestors and MA lines (Table S1). As such, there are likely not many reads that would erroneously map to an incorrect location because their true source of origin was missing from the reference genome. Second, as explained in the introduction, repetitive sequences tend to diverge from each other in *N. crassa* due to the action of RIP. RIP does not induce the exact same mutations to the duplications, so over time duplicated arrays, such as those often found in centromeric regions, tend to diverge from one another, to the extent that short reads can be mapped to the genome in regions where it is not often possible to the same extent in other species (Smith et al., 2011). Third, the small genome of *N. crassa* made it possible to sequence the samples to a high depth, on average over 50x in many samples (Table S1). This allowed us to discriminate between true mutations and mapping errors. With this kind of sequencing depth, sequencing errors are simply not an issue anymore and they have no impact on calling the mutations, e.g. Figure S18 shows a mutation in repetitive region that as a consequence has higher frequency of sequencing errors, but with so many reads identifying the real mutation is not a problem. Finally, *N. crassa* is haploid. Combined with high sequencing depth, this makes identifying mutations easy. The only important errors are read mapping errors that may cause some sites to appear as heterozygous. But as heterozygous sites are not expected to occur in our experiment we can filter out sites called as heterozygous. We did inspect heterozygous sites manually, as it is possible that some mutations could have been present in a heterokaryotic state (nuclei with different genotypes in the same mycelium). However, we did not find any evidence of true mutations in heterokaryotic state. Whenever sites appeared as heterozygous, multiple sites were found close together (Figure S19), indicating that read mapping errors were the more likely explanation. Because of these factors, our study differs substantially from studies that need to call heterozygous sites from data with low sequencing depth and the problem of calling genotypes correctly is of different nature.

In summary, overwhelming majority of mutations that our pipeline detected had the highest possible genotype quality of 99, and this was true in regions of the genome with potentially more repetitive and duplicated regions like in centromeric regions and regions marked by H3K9 methylation. Those mutations that had genotype quality of 99 were unambiguously real mutations. Thus, even if we would filter out every mutation with genotype quality less than 99, we would still detect the observed pattern that mutation rate was higher in regions marked by H3K9 trimethylation and in centromeric regions. Differential mutation calling in different regions of the genome cannot explain the observed results.

#### Simulating variation in mutation rate

Despite our very high genotype qualities, we attempted to further understand could repetitive sequences or other sequence features of heterochromatin in the *N. crassa* genome hinder our ability to correctly estimate differences in mutation rates in different regions of the genome. We simulated data under two different scenarios. First, we simulated a scenario where mutation rate was set to be higher in H3K9me3 domains, with a rate of 2 × 10^−5^ mutations per site, compared to the rest of the genome, with a rate of 3 × 10^−6^ mutations per site. In the second scenario, we simulated a uniform mutation rate across the genome, with a rate of 2 × 10^−6^ mutations per site. We simulated mutations to the *N. crassa* genome using the program Mutation-Simulator (Kühl et al., 2021). We simulated 40 different MA lines for each scenario with a transition / transversion rate of 1.08. We then generated simulated reads from these simulated genomes, using DWGSM (Homer, 2021), with 30X sequencing depth and read length of 150 bp. We tried to imitate the conditions of our real sequenced data, so we set the standard deviation of the base quality scores to two and the per base sequencing error rate to 0.003. The ancestor of the MA lines was simulated by generating reads from the reference genome of *N. crassa*. To call the simulated mutations from the simulated reads, we ran the same pipeline as we used for the experimental data. Thus, we had two simulated scenarios, and for each scenario we had information about the true number of mutations that happened in the simulation, and number of mutations we called with our pipeline from the simulated read data. In the scenario with the higher mutation rate in H3K9me3 regions, we ended up with a total of 1759 mutations, of which 719 were in H3K9me3 domains, 990 in euchromatin, and 50 in unspecific domains. With our pipeline we detected a total of 1705 mutations, of which 692 were in H3K9me3 domains, 964 in euchromatin and 49 in unspecific domains. All of the called mutations were true positives. However, we failed to call 54 true mutations, that is, these were false negatives. In a similar manner, in the scenario with the uniform mutation rate, we ended up with a total of 3078 mutations, of which 562 were in H3K9me3 domains, 2245 in euchromatin, and 271 in unspecific domains. Our pipeline detected 2978 mutations in total, of which 535 were in H3K9 domains, 2177 in euchromatin, and 266 in unspecific domains. Again, there were no false positive calls. We failed to detect 100 mutations in this set. In general, the number of false negatives was higher in H3K9me3 regions, with proportion of false negatives 3.75% and 4.80% in H3K9me3 regions, and 2.62% and 3.02% in euchromatin in for the different and uniform mutation rate scenarios respectively.

We found that the estimated mutation rate was higher in H3K9me3 regions in the scenario where the true mutation rate was higher in H3K9me3 (Figure S23), the mutation rate ratio of H3K9me3 / euchromatin was 3.39 [3.06, 3.72]. This mutation rate ratio was not statistically different from the one calculated from the true simulated mutations: the difference was 0.28 [-0.15, 0.74], which includes zero in the interval estimate. Furthermore, when we simulated a uniform mutation rate across the genome, we found no difference among called and true datasets (Figure S23). The mutation rate ratio of H3K9me3 / euchromatin was 1.15 [1.05, 1.27], there was no statistical difference in the rate ratios between called and true simulated mutations: difference was 0.08 [-0.05, 0.23], which includes zero in the interval estimate.

With this simulation data we show that our pipeline can confidently detect a difference in mutation rates in different regions of the genome. This shows that sequence features of the H3K9me3 regions, such as repetitive sequences, do not interfere with mutation calling in a manner that would lead to gross biases in mutation rate estimates in the different domains. While simulated read data cannot capture all of the properties of real data, because of sequences missing from the reference or assembly errors, it does give us confidence that we will be able to detect a real difference in mutation rates. Moreover, since we did not observe any false positive mutations, we are confident that mutation calling cannot generate spurious results in our case. We did observe slightly higher proportions of false negative mutations in H3K9me3 regions. However, if this bias is true for real data, this would make our estimate of the elevated mutation rate in H3K9me3 regions more conservative.

#### Robustness of relationship between *θ_W_* and predicted mutation rate

We wanted to evaluate the robustness of the observed relationship between *θ* and the predicted mutation rate. One potential issue is that there are windows in the genome, especially for small window sizes, where the observed *θ* is zero. Since zero is the minimum value that *θ* can obtain, and there is a clumping of *θ* = 0 observations in the data, this violates the assumption that response is gaussian and could lead to biased estimates. However, since there so many data points, the model may be robust to observations where *θ* = 0. First, we tested the effect of window size, calculating *θ* over longer windows reduced the number of windows where *θ* = 0. Increasing window size slightly improves the amount of variation explained by the model (Figure S4). Thus, results are robust the to different window sizes.

Then we tested whether the results were robust to different models. Data that can take zero or positive values, but is clumped at zero, can be modeled in different ways. One possibility is Tobit regression. Tobit regression is a type of censored regression, where observations are assumed to have an underlying gaussian distribution, but appear as zeros if *y_i_ ≤* 0 (Min and Agresti, 2002). We used a conventional Tobit regression and robust Tobit regression, for both cases the results were very similar to an ordinary regression model (Figure S5). Then, we tested a log-normal hurdle model. In this model the response distribution is a mixture of two processes, one models the probability that the observation is larger then zero, and the other is a log-normal gaussian model (Min and Agresti, 2002). For the hurdle model, we also observed that that the relationship between *θ* and predicted mutation rate was positive (Figure S5). Therefore, our results are robust to the clumping at zero phenomenon.

Next, we tested whether the action of RIP could explain the relationship between *θ* and predicted mutation rate. If level of genetic diversity is very high in H3K9me3 regions due to C → T transitions induced by RIP, we want to make sure that this phenomenon does not solely cause the relationship between *θ* and predicted mutation rate. We cannot determine the exact contribution of RIP to genetic diversity, because we do not know the ancestral states of the SNPs and therefore cannot distinguish between C:G → T:A and A:T → G:C transitions. Furthermore, we would need to know the population recombination rate to estimate the number of meiotic divisions for every mitosis and thus the frequency of RIP. Therefore, we looked at the relationship between *θ* and predicted mutation rate within each of the genomic domains, and observed a positive relationship between *θ* and predicted mutation rate within each of the domains. Although, the effect was weak within centromeric domains (Figure S6A). We then filtered the SNP dataset to include only transversions and calculated *θ* across the genome. There was a positive relationship between *θ* for transversions only and the predicted mutation rate within all domains except H3K9me3 (Figure S6B). These results show that while RIP probably has a large contribution to genetic diversity in regions of H3K9me3, it does not solely drive the relationship between *θ* and predicted mutation rate.

#### Re-analysis of data from Wang et al. 2020

Wang et al. (2020) estimated the rate of spontaneous mutation during meiosis in *N. crassa*. During meiosis a genome defence mechanism called repeat-induced point mutation (RIP) induces mainly C → T transitions in duplicated regions of the genome resulting in a very high overall mutation rate (Wang et al., 2020). While not made explicit by Wang et al. (2020), the duplicated regions correspond almost completely to the H3K9 trimethylated domains. In order to better compare our results for asexual mutation rate in different domains to the sexual mutation rate estimated in their study, we re-analyzed the data from Wang et al. (2020) provided in their supplementary material, and included the information about chromatin domains. Their data are comprised of mutations in sequenced tetrads, which correspond to the products of a single meiosis. We included only those tetrads originating from crosses between non-mutant strains. This leaves 67 tetrads in the data that originate from five different crosses.

First we split the mutations to those that occurred in euchromatin and to those that occurred in H3K9 trimethylated domains. We observed that the numbers of mutations occurring in euchromatin and H3K9me3 domains for a given tetrad had very different distributions (Figure S10A), number of mutations occurring per tetrad in the H3K9me3 domains had a very long tail. When we examined the number of mutations per tetrad by cross, we observed a median of 22 mutations that occurred in euchromatic regions per tetrad, with some differences among the five crosses. However, the variation among tetrads from the different crosses was similar (Figure S10B). However, there were a median of 38 mutations that occurred in the H3K9me3 domains per tetrad, but a huge variation among tetrads, even within tetrads from a single cross (Figure S10B). For example, some tetrads from the same cross had 20 to 40 mutations, while others could have hundreds. In cross E the range of mutations was from 27 in one tetrad to 1187 in another. Variation among mutations in the H3K9me3 domains per tetrad suggest that while there probably were some genetic influences on the mutation rate in the different crosses, there was substantial heterogeneity in the activation of RIP that was independent of genetic effects.

We calculated the mutation rate per meiosis for the euchromatic regions of the genome using a multilevel model with cross as a random factor. The model was

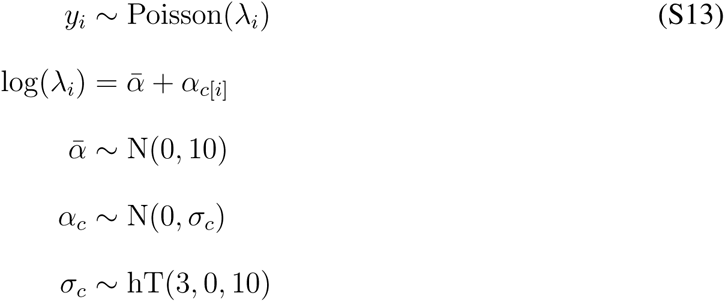

where *y_i_* is the number of mutations in euchromatic regions in the *i*th tetrad, *ᾱ* is the average intercept, *α_c_* is deviation from average intercept for each cross, and *σ_c_* is the cross standard deviation. Prior for *σ_c_* was the half-location scale version of Student’s t-distribution, with 3 degrees of freedom, location 0, and scale 10. Based on posterior predictive checks, this model fitted the data. Mutation rate was calculated from posterior distribution of *ᾱ* as

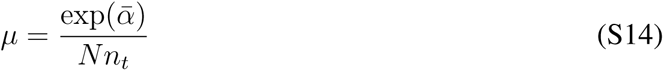

where *N* is the number of called nucleotides, and *n_t_* is the number of tetrads. The mutation rate in euchromatic regions during sexual reproduction was 1.07 [0.6, 1.67] ×10^−8^ mutations / meiosis / bp.

The data for mutations that occurred in the H3K9me3 domains are clearly overdispersed. To calculate the mutation rate per meiosis for the H3K9me3 domains we also modelled the heterogeneity among the tetrads. We fitted a gamma-poisson model, also called a negative binomial model, to the data. A gamma-poisson model allows each observation, a tetrad in our case, to have a different poisson rate allowing us to model this heterogeneity in observed rates (McElreath, 2015). We fitted a model

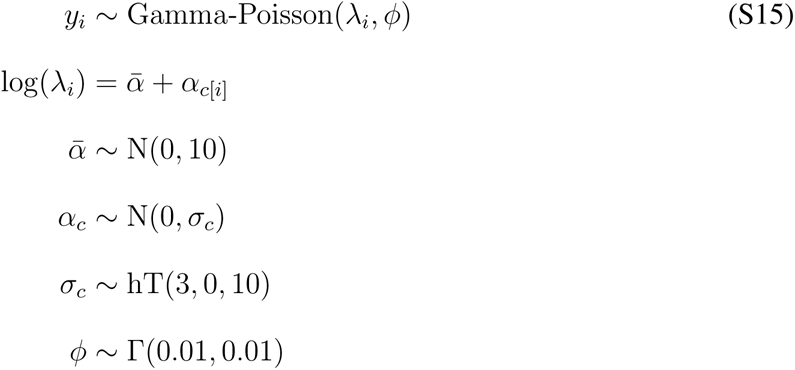

where *y_i_* is the number of mutations in the H3K9me3 domains in the *i*th tetrad, *ϕ* is the dispersion parameter, and other parameters were same as above. The prior for *ϕ* was a gamma distribution with shape of 0.01 and scale 0.01. Posterior predictive check indicated that the model fit the data reasonably well. The mutation rate was calculated from the average intercept as above. The mutation rate in H3K9 trimethylated regions during sexual reproduction was 2.54 [0.11, 7.55] ×10^−7^ mutations / meiosis / bp. As a result of rate heterogeneity there is quite a bit of uncertainty in the estimate. The ratio of mutation rates in the H3K9me3 regions over the euchromatic regions was 23.7 [0.99, 76.38]. While the 95% interval of the ratio slightly overlaps one due to large uncertainly in mutation rate in the H3K9me3 regions, mutation rate those regions seems higher.

We examined the spectrum of mutations that occurred in the euchromatic and the H3K9me3 regions separately, in the same way we did for asexual mutations. We observed that in the H3K9me3 regions there was a substantial over-representation of C:G → T:A transitions due to the action of RIP (Figure S10C). However, the mutation spectra that occurred in euchromatic regions was much more similar to the one we observed during asexual reproduction in euchromatic regions. There was no difference in the relative mutation rate of C:G → T:A transitions during sexual and asexual reproduction in euchromatic regions. Some of the transversions did have different relative rates: A:T → C:G, and C:G → G:C transversions had higher rate during sexual reproduction, while C:G → A:T transversions had a lower relative mutation rate during sexual reproduction (Figure S11).

Our analysis gives somewhat different results compared to those of Wang et al. (2020), who only calculated mutation rates across the whole genome, and did not take variation among tetrads or crosses into account. We do find higher mutation rates during sexual reproduction than during asexual reproduction, suggesting that in *N. crassa* meiosis is mutagenic in addition to the RIP effect in the H3K9me3 domains. However, the mutation rate per meiosis was much smaller than that estimated by Wang et al. (2020). The H3K9 trimethylated regions contain mainly degraded transposable elements, and are quite gene poor. If we compare non-synonymous mutations in euchromatic and H3K9me3 regions, of those mutations that occurred in euchromatic regions 22.16% were non-synonymous, while only 0.17% of mutations were non-synonymous in H3K9 methylated regions. Thus, the very high mutation rate observed in H3K9 regions due to action of RIP, does not necessarily translate into a high genetic load. We suggest that the mutation load during sexual reproduction in *N. crassa* may not be as high as it has been suggested by Wang et al. (2020).

## Supplementary Figures

**Figure S1:**
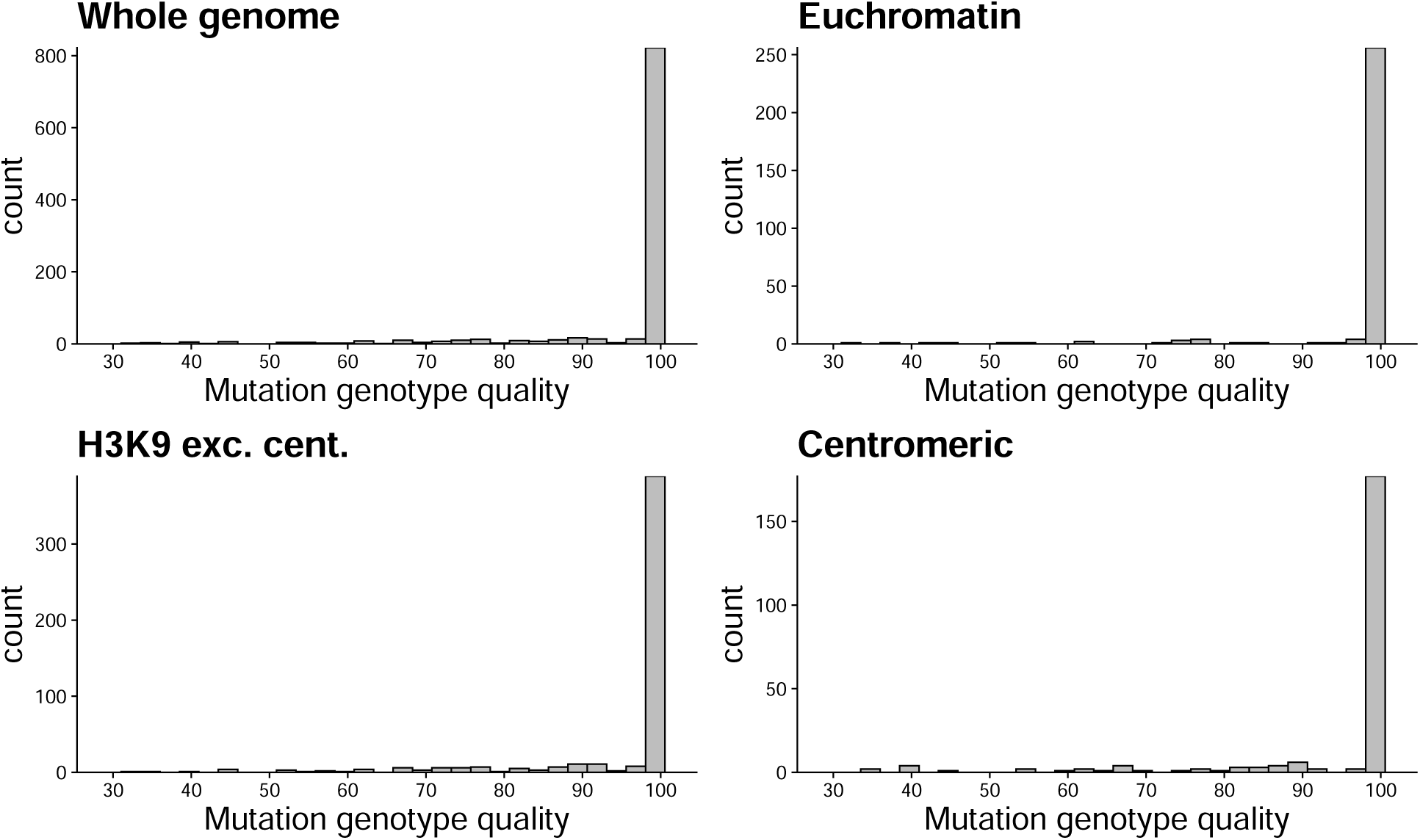
Distribution of genotype qualities of observed mutations given by GATK. Distributions are shown for the whole genome, euchromatin, H3K9me3 domains excluding centromeric regions, and centromeric regions.

**Figure S2:**
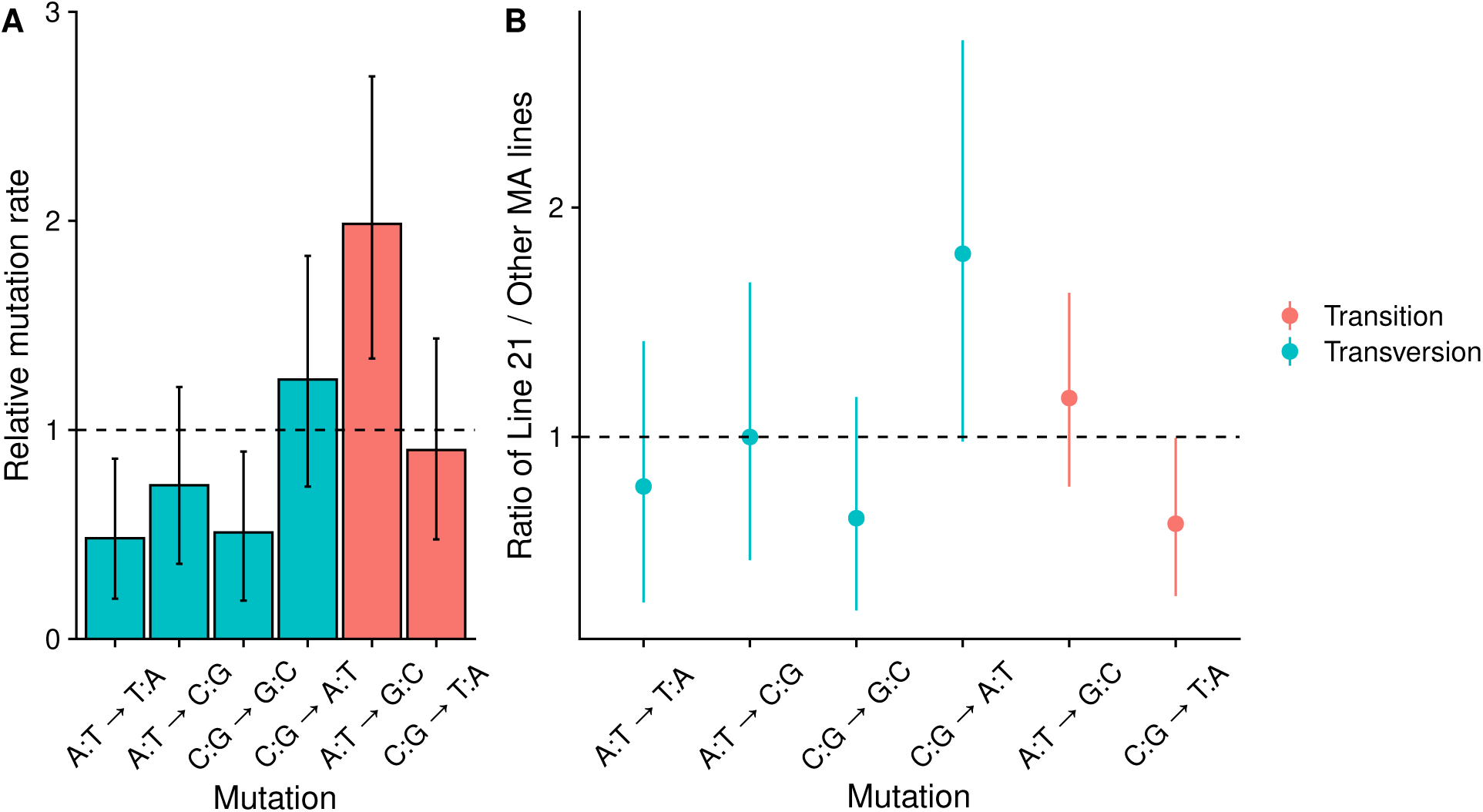
A) Mutation spectra for the MA line 21, this line had an excess number of mutations. B) Ratios of relative mutation rates for line 21 / rest of the MA lines. Intervals for C:G → A:T transversions and C:G → T:A transitions barely overlap one.

**Figure S3:**
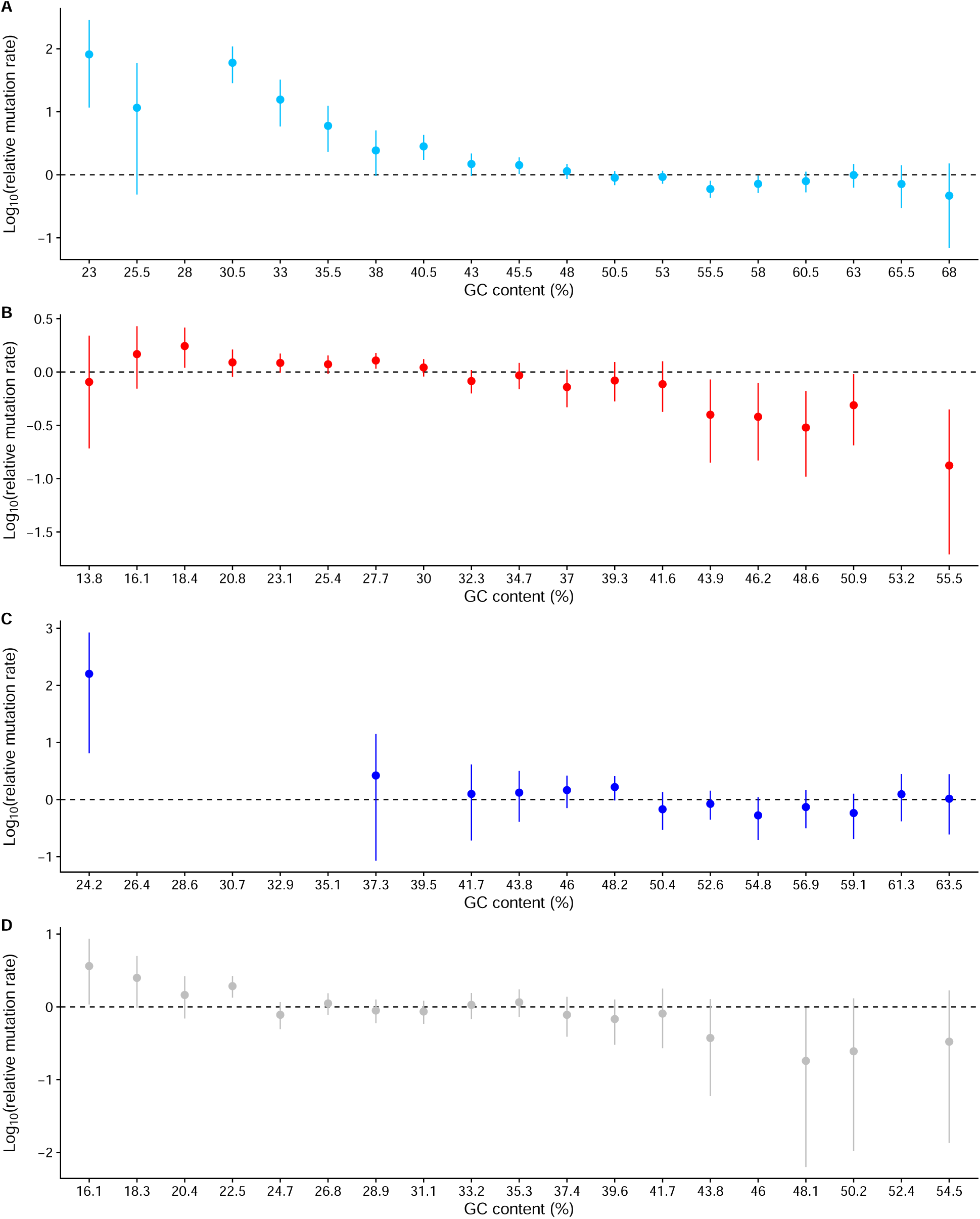
Relative mutation rates for windows of 200 bp binned for GC-content at 2.5 percentage point intervals. Ticks on the horizontal axis are at the end points of intervals. Note that y-axis is on a log_10_ scale, the dashed line indicates relative mutation rate of one. Some bins did not contain any mutations, so estimates are missing for those bins. A) Euchromatic regions B) H3K9me3 domains C) H3K27me3 domains D) Centromeric regions.

**Figure S4:**
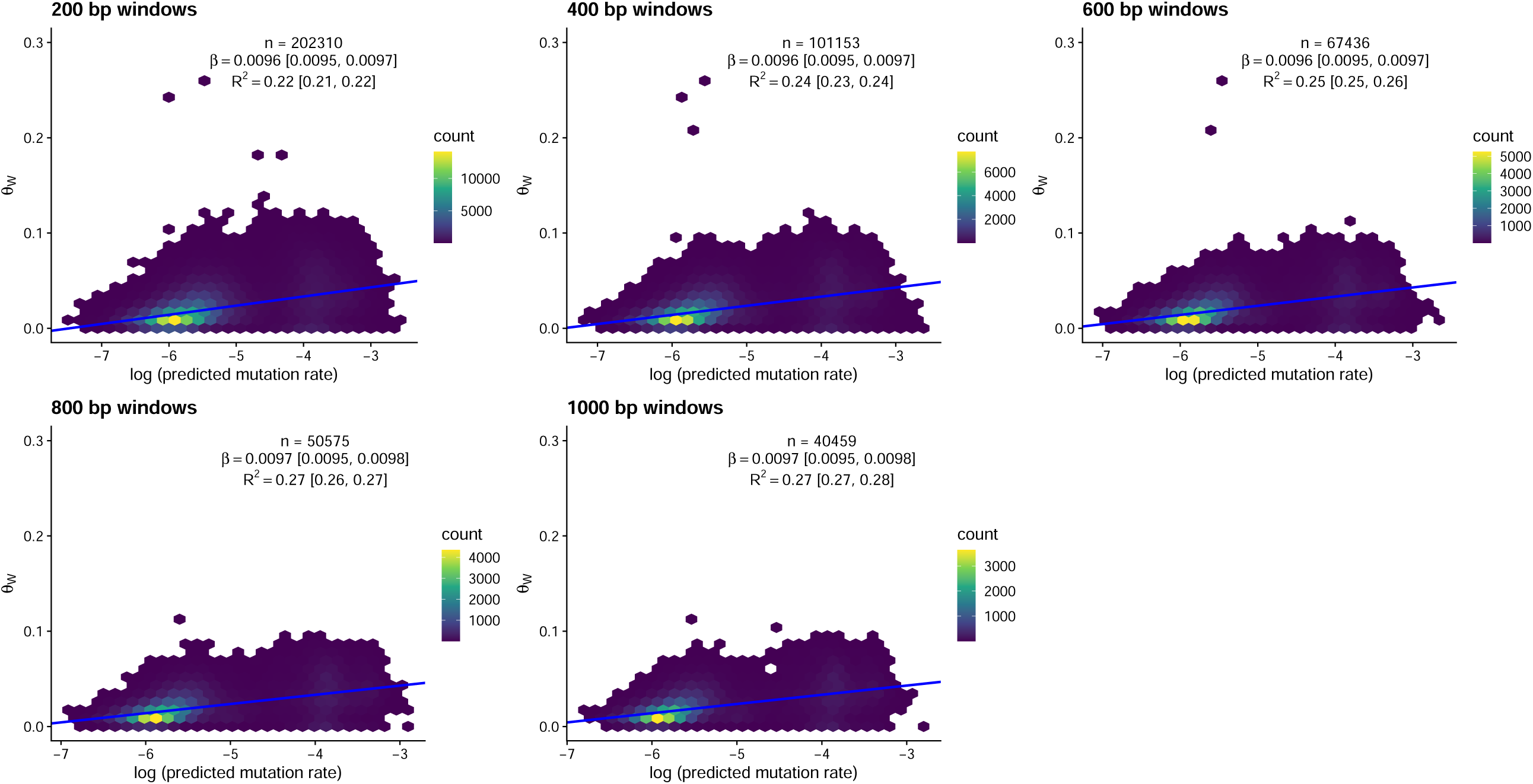
Regression between the predicted mutation rate and the observed nucleotide polymorphism in a sample of strains from natural populations. Results have been calculated for different window sizes. *n* is the number of windows, *β* is the slope of the regression line, and *R*^2^ is the Baysian *R*^2^ value, a measure of model fit.

**Figure S5:**
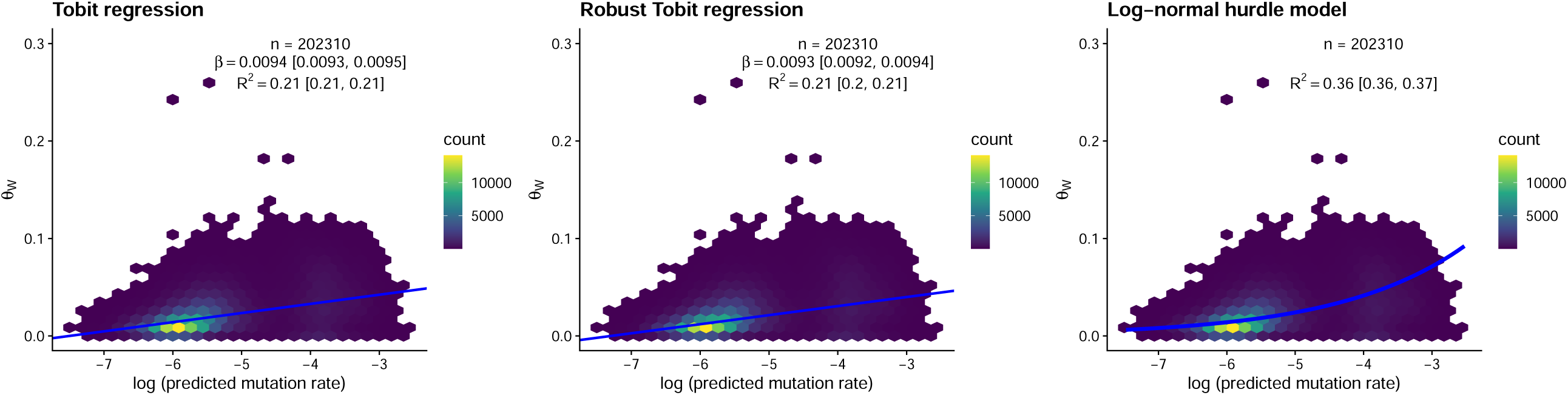
Regression between the predicted mutation rate and the observed nucleotide polymorphism. To check the robustness of results to windows where *θ* = 0, different models were used. Window size = 200 bp, *n* is the number of windows, *β* is the slope of the regression line, and *R*^2^ is the Baysian *R*^2^ value, a measure of model fit.

**Figure S6:**
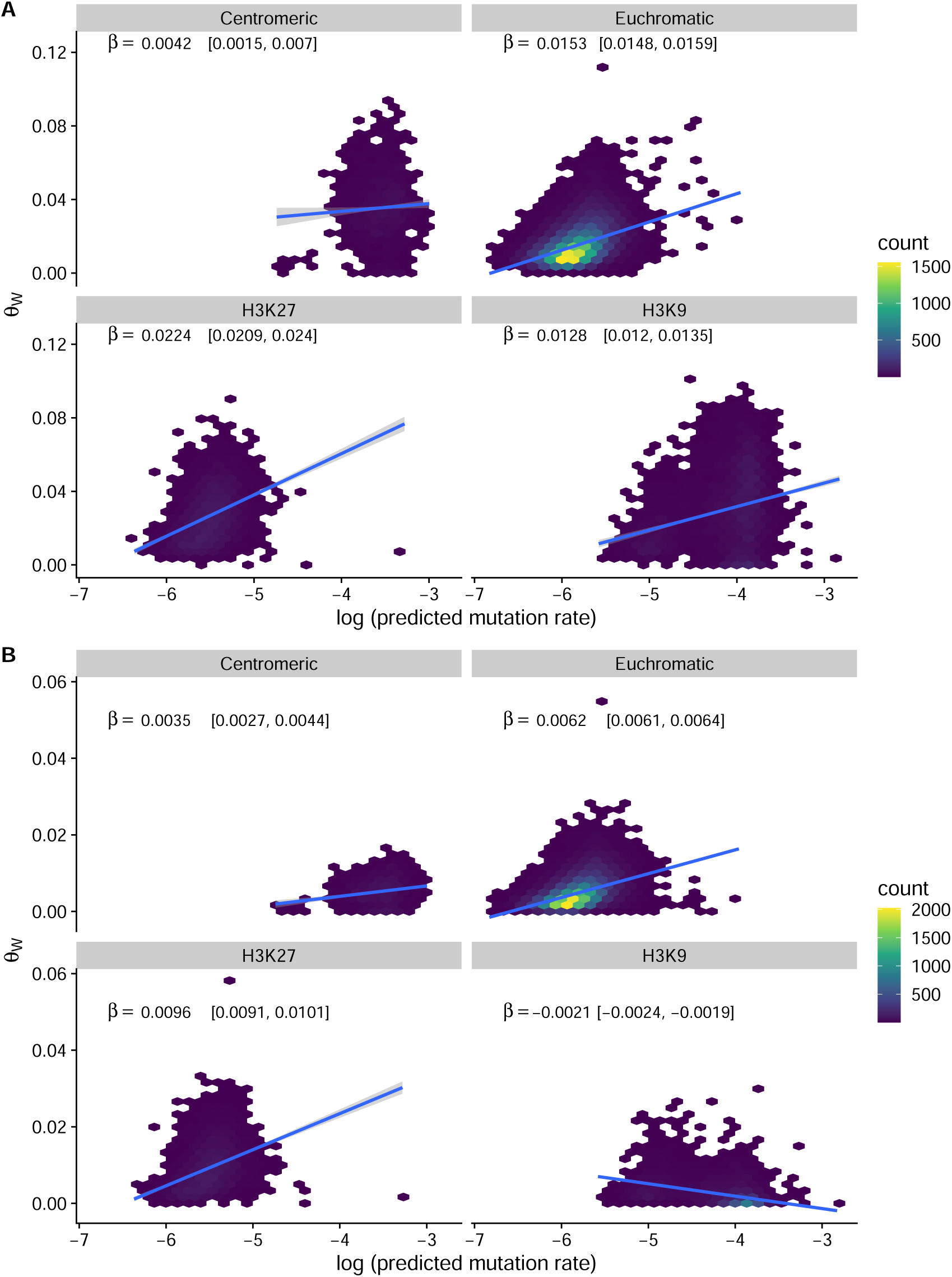
Regression between the predicted mutation rate and the observed nucleotide polymorphism within different regions of the genome. Window size was set to 1000 bp in both panels, as there are large number of windows where *θ* = 0 for small window sizes in the transversions only data. *n* = 40459, *β* is the slope of the regression line. A) *θ* has been calculated for all SNPs, *R*^2^ = 0.32 [0.32, 0.33]. B) *θ* has been calculated only for SNPs that represent transversions, *R*^2^ = 0.29 [0.28, 0.30].

**Figure S7:**
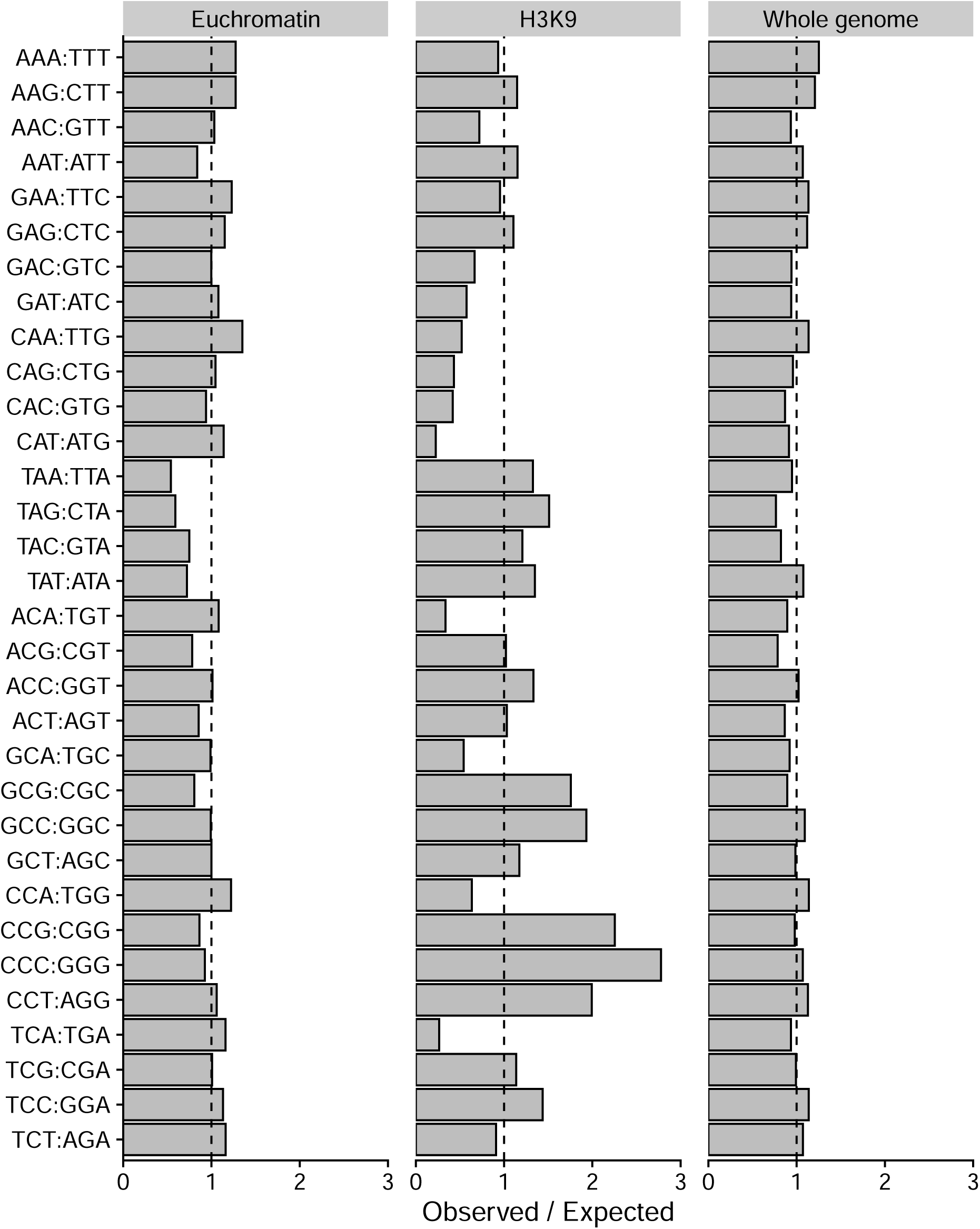
Observerved deviations of trinucleotide frequencies from expectations for different parts of the genome. Observed trinucleotide frequencies were divided by their expected frequencies based on GC-content. The dashed line shows the expected ratio of one.

**Figure S8:**
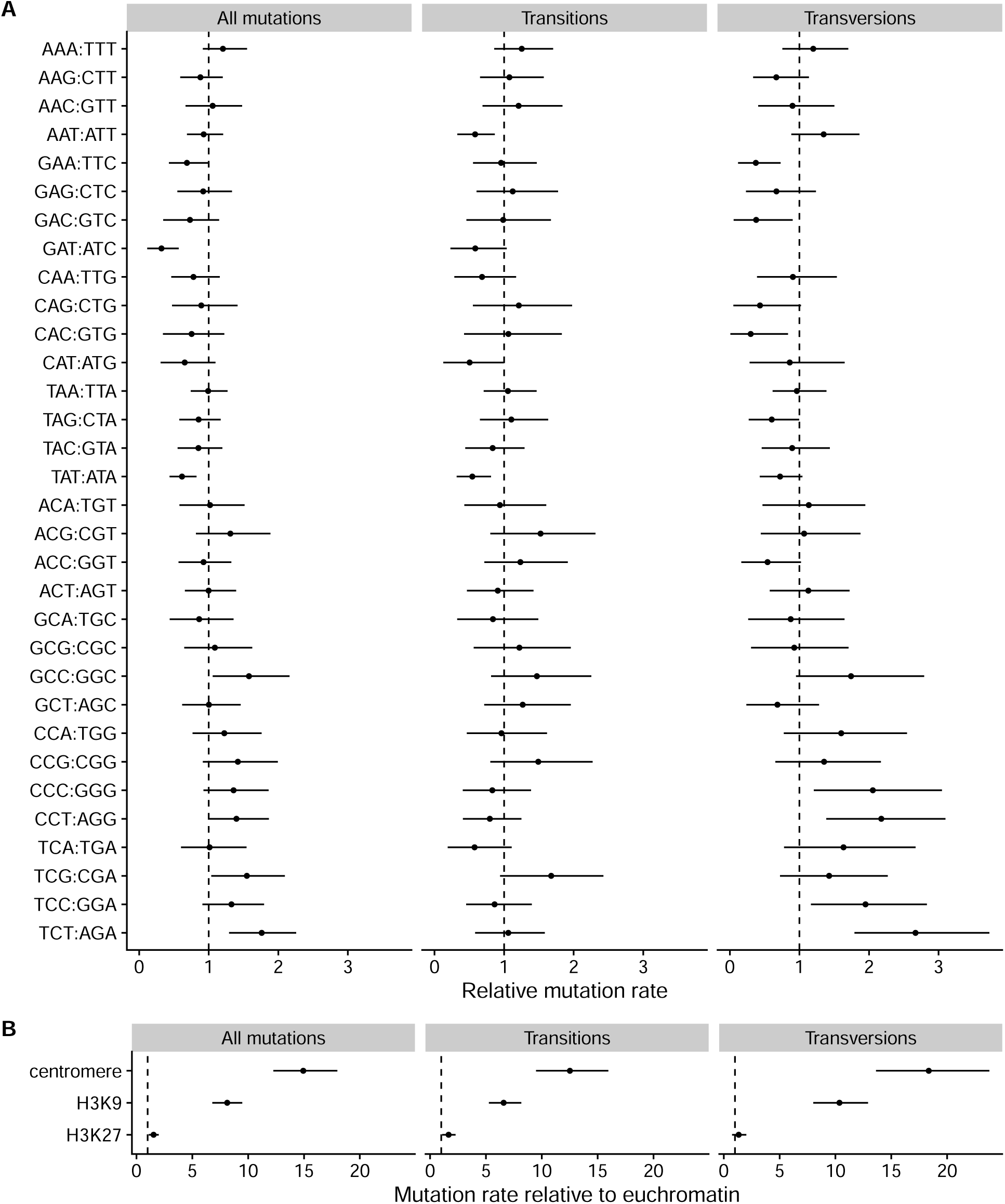
A) Relative mutation rates for the 32 different trinucleotide classes. B) Model estimates for relative mutation rates for centromeric, H3K9me3 and H3K27me3 domains from the trinucleotide model. Estimates are medians and range shows 95% HPD intervals.

**Figure S9:**
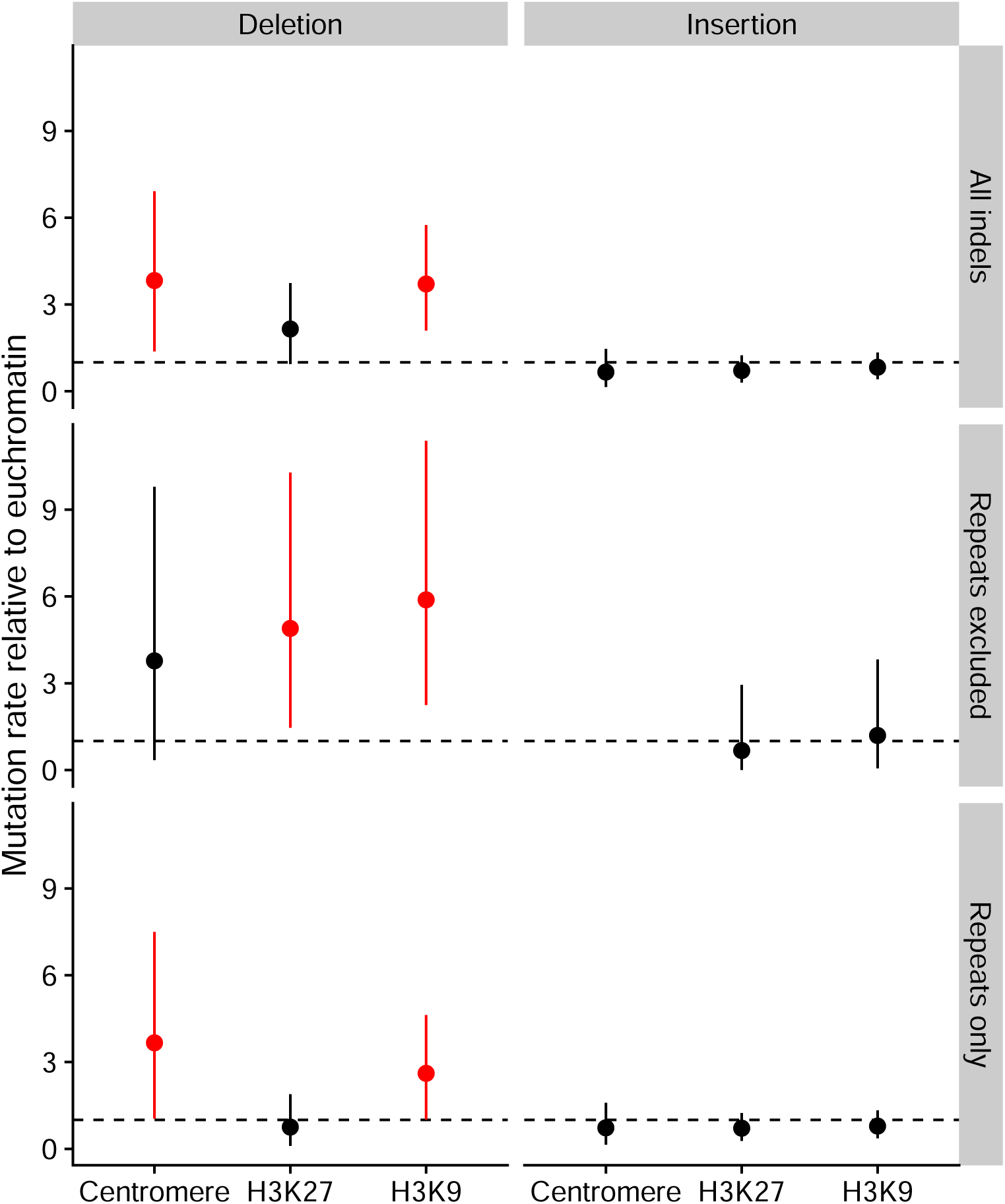
Mutation rates for indels in the different domains relative to euchromatin. The dashed line shows the relative mutation rate of one. Facets show deletions and insertions for all indels, for indels that did not occur in repeats, and for indels that occurred in repeated sequences. Estimates are medians and ranges show 95% HPD intervals. Intervals that do not overlap with one, that is, those where mutation rate is higher than in euchromatin are colored red.

**Figure S10:**
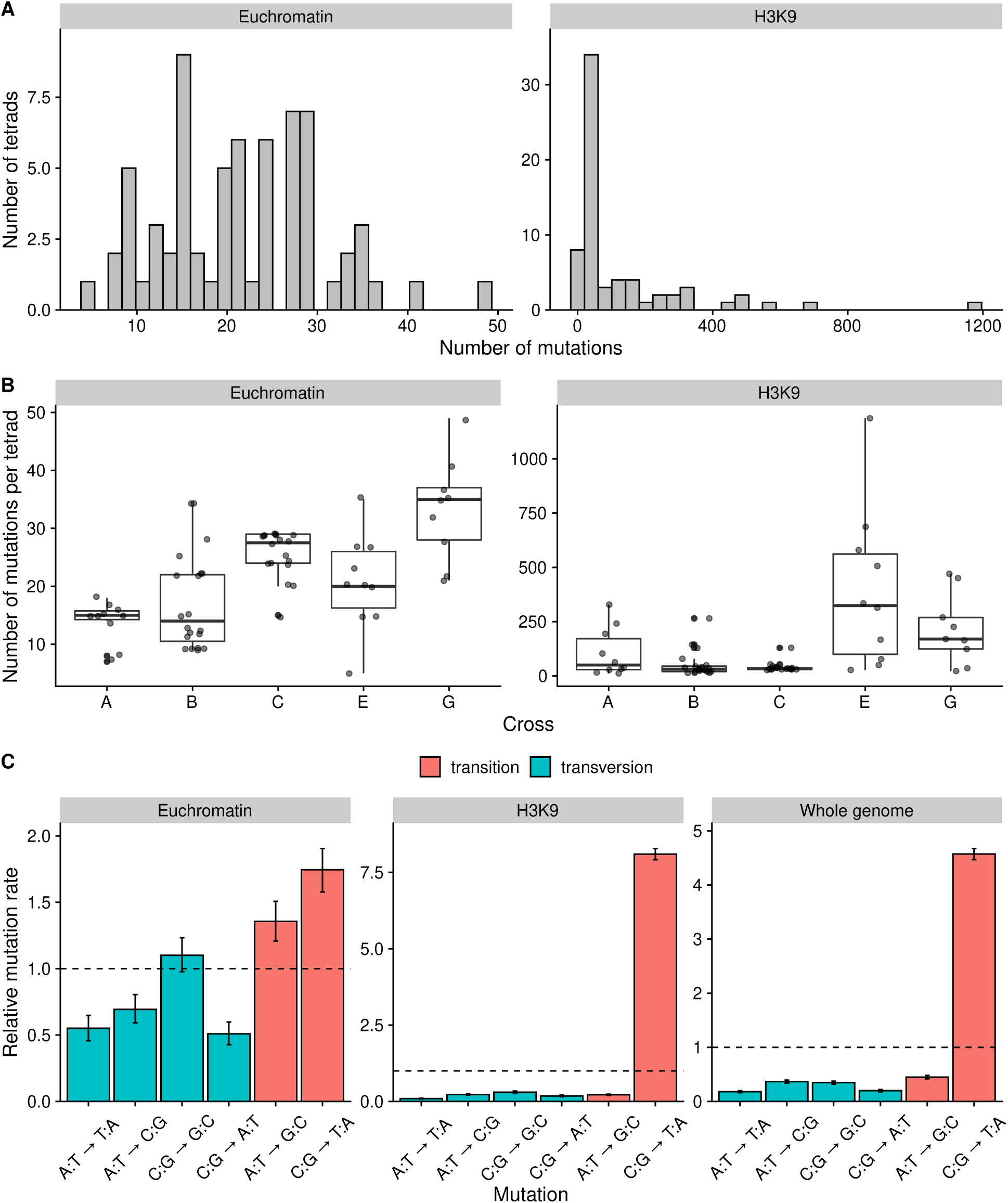
Mutations that occurred during sexual reproduction. Data is from Wang et al. (2020). Note that y-axis scales are different in different panels. A) The distribution of the number of mutations in the tetrads in euchromatin and H3K9 methylated domains. B) The number of mutations per tetrad for the different crosses. C) Spectrum of mutations for different regions of the genome. Error bars are 95% HPD intervals.

**Figure S11:**
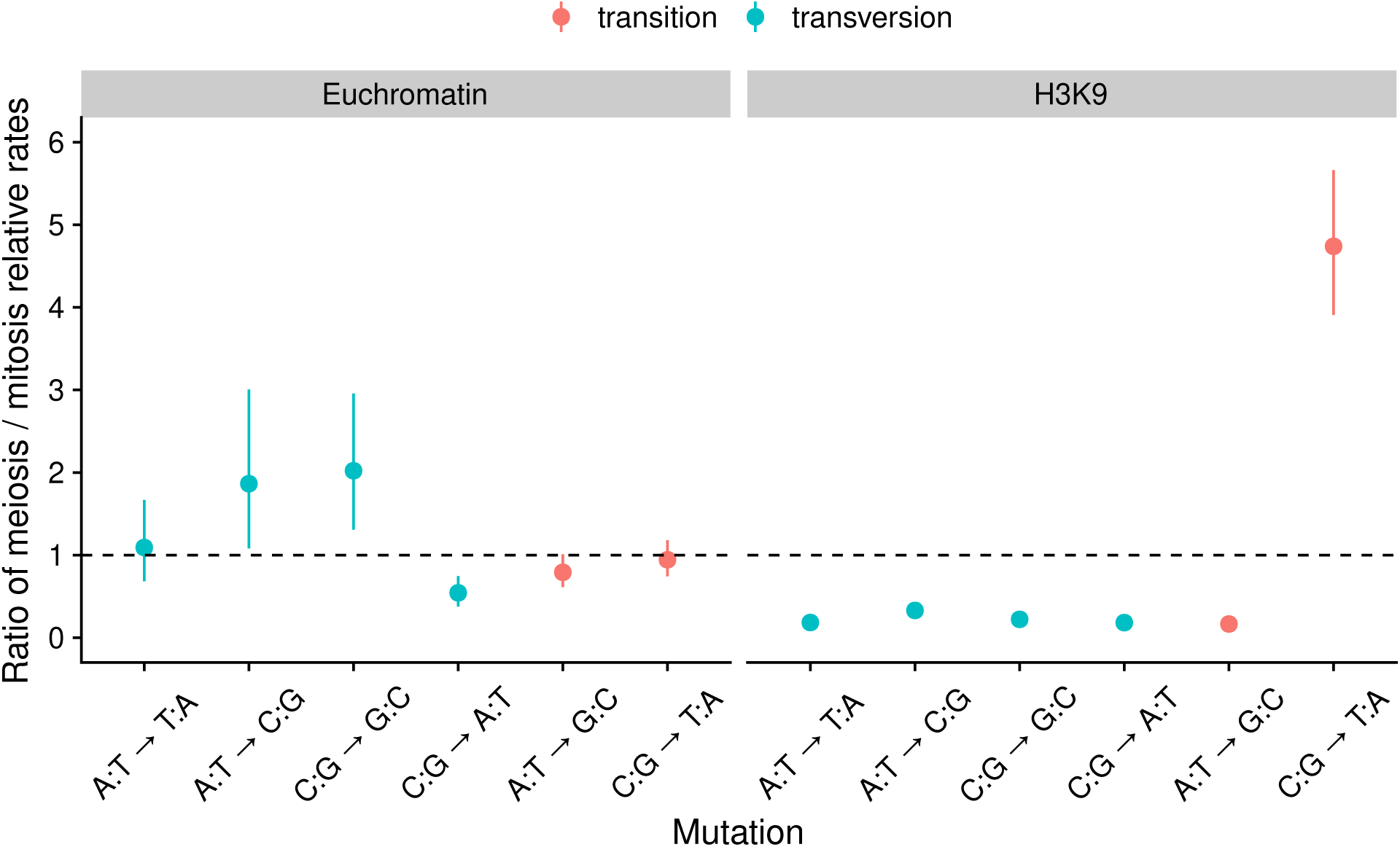
Ratios of the relative mutation rates during meiosis over mitosis. Points are medians and ranges show 95% HPD interval of the ratios. If the interval estimate is higher than one, mutation rate in meiosis is higher, if the interval estimate is lower than one, mutation rate in mitosis is higher.

**Figure S12:**
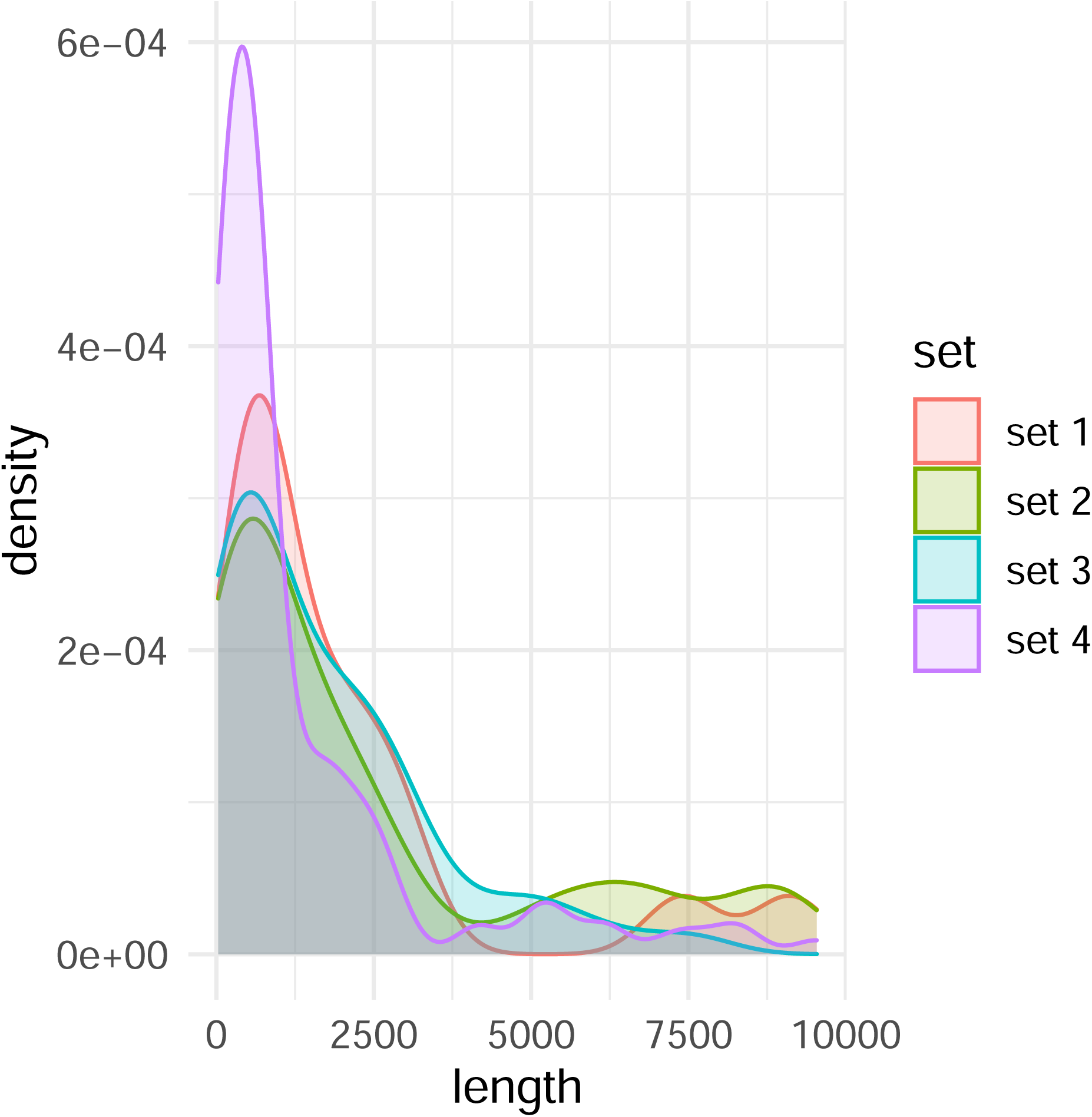
Densities for the length of distributions of SVs simulated using survivor. The characteristics of each simulated set are specified in the supplementary table S6.

**Figure S13:**
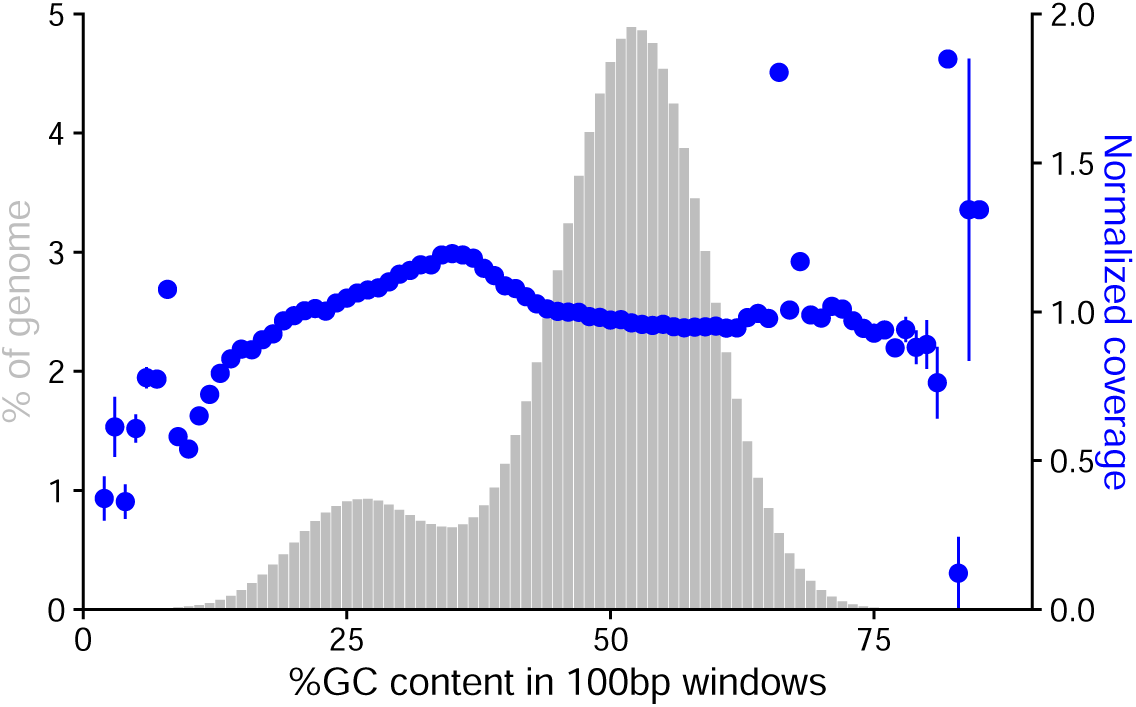
Sequencing coverage plotted against GC-content of the genome for the *mat A* ancestor. Other samples had similar profiles.

**Figure S14:**
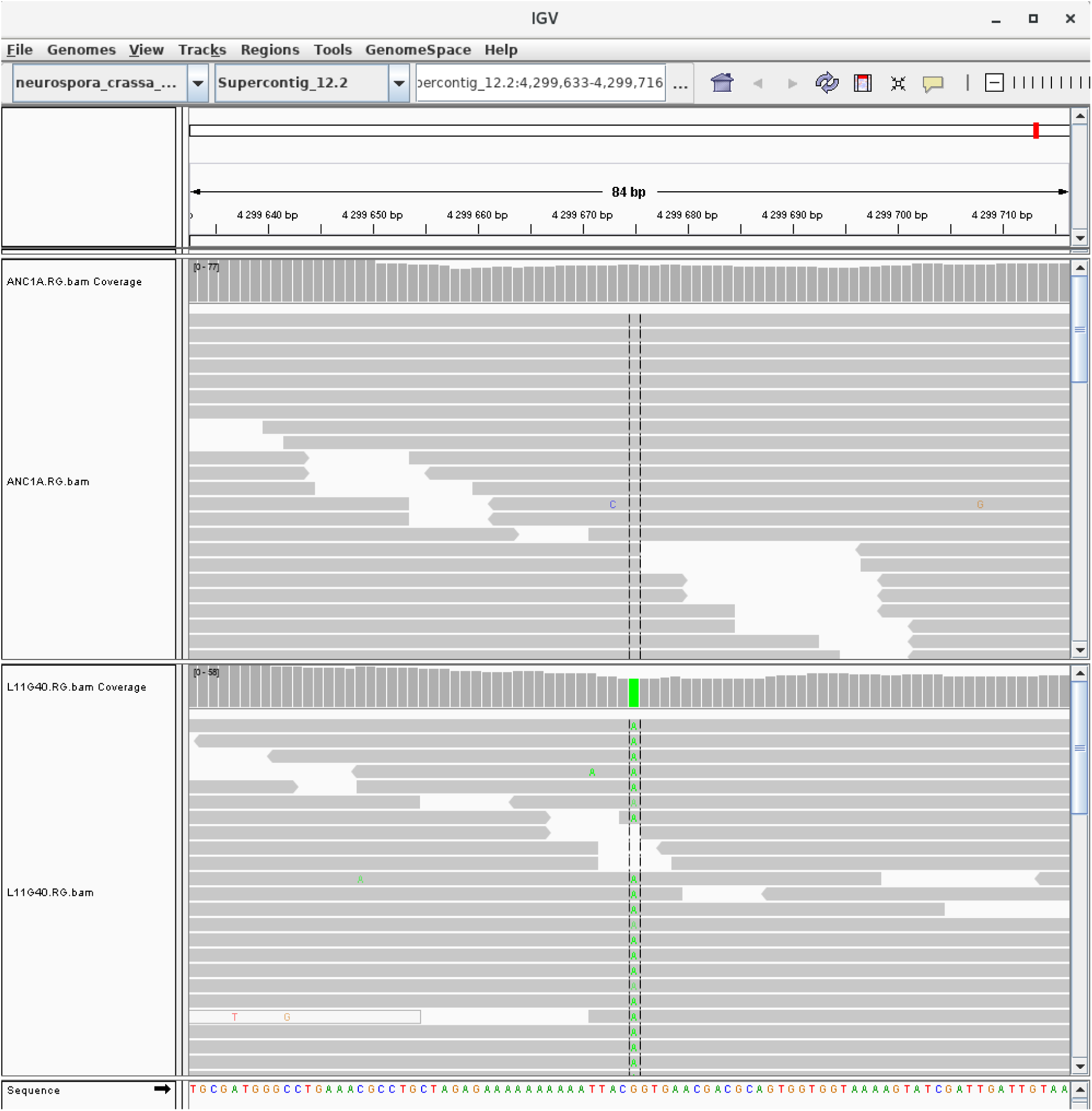
Screenshot of a mutation viewed in IGV. Upper track is the ancestor and lower track is MA line 11. Mutation is in chromosome 2, position 4 299 675, in euchromatin. Genotype quality of the mutation is 99.

**Figure S15:**
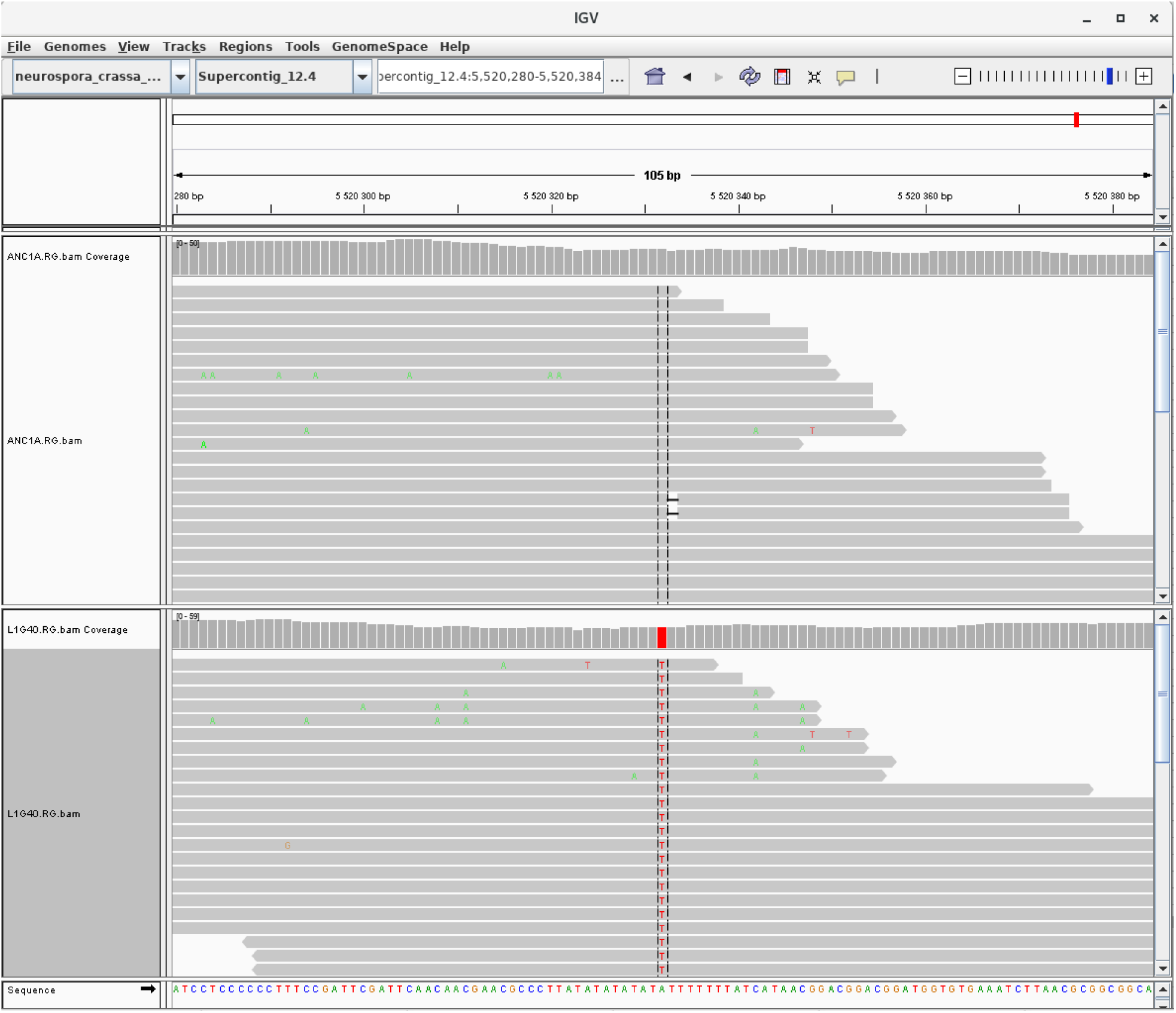
Screenshot of a mutation viewed in IGV. Upper track is the ancestor and lower track is MA line 1. Mutation is in chromosome 4, position 5 520 332, in euchromatin. Genotype quality of the mutation is 75.

**Figure S16:**
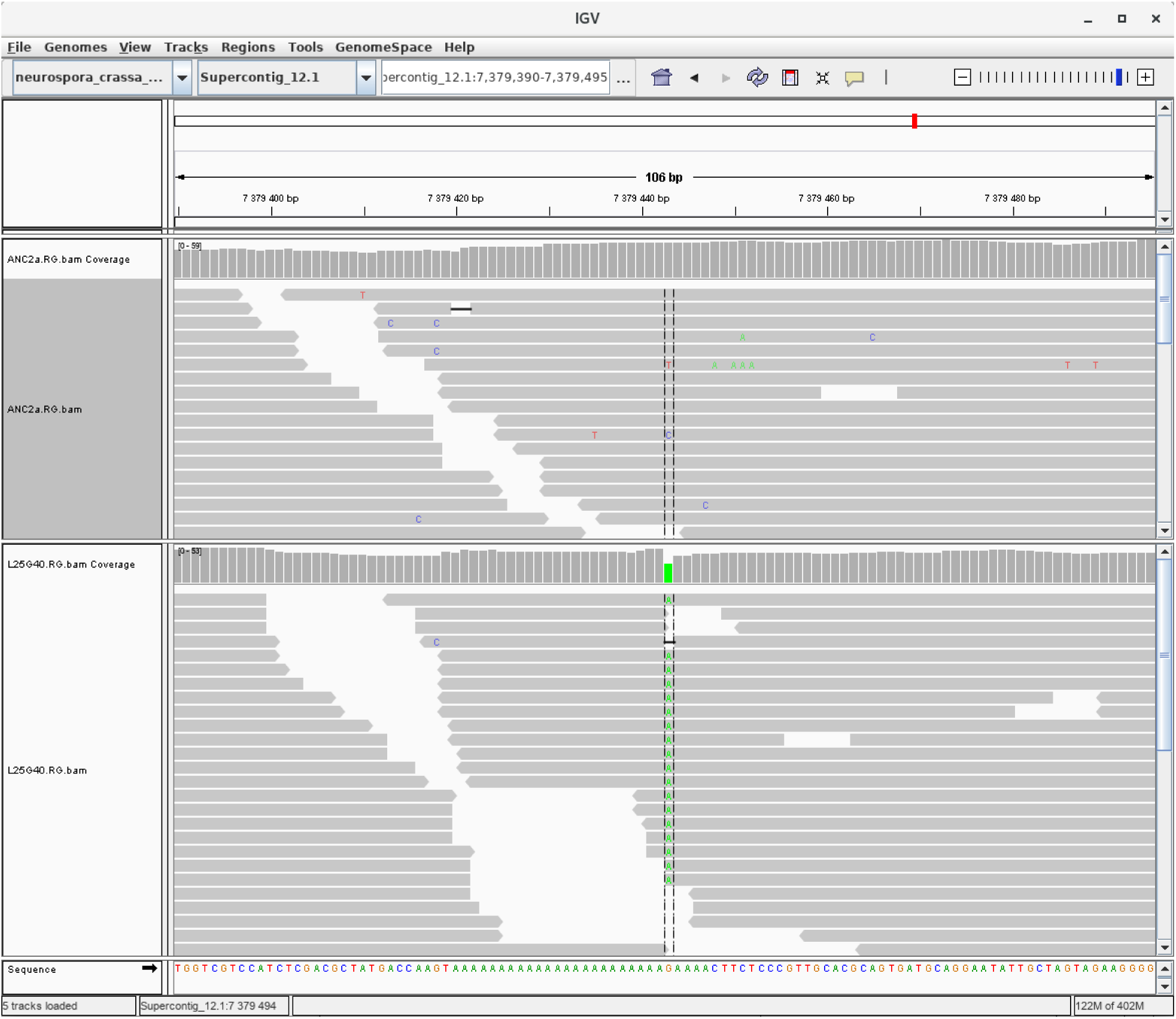
Screenshot of a mutation viewed in IGV. Upper track is the ancestor and lower track is MA line 25. Mutation is in chromosome 1, position 7 379 443, in euchromatin. Genotype quality of the mutation is 45 as the mutation is located in a repetitive region. This mutation was confirmed by Sanger sequencing.

**Figure S17:**
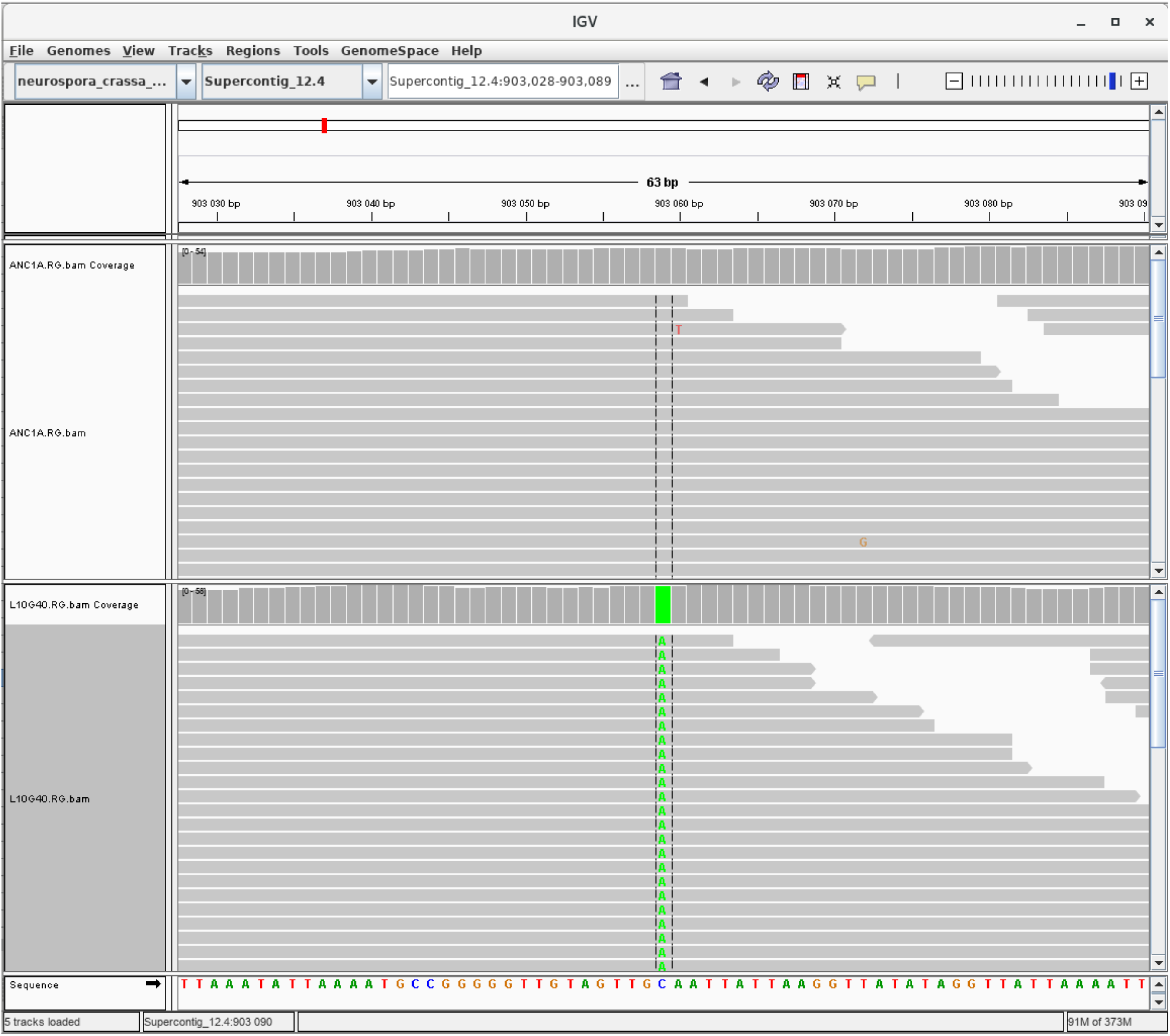
Screenshot of a mutation viewed in IGV. Upper track is the ancestor and lower track is MA line 10. Mutation is in chromosome 4, position 903 059, in centromeric region. Genotype quality of the mutation is 99.

**Figure S18:**
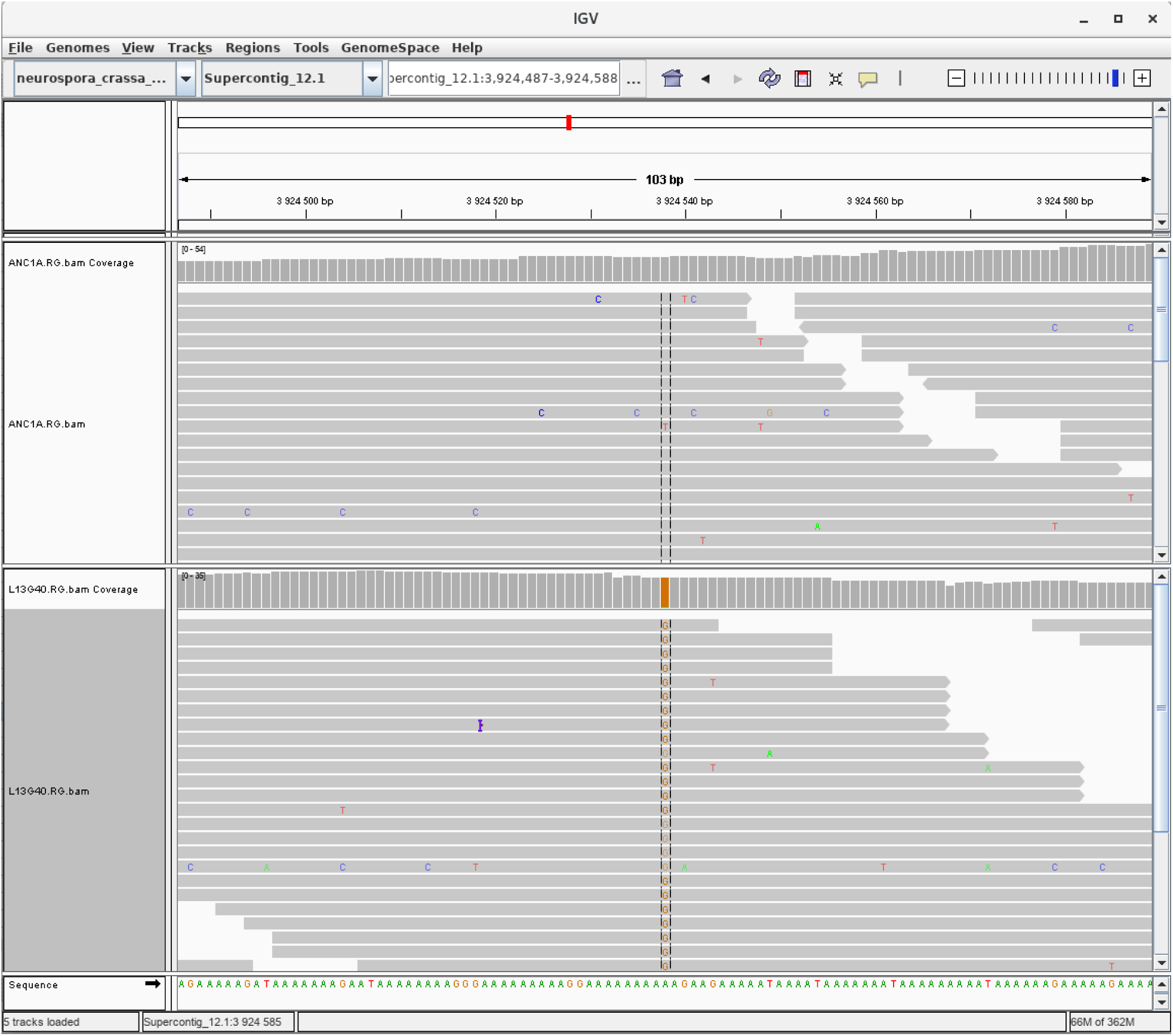
Screenshot of a mutation viewed in IGV. Upper track is the ancestor and lower track is MA line 13. Mutation is in chromosome 1, position 3 924 538, in centromeric region. Genotype quality of the mutation is 67 as the mutation is located in a repetitive region.

**Figure S19:**
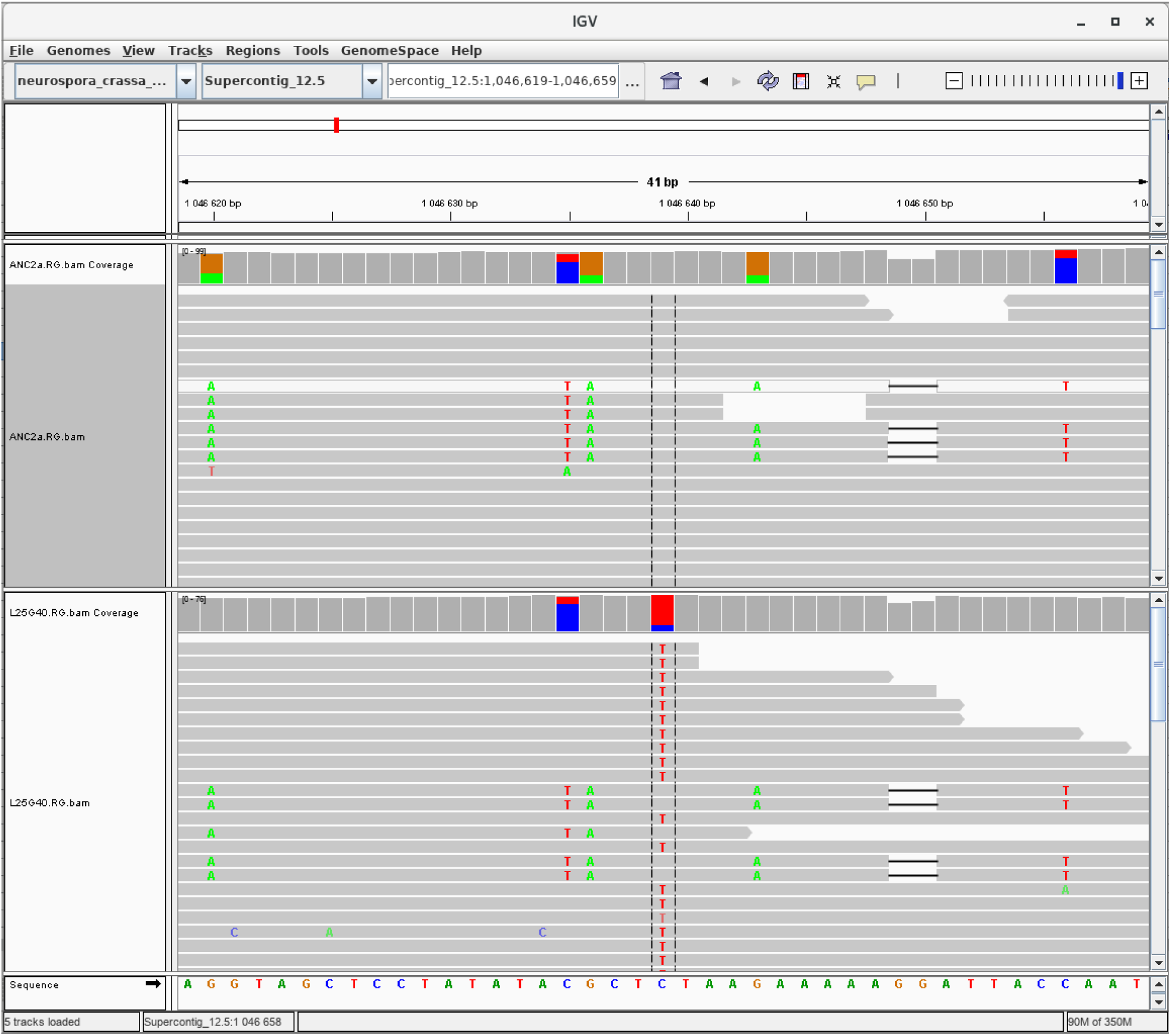
Screenshot of a mutation viewed in IGV. Upper track is the ancestor and lower track is MA line 25. Mutation is in chromosome 5, position 1 046 639, in centromeric region. Genotype quality of the mutation is 45 as the mutation is located in region with reduced mapping quality. Some reads that do not support the mutation map to this location. However, those reads also have other changes that are not supported by other reads. This suggest that reads not supporting the mutation are mapping errors. This mutation was confirmed by Sanger sequencing.

**Figure S20:**
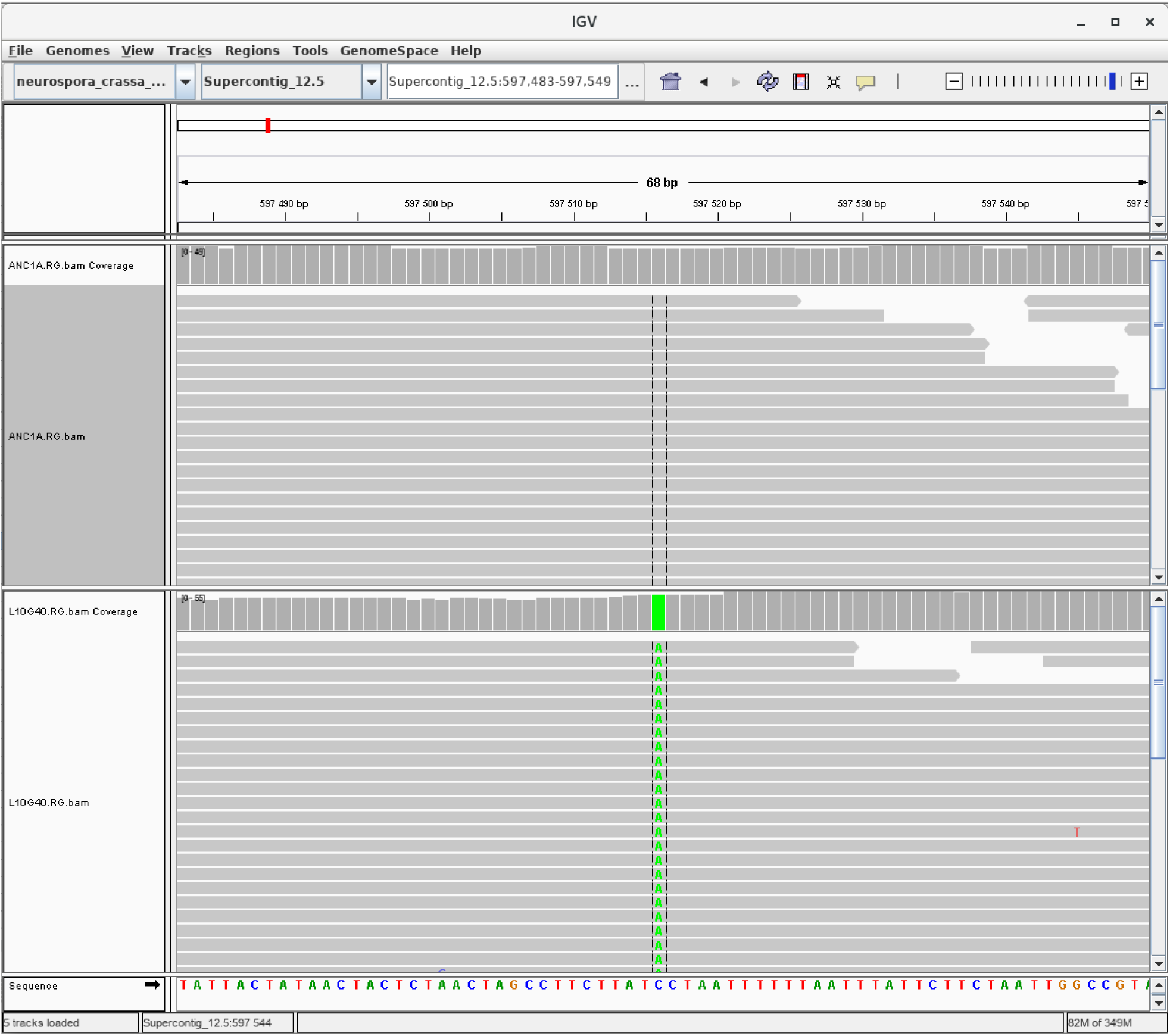
Screenshot of a mutation viewed in IGV. Upper track is the ancestor and lower track is MA line 10. Mutation is in chromosome 5, position 597 516, in region marked by H3K9 methylation. Genotype quality is 99.

**Figure S21:**
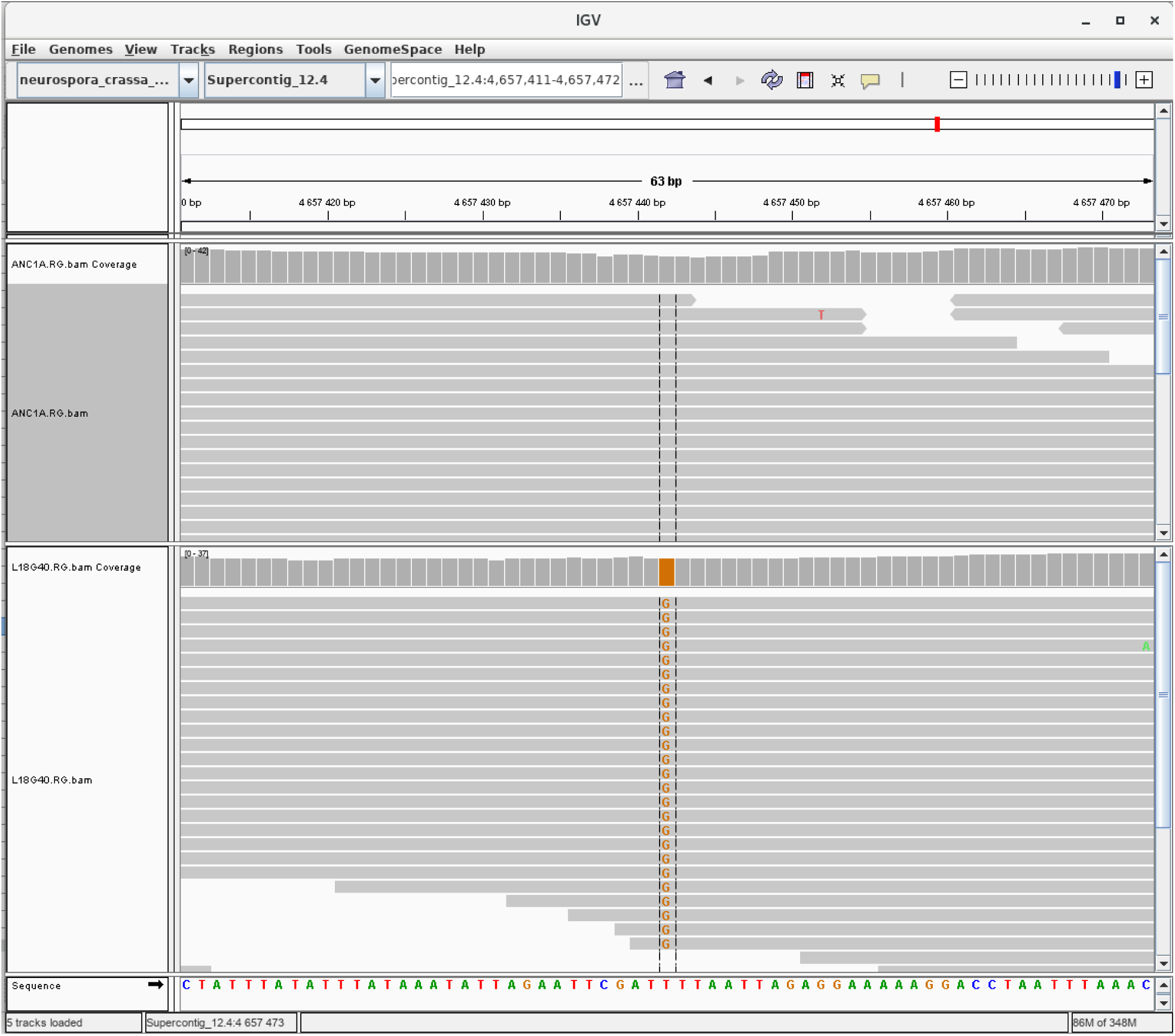
Screenshot of a mutation viewed in IGV. Upper track is the ancestor and lower track is MA line 18. Mutation is in chromosome 4, position 5 657 442, in region marked by H3K9 methylation. Genotype quality is 72.

**Figure S22:**
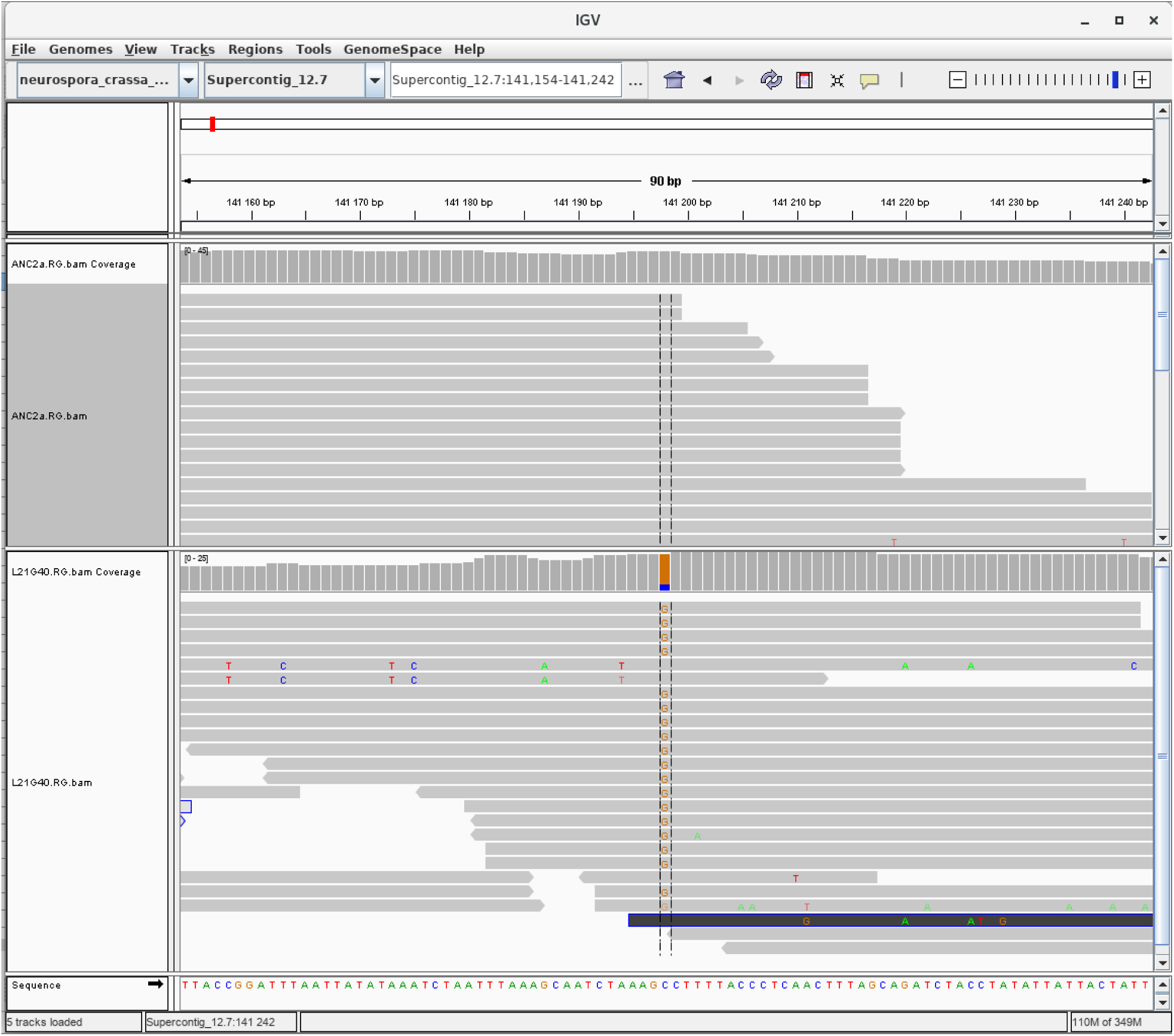
Screenshot of a mutation viewed in IGV. Upper track is the ancestor and lower track is MA line 21. Mutation is in chromosome 7, position 141 198, in region marked by H3K9 methylation. Genotype quality is 45. Some reads do not support the mutation. However, those reads have other changes that suggest a read mapping error. This mutation was confirmed by Sanger sequencing.

**Figure S23:**
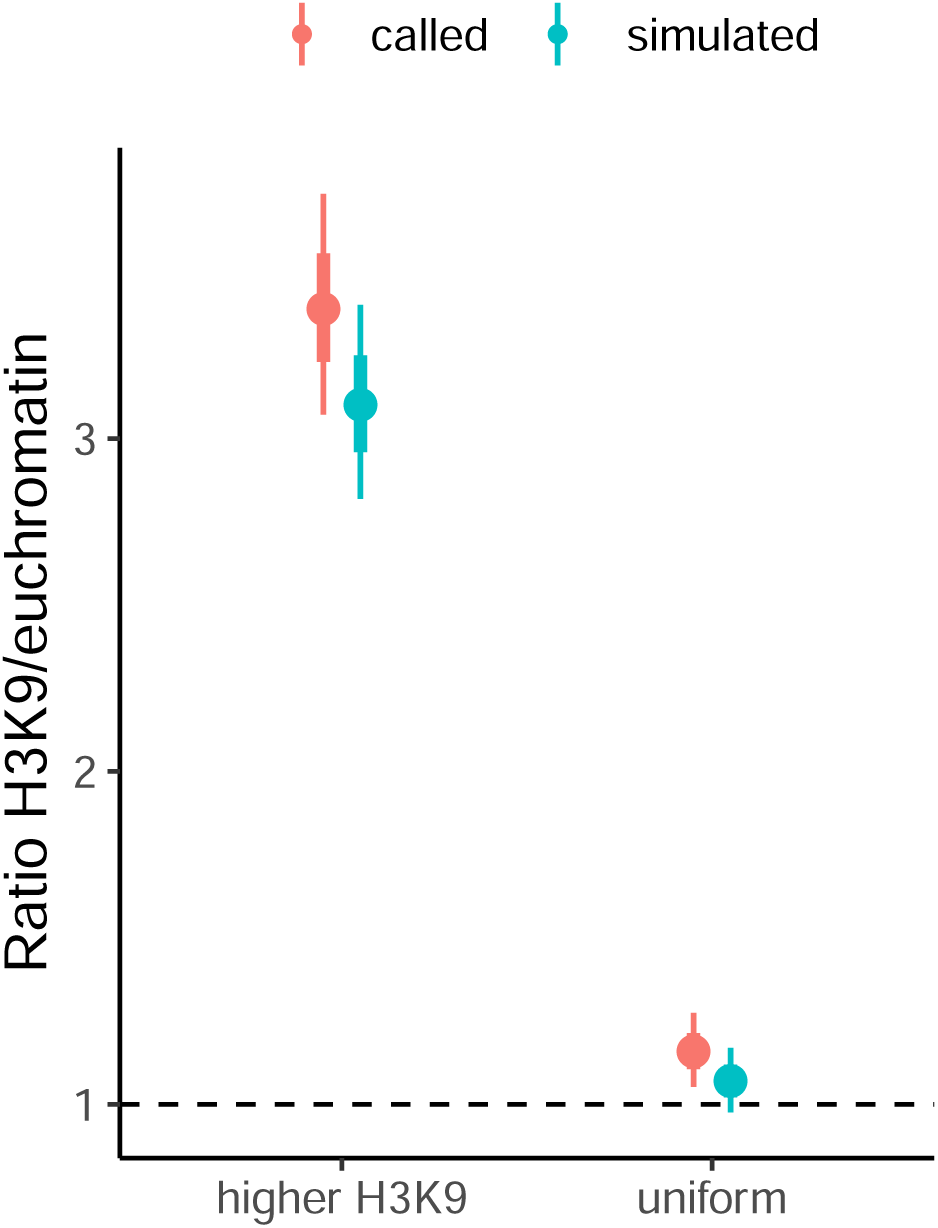
H3K9me3 / euchromatin mutation rate ratio in simulated data. Estimates calculated from called mutations from the simulated reads are in red, and estimates calculated using the true simulated mutations are in blue. The two different scenarios are: mutation rate was higher in regions of the genome marked by H3K9me3, and mutation rate was uniform across the genome. Points are means and range shows the 95% HPD interval of the ratio. Interval estimates overlap in both scenarios.

## Supplementary Tables

**Table S1:** Summary of alignment metrics for genomes used in this study. The ancestors used to start the MA experiment were: B 26708, which is 2489 *mat A*, and B 26709, which is 2489 *mat a*. Lines L1–L20 are *mat A* and L21–L40 are *mat a*.

| Natural strains |  |  |  | MA lines |  |  |  |
| --- | --- | --- | --- | --- | --- | --- | --- |
| Line | Number of reads | Depth | Mapped reads (%) | Line | Number of reads | Depth | Mapped reads (%) |
| 10882 | 30683221 | 112 | 87.2 | B 26708 | 15183018 | 55 | 98.5 |
| 10883 | 20920000 | 76 | 69.7 | B 26709 | 18679696 | 68 | 98.3 |
| 10884 | 32294020 | 118 | 72.8 | L1 | 13215012 | 48 | 98.2 |
| 10886 | 15265173 | 56 | 92.7 | L2 | 15658340 | 57 | 98.2 |
| 10892 | 32124218 | 117 | 62.3 | L3 | 14699884 | 54 | 98.3 |
| 10904 | 14718008 | 54 | 91.1 | L4 | 16424902 | 60 | 98.2 |
| 10906 | 14935778 | 55 | 89.7 | L5 | 16328002 | 60 | 97.6 |
| 10907 | 32213115 | 118 | 68.1 | L6 | 15189850 | 55 | 98.3 |
| 10908 | 13924689 | 51 | 91.7 | L7 | 14006386 | 51 | 98.4 |
| 10912 | 14829204 | 54 | 92 | L8 | 13888092 | 51 | 98 |
| 10914 | 30158241 | 110 | 61.1 | L9 | 14386986 | 53 | 98.3 |
| 10915 | 24410441 | 89 | 61.1 | L10 | 15458492 | 56 | 98.6 |
| 10918 | 40163515 | 147 | 65.4 | L11 | 13884490 | 51 | 98.1 |
| 10923 | 16231892 | 59 | 92 | L12 | 15913482 | 58 | 98.2 |
| 10925 | 37031022 | 135 | 85.2 | L13 | 20192750 | 74 | 97.8 |
| 10926 | 32682749 | 119 | 66 | L14 | 17445964 | 64 | 98.1 |
| 10927 | 17741573 | 65 | 73.1 | L15 | 14776588 | 54 | 98.5 |
| 10928 | 15048163 | 55 | 92 | L16 | 14562124 | 53 | 98 |
| 10932 | 13402421 | 49 | 91.1 | L17 | 16043040 | 59 | 97.9 |
| 10935 | 22304835 | 81 | 81.1 | L18 | 14755826 | 54 | 97.7 |
| 10937 | 29757556 | 109 | 85.4 | L19 | 13712542 | 50 | 98.1 |
| 10943 | 32838278 | 120 | 72.8 | L20 | 18685746 | 68 | 96.9 |
| 10946 | 17261401 | 63 | 90.1 | L21 | 14814794 | 54 | 98.5 |
| 10948 | 13835447 | 50 | 91 | L22 | 15849832 | 58 | 98 |
| 10950 | 14364201 | 52 | 98 | L23 | 17528602 | 64 | 98.3 |
| 10951 | 13354998 | 49 | 89.5 | L24 | 14415798 | 53 | 97.8 |
| 10983 | 38349194 | 140 | 72.8 | L25 | 14773870 | 54 | 98.1 |
| 1131 | 16974127 | 62 | 88.7 | L26 | 17436754 | 64 | 98.2 |
| 1133 | 14540257 | 53 | 88.3 | L27 | 14047860 | 51 | 98.2 |
| 1165 | 15095250 | 55 | 90 | L28 | 16295790 | 59 | 98.3 |
| 3210 | 13681263 | 50 | 90.4 | L29 | 16244750 | 59 | 98.1 |
| 3211 | 13908534 | 51 | 89.4 | L30 | 19847786 | 72 | 98.1 |
| 3223 | 14095397 | 51 | 92.1 | L31 | 19613222 | 72 | 97.9 |
| 3943 | 143993668 | 525 | 82.6 | L33 | 33430658 | 122 | 97.9 |
| 3975 | 13671543 | 50 | 88.8 | L34 | 15456254 | 56 | 97.9 |
| 4708 | 14310459 | 52 | 87.7 | L35 | 14405808 | 53 | 98 |
| 4712 | 12602174 | 46 | 86 | L36 | 15567642 | 57 | 97.8 |
| 4716 | 18336337 | 67 | 86.7 | L37 | 15232240 | 56 | 98 |
| 4730 | 15588817 | 57 | 87.4 | L38 | 14265830 | 52 | 98 |
| 4824 | 12921275 | 47 | 88.3 | L39 | 16540462 | 60 | 98.1 |
| 5910 | 12494976 | 46 | 83.4 | L40 | 18856392 | 69 | 98.3 |
| 6203 | 13266522 | 48 | 89.3 |  |  |  |  |
| 851 | 12956372 | 47 | 89.7 |  |  |  |  |
| 8783 | 15381794 | 56 | 88 |  |  |  |  |
| 8790 | 13205097 | 48 | 90.1 |  |  |  |  |
| 8816 | 15452652 | 56 | 89.6 |  |  |  |  |
| 8819 | 13736103 | 50 | 87.5 |  |  |  |  |
| 8845 | 15868843 | 58 | 86.9 |  |  |  |  |
| 8850 | 12695261 | 46 | 89.3 |  |  |  |  |
| P4452 | 172179164 | 628 | 86.9 |  |  |  |  |
| P4463 | 21708426 | 79 | 69.7 |  |  |  |  |
| P4468 | 21129155 | 77 | 68.4 |  |  |  |  |
| P4471 | 24828717 | 91 | 72.4 |  |  |  |  |
| P4476 | 92452134 | 337 | 86.8 |  |  |  |  |
| P4479 | 22220857 | 81 | 72.5 |  |  |  |  |
| P4489 | 34624044 | 126 | 72.5 |  |  |  |  |

**Table S2:** Model comparisons among different models that predict the mutation rate by GC-content and chromatin modifications. Model terms are different linear model parts, *α* is the intercept, *β_GC_* is the slope effect of GC-content, *β_K9_* is the effect of H3K9 domain, *β_K27_* is the effect of H3K27 domain, *β_C_* is the effect of centromeric domain, *β_I_* is the interaction effect between GC-content and H3K9 domain, *β_I2_* is the interaction effect between GC-content and centromeric domain, *β_I3_* is the interaction effect between GC-content and H3K27 domain. *d_i_*, *g_i_*, and *c_i_* are indicator variables, and *x_i_* is GC-content in percentage points. WAIC = widely applicable information criterion, SE = standard error.

| Model terms | WAIC | diff (± SE) | weight |
| --- | --- | --- | --- |
| $\alpha + \beta_{GC}x_i + \beta_{K9}d_i + \beta_{K27}g_i + \beta_Cc_i + \beta_Ix_id_i$ | 454.47 | 0 (0) | 0.63 |
| $\alpha + \beta_{GC}x_i + \beta_{K9}d_i + \beta_{K27}g_i + \beta_Cc_i + \beta_Ix_id_i + \beta_{I3}x_ig_i$ | 456.86 | 2.39 (2.19) | 0.19 |
| $\alpha + \beta_{GC}x_i + \beta_{K9}d_i + \beta_Cc_i + \beta_Ix_id_i$ | 458.67 | 4.2 (6.94) | 0 |
| $\alpha + \beta_{GC}x_i + \beta_{K9}d_i$ | 458.84 | 4.37 (10.59) | 0 |
| $\alpha + \beta_{GC}x_i + \beta_{K9}d_i + \beta_Cc_i + \beta_Ix_id_i + \beta_{I2}x_ic_i$ | 460.35 | 5.88 (7.17) | 0 |
| $\alpha + \beta_{GC}x_i + \beta_{K9}d_i + \beta_Ix_id_i$ | 495.86 | 41.39 (19.86) | 0 |
| $\alpha + \beta_{GC}x_i + \beta_{K9}d_i$ | 496.65 | 42.18 (22.43) | 0 |
| $\alpha + \beta_{GC}x_i$ | 546.62 | 92.15 (33.84) | 0 |
| $\alpha + \beta_{K9}d_i + \beta_Cc_i$ | 614.83 | 160.36 (42.53) | 0 |
| $\alpha + \beta_{K9}d_i$ | 645.82 | 191.35 (49.19) | 0 |
| $\alpha + \beta_Cc_i$ | 1290.02 | 835.55 (255.52) | 0 |
| $\alpha$ | 1689.56 | 1235.09 (264.15) | 0 |

**Table S3:** Model estimates for a model predicting mutation rate by GC-content, centromeric, H3K9, and H3K27 domains, **α** is the intercept, *β_GC_* is the slope effect of GC-content, *β_K_*_9_ is the effect of H3K9me domain, *β_K_*_27_ is the effect of the H3K27me3 domain, *β_C_* is the effect of centromeric domain, and *β_I_* is the interaction effect between GC-content and H3K9me domain.

| Parameter | Estimate [95% HPDI] |
| --- | --- |
| $\alpha$ | -2.61 [-3.34, -1.88] |
| $\beta_C$ | 0.51 [0.35, 0.67] |
| $\beta_{K9}$ | -0.14 [-0.93, 0.63] |
| $\beta_{K27}$ | 0.32 [0.08, 0.55] |
| $\beta_{GC}$ | -0.06 [-0.08, -0.05] |
| $\beta_I$ | 0.02 [0.00, 0.04] |

**Table S4:**
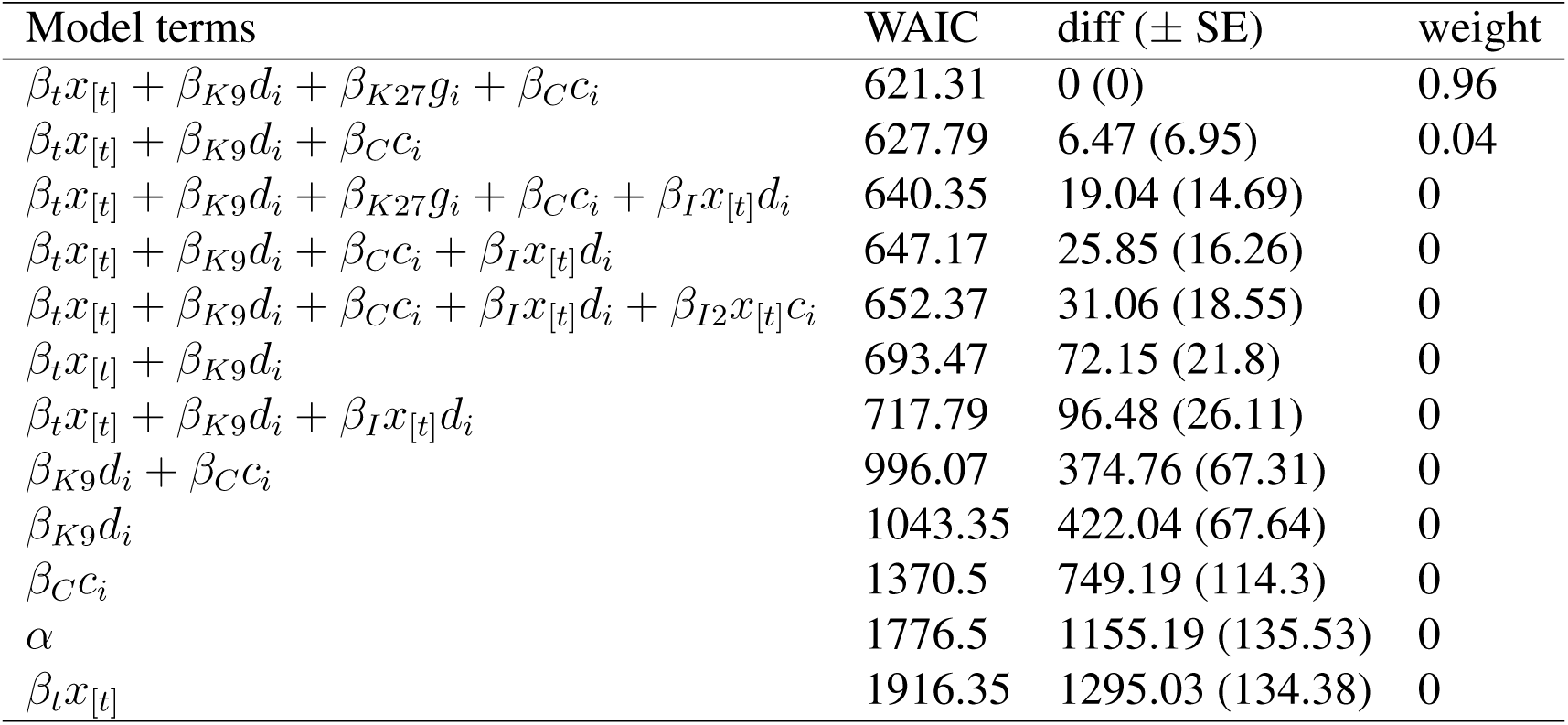
Model comparison among different models that predict the mutation rate by trinucleotide class and chromatin modifications. Model terms are different linear model parts. *α* is the intercept, *β_t_* is a vector of effects for the 32 trinucleotide classes, *β_K9_* is the effect of H3K9 domain, *β_K27_* is the effect of H3K27 domain, *β_C_* is the effect of centromeric domain, *β_I_* is the interaction effect between trinucleotide class and H3K9 domain, *β_I2_* is the interaction effect between trinucleotide class and centromeric domain. *d_i_*, *g_i_*, and *c_i_* are indicator variables, and *x*_[*t*]_ is the trinucleotide class. WAIC = widely applicable information criterion, SE = standard error.

| Model terms | WAIC | diff (± SE) | weight |
| --- | --- | --- | --- |
| $\beta_t x_{[t]} + \beta_{K9} d_i + \beta_{K27} g_i + \beta_C c_i$ | 621.31 | 0 (0) | 0.96 |
| $\beta_t x_{[t]} + \beta_{K9} d_i + \beta_C c_i$ | 627.79 | 6.47 (6.95) | 0.04 |
| $\beta_t x_{[t]} + \beta_{K9} d_i + \beta_{K27} g_i + \beta_C c_i + \beta_I x_{[t]} d_i$ | 640.35 | 19.04 (14.69) | 0 |
| $\beta_t x_{[t]} + \beta_{K9} d_i + \beta_C c_i + \beta_I x_{[t]} d_i$ | 647.17 | 25.85 (16.26) | 0 |
| $\beta_t x_{[t]} + \beta_{K9} d_i + \beta_C c_i + \beta_I x_{[t]} d_i + \beta_{I2} x_{[t]} c_i$ | 652.37 | 31.06 (18.55) | 0 |
| $\beta_t x_{[t]} + \beta_{K9} d_i$ | 693.47 | 72.15 (21.8) | 0 |
| $\beta_t x_{[t]} + \beta_{K9} d_i + \beta_I x_{[t]} d_i$ | 717.79 | 96.48 (26.11) | 0 |
| $\beta_{K9} d_i + \beta_C c_i$ | 996.07 | 374.76 (67.31) | 0 |
| $\beta_{K9} d_i$ | 1043.35 | 422.04 (67.64) | 0 |
| $\beta_C c_i$ | 1370.5 | 749.19 (114.3) | 0 |
| $\alpha$ | 1776.5 | 1155.19 (135.53) | 0 |
| $\beta_t x_{[t]}$ | 1916.35 | 1295.03 (134.38) | 0 |

**Table S5:**
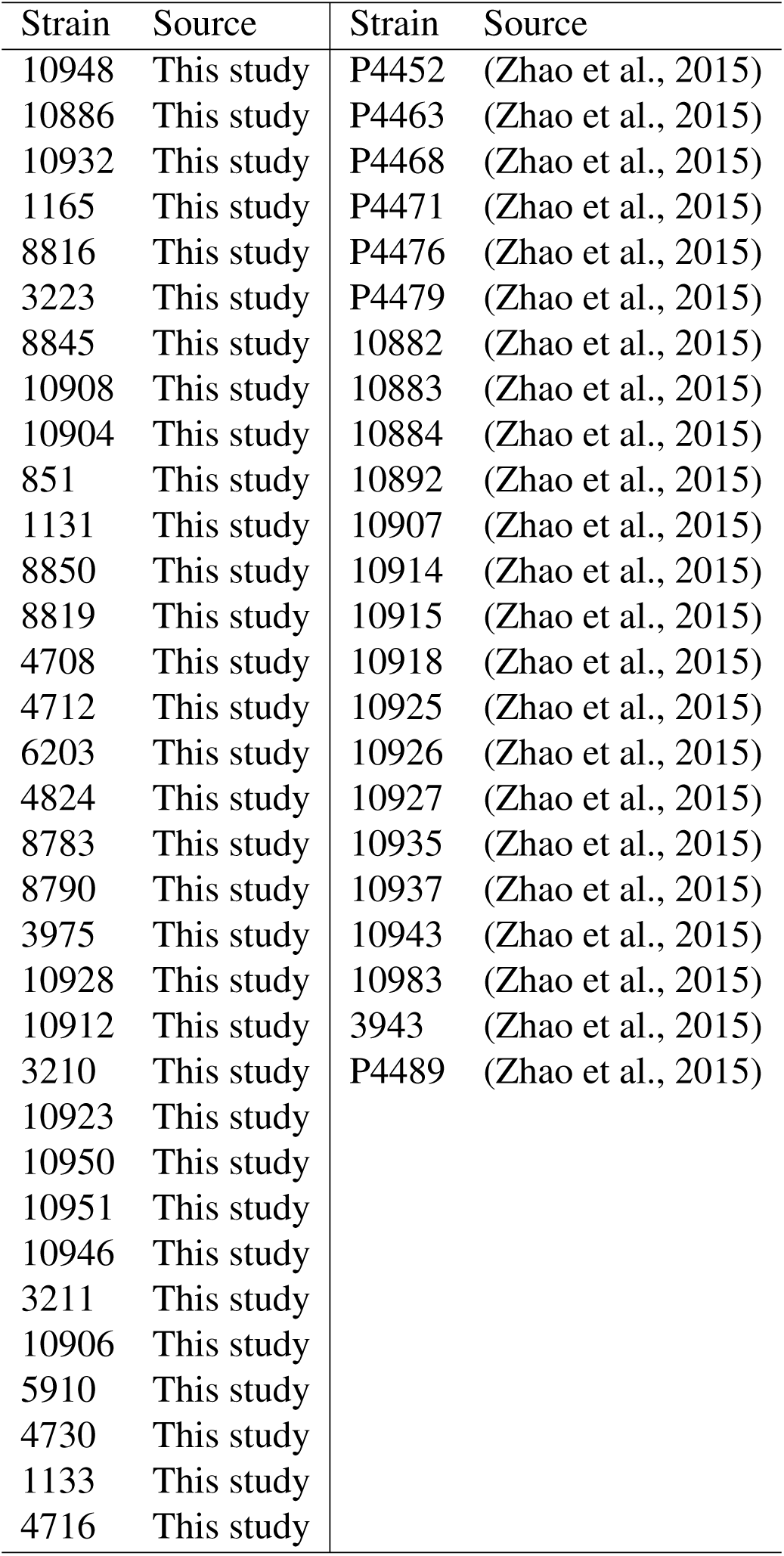
Natural strains with sequencing data included in this study. Strains were obtained from FGSC. 33 strains were sequenced in this study and data for 23 strains were obtained from Zhao et al. (2015).

| Strain | Source | Strain | Source |
| --- | --- | --- | --- |
| 10948 | This study | P4452 | (Zhao et al., 2015) |
| 10886 | This study | P4463 | (Zhao et al., 2015) |
| 10932 | This study | P4468 | (Zhao et al., 2015) |
| 1165 | This study | P4471 | (Zhao et al., 2015) |
| 8816 | This study | P4476 | (Zhao et al., 2015) |
| 3223 | This study | P4479 | (Zhao et al., 2015) |
| 8845 | This study | 10882 | (Zhao et al., 2015) |
| 10908 | This study | 10883 | (Zhao et al., 2015) |
| 10904 | This study | 10884 | (Zhao et al., 2015) |
| 851 | This study | 10892 | (Zhao et al., 2015) |
| 1131 | This study | 10907 | (Zhao et al., 2015) |
| 8850 | This study | 10914 | (Zhao et al., 2015) |
| 8819 | This study | 10915 | (Zhao et al., 2015) |
| 4708 | This study | 10918 | (Zhao et al., 2015) |
| 4712 | This study | 10925 | (Zhao et al., 2015) |
| 6203 | This study | 10926 | (Zhao et al., 2015) |
| 4824 | This study | 10927 | (Zhao et al., 2015) |
| 8783 | This study | 10935 | (Zhao et al., 2015) |
| 8790 | This study | 10937 | (Zhao et al., 2015) |
| 3975 | This study | 10943 | (Zhao et al., 2015) |
| 10928 | This study | 10983 | (Zhao et al., 2015) |
| 10912 | This study | 3943 | (Zhao et al., 2015) |
| 3210 | This study | P4489 | (Zhao et al., 2015) |
| 10923 | This study |  |  |
| 10950 | This study |  |  |
| 10951 | This study |  |  |
| 10946 | This study |  |  |
| 3211 | This study |  |  |
| 10906 | This study |  |  |
| 5910 | This study |  |  |
| 4730 | This study |  |  |
| 1133 | This study |  |  |
| 4716 | This study |  |  |

**Table S6:** Detecting structural variants with different callers from simulated data. Callers tested were Delly, Lumpy, SVaba, and Pindel. Different sets are simulations with different numbers of structural variants.

| set | Deletion | Duplication | Inversion | Translocation | Insertion | Inv-del | Total number of SV | Delly |  | Lumpy |  | SVaba |  | Pindel |  |
| --- | --- | --- | --- | --- | --- | --- | --- | --- | --- | --- | --- | --- | --- | --- | --- |
|  |  |  |  |  |  |  |  | sensitivity | FDR | sensitivity | FDR | sensitivity | FDR | sensitivity | FDR |
| 1 | 1 | 4 | 4 | 4 | 5 | 0 | 18 | 0.90 | 0 | 0.90 | 0.55 | 0.60 | 0.60 | 0.55 | 0.65 |
| 2 | 7 | 15 | 5 | 8 | 5 | 0 | 40 | 0.84 | 0.02 | 0.84 | 0.54 | 0.51 | 0.55 | 0.48 | 0.60 |
| 3 | 8 | 10 | 10 | 15 | 7 | 0 | 50 | 0.88 | 0 | 0.88 | 0.54 | 0.64 | 0.52 | 0.38 | 0.61 |
| 4 | 6 | 20 | 20 | 20 | 14 | 0 | 100 | 0.54 | 0.33 | 0.54 | 0.60 | 0.36 | 0.77 | 0.28 | 0.94 |

**Table S7:** Calling CNVs on simulated data using either CNVnator with two different bin sizes, CNV-seq, or both callers together.

|  | Sensitivity score | FDR score |
| --- | --- | --- |
| CNVnator (1670 bin size) | 0.375 | 0.556 |
| CNVnator (75 bin size) | 0.968 | 0.797 |
| CNV-seq | 0.937 | 0.999 |
| Both callers | 0.906 | 0.482 |

## References

Beier, S., Thiel, T., Münch, T., Scholz, U., and Mascher, M., 2017. MISA-web: a web server for microsatellite prediction. Bioinformatics, 33(16):2583–2585.

Besenbacher, S., Sulem, P., Helgason, A., Helgason, H., Kristjansson, H., Jonasdottir, A., Jonasdottir, A., Magnusson, O. T., Thorsteinsdottir, U., Masson, G., et al., 2016. Multi-nucleotide *de novo* mutations in humans. PLOS Genetics, 12(11):1–15.

Bicocca, V. T., Ormsby, T., Adhvaryu, K. K., Honda, S., and Selker, E. U., 2018. ASH1-catalyzed H3K36 methylation drives gene repression and marks H3K27me2/3-competent chromatin. eLife, 7:e41497.

Bürkner, P.-C., 2017. brms: An R package for Bayesian multilevel models using Stan. Journal of Statistical Software, 80(1):1–28.

Cambareri, E. B., Singer, M. J., and Selker, E. U., 1991. Recurrence of repeat-induced point mutation (RIP) in *Neurospora crassa*. Genetics, 127(4):699–710.

Campos, J. L., Zhao, L., and Charlesworth, B., 2017. Estimating the parameters of background selection and selective sweeps in *Drosophila* in the presence of gene conversion. Proceedings of the National Academy of Sciences, 114(24):E4762–E4771.

Carpenter, B., Gelman, A., Hoffman, M., Lee, D., Goodrich, B., Betancourt, M., Brubaker, M., Guo, J., Li, P., and Riddell, A., et al., 2017. Stan: A probabilistic programming language. Journal of Statistical Software, 76(1):1–32.

Castellano, D., Eyre-Walker, A., and Munch, K., 2020. Impact of Mutation Rate and Selection at Linked Sites on DNA Variation across the Genomes of Humans and Other Homininae. Genome Biology and Evolution, 12(1):3550–3561.

Chen, X., Chen, Z., Chen, H., Su, Z., Yang, J., Lin, F., Shi, S., and He, X., 2012. Nucleosomes suppress spontaneous mutations base-specifically in eukaryotes. Science, 335(6073):1235–1238.

Cheng, K. C., Cahill, D. S., Kasai, H., Nishimura, S., and Loeb, L. A., 1992. 8-Hydroxyguanine, an abundant form of oxidative DNA damage, causes G-T and A-C substitutions. Journal of Biological Chemistry, 267(1):166–172.

Cooper, D. N., Mort, M., Stenson, P. D., Ball, E. V., and Chuzhanova, N. A., 2010. Methylation-mediated deamination of 5-methylcytosine appears to give rise to mutations causing human in-herited disease in CpNpG trinucleotides, as well as in CpG dinucleotides. Human Genomics, 4(6):406.

Denver, D. R., Morris, K., Kewalramani, A., Harris, K. E., Chow, A., Estes, S., Lynch, M., and Thomas, W. K., 2004. Abundance, distribution, and mutation rates of homopolymeric nucleotide runs in the genome of *Caenorhabditis elegans*. Journal of Molecular Evolution, 58(5):584–595.

Drake, J. W., 1991. A constant rate of spontaneous mutation in DNA-based microbes. Proceedings of the National Academy of Sciences, 88(16):7160–7164.

Eyre-Walker, A. and Keightley, P. D., 2007. The distribution of fitness effects of new mutations. Nature Reviews Genetics, 8(8):610–618.

Ferretti, L., Raineri, E., and Ramos-Onsins, S., 2012. Neutrality tests for sequences with missing data. Genetics, 191(4):1397–1401.

Habig, M., Lorrain, C., Feurtey, A., Komluski, J., and Stukenbrock, E. H., 2021. Epigenetic modifications affect the rate of spontaneous mutations in a pathogenic fungus. Nature Communications, 12(1):5869.

Halligan, D. L. and Keightley, P. D., 2009. Spontaneous mutation accumulation studies in evolutionary genetics. Annual Review of Ecology, Evolution, and Systematics, 40(1):151–172.

Hanrahan, C. J., Bacolod, M. D., Vyas, R. R., Liu, T., Geacintov, N. E., Loechler, E. L., and Basu, A. K., 1997. Sequence specific mutagenesis of the major (+)-anti-benzo[a]pyrene diol epoxide– DNA adduct at a mutational hot spot *in vitro* and in *Escherichia coli* cells. Chem. Res. Toxicol., 10(4):369–377.

Harris, K. and Nielsen, R., 2014. Error-prone polymerase activity causes multinucleotide mutations in humans. Genome Research, 24(9):1445–1454.

Homer, N., 2021. DWGSIM: Whole genome simulator for next-generation sequencing. github repository.

Hosseini, S., Meunier, C., Nguyen, D., Reimegård, J., and Johannesson, H., 2020. Comparative analysis of genome-wide DNA methylation in *Neurospora*. Epigenetics, 15(9):972–987. PMID: 32228351.

Jamieson, K., Rountree, M. R., Lewis, Z. A., Stajich, J. E., and Selker, E. U., 2013. Regional control of histone H3 lysine 27 methylation in *Neurospora*. Proceedings of the National Academy of Sciences, 110(15):6027–6032.

Jeffares, D. C., Jolly, C., Hoti, M., Speed, D., Shaw, L., Rallis, C., Balloux, F., Dessimoz, C., Bähler, J., and Sedlazeck, F. J., et al., 2017. Transient structural variations have strong effects on quantitative traits and reproductive isolation in fission yeast. Nature Communications, 8:14061.

Johri, P., Charlesworth, B., and Jensen, J. D., 2020. Toward an evolutionarily appropriate null model: Jointly inferring demography and purifying selection. Genetics, 215(1):173–192.

Kadyrova, L. Y., Mertz, T. M., Zhang, Y., Northam, M. R., Sheng, Z., Lobachev, K. S., Shcherbakova, P. V., and Kadyrov, F. A., 2013. A reversible histone H3 acetylation cooperates with mismatch repair and replicative polymerases in maintaining genome stability. PLOS Genetics, 9(10):1–16.

Katju, V. and Bergthorsson, U., 2018. Old trade, new tricks: Insights into the spontaneous mutation process from the partnering of classical mutation accumulation experiments with high-throughput genomic approaches. Genome Biology and Evolution, 11(1):136–165.

Keightley, P. D. and Eyre-Walker, A., 2007. Joint inference of the distribution of fitness effects of deleterious mutations and population demography based on nucleotide polymorphism frequencies. Genetics, 177(4):2251–2261.

Keightley, P. D., Ness, R. W., Halligan, D. L., and Haddrill, P. R., 2014. Estimation of the spontaneous mutation rate per nucleotide site in a *Drosophila melanogaster* full-sib family. Genetics, 196(1):313–320.

Keightley, P. D., Pinharanda, A., Ness, R. W., Simpson, F., Dasmahapatra, K. K., Mallet, J., Davey, J. W., and Jiggins, C. D., 2015. Estimation of the spontaneous mutation rate in *Heliconius melpomene*. Molecular Biology and Evolution, 32(1):239–243.

Keith, N., Tucker, A. E., Jackson, C. E., Sung, W., Lucas Lledó, J. I., Schrider, D. R., Schaack, S., Dudycha, J. L., Ackerman, M., Younge, A. J., et al., 2016. High mutational rates of large-scale duplication and deletion in *Daphnia pulex*. Genome Research, 26(1):60–69.

Kelleher, J., Ness, R. W., and Halligan, D. L., 2013. Processing genome scale tabular data with wormtable. BMC Bioinformatics, 14(1):1–5.

Kouzarides, T., 2007. Chromatin modifications and their function. Cell, 128(4):693 – 705.

Kronholm, I., Bassett, A., Baulcombe, D., and Collins, S., 2017. Epigenetic and genetic contributions to adaptation in *Chlamydomonas*. Molecular Biology and Evolution, 34(9):2285–2306.

Law, Y. K., Forties, R. A., Liu, X., Poirier, M. G., and Kohler, B., 2013. Sequence-dependent thymine dimer formation and photoreversal rates in double-stranded DNA. Photochem. Photobiol. Sci., 12:1431–1439.

Layer, R. M., Chiang, C., Quinlan, A. R., and Hall, I. M., 2014. LUMPY: a probabilistic framework for structural variant discovery. Genome Biology, 15(6):R84.

Leushkin, E. V., Bazykin, G. A., and Kondrashov, A. S., 2013. Strong mutational bias toward deletions in the *Drosophila melanogaster* genome is compensated by selection. Genome Biology and Evolution, 5(3):514–524.

Li, C. and Luscombe, N. M., 2020. Nucleosome positioning stability is a modulator of germline mutation rate variation across the human genome. Nature Communications, 11(1):1363.

Li, H., 2013. Aligning sequence reads, clone sequences and assembly contigs with BWA-MEM. arXiv:1303.3997 [q-bio.GN],.

Li, H., 2014. Toward better understanding of artifacts in variant calling from high-coverage samples. Bioinformatics, 30(20):2843–2851.

Lynch, M., Ackerman, M. S., Gout, J.-F., Long, H., Sung, W., Thomas, W. K., and Foster, P. L., 2016. Genetic drift, selection and the evolution of the mutation rate. Nat Rev Genet, 17(11):704– 714.

Makova, K. D. and Hardison, R. C., 2015. The effects of chromatin organization on variation in mutation rates in the genome. Nature Reviews Genetics, 16(4):213–223.

McBride, T. J., Preston, B. D., and Loeb, L. A., 1991. Mutagenic spectrum resulting from DNA damage by oxygen radicals. Biochemistry, 30(1):207–213.

McCluskey, K., Wiest, A., and Plamann, M., 2010. The fungal genetics stock center: a repository for 50 years of fungal genetics research. J Biosci, 35(1):119–126.

McKenna, A., Hanna, M., Banks, E., Sivachenko, A., Cibulskis, K., Kernytsky, A., Garimella, K., Altshuler, D., Gabriel, S., Daly, M., et al., 2010. The Genome Analysis Toolkit: a MapReduce framework for analyzing next-generation DNA sequencing data. Genome Research, 20(9):1297– 1303.

Monroe, J. G., Srikant, T., Carbonell-Bejerano, P., Becker, C., Lensink, M., Exposito-Alonso, M., Klein, M., Hildebrandt, J., Neumann, M., Kliebenstein, D., et al., 2022. Mutation bias reflects natural selection in *Arabidopsis thaliana*. Nature, 602:101–105.

Ness, R. W., Morgan, A. D., Colegrave, N., and Keightley, P. D., 2012. Estimate of the spontaneous mutation rate in *Chlamydomonas reinhardtii*. Genetics, 192:1447–1454.

Ness, R. W., Morgan, A. D., Vasanthakrishnan, R. B., Colegrave, N., and Keightley, P. D., 2015. Extensive de novo mutation rate variation between individuals and across the genome of *Chlamydomonas reinhardtii*. Genome Research, 25:1739–1749.

Ossowski, S., Schneeberger, K., Lucas-Lledó, J. I., Warthmann, N., Clark, R. M., Shaw, R. G., Weigel, D., and Lynch, M., 2010. The rate and molecular spectrum of spontaneous mutations in *Arabidopsis thaliana*. Science, 327(5961):92–94.

Polak, P., Karlić, R., Koren, A., Thurman, R., Sandstrom, R., Lawrence, M. S., Reynolds, A., Rynes, E., Vlahoviček, K., Stamatoyannopoulos, J. A., et al., 2015. Cell-of-origin chromatin organization shapes the mutational landscape of cancer. Nature, 518(7539):360–364.

R Core Team, 2019. R: A language and environment for statistical computing. R Foundation for Statistical Computing, Vienna, Austria.

Rausch, T., Zichner, T., Schlattl, A., Stütz, A. M., Benes, V., and Korbel, J. O., 2012. DELLY: structural variant discovery by integrated paired-end and split-read analysis. Bioinformatics, 28(18):i333–i339.

Sasaki, S., Mello, C. C., Shimada, A., Nakatani, Y., Hashimoto, S.-i., Ogawa, M., Matsushima, K., Gu, S. G., Kasahara, M., Ahsan, B., et al., 2009. Chromatin-associated periodicity in genetic variation downstream of transcriptional start sites. Science, 323(5912):401–404.

Schuster-Böckler, B. and Lehner, B., 2012. Chromatin organization is a major influence on regional mutation rates in human cancer cells. Nature, 488(7412):504–507.

Selker, E. U., 1990. Premeiotic instability of repeated sequences in *Neurospora crassa*. Annual Review of Genetics, 24(1):579–613. PMID: 2150906.

Smith, K. M., Phatale, P. A., Sullivan, C. M., Pomraning, K. R., and Freitag, M., 2011. Heterochromatin is required for normal distribution of *Neurospora crassa* CenH3. Molecular and Cellular Biology, 31(12):2528–2542.

Stone, J. E., Lujan, S. A., and Kunkel, T. A., 2012. DNA polymerase zeta generates clustered mutations during bypass of endogenous DNA lesions in *Saccharomyces cerevisiae*. Environmental and Molecular Mutagenesis, 53(9):777–786.

Sung, W., Ackerman, M. S., Gout, J.-F., Miller, S. F., Williams, E., Foster, P. L., and Lynch, M., 2015. Asymmetric context-dependent mutation patterns revealed through mutation–accumulation experiments. Molecular Biology and Evolution, 32(7):1672–1683.

Supek, F. and Lehner, B., 2015. Differential DNA mismatch repair underlies mutation rate variation across the human genome. Nature, 521(7550):81–84.

Tamaru, H. and Selker, E. U., 2001. A histone H3 methyltransferase controls DNA methylation in *Neurospora crassa*. Nature, 414(6861):277–283.

Thorvaldsdóttir, H., Robinson, J. T., and Mesirov, J. P., 2013. Integrative Genomics Viewer (IGV): high-performance genomics data visualization and exploration. Briefings in Bioinformatics, 14(2):178–192.

Tolstorukov, M. Y., Volfovsky, N., Stephens, R. M., and Park, P. J., 2011. Impact of chromatin structure on sequence variability in the human genome. Nature Structural & Molecular Biology, 18(4):510–515.

Wang, L., Sun, Y., Sun, X., Yu, L., Xue, L., He, Z., Huang, J., Tian, D., Hurst, L. D., and Yang, S., et al., 2020. Repeat-induced point mutation in *Neurospora crassa* causes the highest known mutation rate and mutational burden of any cellular life. Genome Biology, 21(1):142.

Washietl, S., Machné, R., and Goldman, N., 2008. Evolutionary footprints of nucleosome positions in yeast. Trends in Genetics, 24(12):583–587.

Weng, M.-L., Becker, C., Hildebrandt, J., Neumann, M., Rutter, M. T., Shaw, R. G., Weigel, D., and Fenster, C. B., 2019. Fine-grained analysis of spontaneous mutation spectrum and frequency in *Arabidopsis thaliana*. Genetics, 211(2):703–714.

Yazdi, P. G., Pedersen, B. A., Taylor, J. F., Khattab, O. S., Chen, Y.-H., Chen, Y., Jacobsen, S. E., and Wang, P. H., 2015. Increasing nucleosome occupancy is correlated with an increasing mutation rate so long as DNA repair machinery is intact. PLoS One, 10(8):e0136574.

Ying, H., Epps, J., Williams, R., and Huttley, G., 2010. Evidence that localized variation in primate sequence divergence arises from an influence of nucleosome placement on DNA repair. Molecular biology and evolution, 27(3):637–649.

Zhao, J., Gladieux, P., Hutchison, E., Bueche, J., Hall, C., Perraudeau, F., and Glass, N. L., 2015. Identification of allorecognition loci in *Neurospora crassa* by genomics and evolutionary approaches. Molecular Biology and Evolution, 32(9):2417–2432.

Zhu, Y. O., Siegal, M. L., Hall, D. W., and Petrov, D. A., 2014. Precise estimates of mutation rate and spectrum in yeast. Proceedings of the National Academy of Sciences, 111(22):E2310– E2318.

## Supplementary References

Abyzov, A., Urban, A. E., Snyder, M., and Gerstein, M., 2011. CNVnator: An approach to discover, genotype, and characterize typical and atypical CNVs from family and population genome sequencing. Genome Research, 21(6):974–984.

Chiang, C., Layer, R. M., Faust, G. G., Lindberg, M. R., Rose, D. B., Garrison, E. P., Marth, G. T., Quinlan, A. R., and Hall, I. M., 2015. Speedseq: ultra-fast personal genome analysis and interpretation. Nature Methods, 12(10):966–968.

Cingolani, P., Platts, A., Coon, M., Nguyen, T., Wang, L., Land, S., Lu, X., and Ruden, D., 2012. A program for annotating and predicting the effects of single nucleotide polymorphisms, SnpEff: SNPs in the genome of *Drosophila melanogaster* strain w1118; iso-2; iso-3. Fly, 6(2):80–92.

Danecek, P., Bonfield, J. K., Liddle, J., Marshall, J., Ohan, V., Pollard, M. O., Whitwham, A., Keane, T., McCarthy, S. A., Davies, R. M., et al., 2021. Twelve years of SAMtools and BCFtools. GigaScience, 10(2). giab008.

Davis, R. H. and de Serres, F. J., 1970. Genetic and microbiological research techniques for *Neurospora crassa*. Methods in Enzymology, 17:79–143.

Faust, G. G. and Hall, I. M., 2014. SAMBLASTER: fast duplicate marking and structural variant read extraction. Bioinformatics, 30(17):2503–2505.

Freitag, M., Hickey, P. C., Raju, N. B., Selker, E. U., and Read, N. D., 2004. GFP as a tool to analyze the organization, dynamics and function of nuclei and microtubules in *Neurospora crassa*. Fungal Genetics and Biology, 41(10):897–910.

Galagan, J. E., Calvo, S. E., Borkovich, K. A., Selker, E. U., Read, N. D., Jaffe, D., FitzHugh, W., Ma, L.-J., Smirnov, S., Purcell, S., et al., 2003. The genome sequence of the filamentous fungus *Neurospora crassa*. Nature, 422(6934):859–868.

Gelman, A., Goodrich, B., Gabry, J., and Vehtari, A., 2019. R-squared for Bayesian regression models. The American Statistician, 73(3):307–309.

Homer, N., 2021. DWGSIM: Whole genome simulator for next-generation sequencing. github repository.

Kosugi, S., Momozawa, Y., Liu, X., Terao, C., Kubo, M., and Kamatani, Y., 2019. Comprehensive evaluation of structural variation detection algorithms for whole genome sequencing. Genome Biology, 20(1):117.

Kronholm, I., Ormsby, T., McNaught, K. J., Selker, E. U., and Ketola, T., 2020. Marked *Neurospora crassa* strains for competition experiments and Bayesian methods for fitness estimates. G3: Genes|Genomes|Genetics, 10:1261–1270.

Kühl, M., Stich, B., and Ries, D., 2021. Mutation-Simulator: fine-grained simulation of random mutations in any genome. Bioinformatics, 37(4):568–569.

Lichius, A. and Zeilinger, S., 2019. Application of membrane and cell wall selective fluorescent dyes for live-cell imaging of filamentous fungi. JoVE, (153):e60613.

McElreath, R., 2015. Statistical Rethinking - A Bayesian course with examples in R and Stan. CRC Press, New York.

Metzenberg, R. L., 2003. Vogel’s medium N salts: Avoiding the need for ammonium nitrate. Fungal Genetics Newsletter, 50:14.

Min, Y. and Agresti, A. a., 2002. Modeling nonnegative data with clumping at zero: A survey. JIRSS, 1(1):7–33.

Rueden, C. T., Schindelin, J., Hiner, M. C., DeZonia, B. E., Walter, A. E., Arena, E. T., and Eliceiri, K. W., 2017. ImageJ2: ImageJ for the next generation of scientific image data. BMC Bioinformatics, 18(1):529.

Schindelin, J., Arganda-Carreras, I., Frise, E., Kaynig, V., Longair, M., Pietzsch, T., Preibisch, S., Rueden, C., Saalfeld, S., Schmid, B., et al., 2012. Fiji: an open-source platform for biological-image analysis. Nature Methods, 9(7):676–682.

Song, Q. and Smith, A. D., 2011. Identifying dispersed epigenomic domains from ChIP-Seq data. Bioinformatics, 27(6):870–871.

Vehtari, A., Gelman, A., and Gabry, J., 2017. Practical Bayesian model evaluation using leave-one-out cross-validation and WAIC. Statistics and Computing, 27(5):1413–1432.

Wala, J. A., Bandopadhayay, P., Greenwald, N. F., O’Rourke, R., Sharpe, T., Stewart, C., Schumacher, S., Li, Y., Weischenfeldt, J., Yao, X., et al., 2018. SvABa: genome-wide detection of structural variants and indels by local assembly. Genome Research, 28(4):581–591.

Xie, C. and Tammi, M. T., 2009. CNV-seq, a new method to detect copy number variation using high-throughput sequencing. BMC Bioinformatics, 10(1):80.

Xing, Y., Dabney, A. R., Li, X., Wang, G., Gill, C. A., and Casola, C., 2020. SECNVs: a simulator of copy number variants and whole-exome sequences from reference genomes. Frontiers in Genetics, 11:82.

Ye, K., Schulz, M. H., Long, Q., Apweiler, R., and Ning, Z., 2009. Pindel: a pattern growth approach to detect break points of large deletions and medium sized insertions from paired-end short reads. Bioinformatics, 25(21):2865–2871.

