## Supplementary figures and images for "Chromatin structure influences rate and spectrum of spontaneous mutations in *Neurospora crassa*"

### mutation_centromer_1.jpg

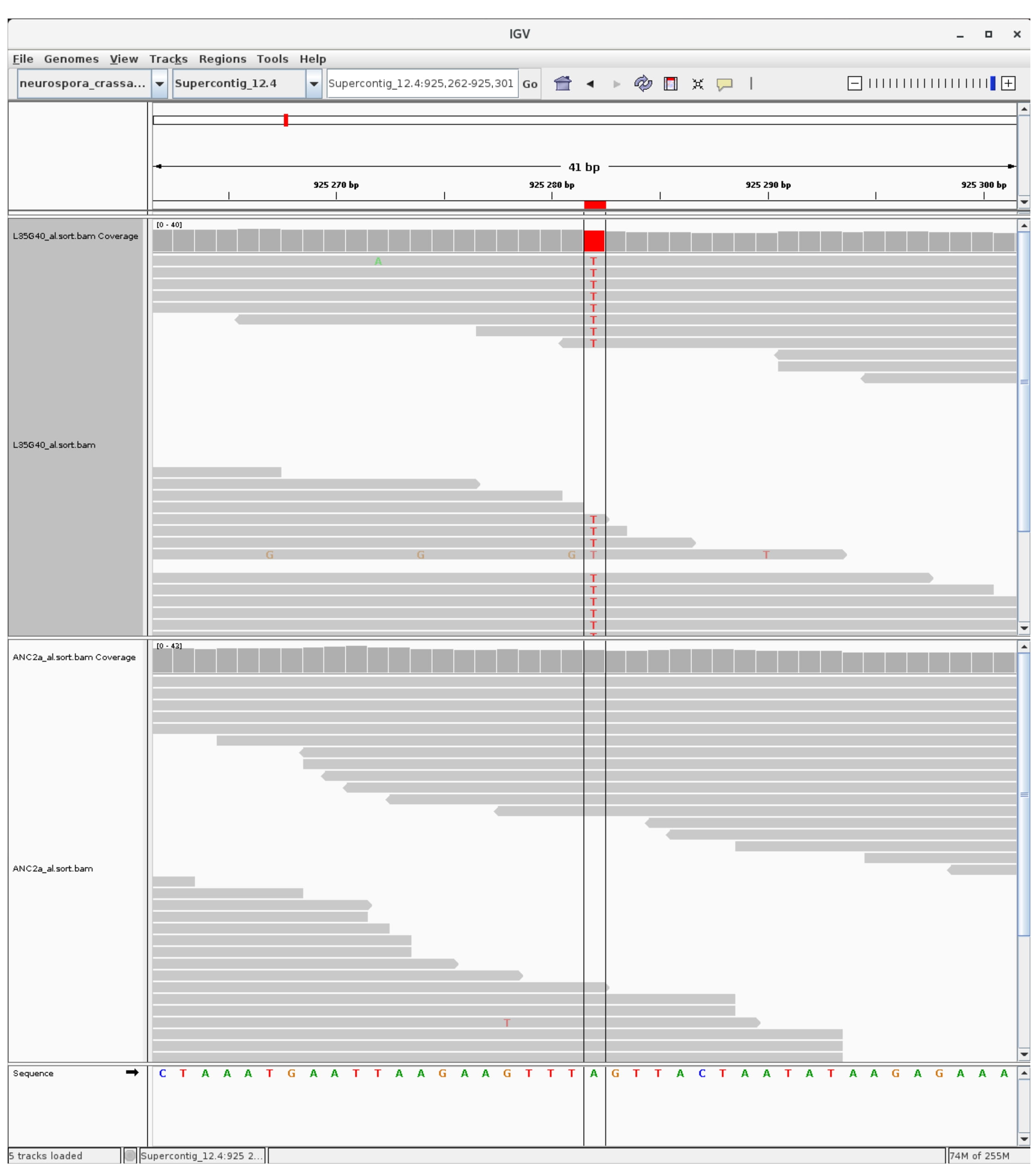

### mutation_centromer_2.jpg

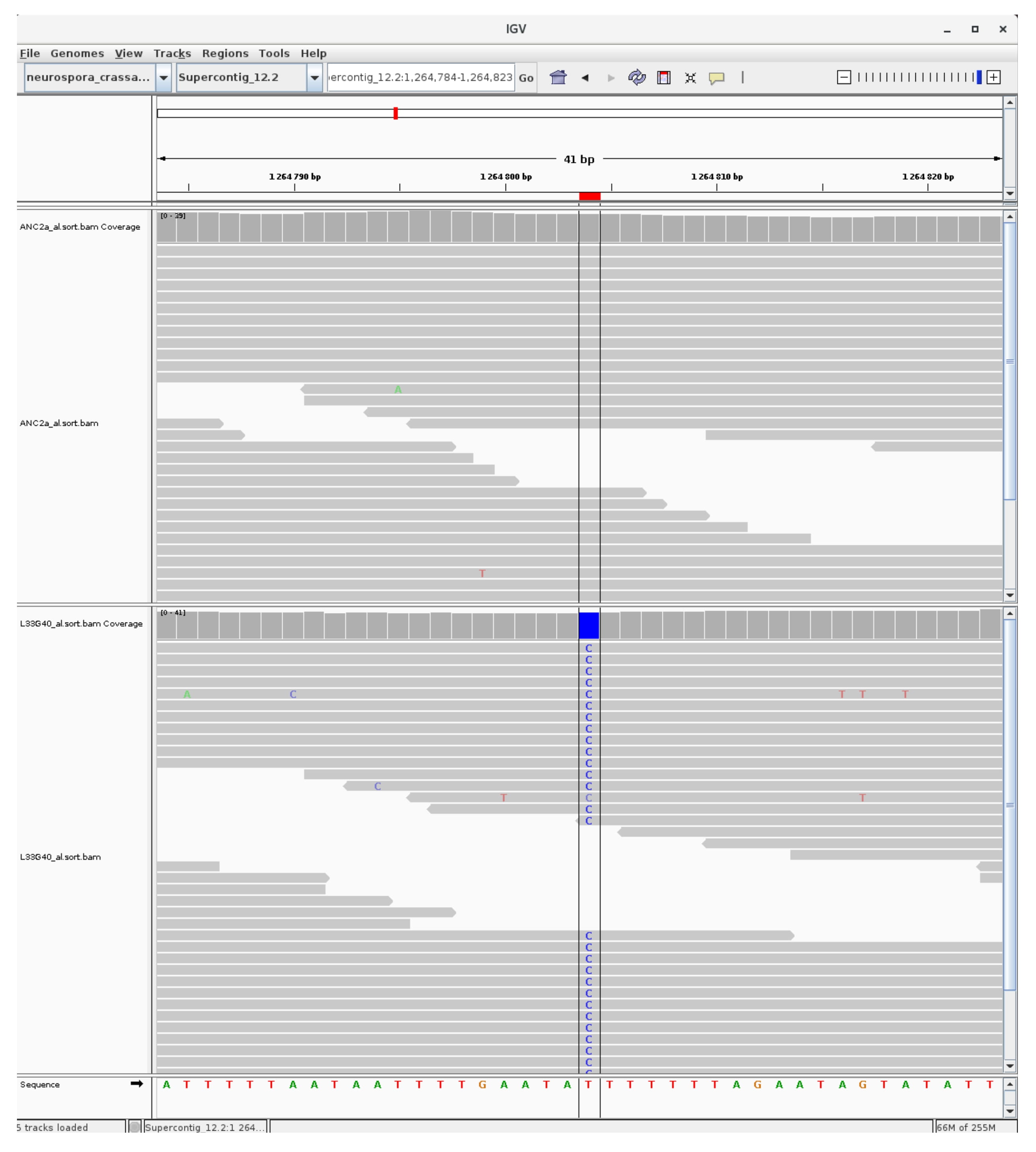

### mutation_centromer_3.jpg

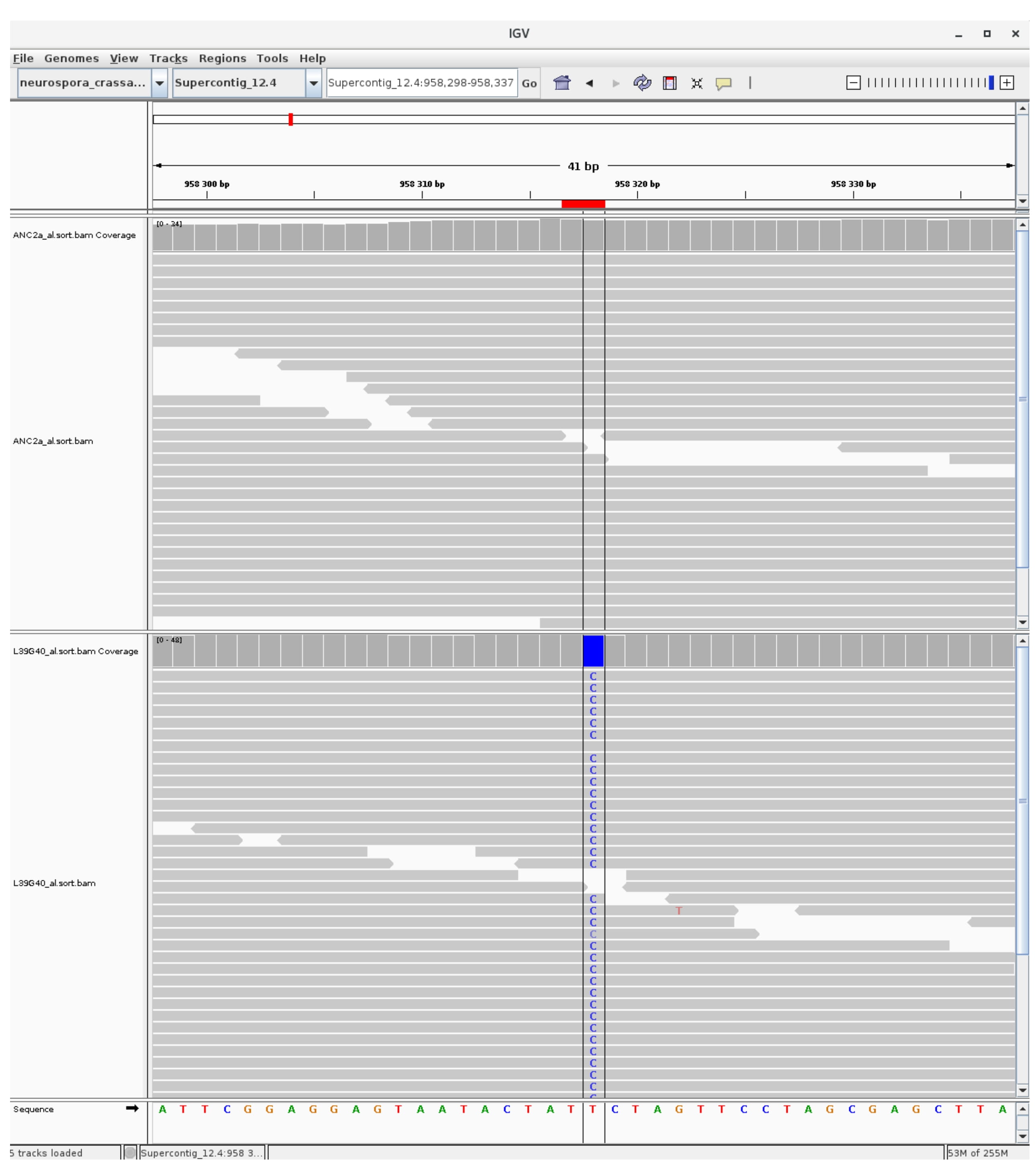

### mutation_centromer_4.jpg

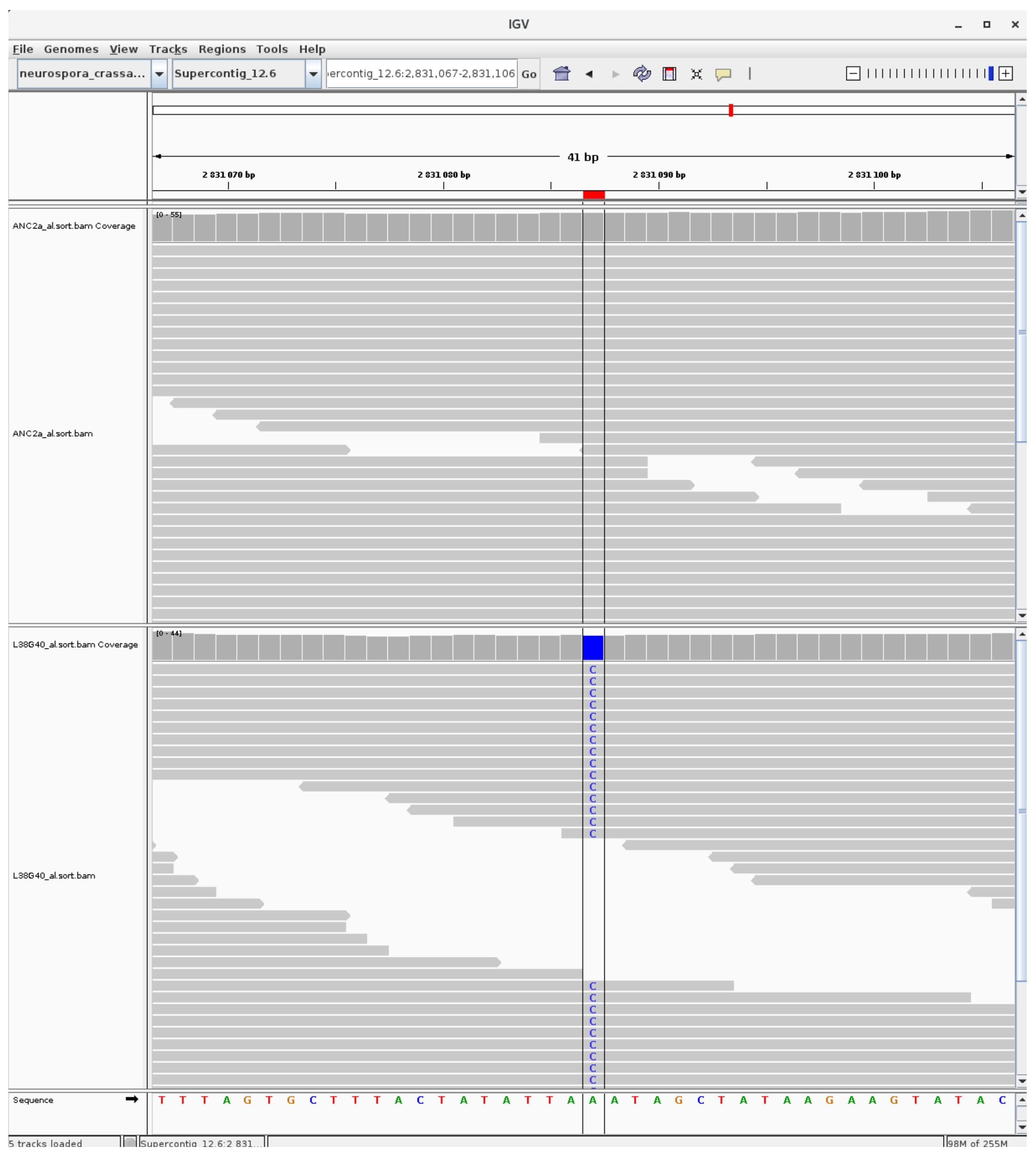

### mutation_centromer_5.jpg

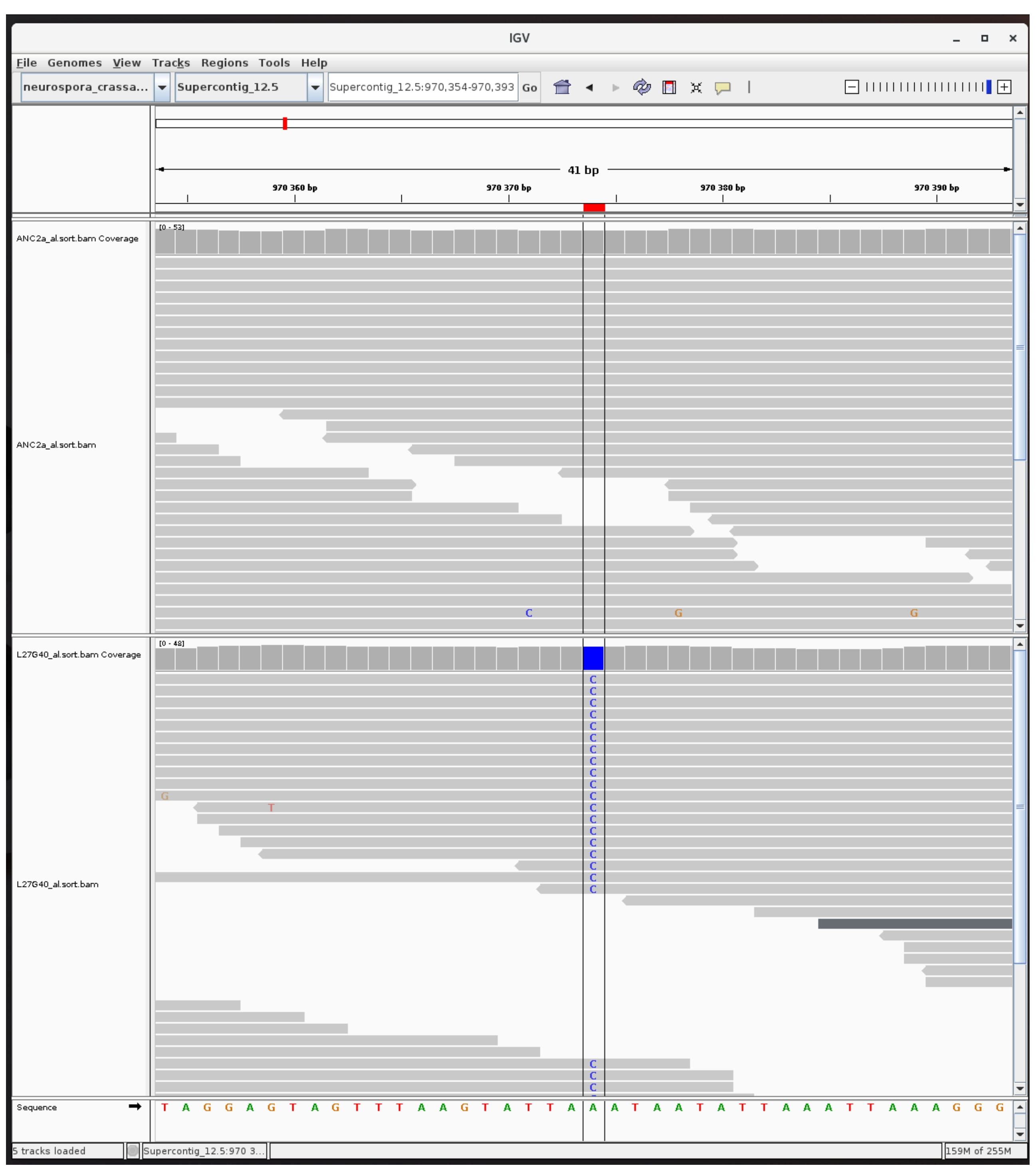

### mutation_centromer_6.jpg

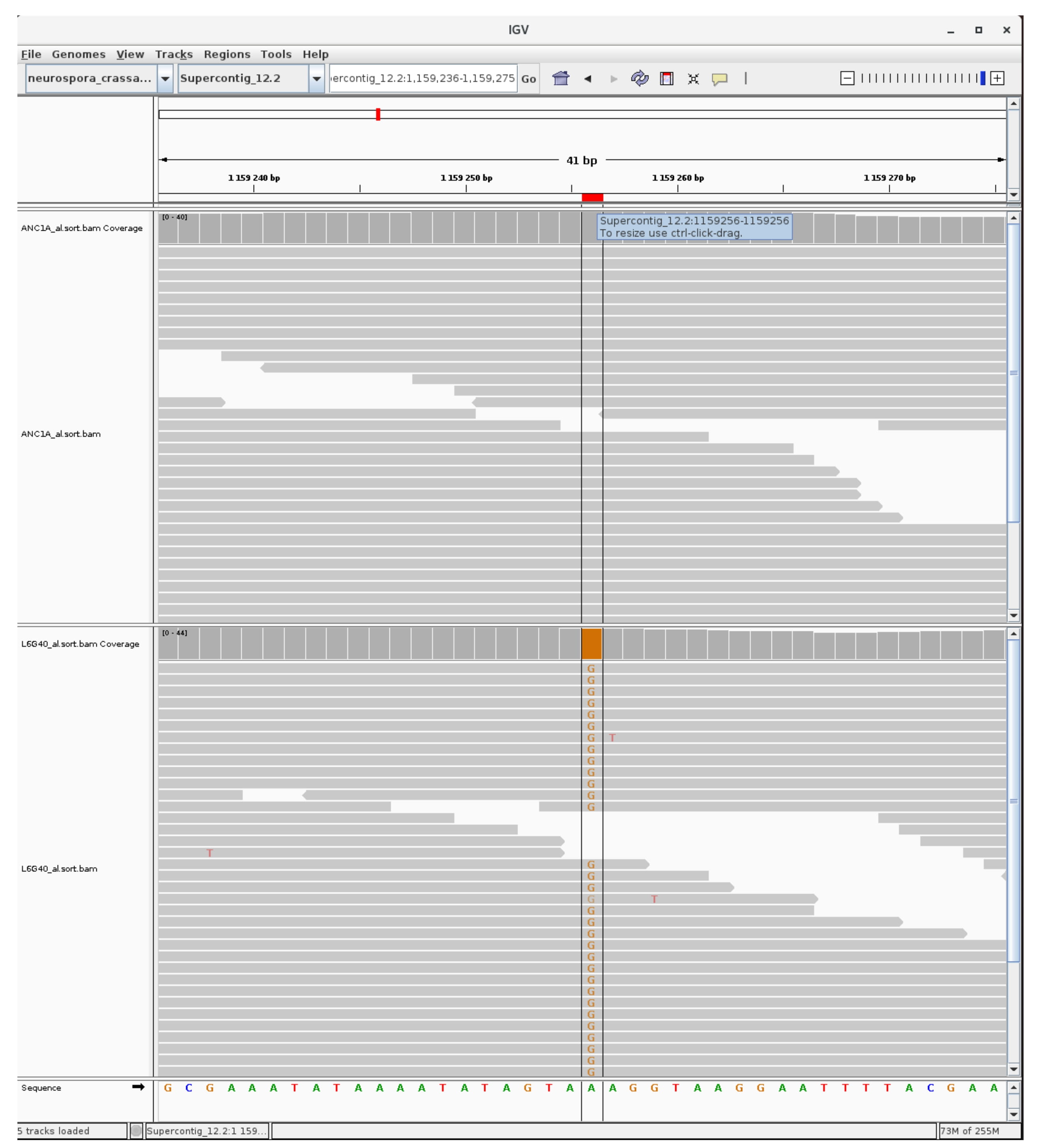

### mutation_centromer_7.jpg

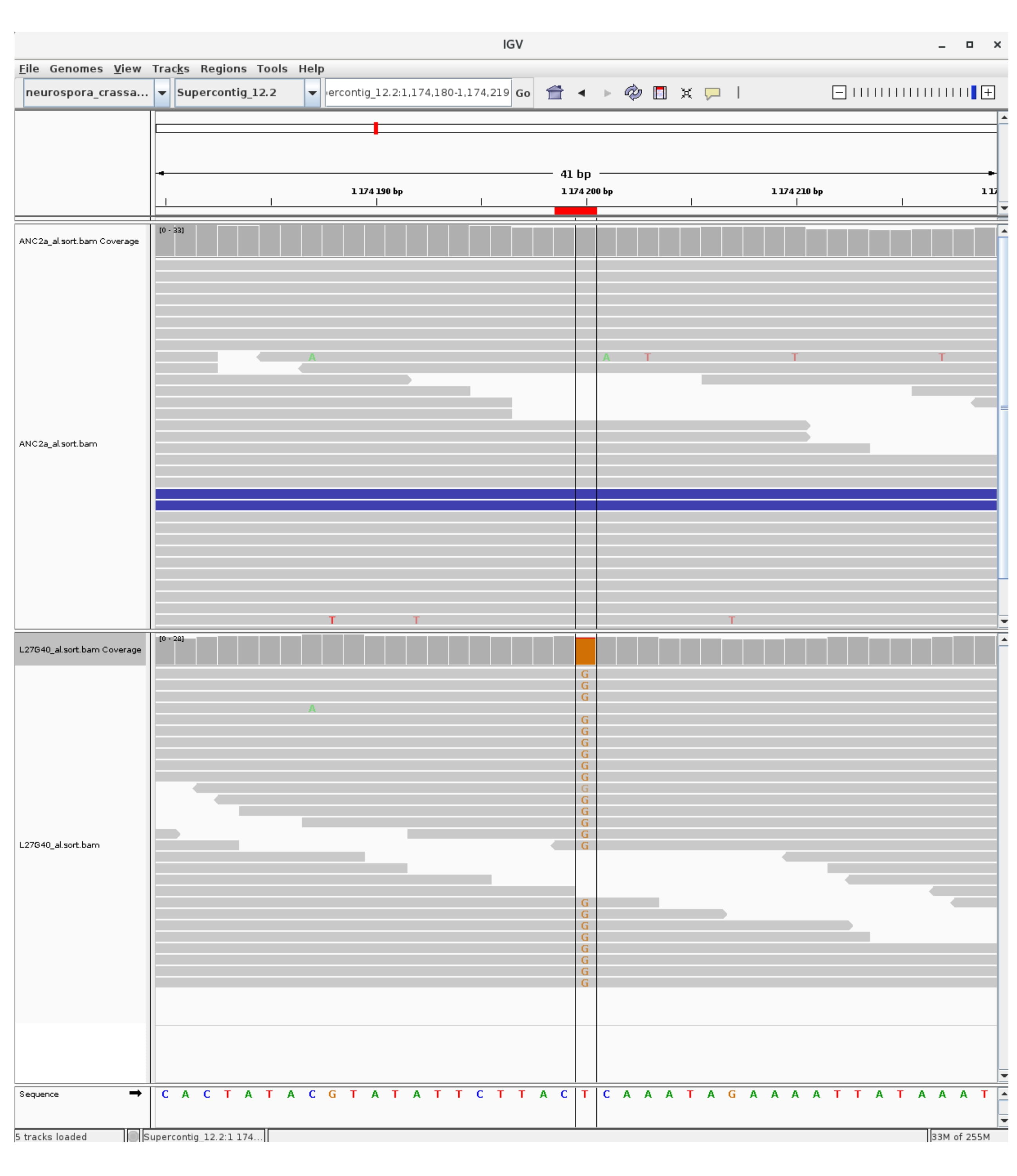

### mutation_centromer_8.jpg

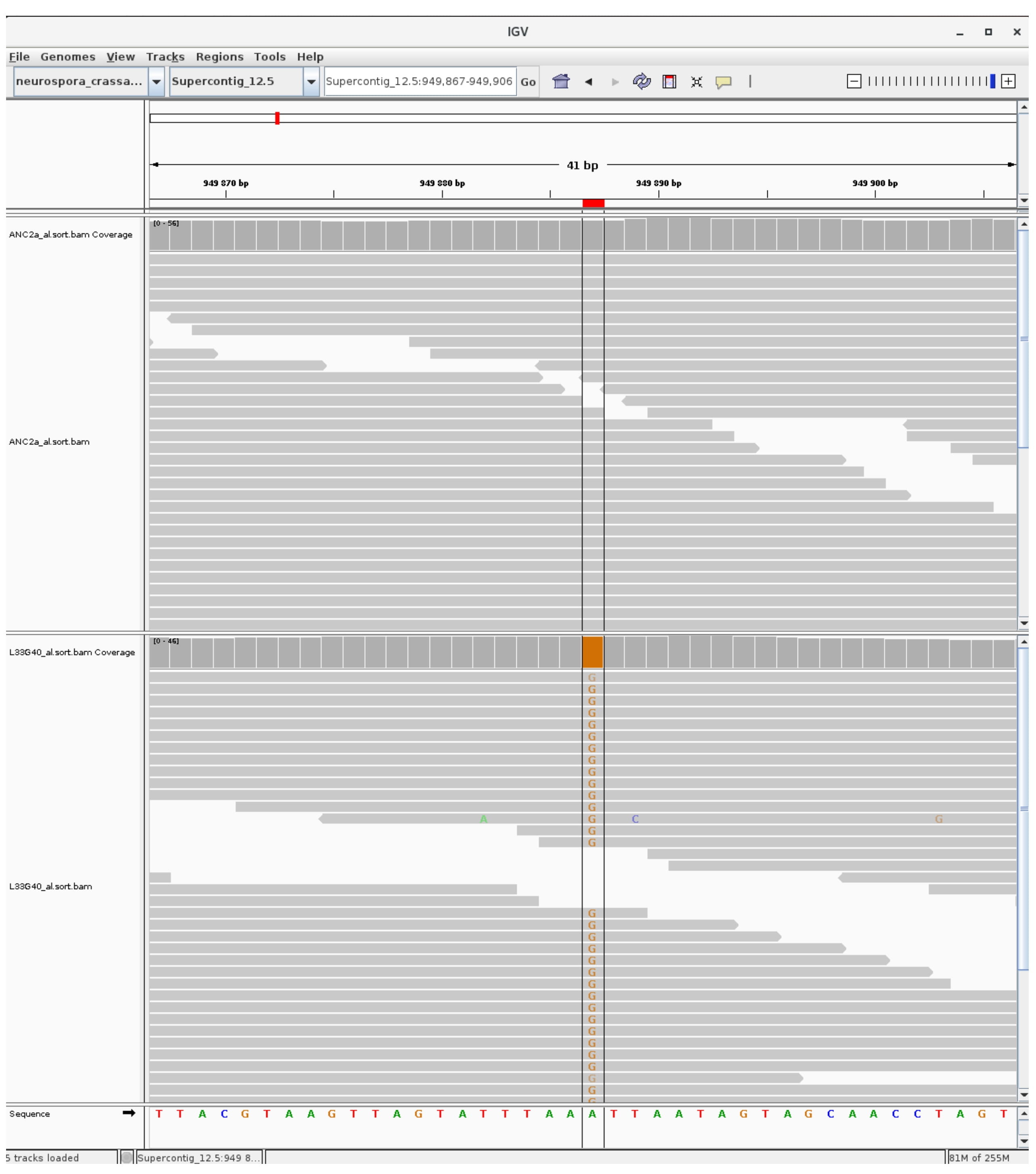

### mutation_centromer_9.jpg

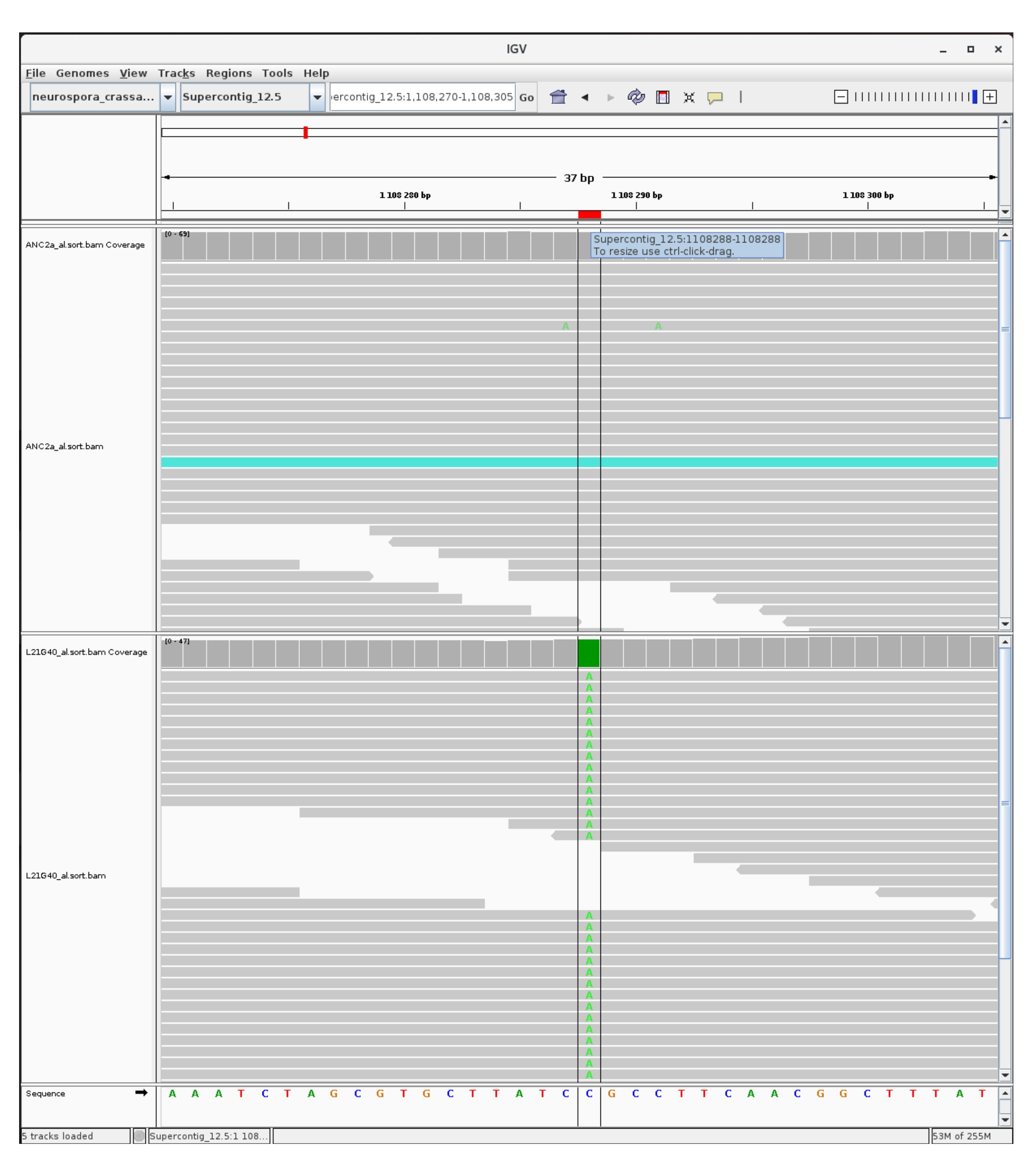

### mutation_centromer_10.jpg

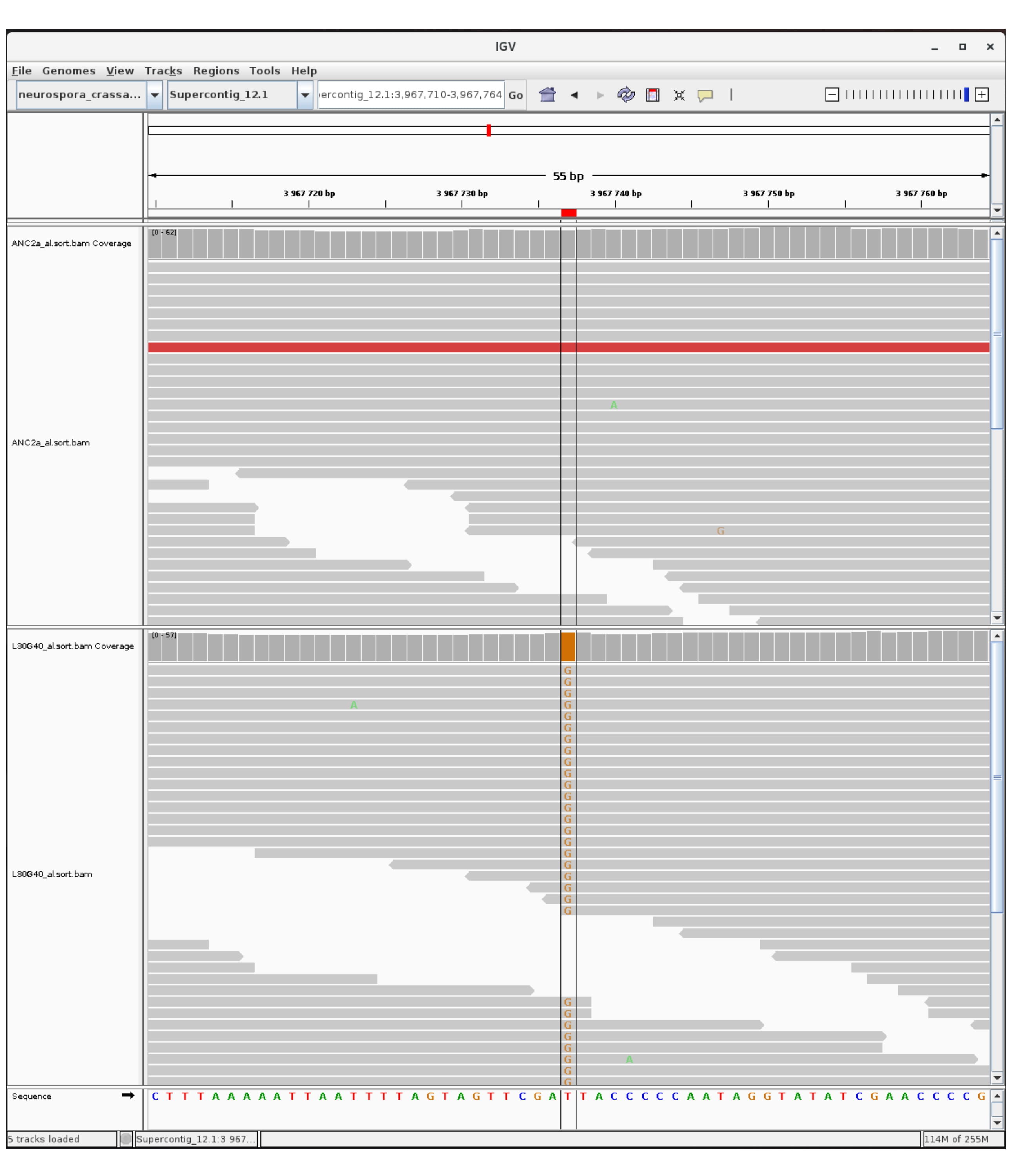

### mutation_centromer_11.jpg

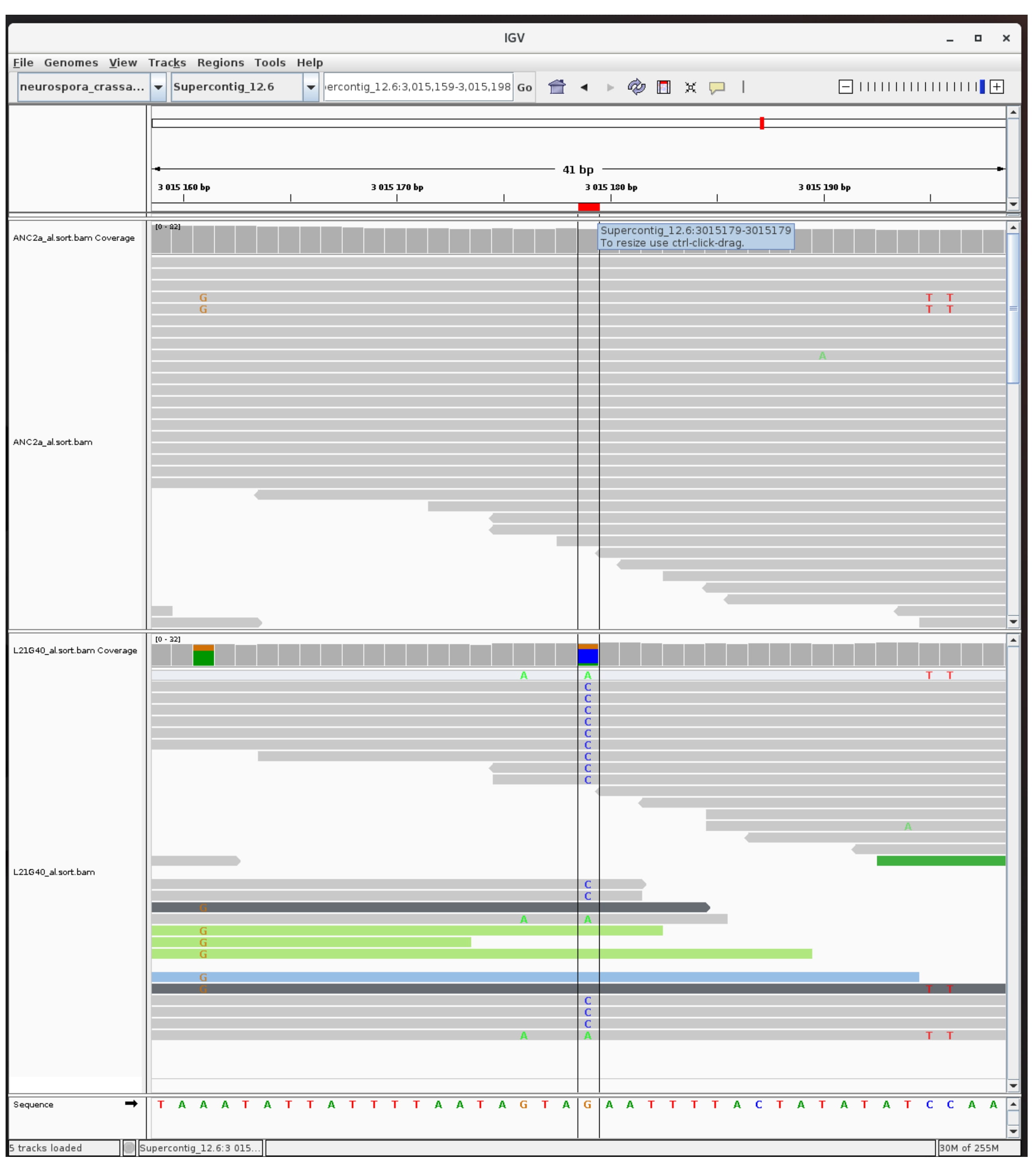

### mutation_centromer_12.jpg

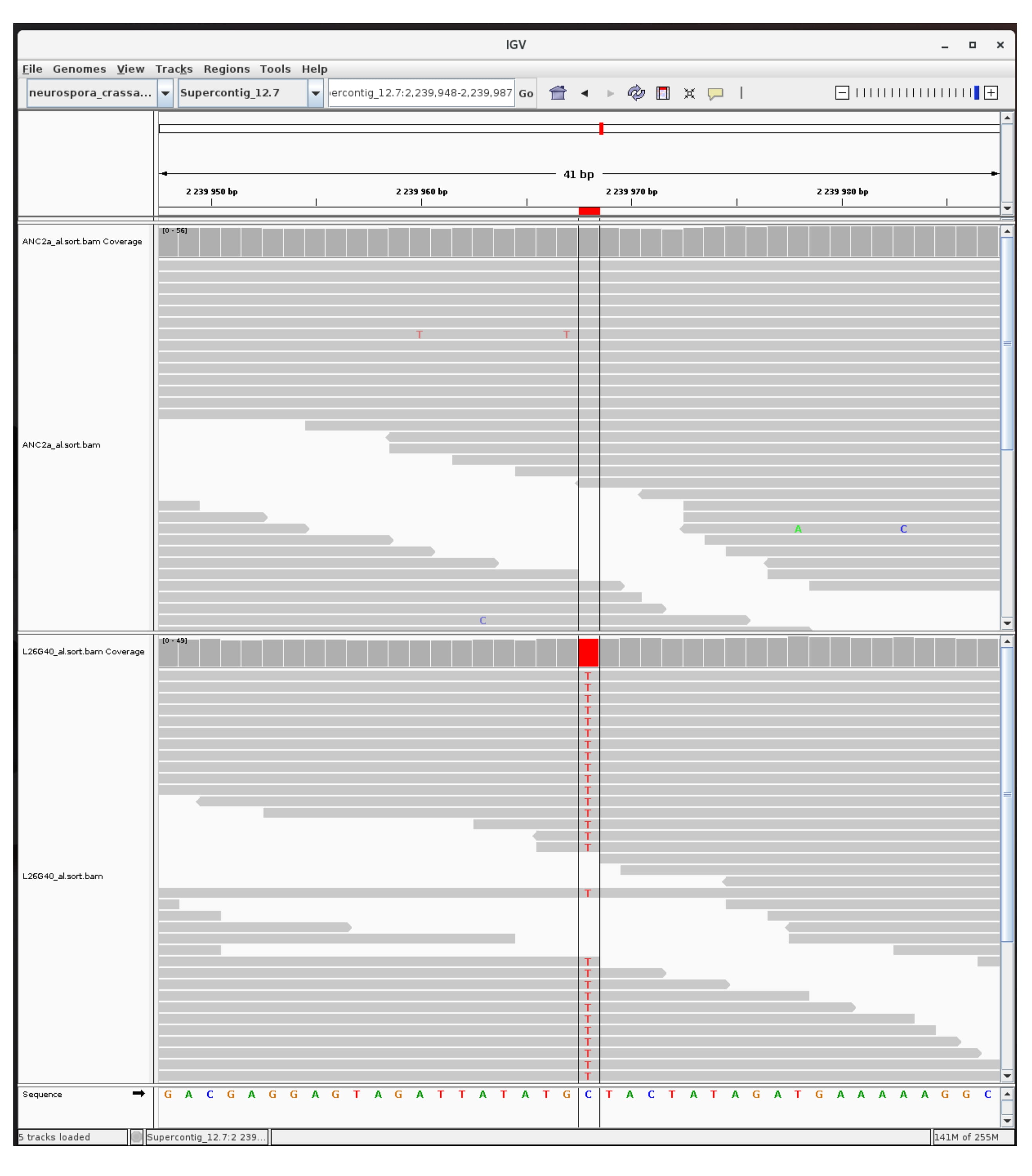

### mutation_centromer_13.jpg

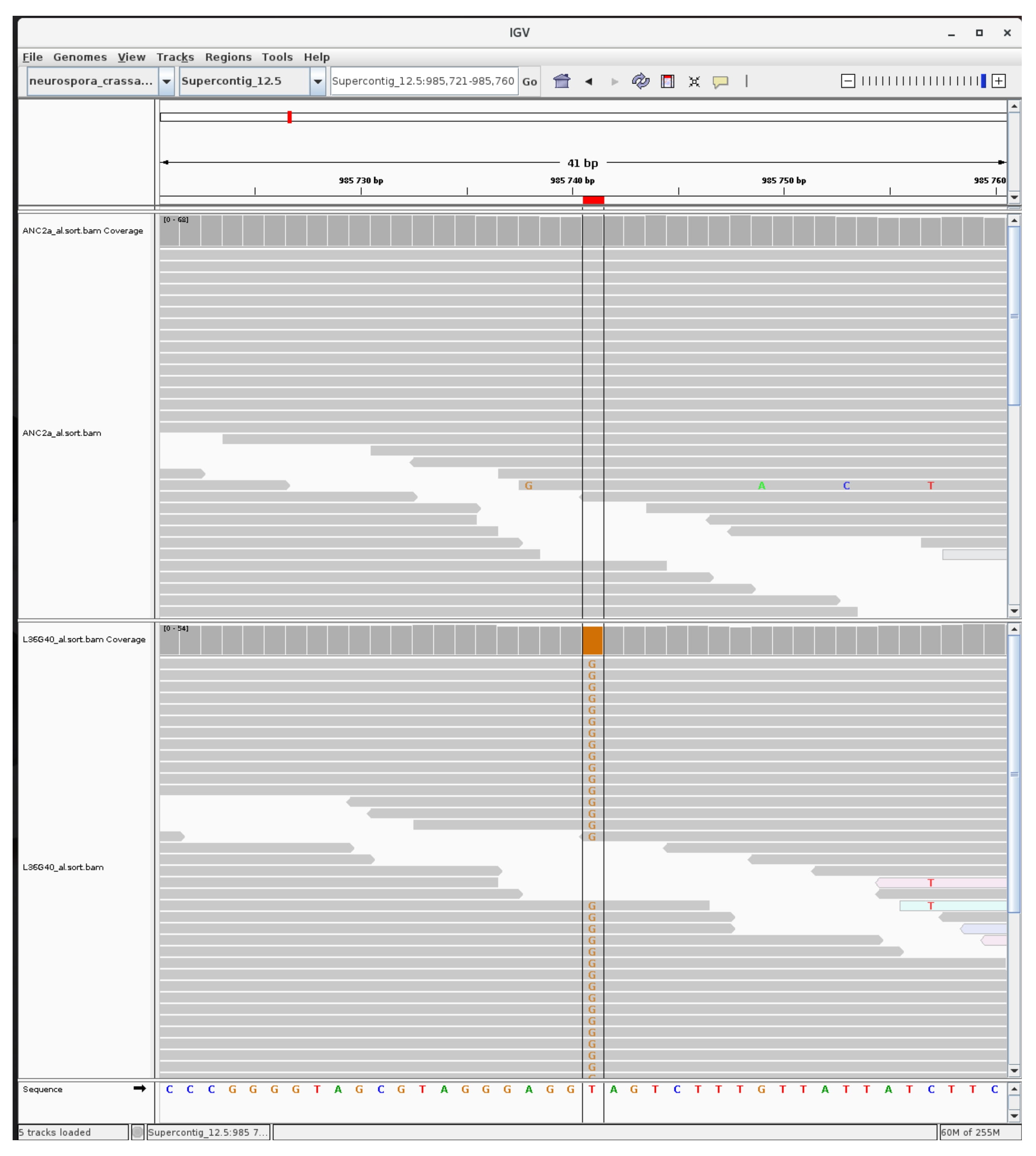

### mutation_centromer_14.jpg

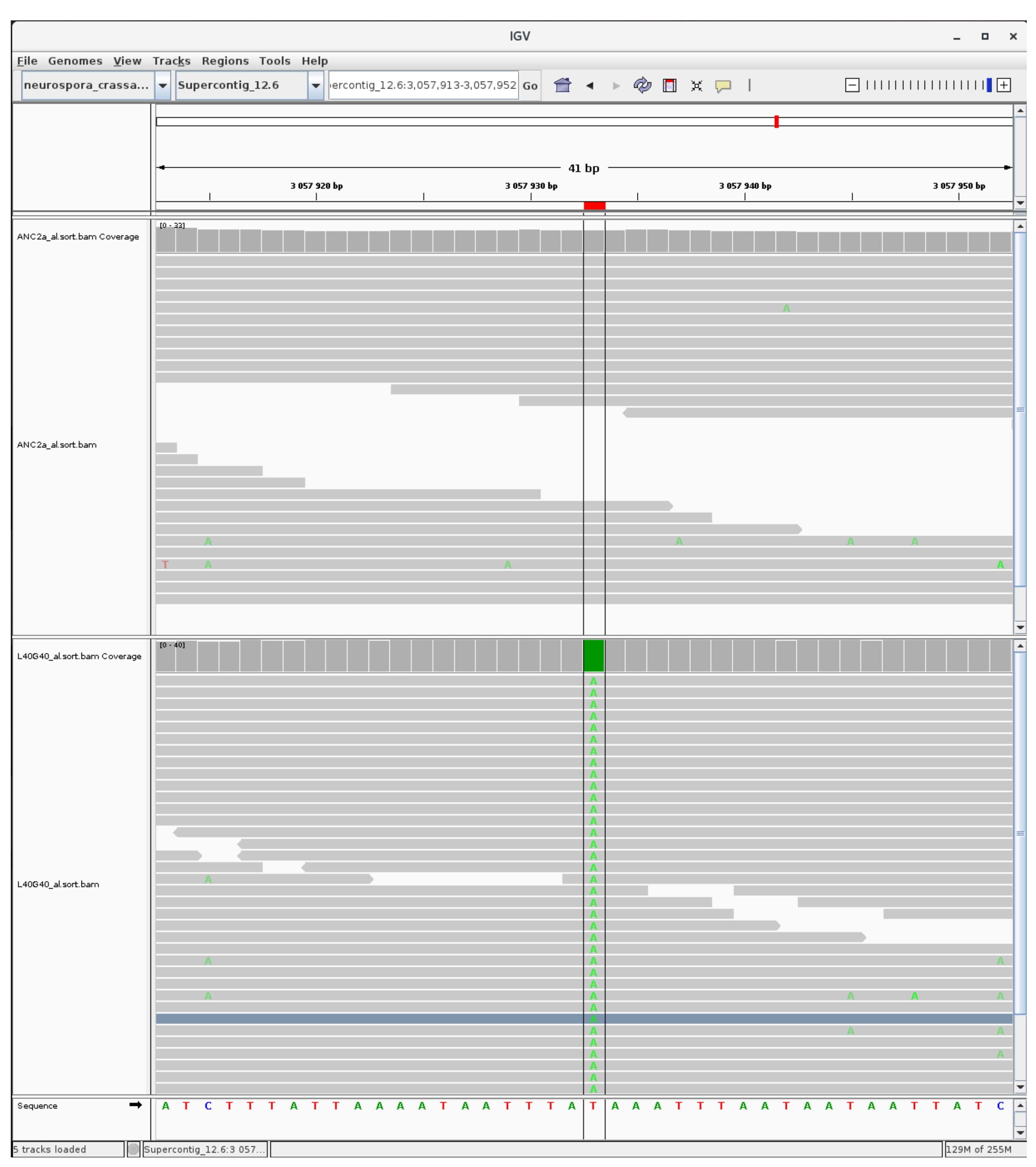

### mutation_centromer_15.jpg

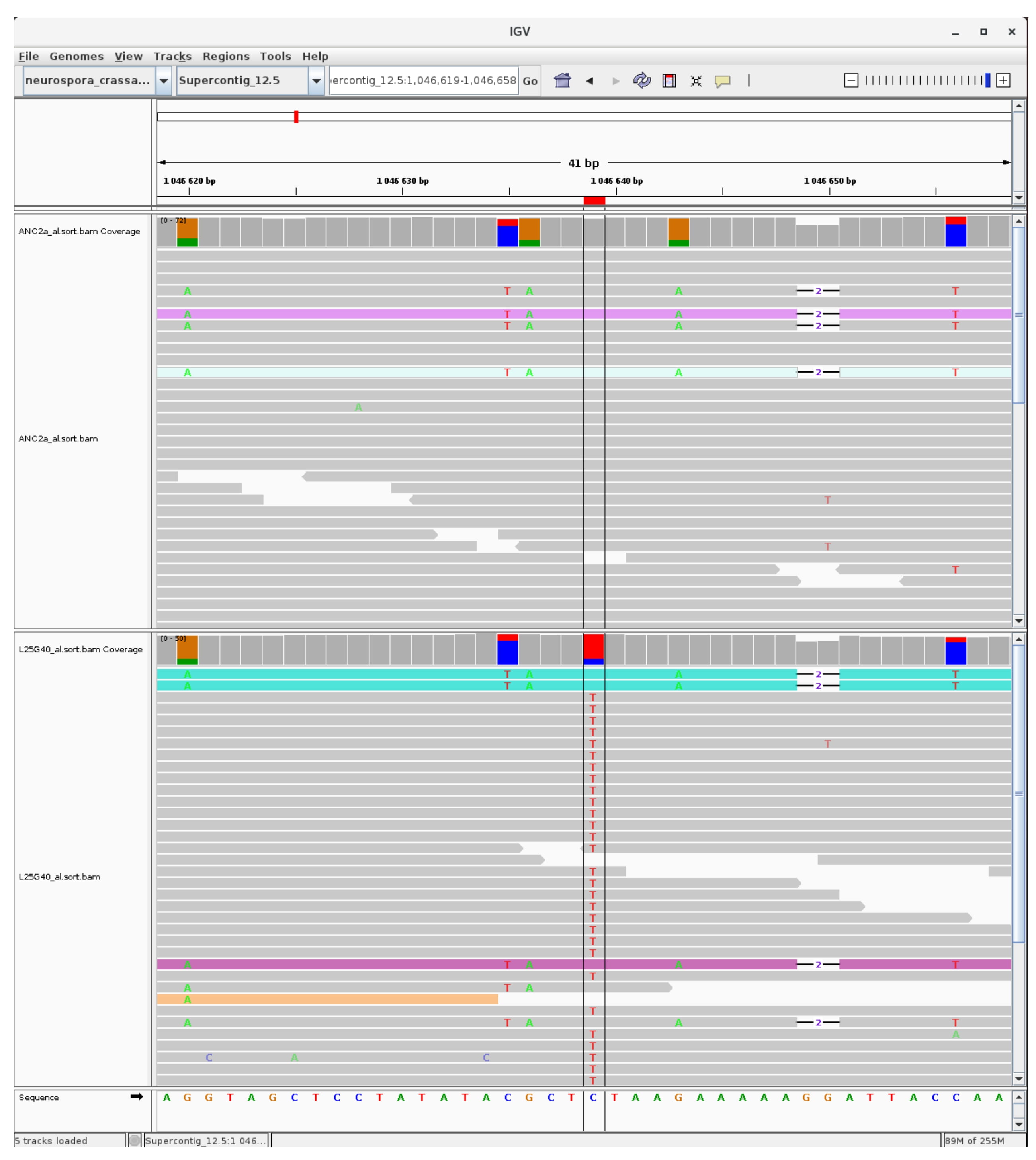

### mutation_centromer_16.jpg

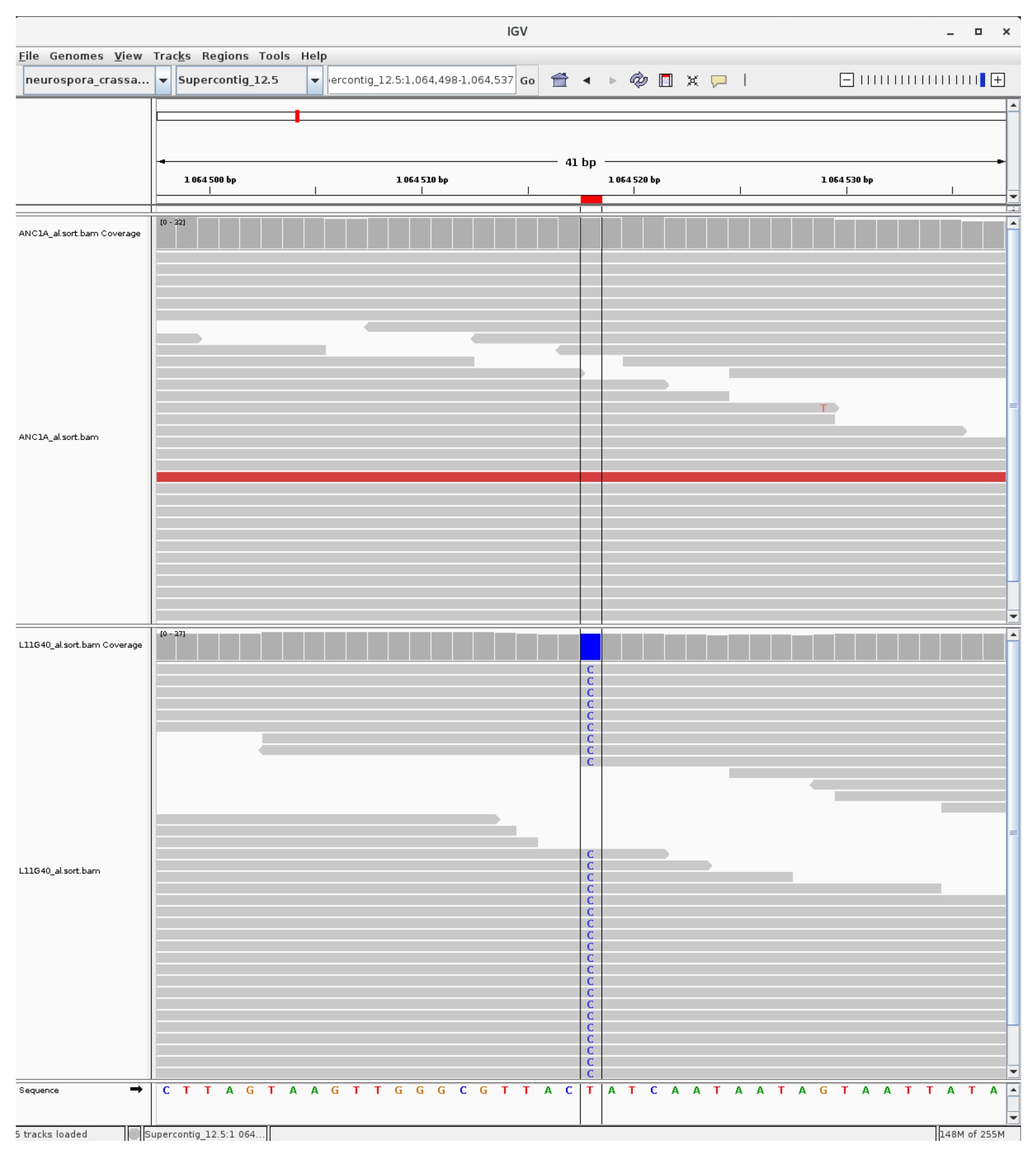

### mutation_centromer_17.jpg

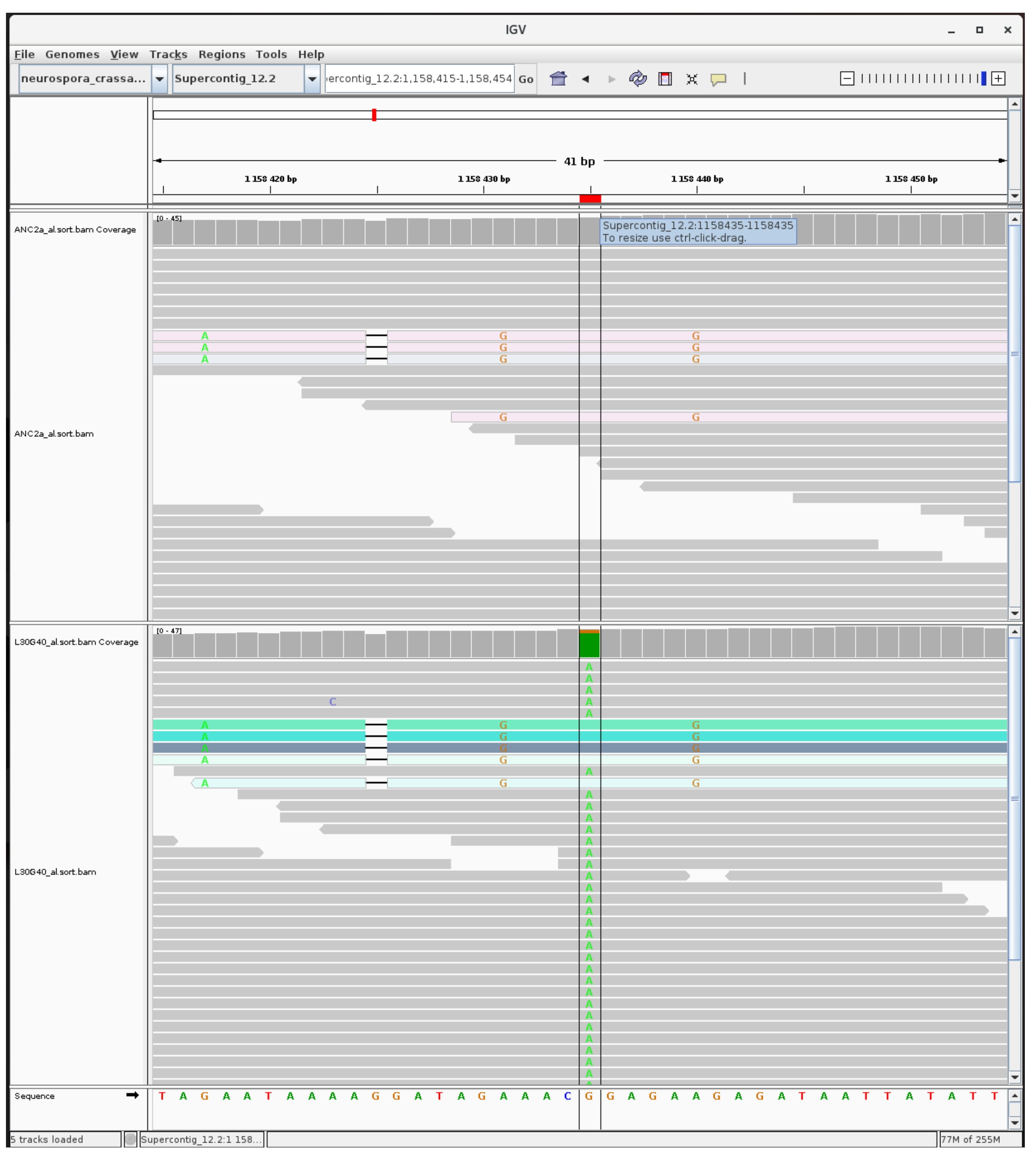

### mutation_centromer_18.jpg

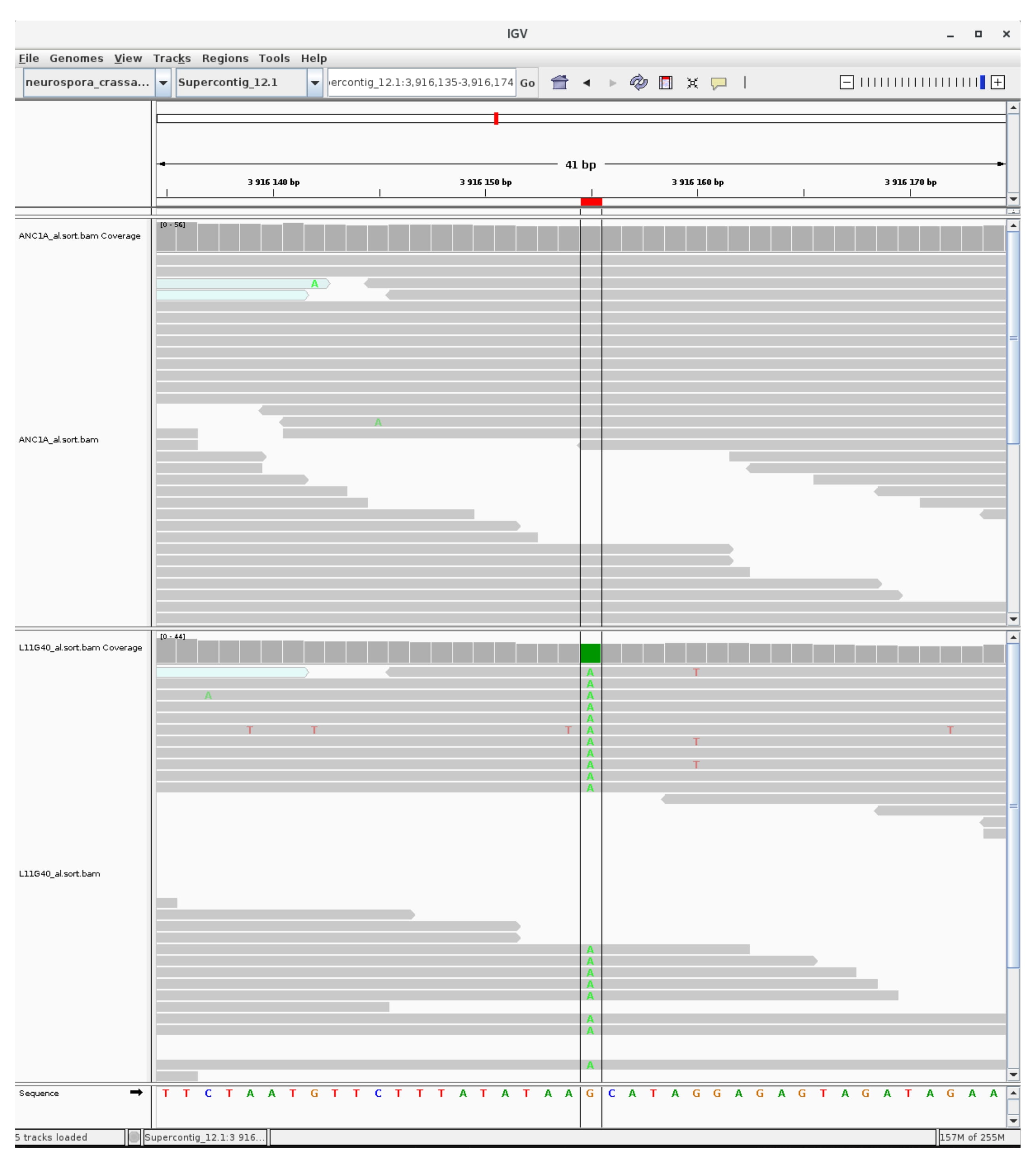

### mutation_centromer_19.jpg

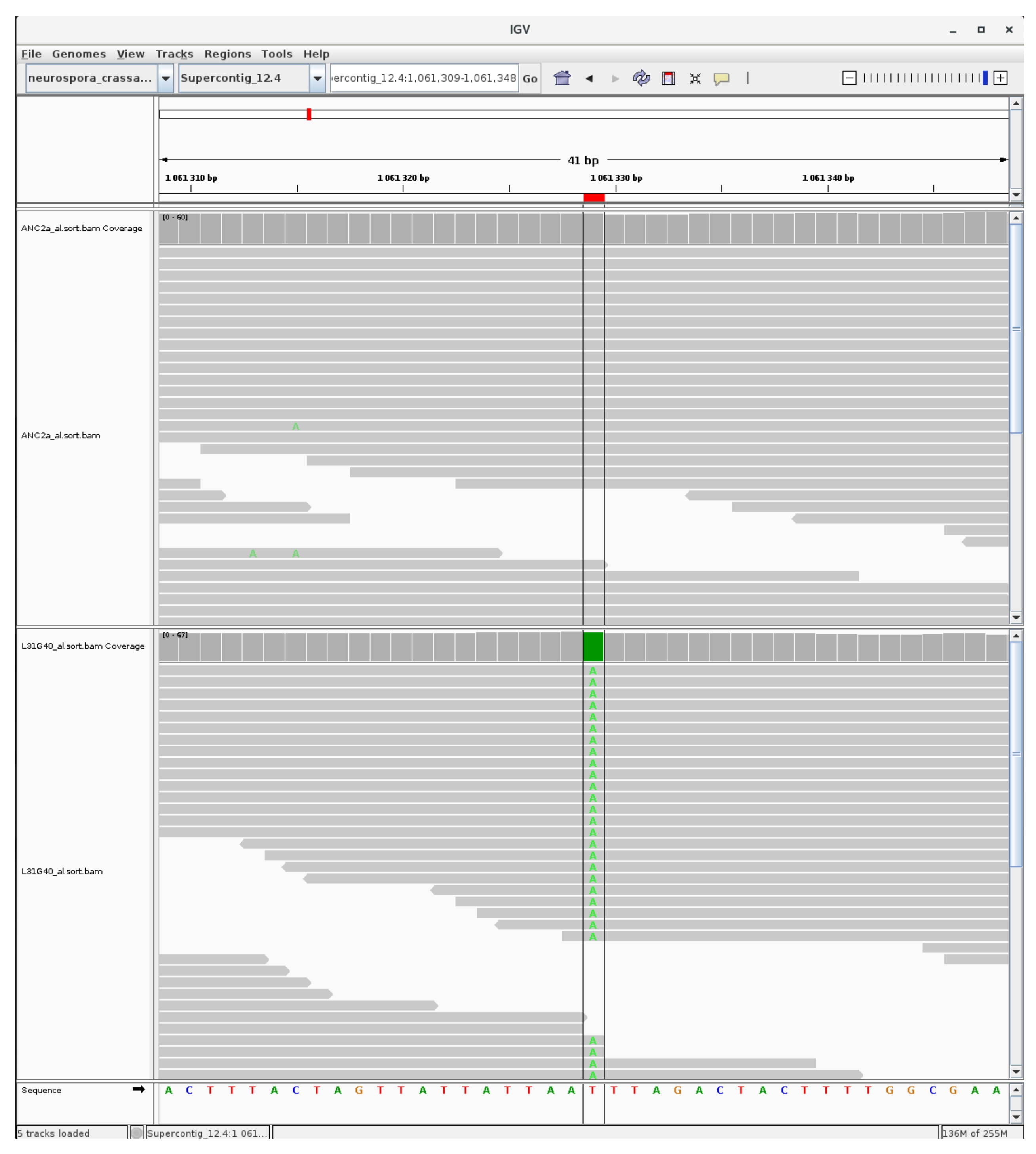

### mutation_centromer_20.jpg

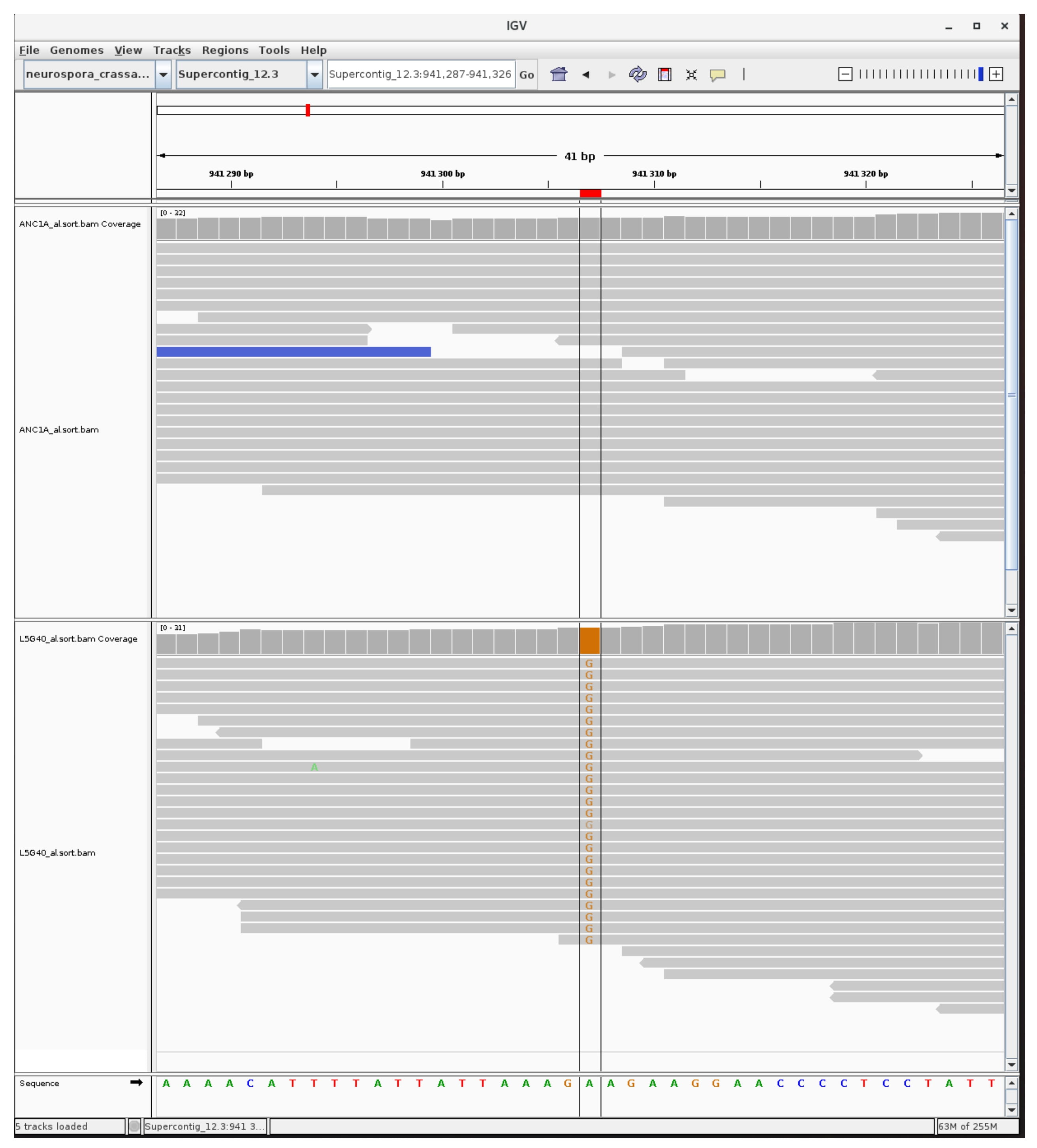

### mutation_centromer_21.jpg

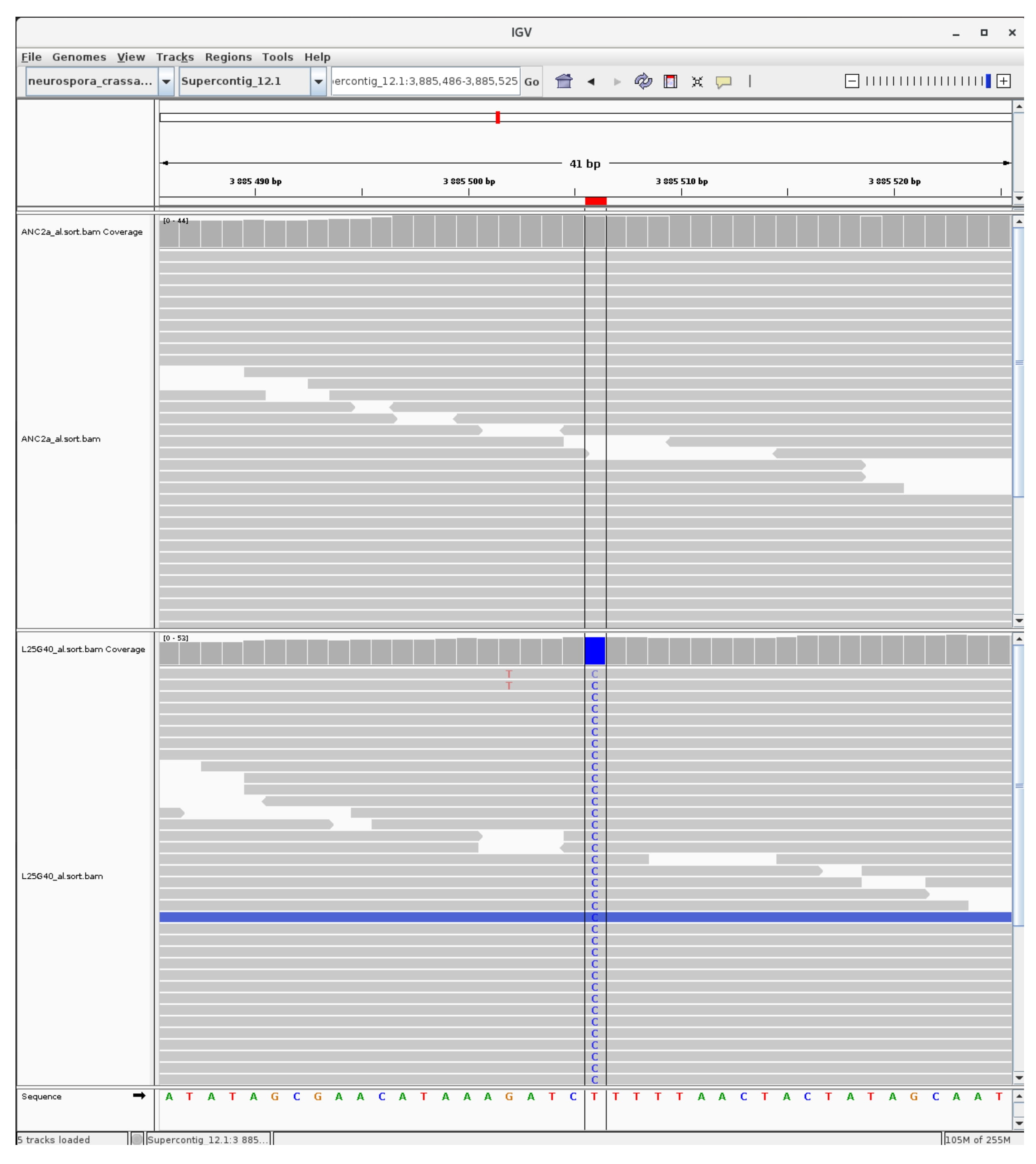

### mutation_centromer_22.jpg

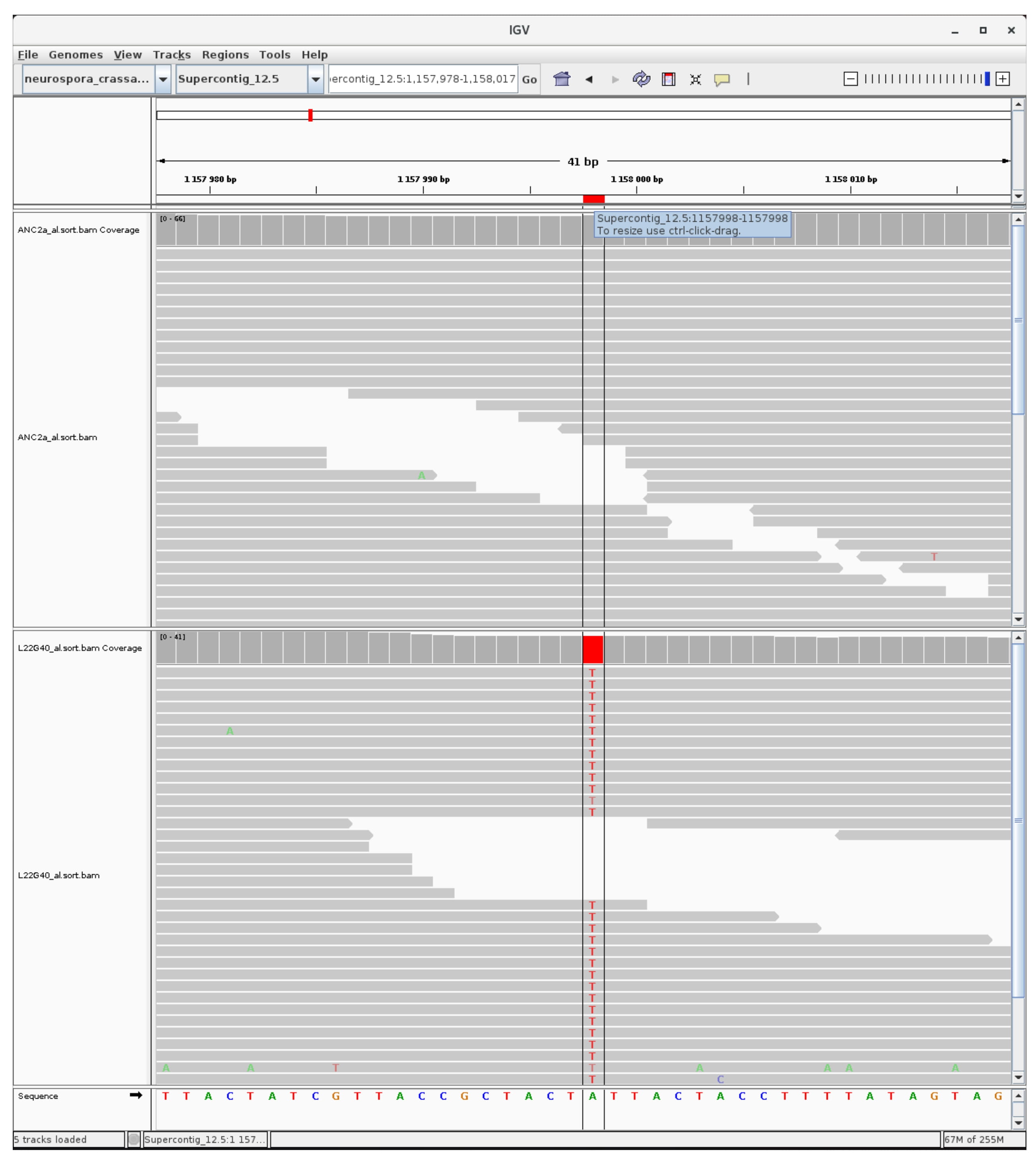

### mutation_centromer_23.jpg

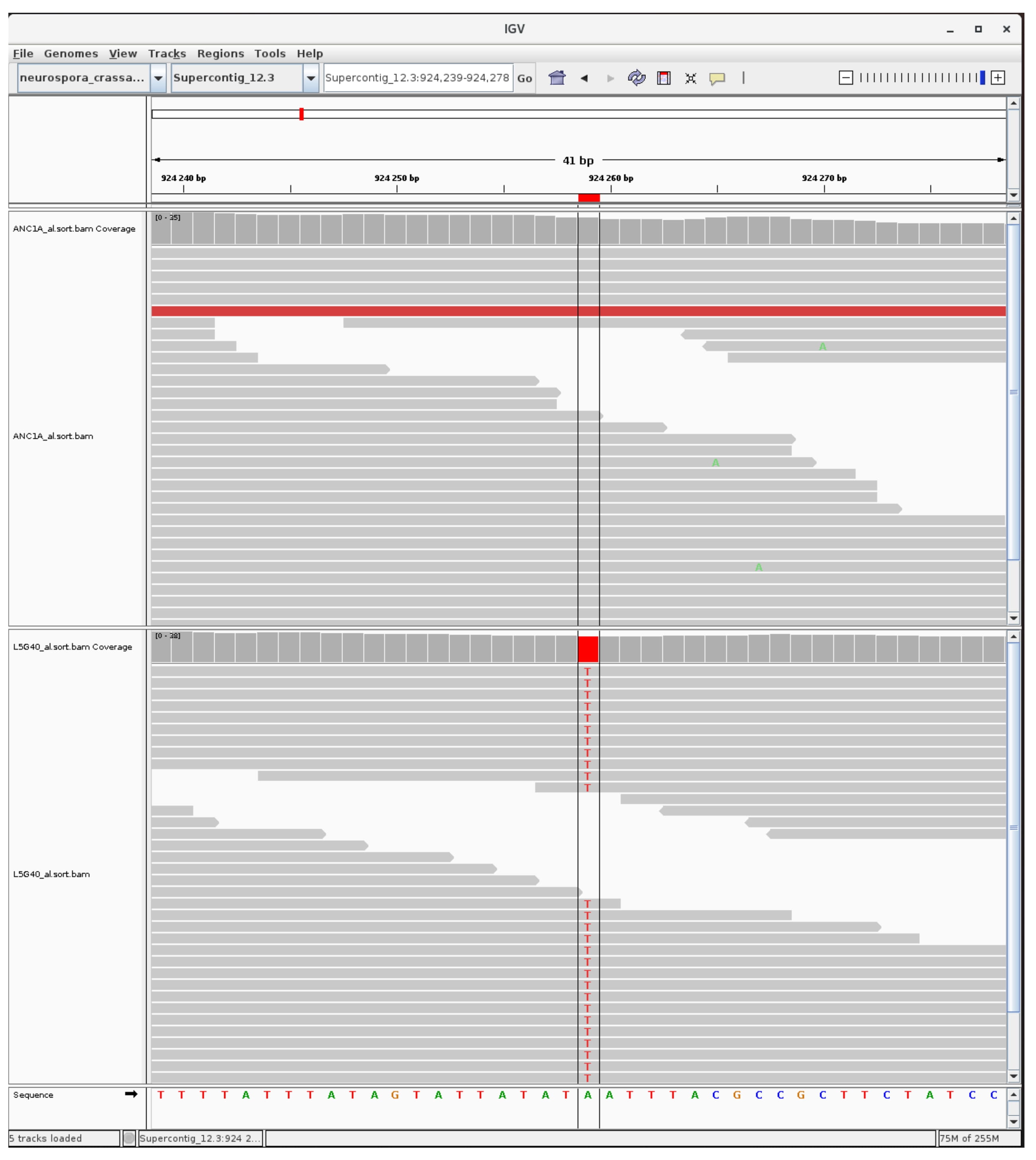

### mutation_centromer_24.jpg

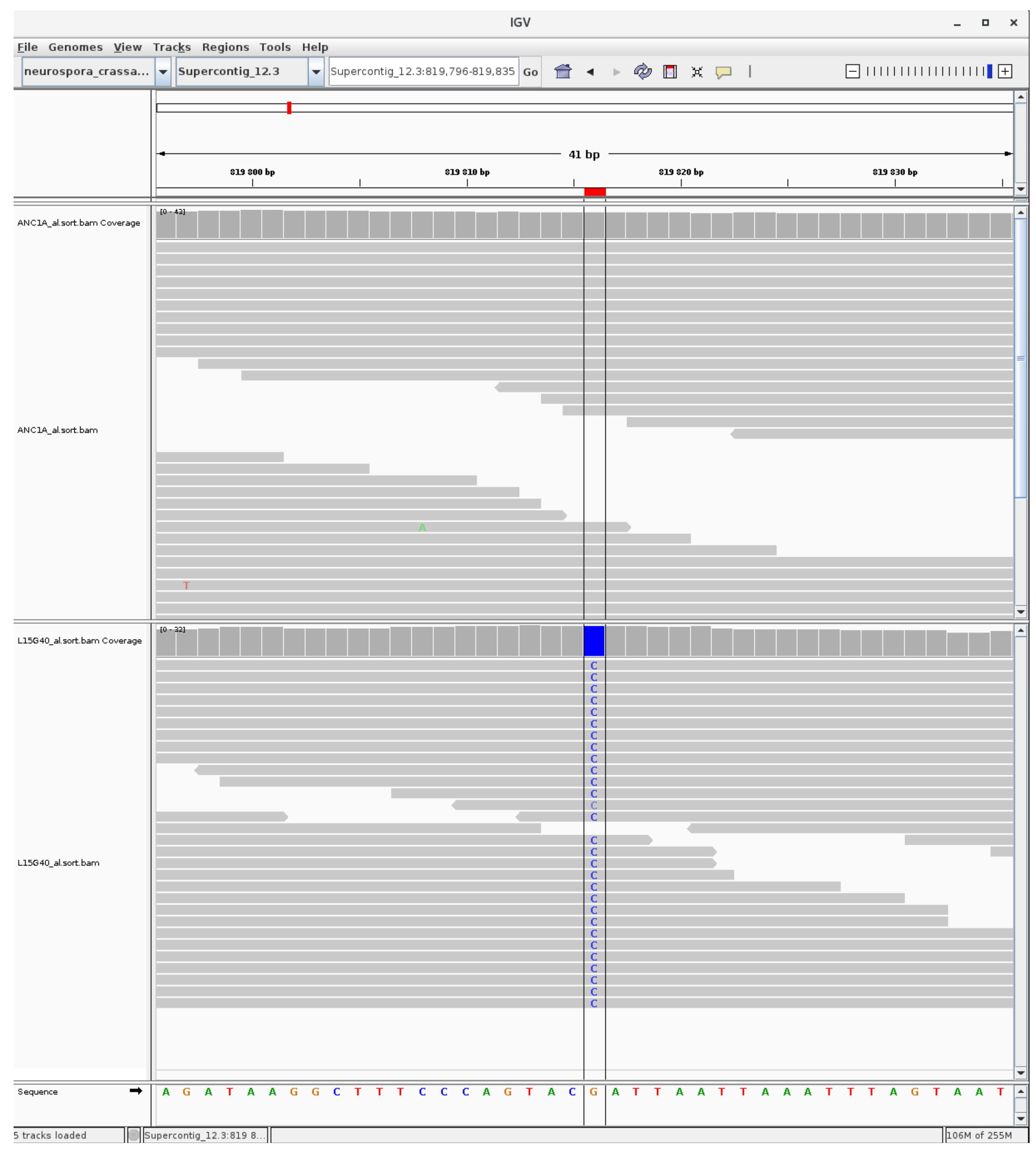

### mutation_centromer_25.jpg

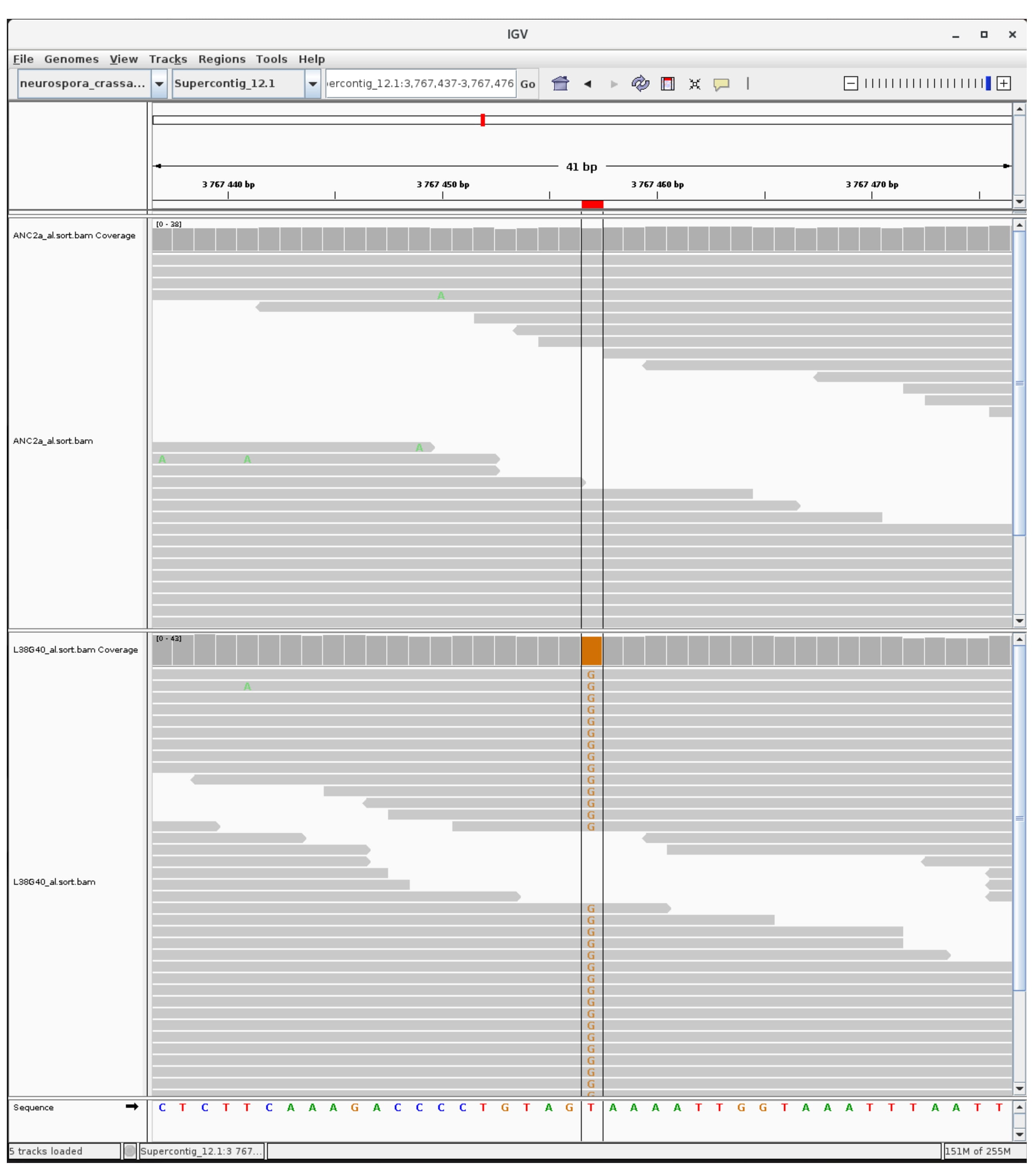

### mutation_centromer_26.jpg

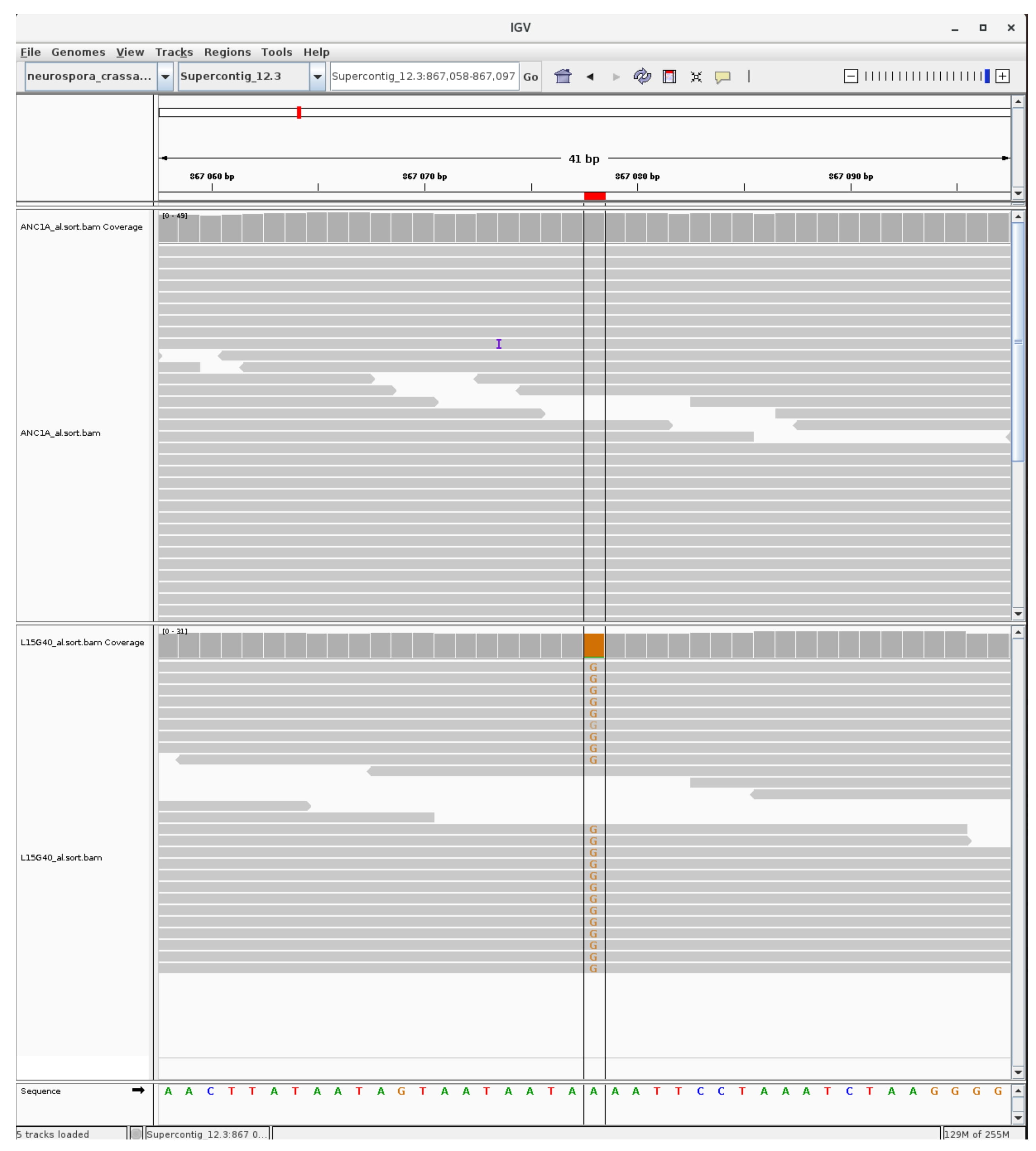

### mutation_centromer_27.jpg

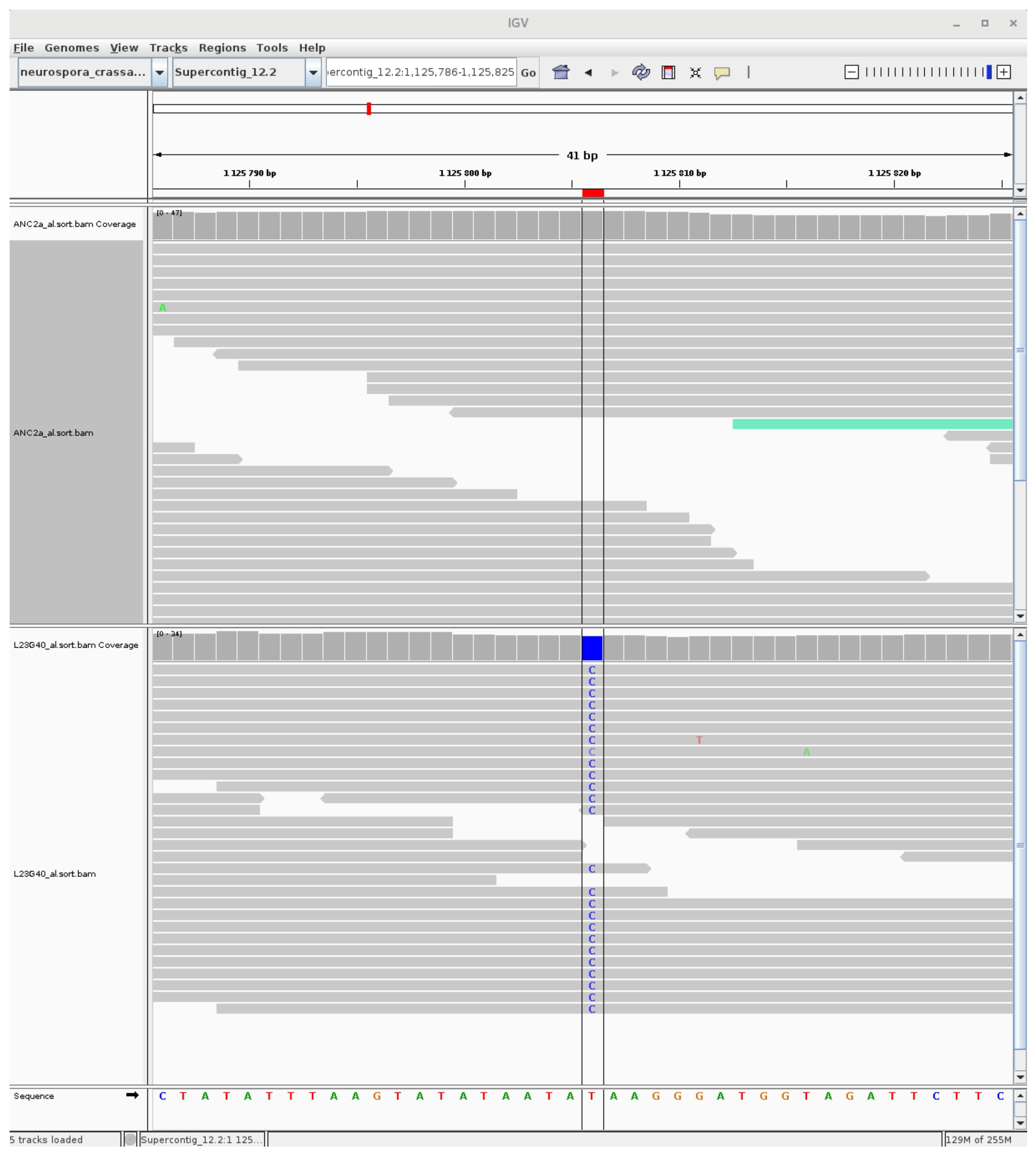

### mutation_centromer_28.jpg

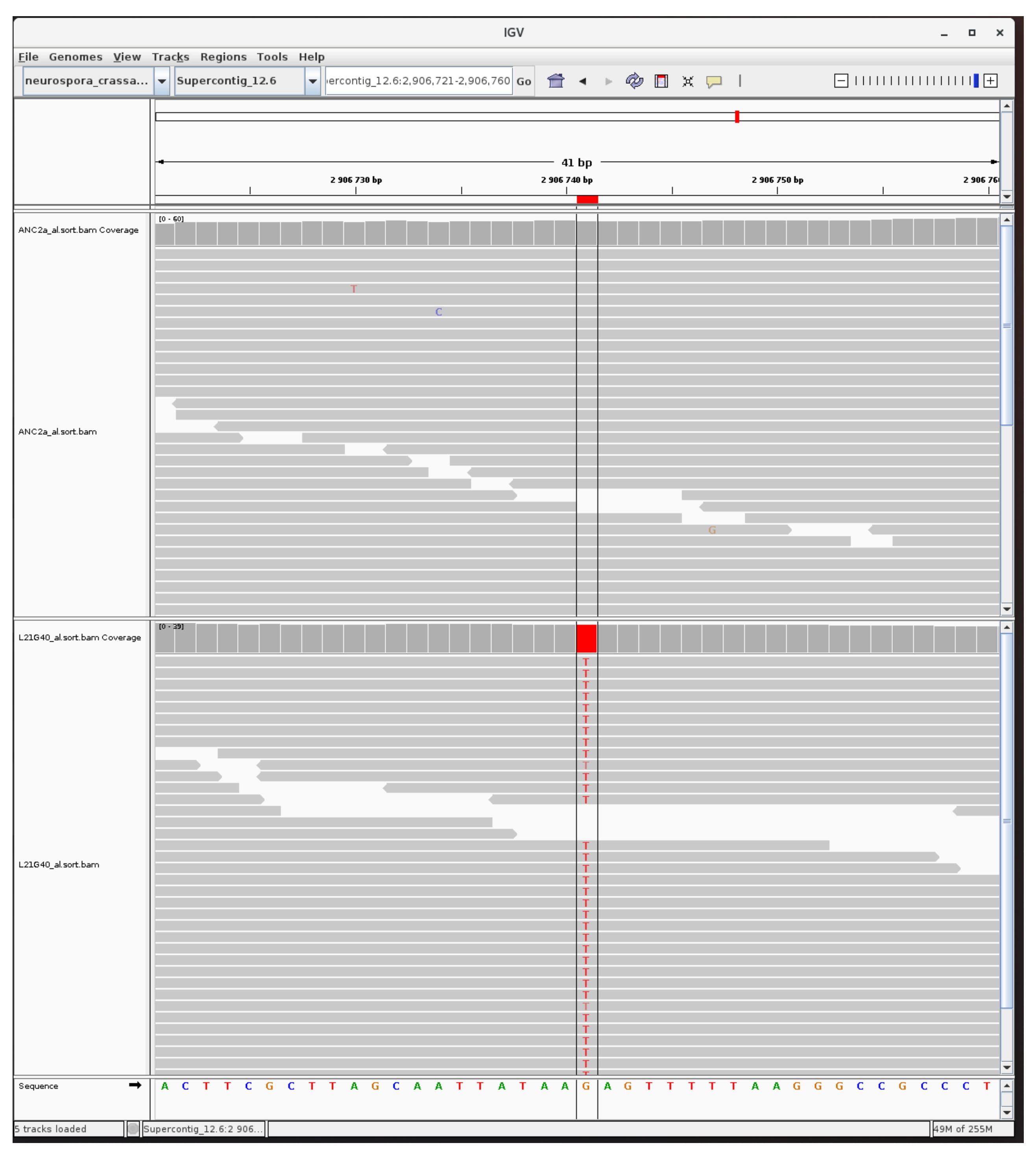

### mutation_centromer_29.jpg

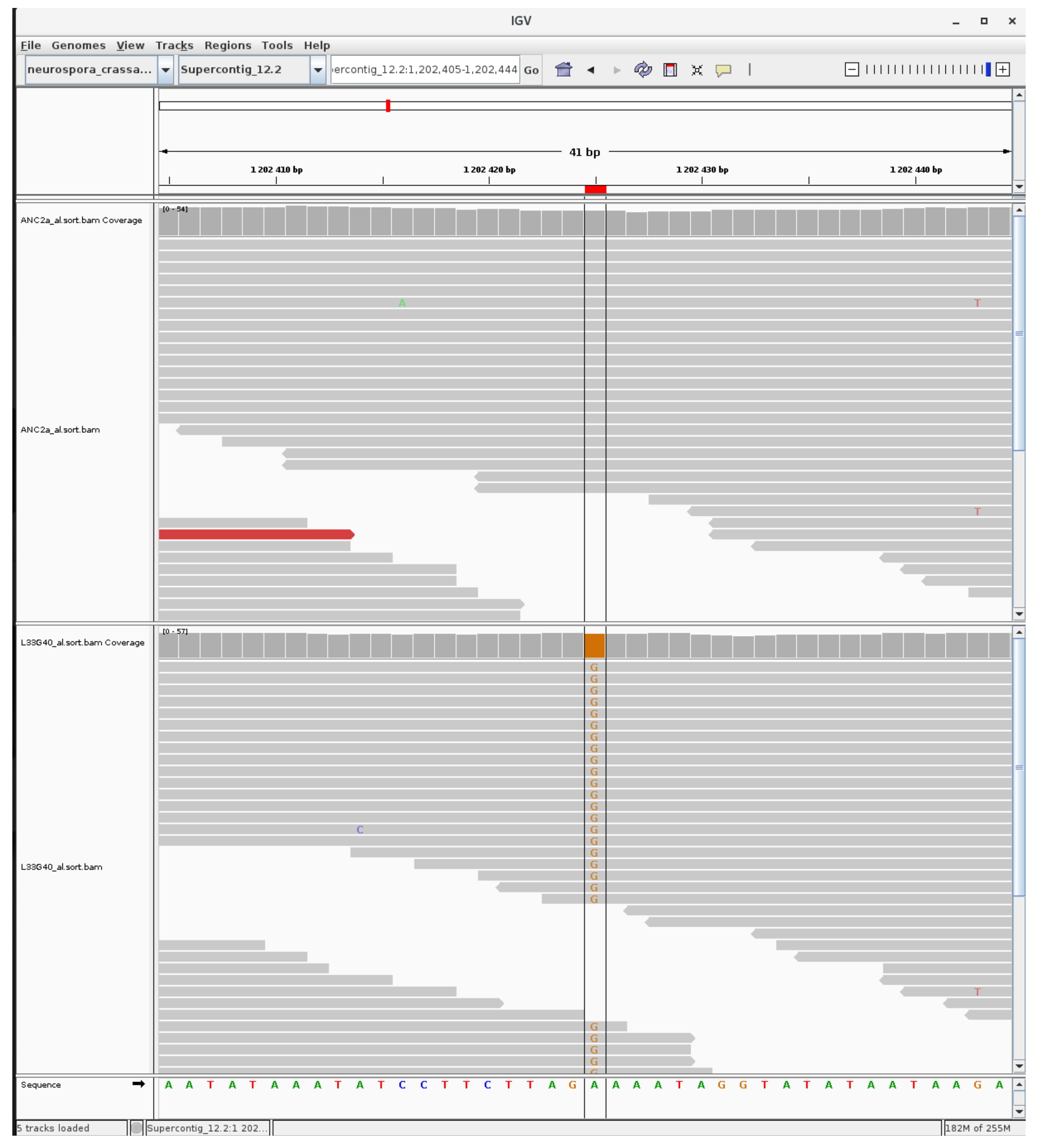

### mutation_centromer_30.jpg

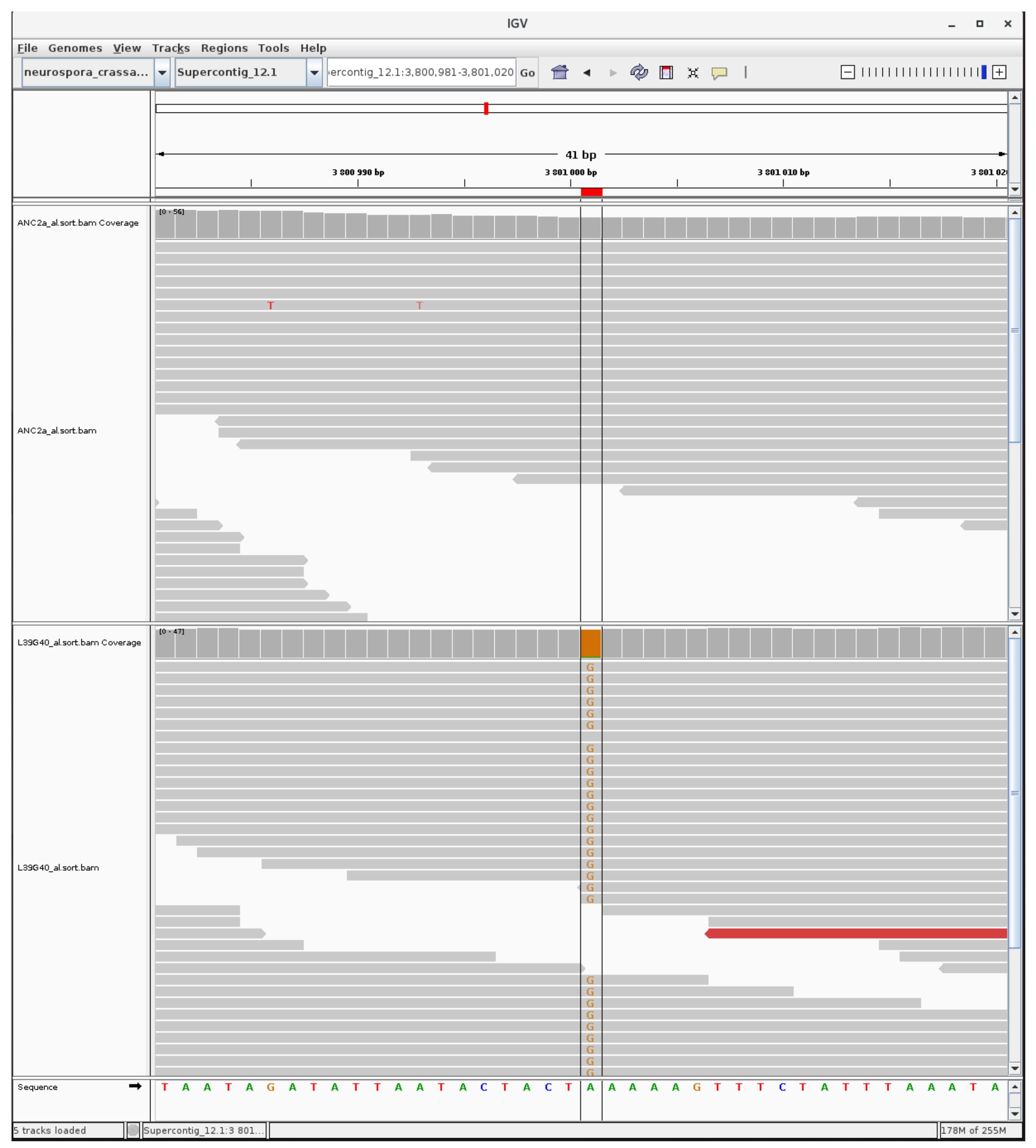

### mutation_euchromatic_1.jpg

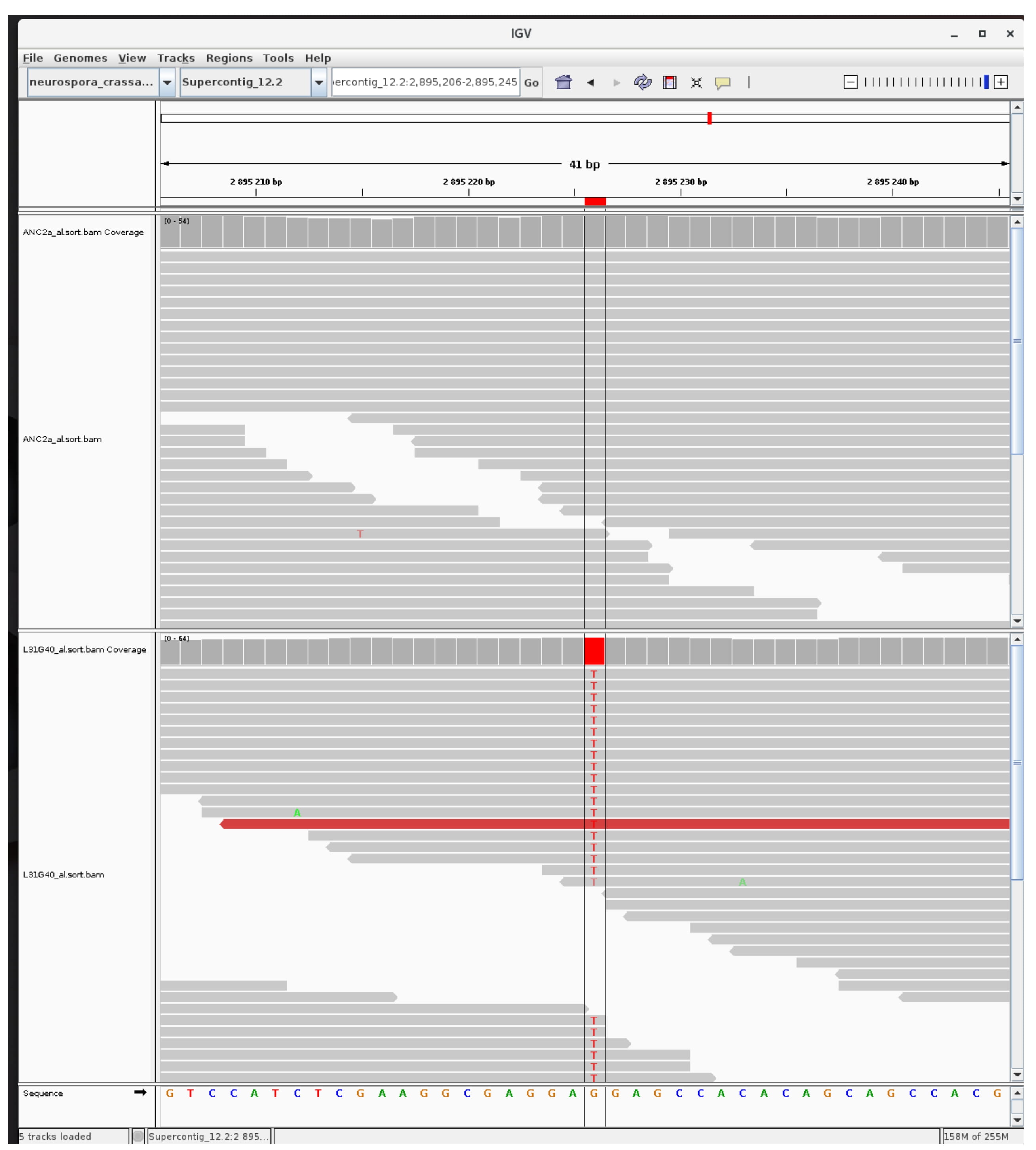

### mutation_euchromatic_2.jpg

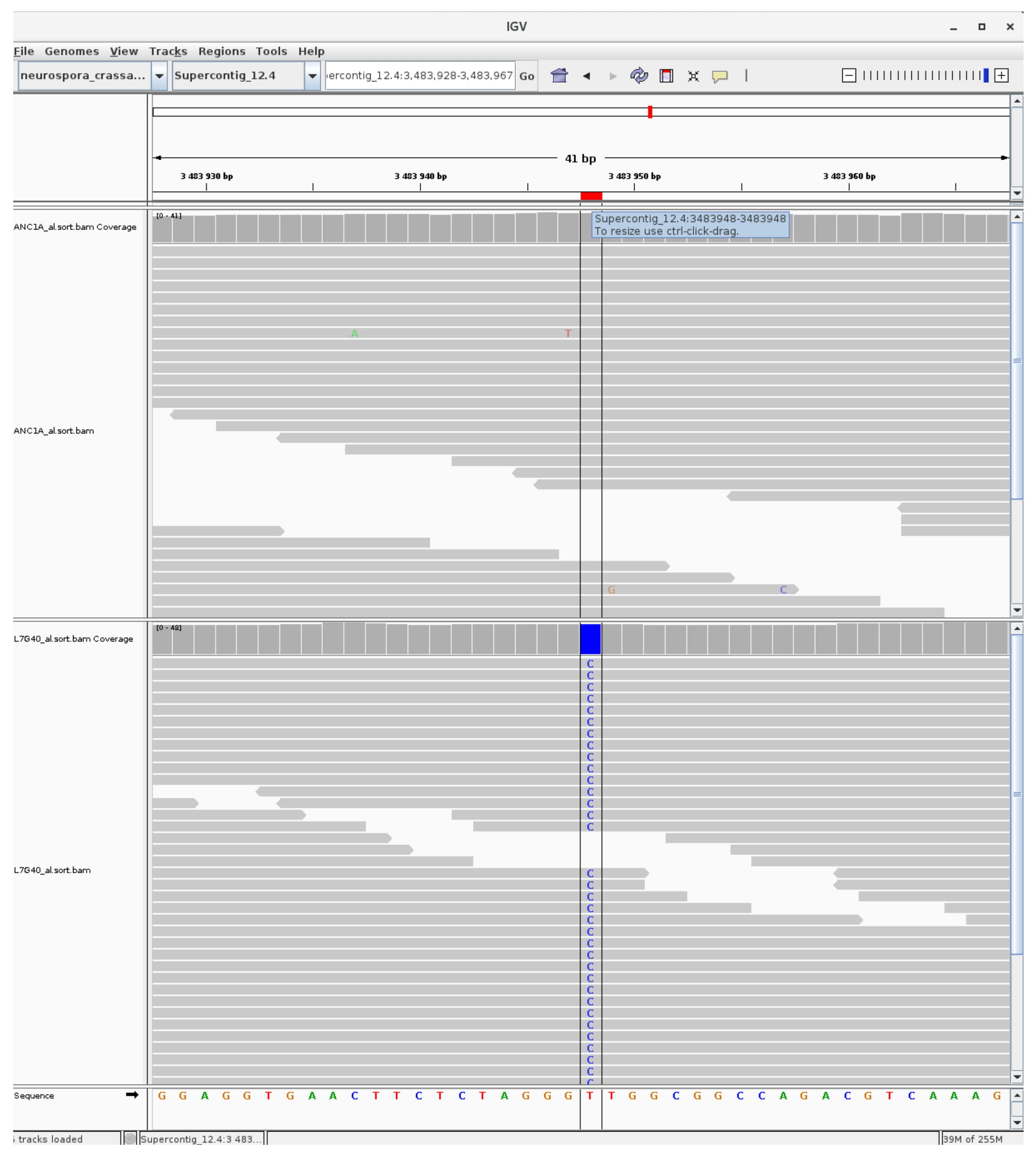

### mutation_euchromatic_3.jpg

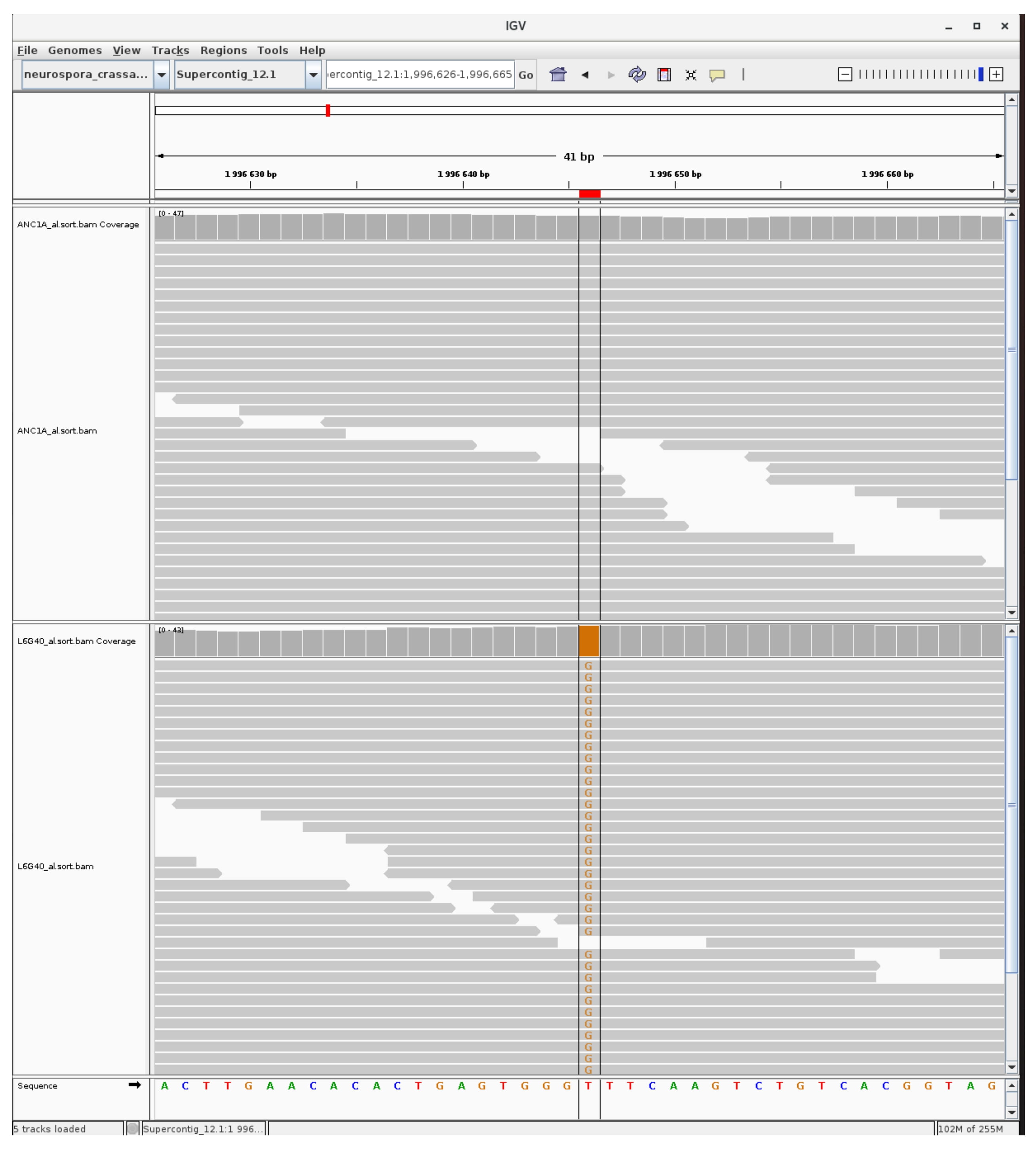

### mutation_euchromatic_4.jpg

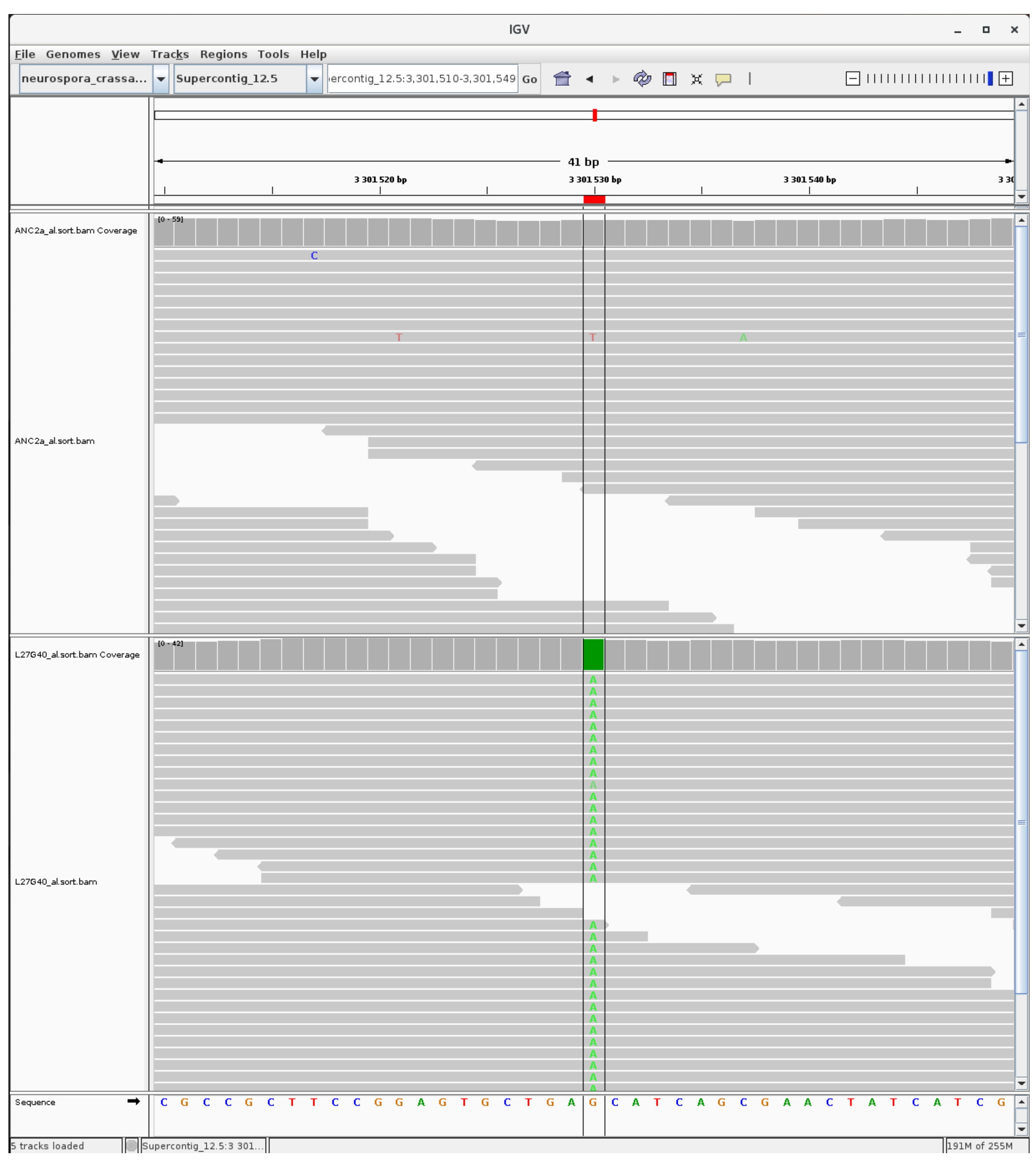

### mutation_euchromatic_5.jpg

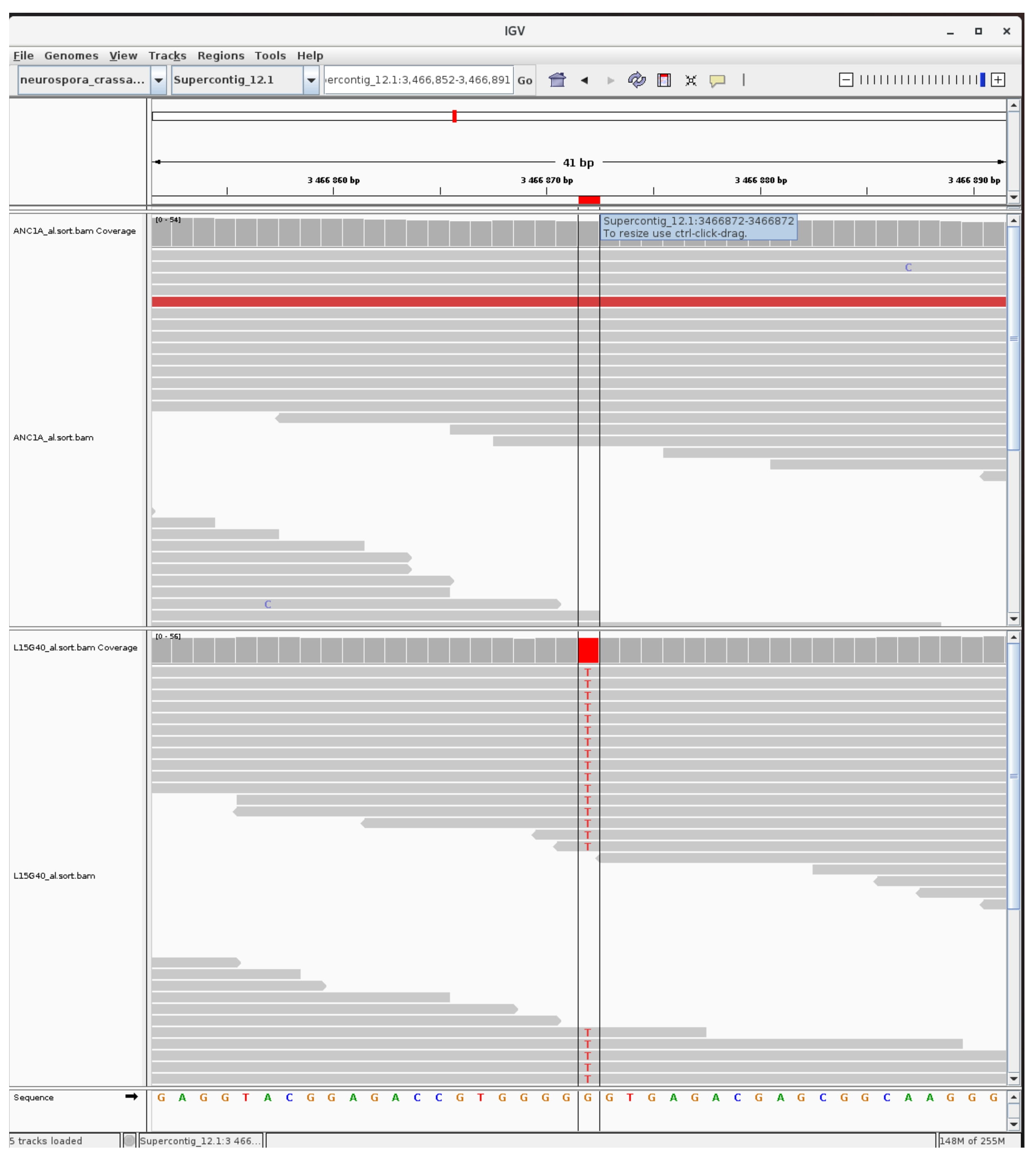

### mutation_euchromatic_6.jpg

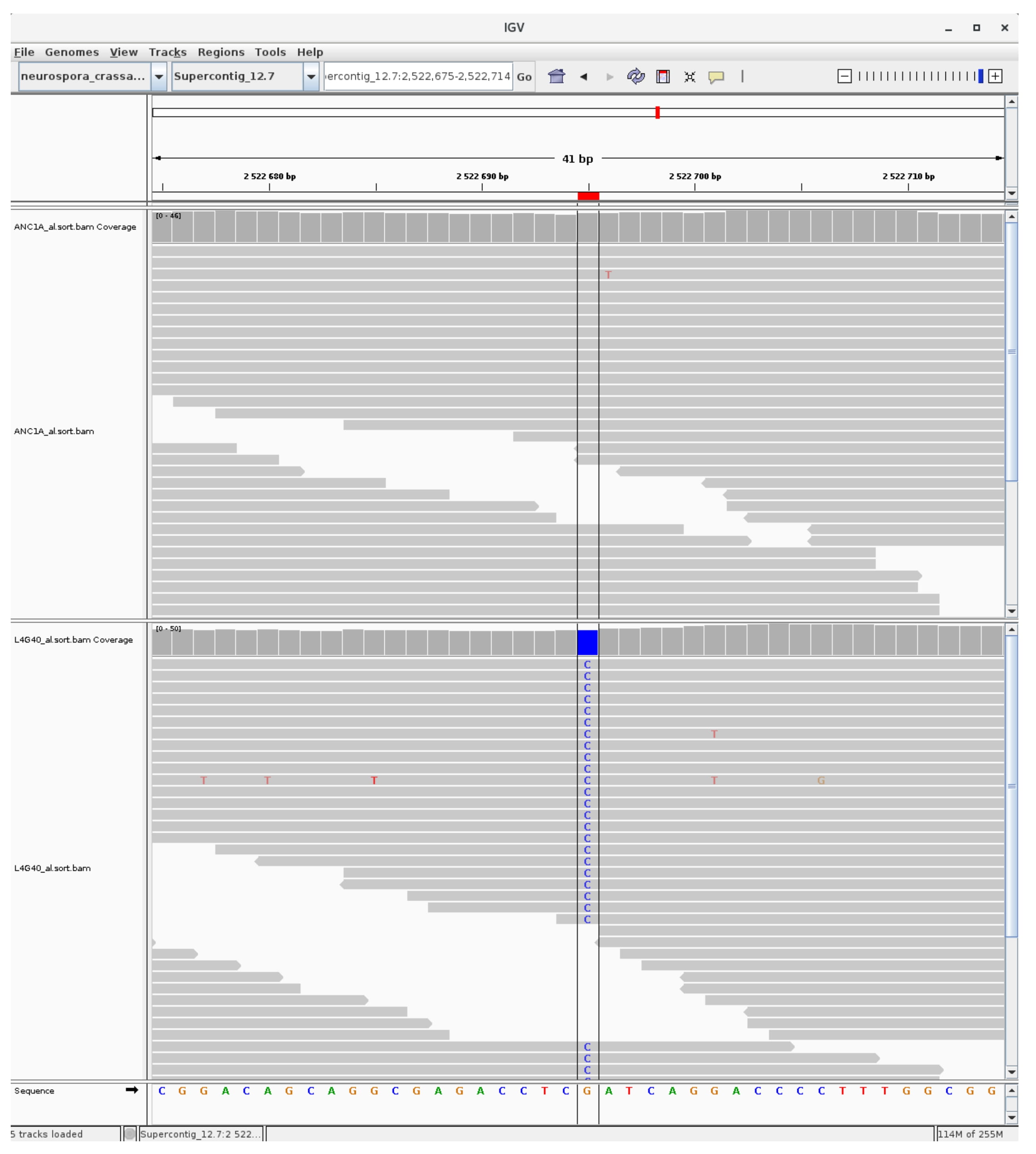

### mutation_euchromatic_7.jpg

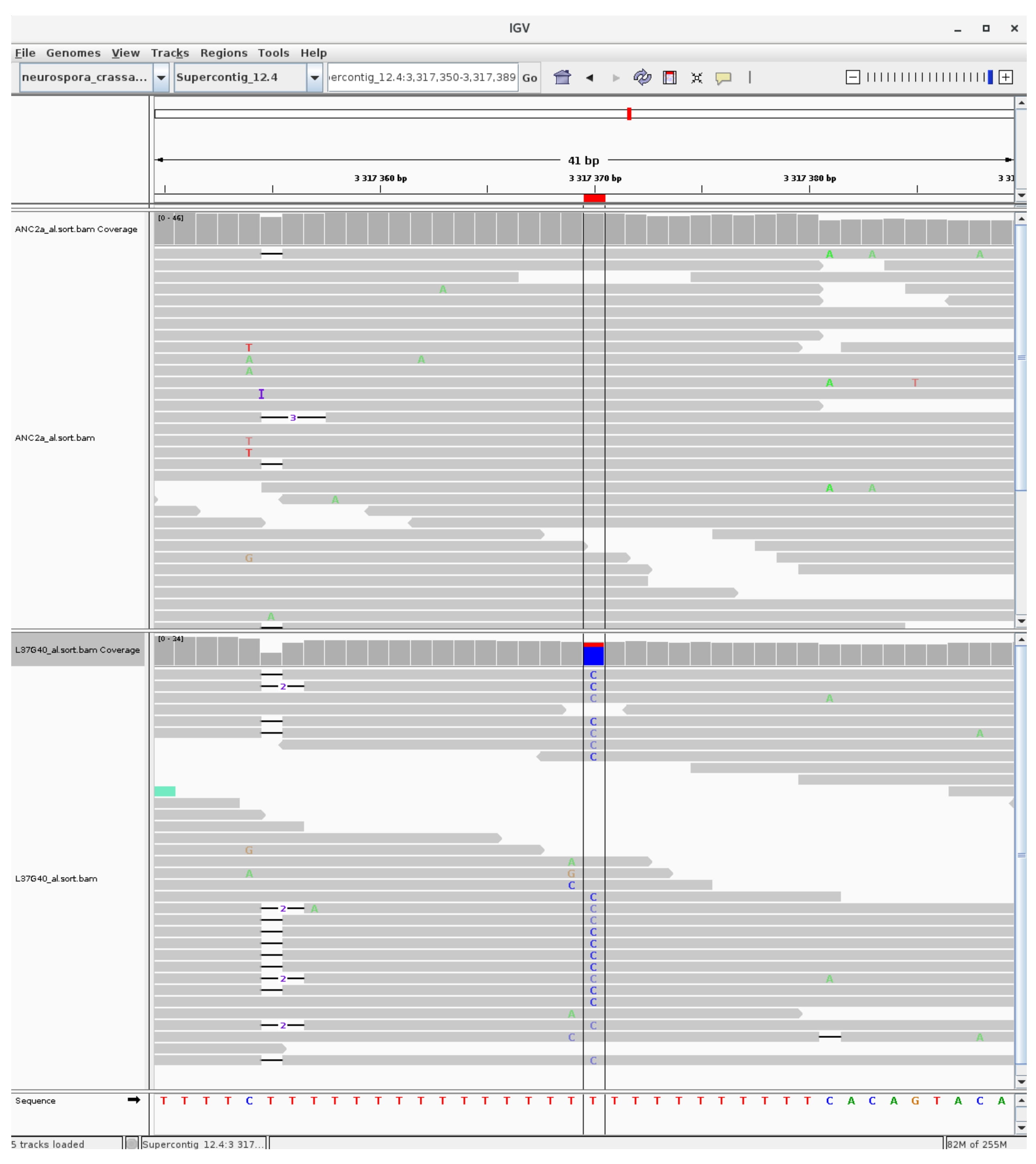

### mutation_euchromatic_8.jpg

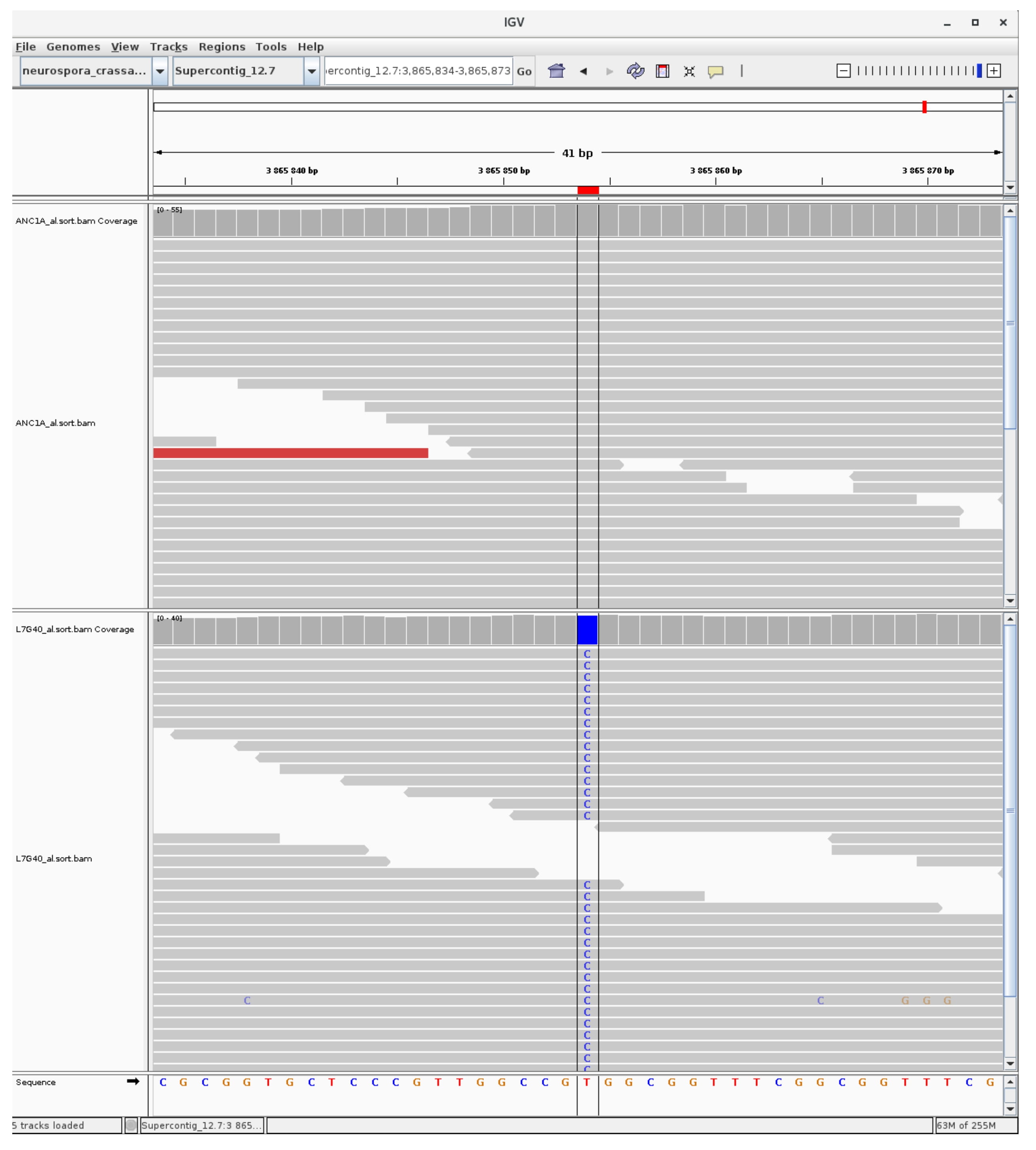

### mutation_euchromatic_9.jpg

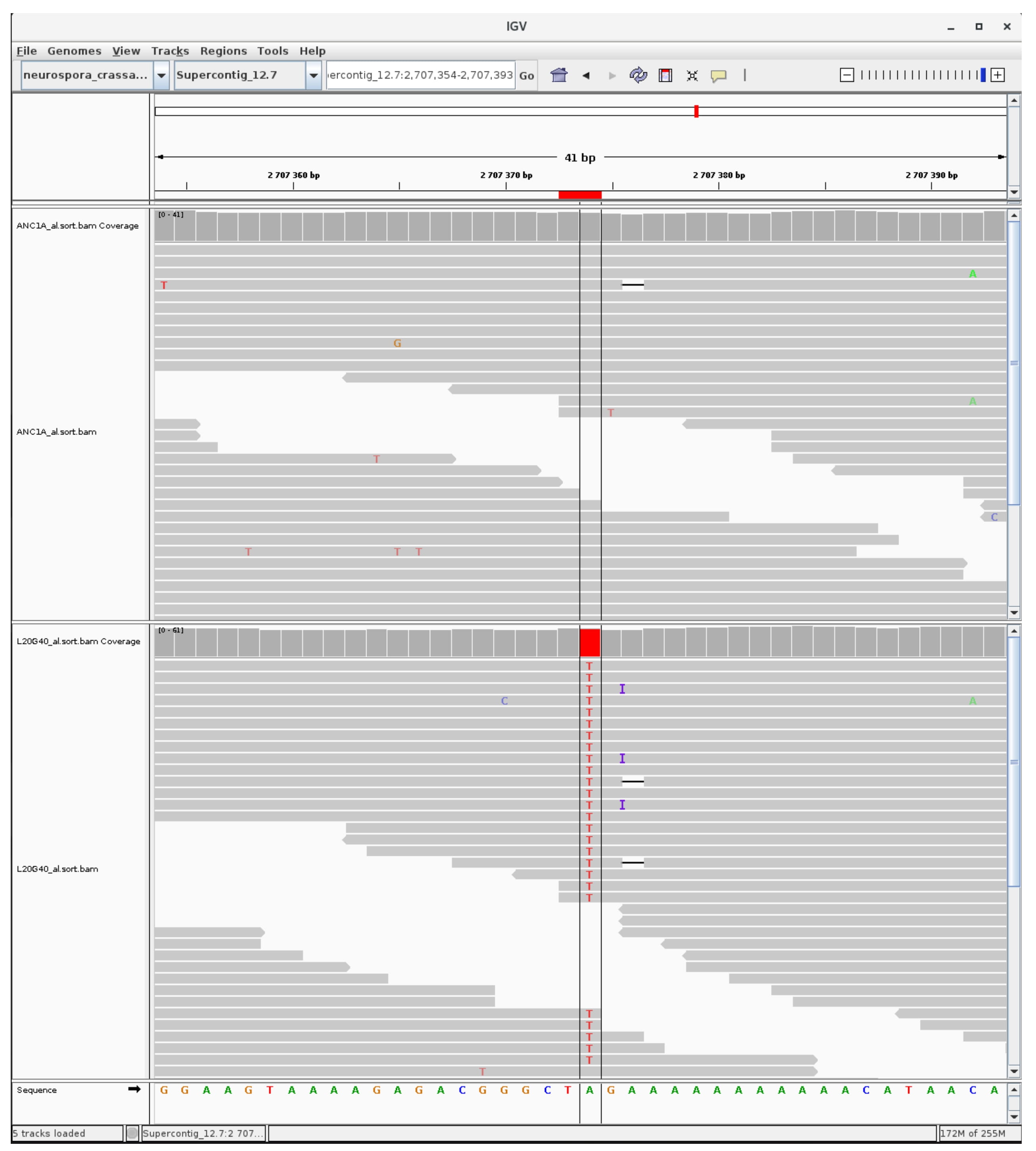

### mutation_euchromatic_10.jpg

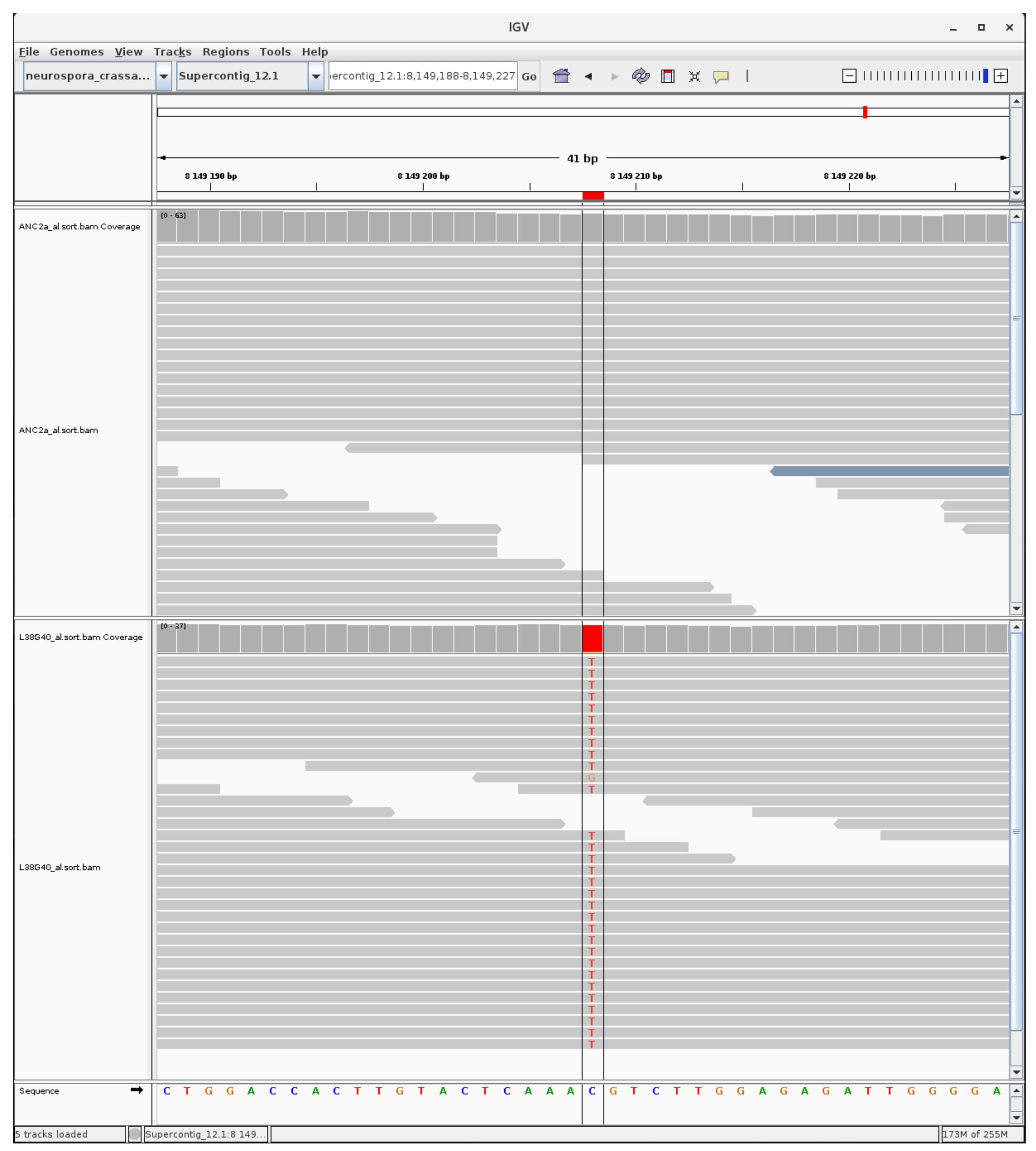

### mutation_euchromatic_11.jpg

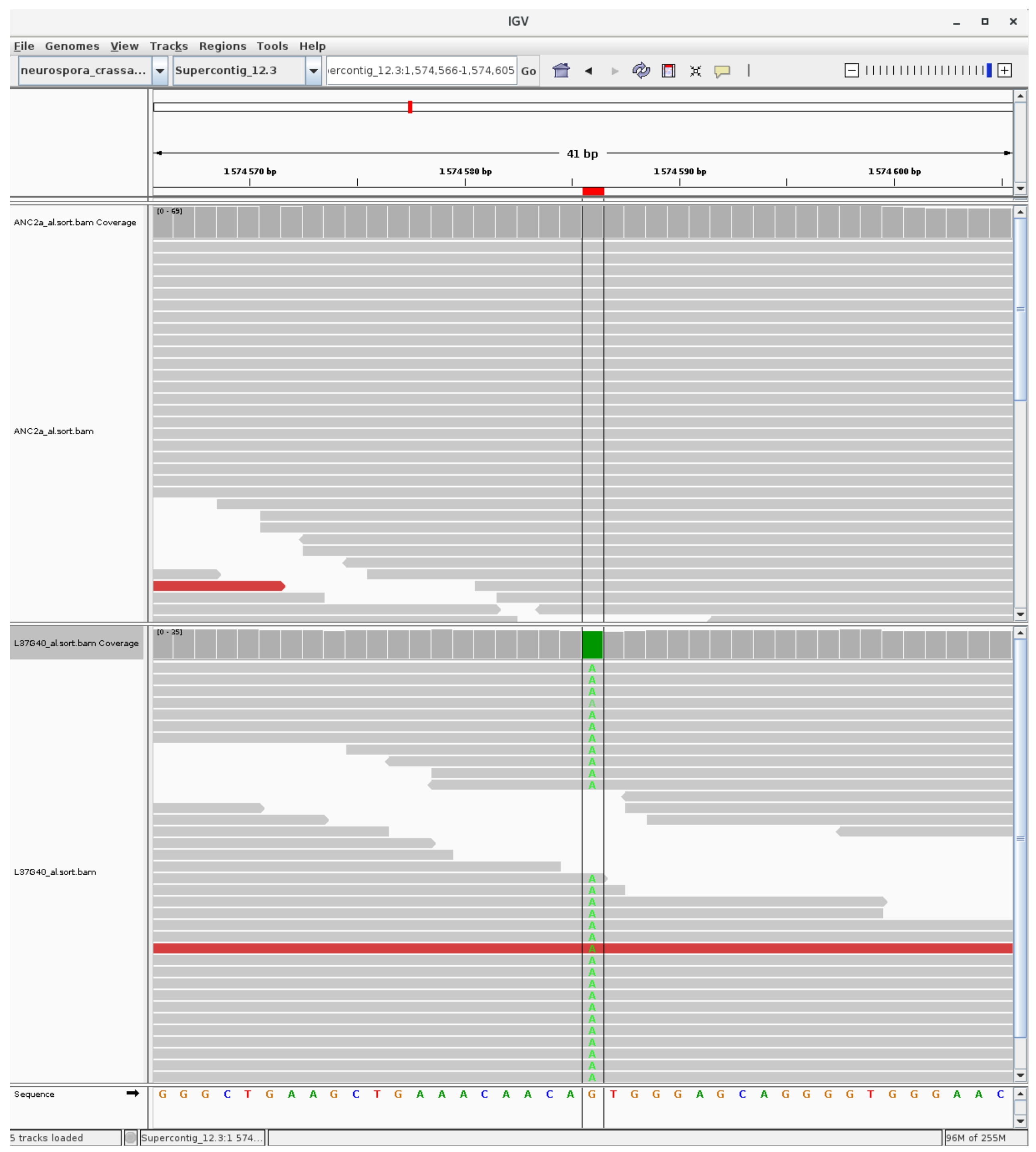

### mutation_euchromatic_12.jpg

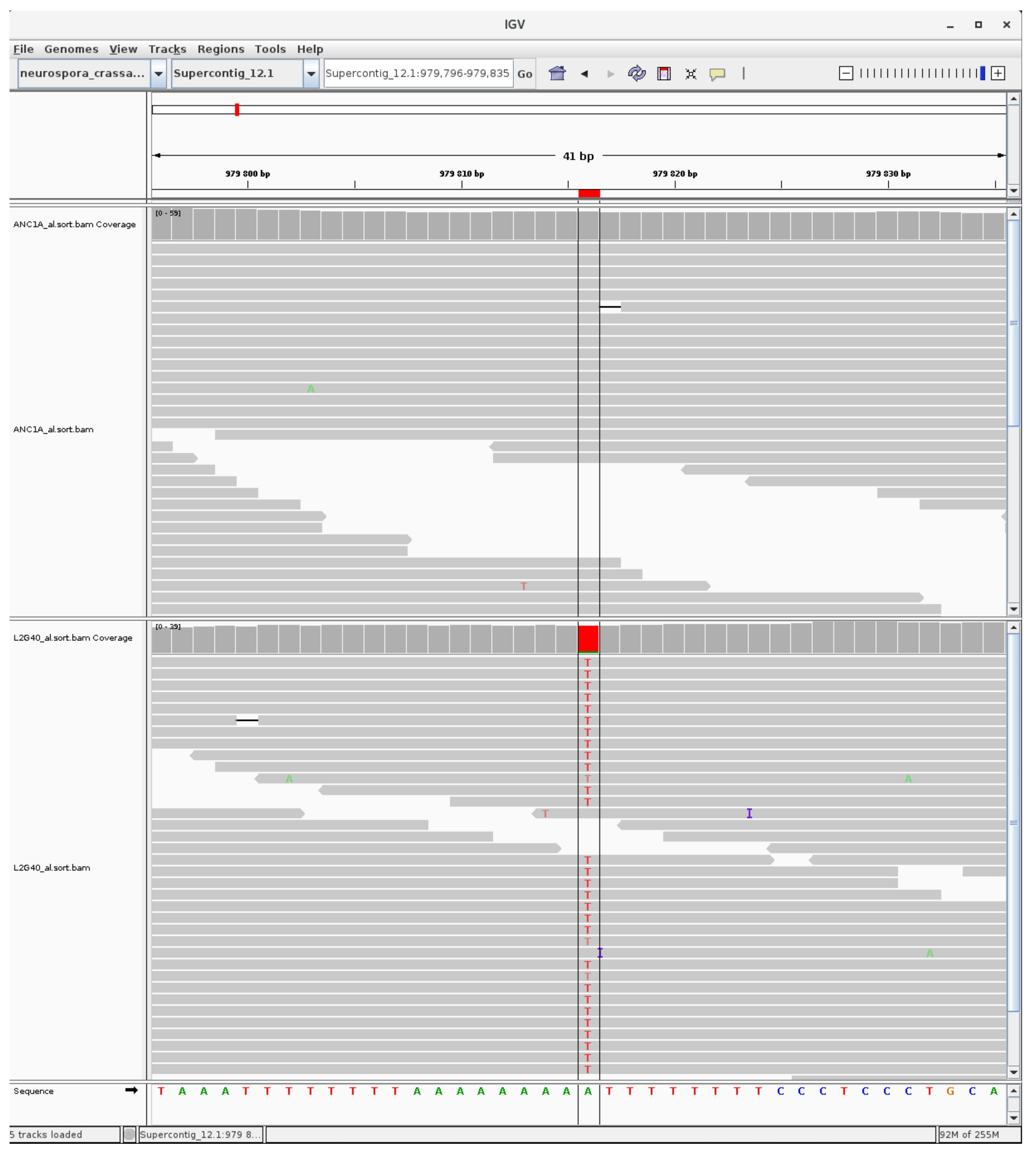

### mutation_euchromatic_13.jpg

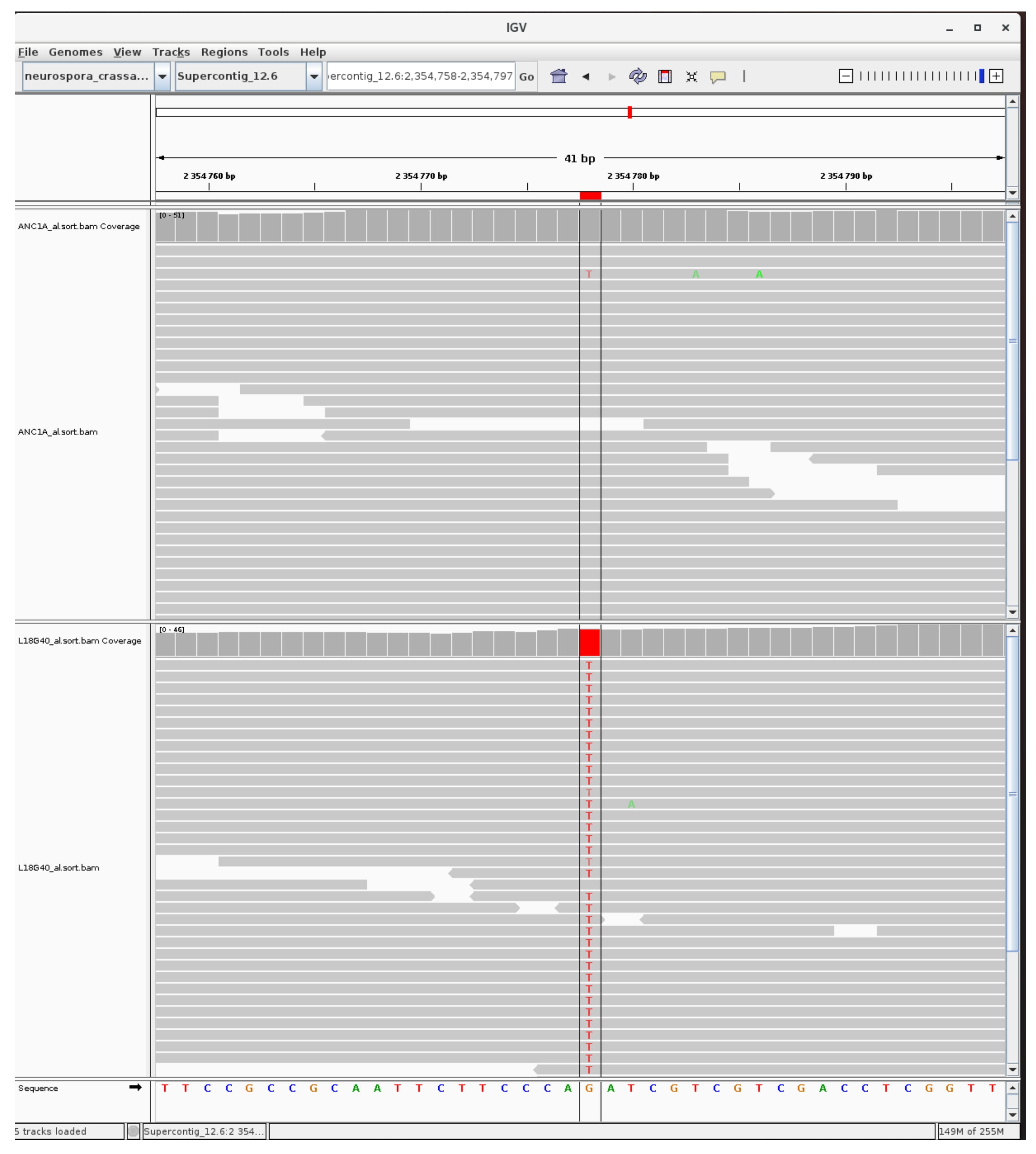

### mutation_euchromatic_14.jpg

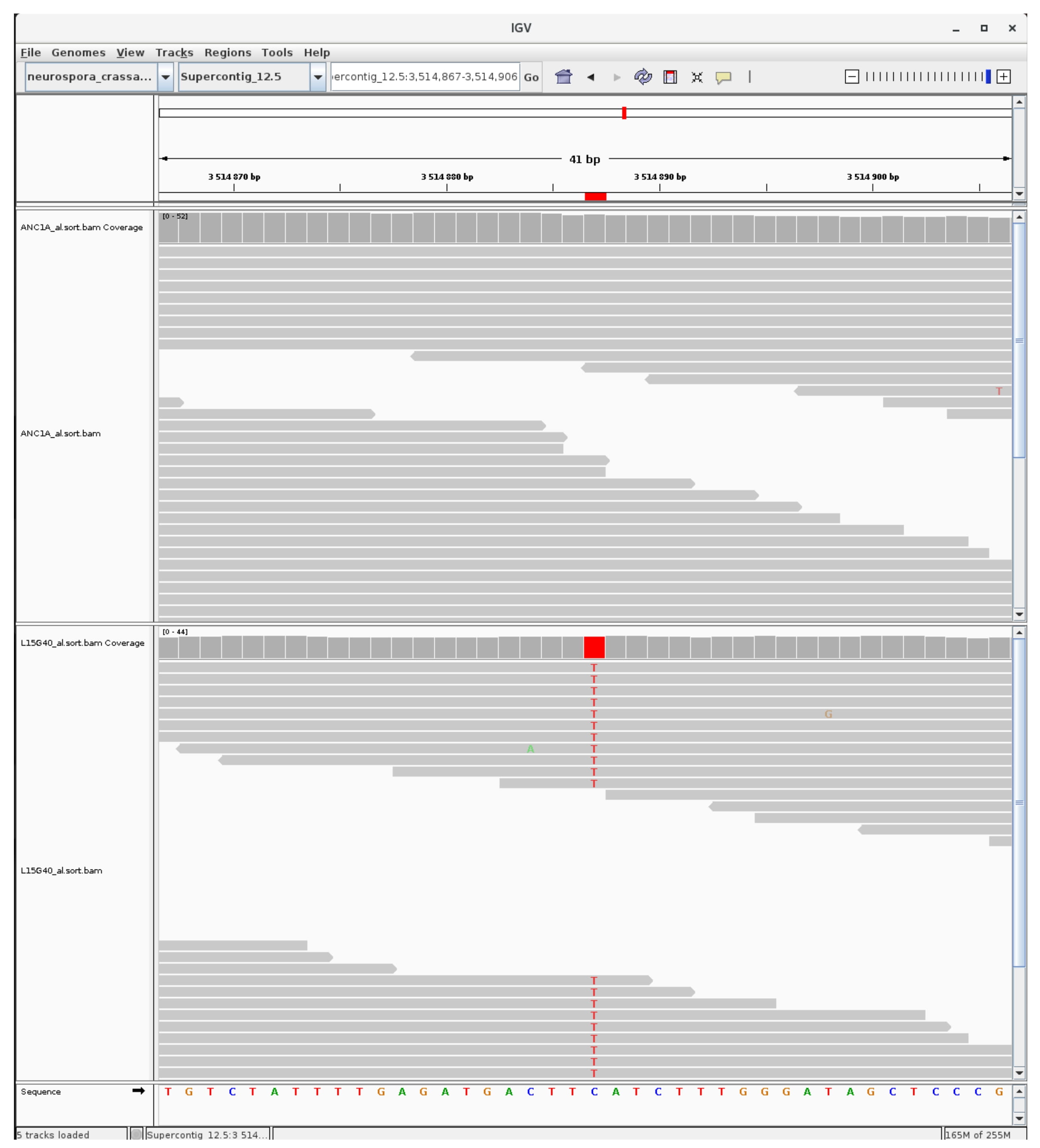

### mutation_euchromatic_15.jpg

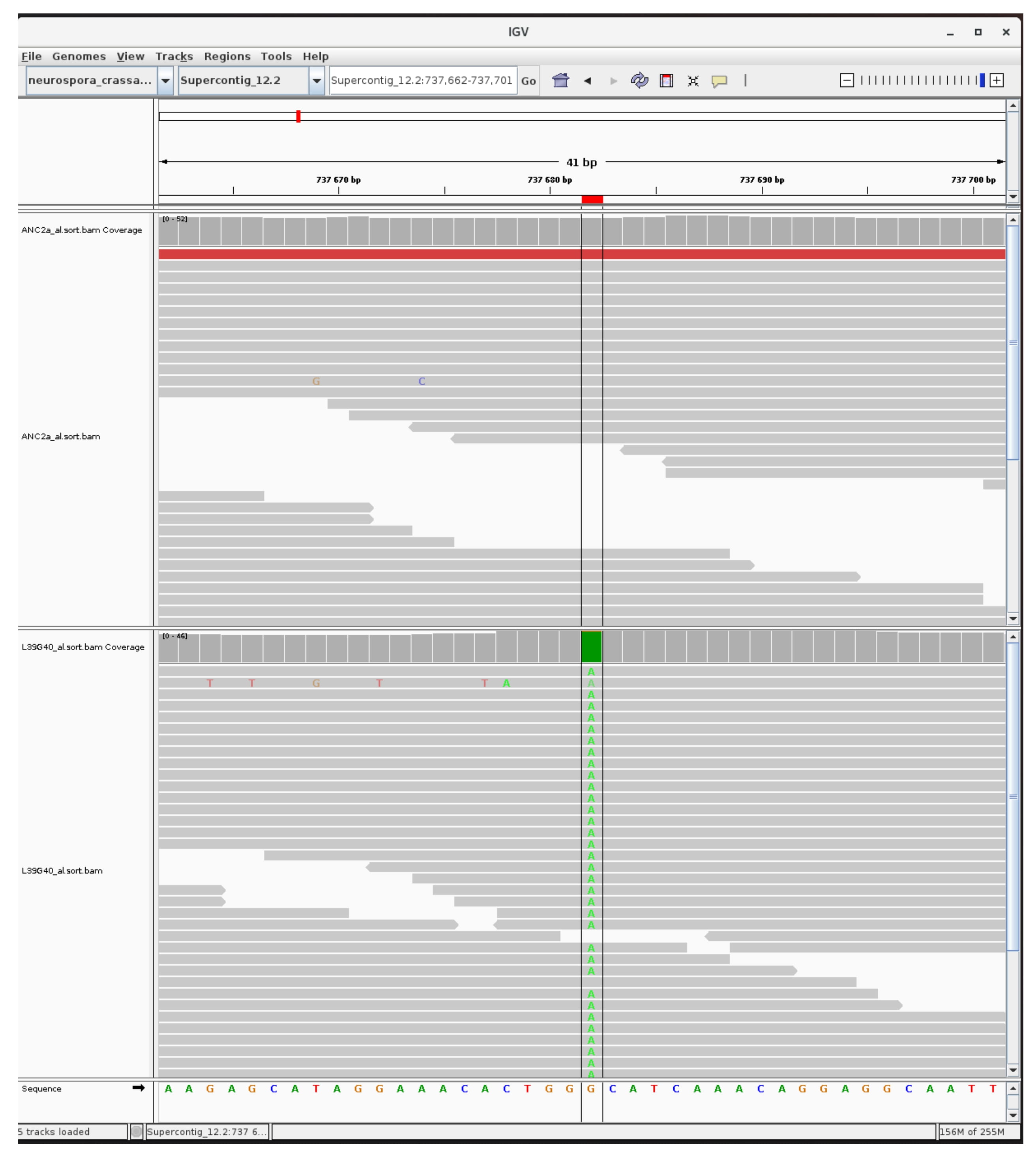

### mutation_euchromatic_16.jpg

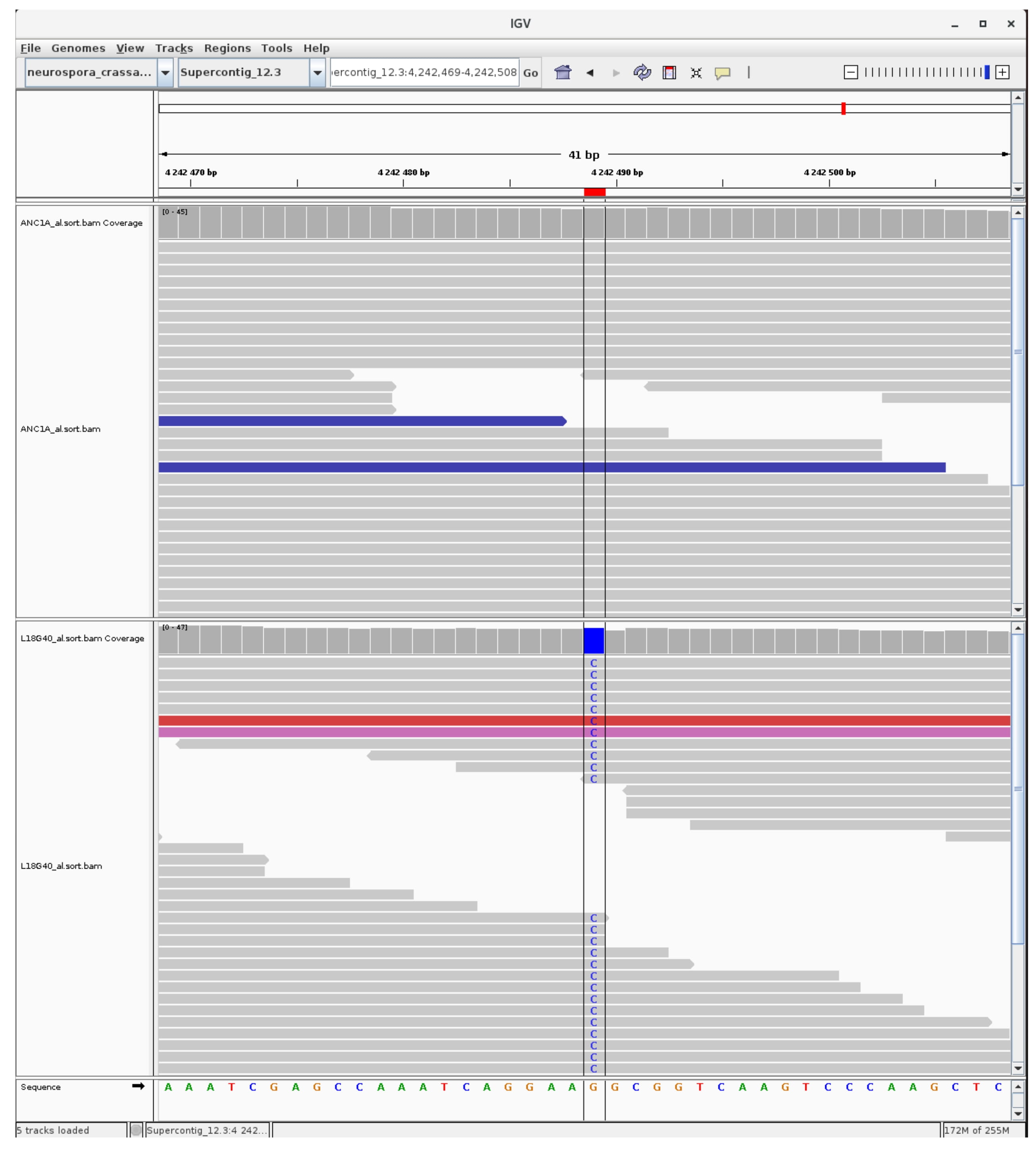

### mutation_euchromatic_17.jpg

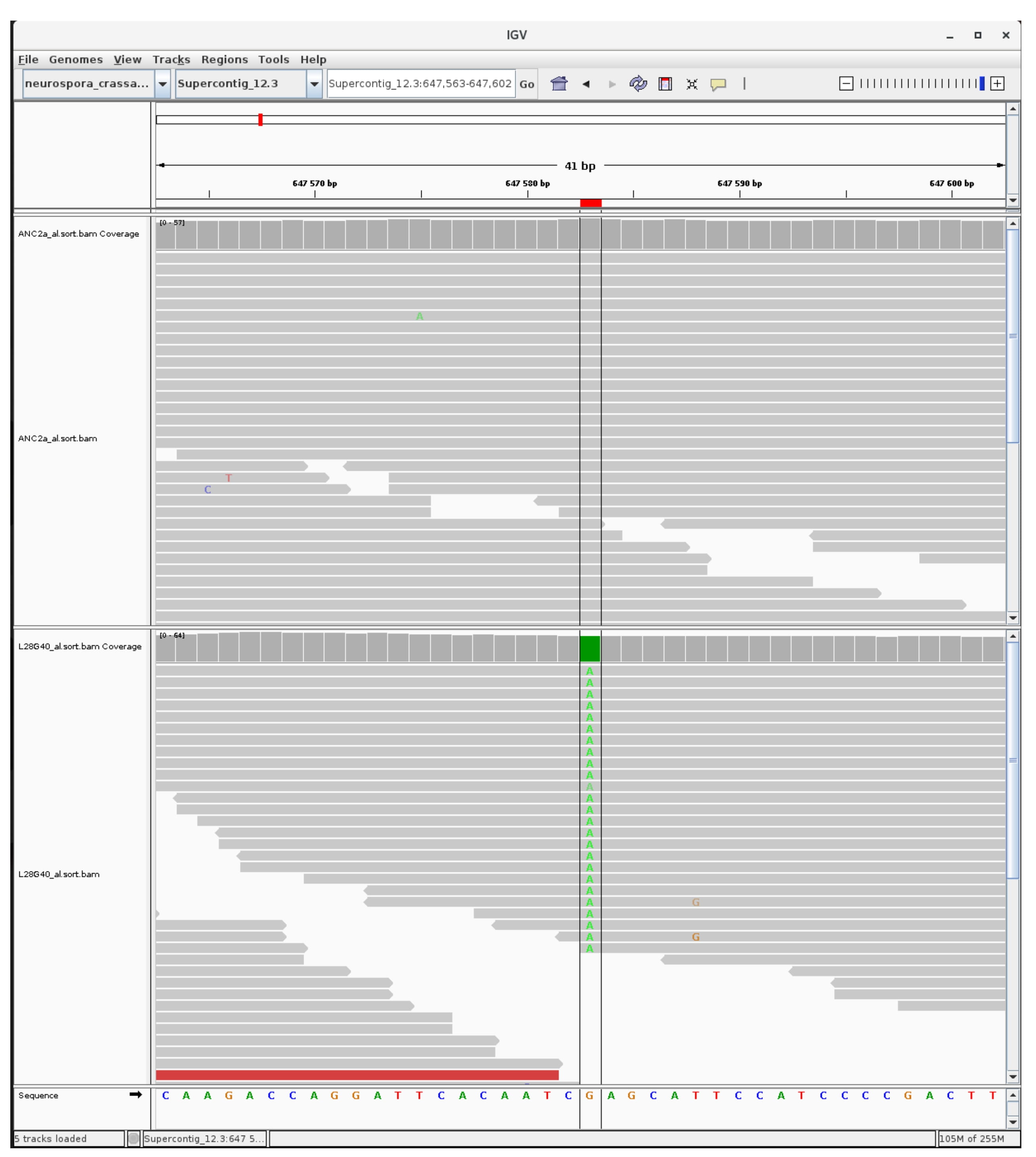

### mutation_euchromatic_18.jpg

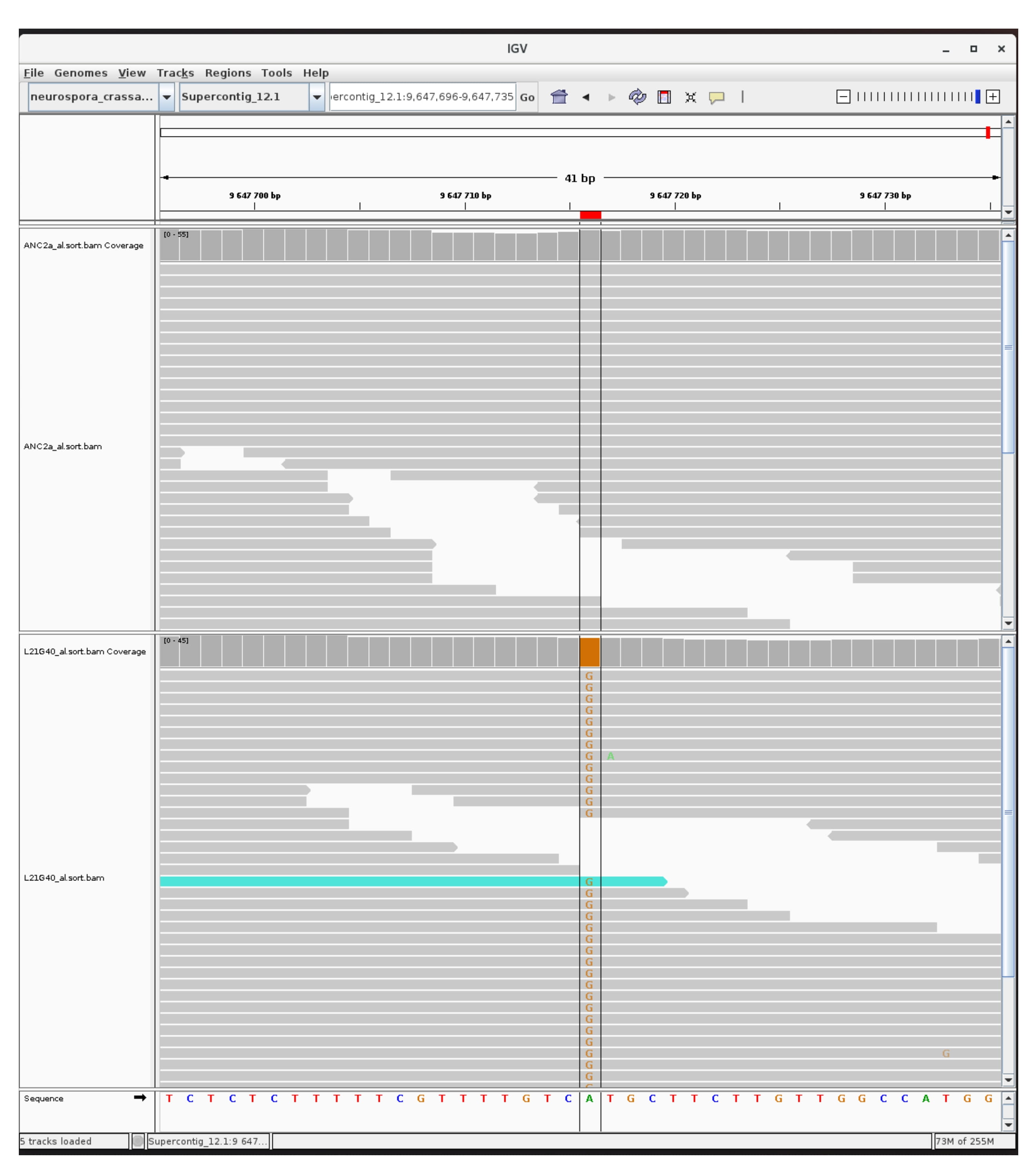

### mutation_euchromatic_19.jpg

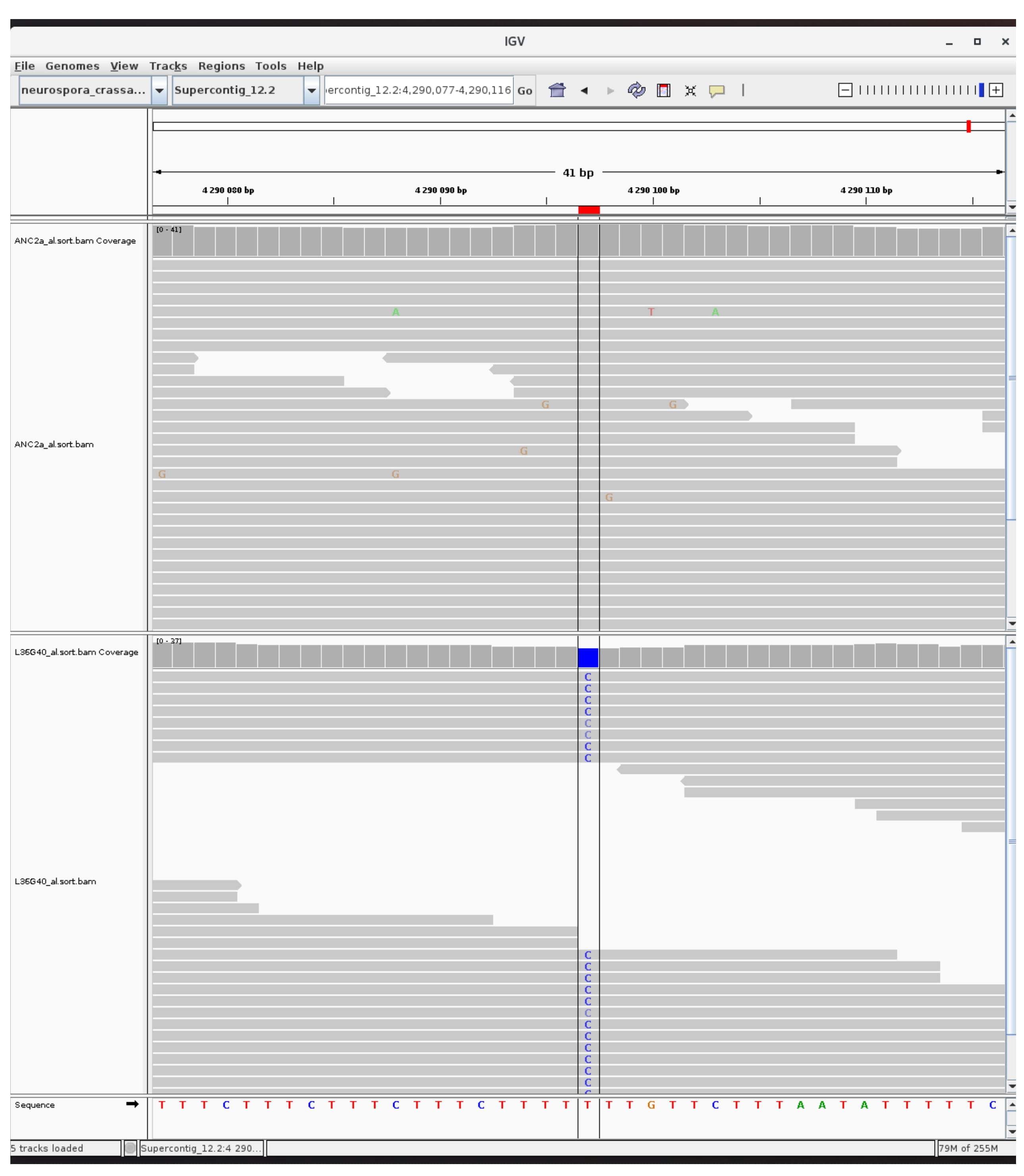

### mutation_euchromatic_20.jpg

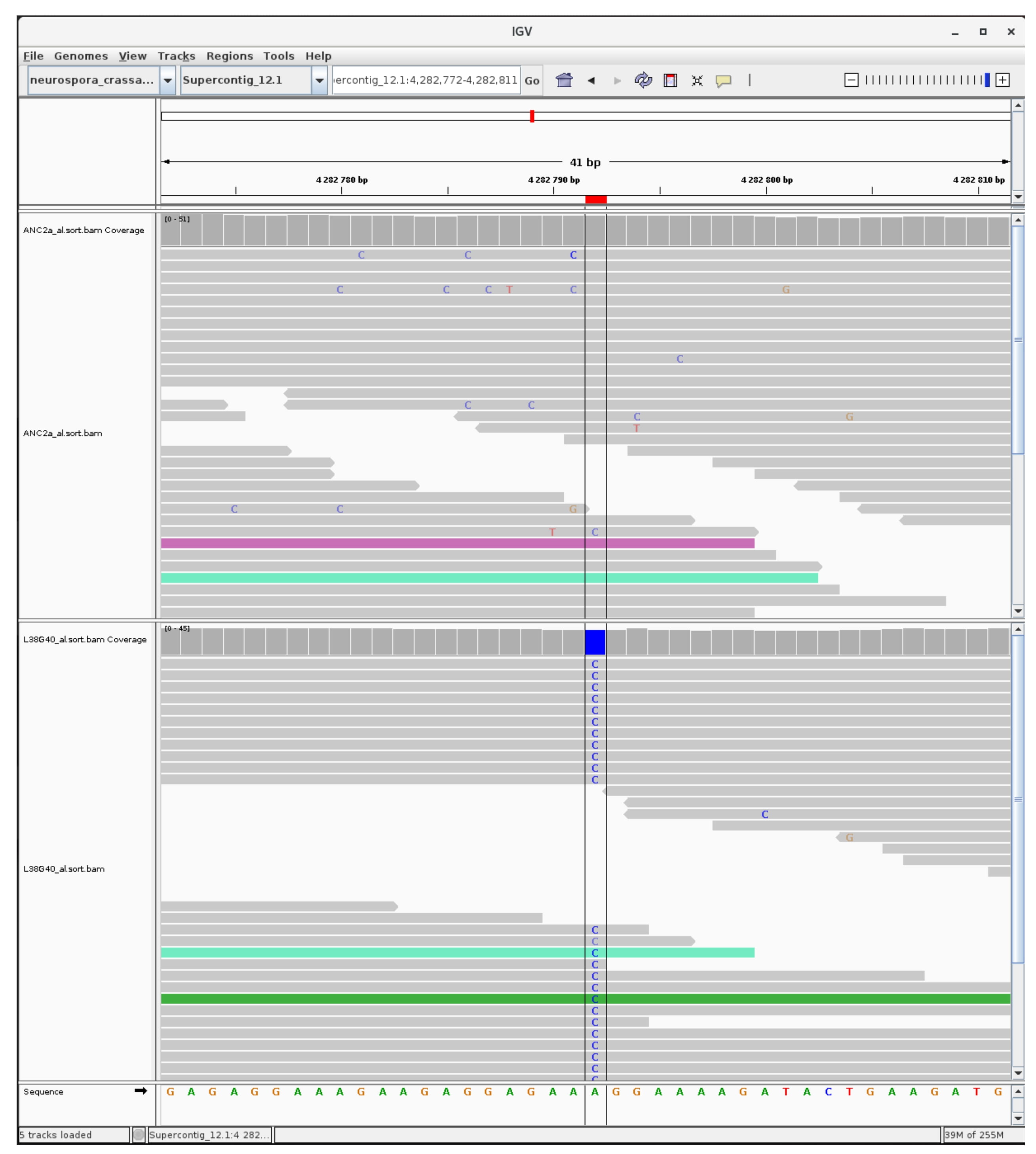

### mutation_euchromatic_21.jpg

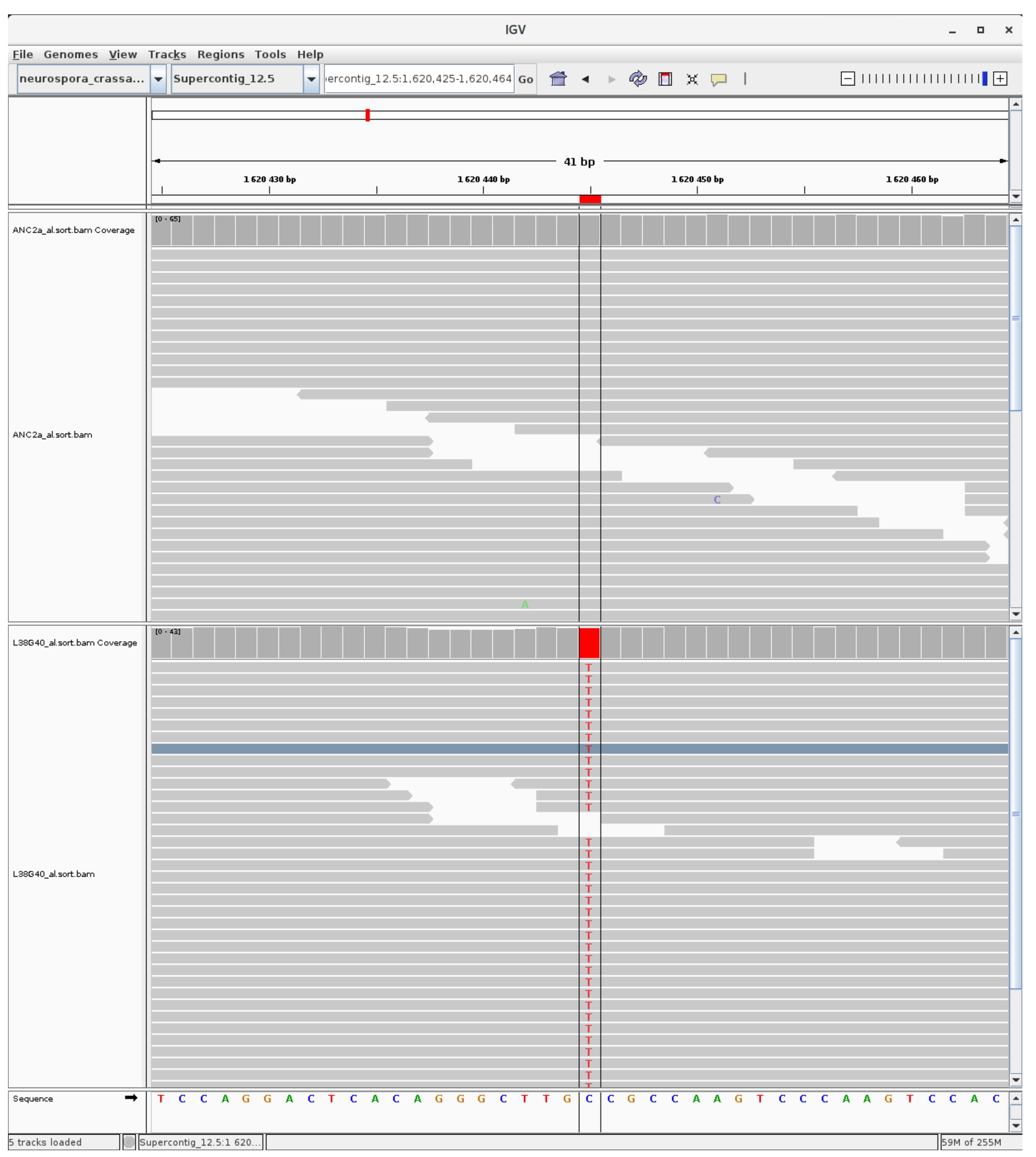

### mutation_euchromatic_22.jpg

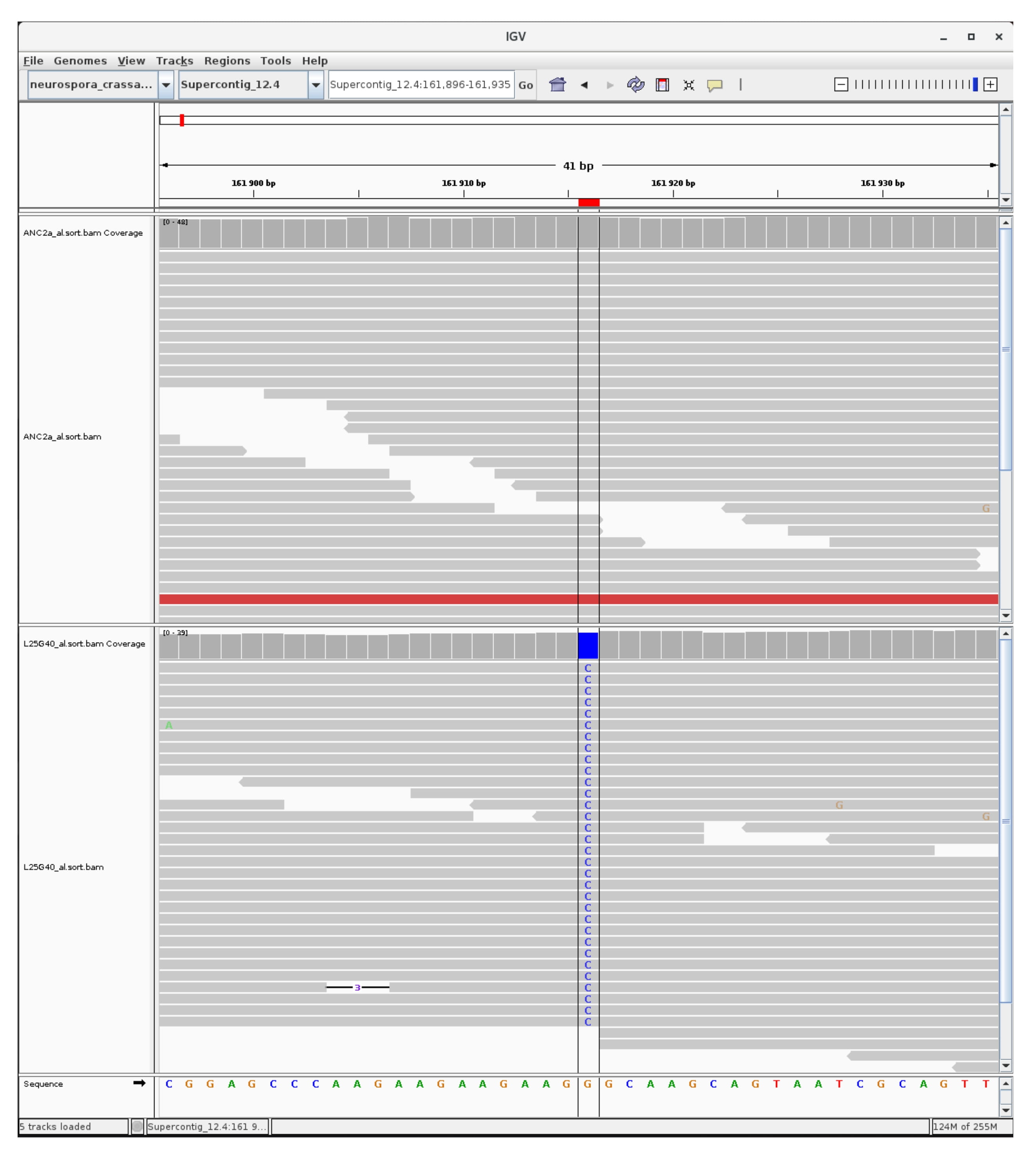

### mutation_euchromatic_23.jpg

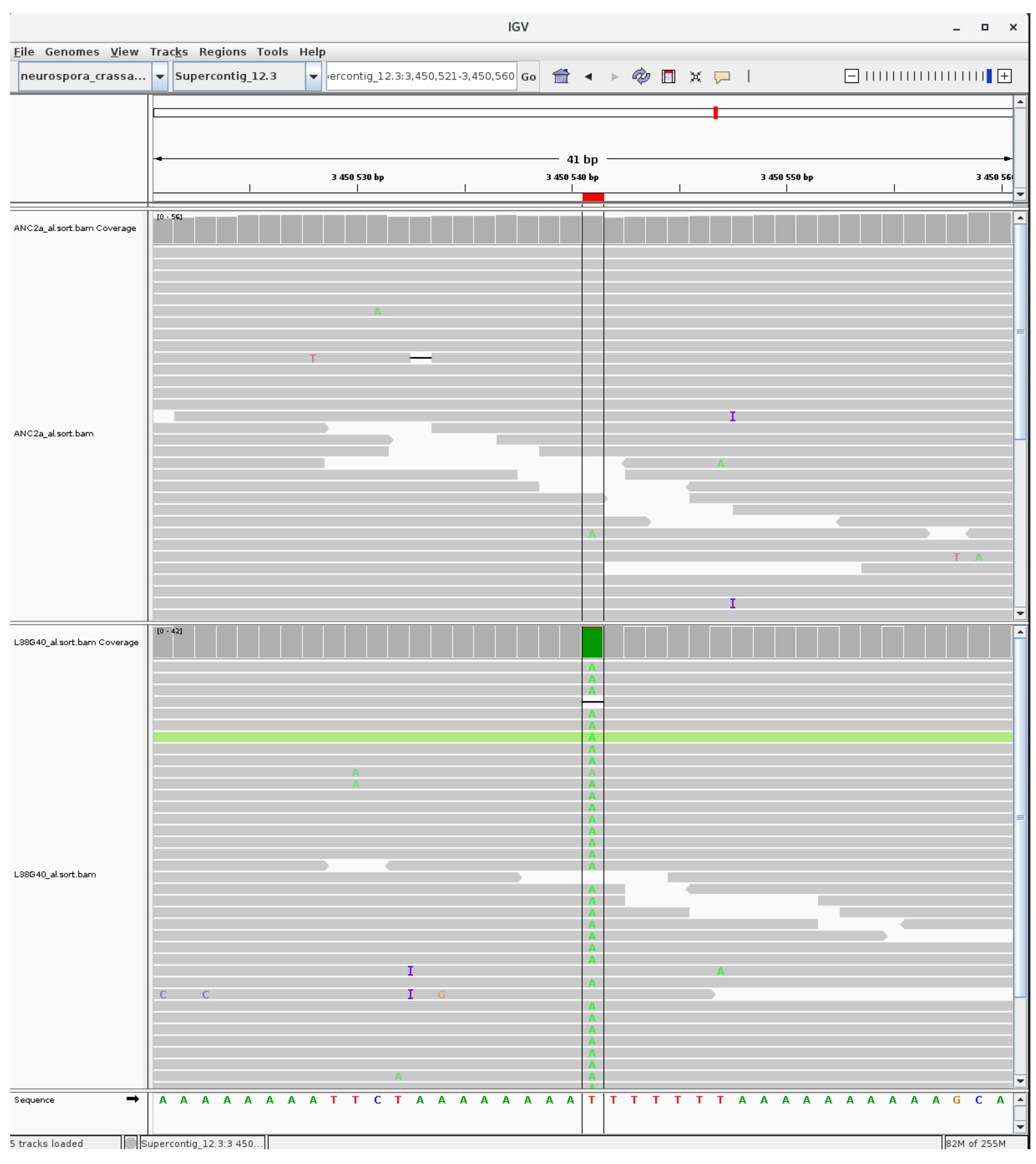

### mutation_euchromatic_24.jpg

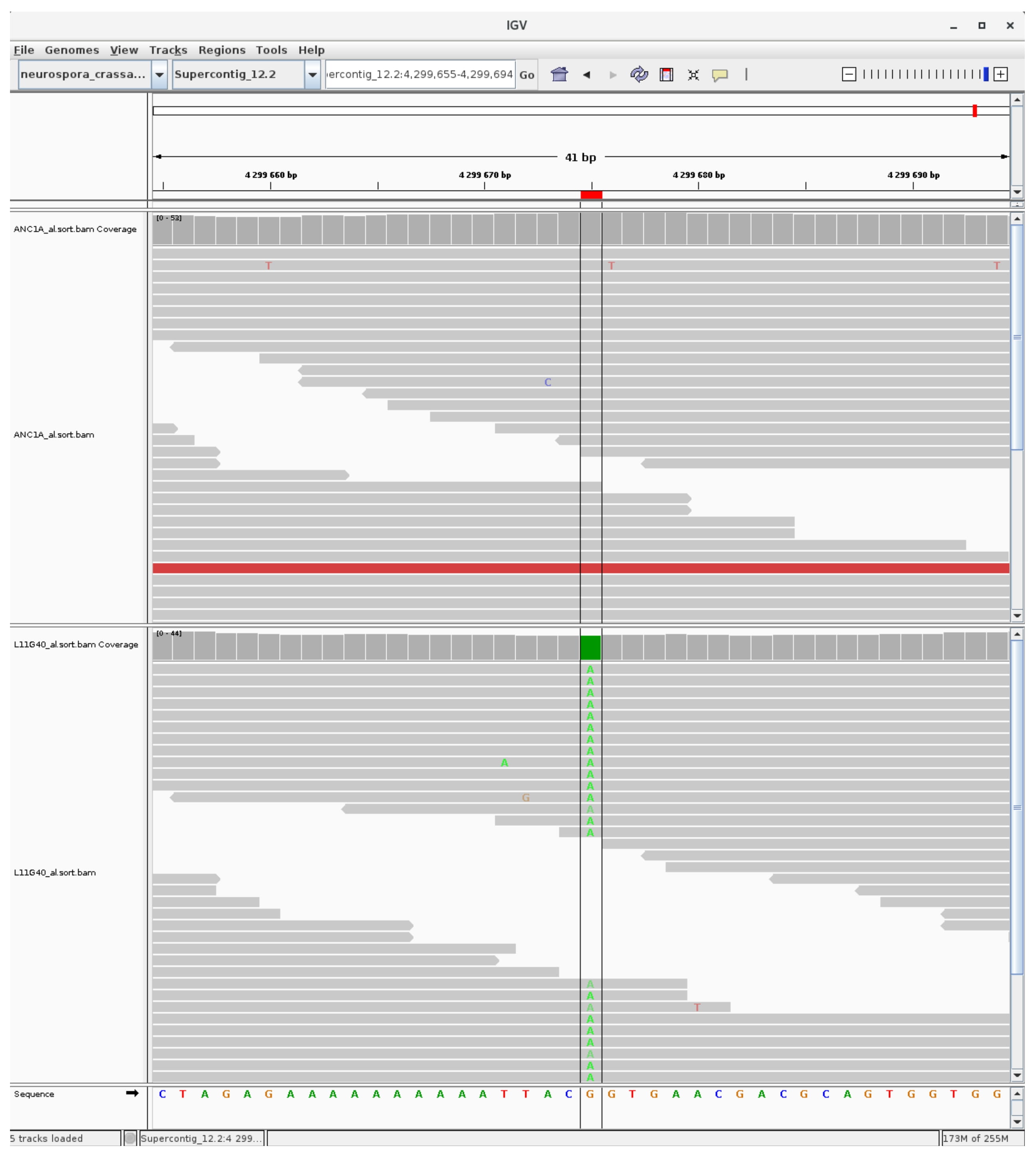

### mutation_euchromatic_25.jpg

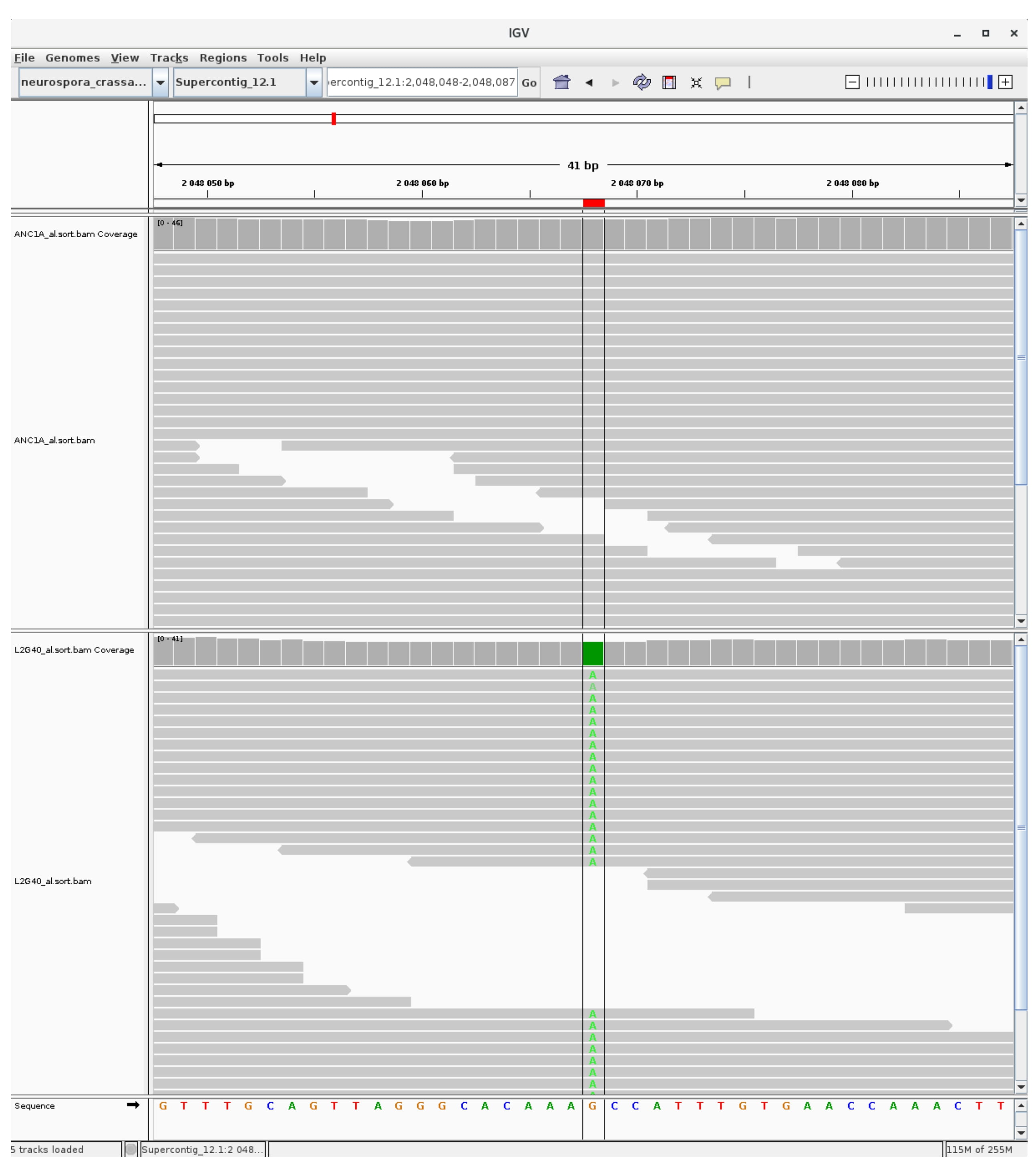

### mutation_euchromatic_26.jpg

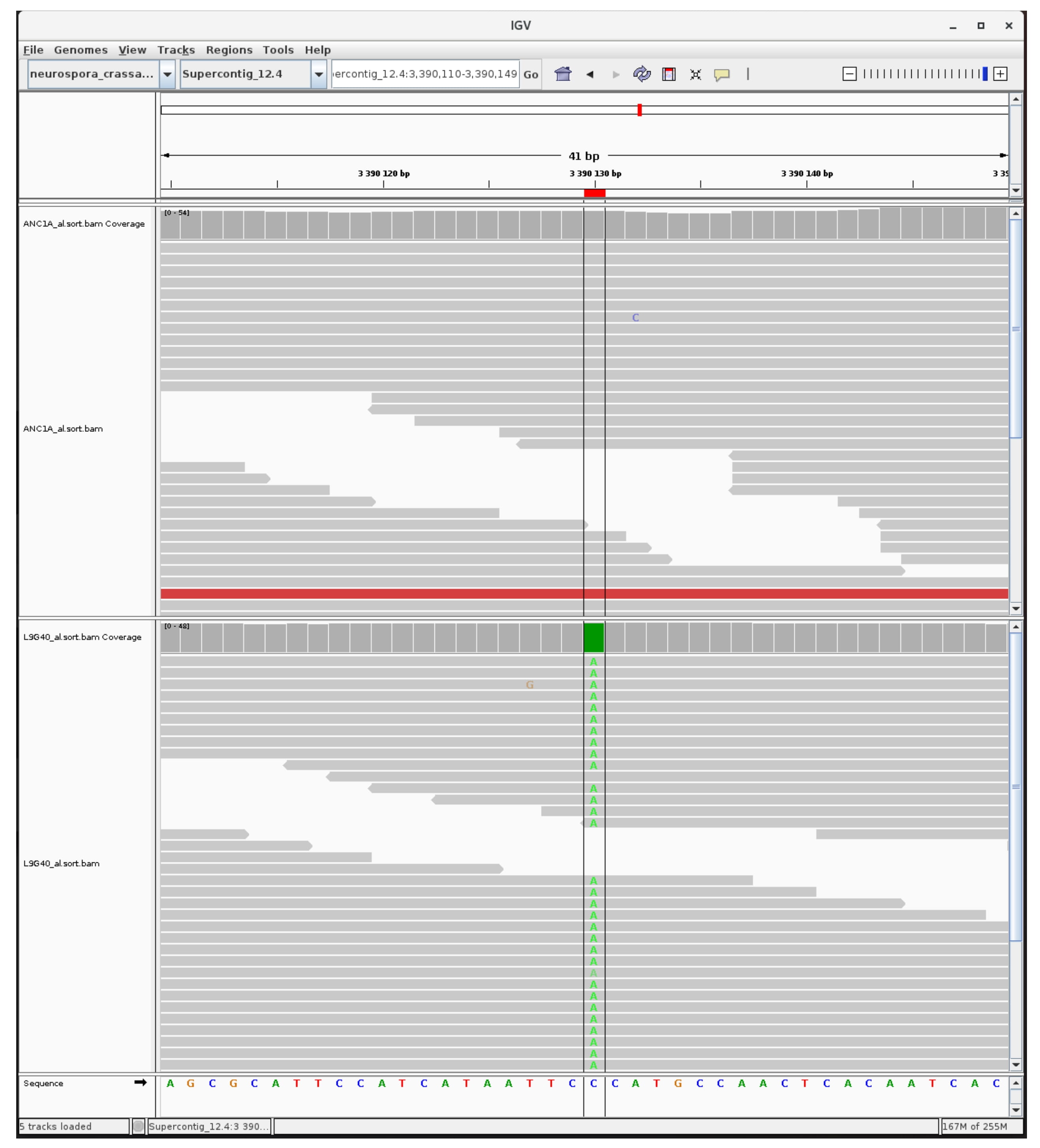

### mutation_euchromatic_27.jpg

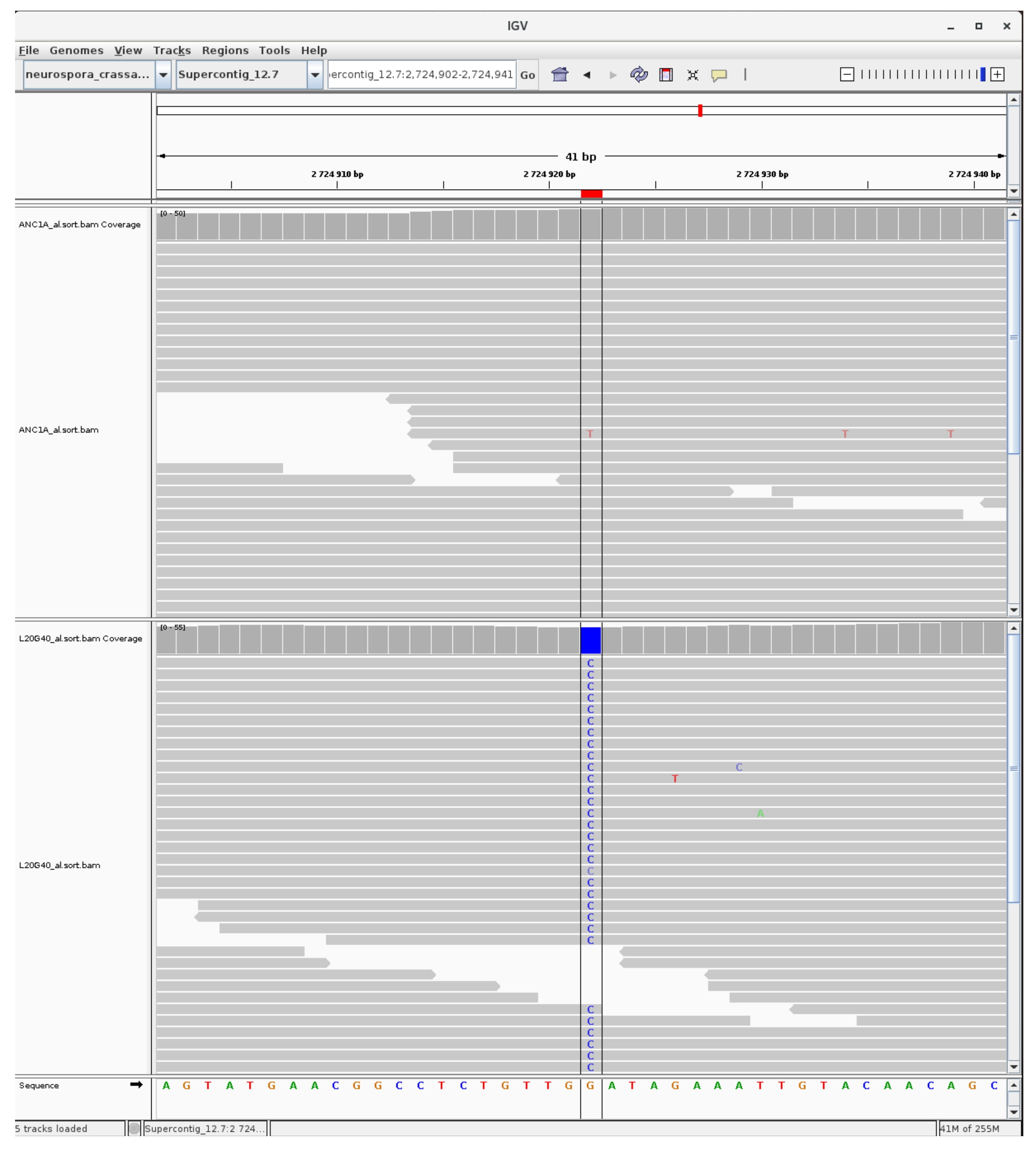

### mutation_euchromatic_28.jpg

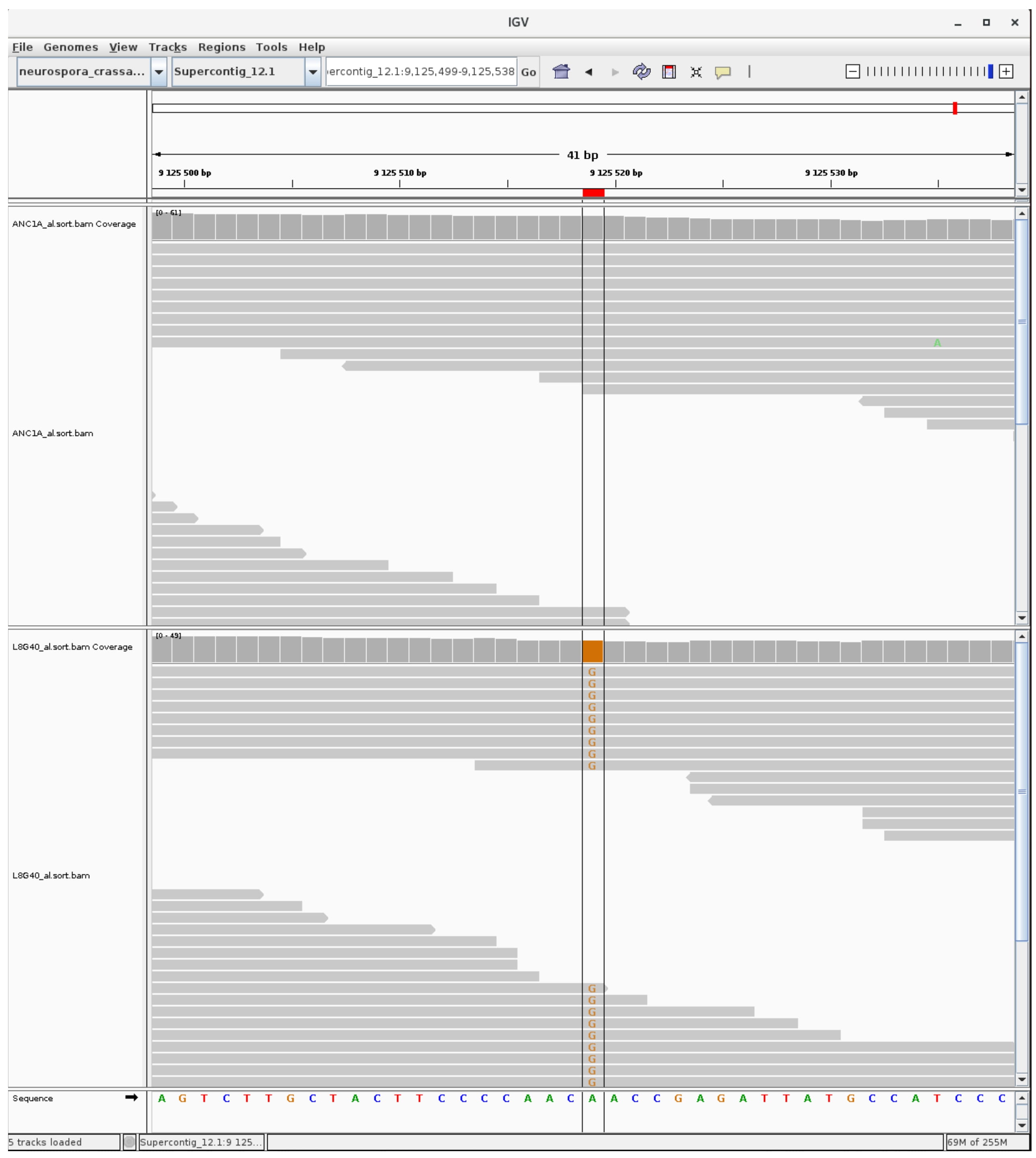

### mutation_euchromatic_29.jpg

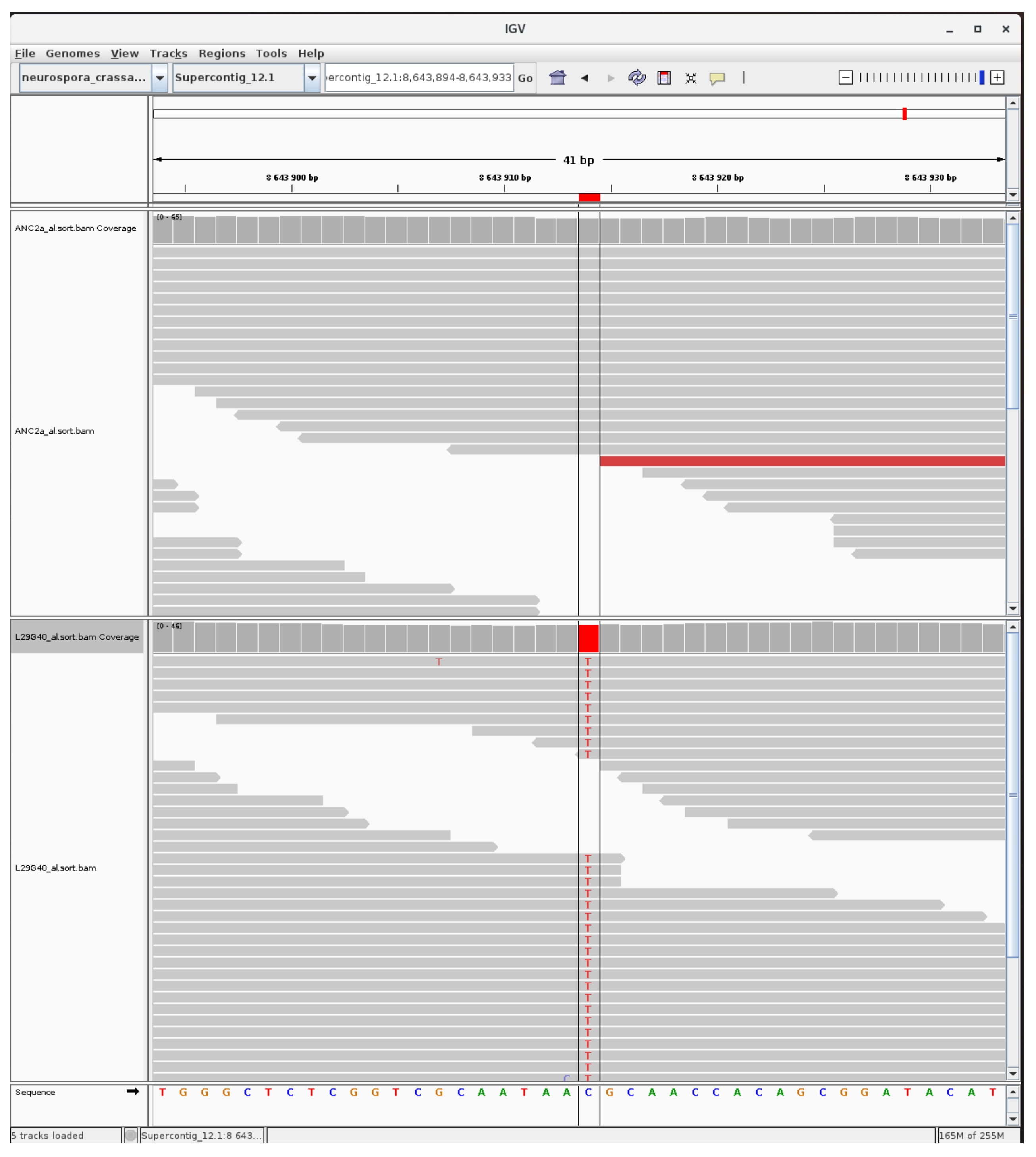

### mutation_euchromatic_30.jpg

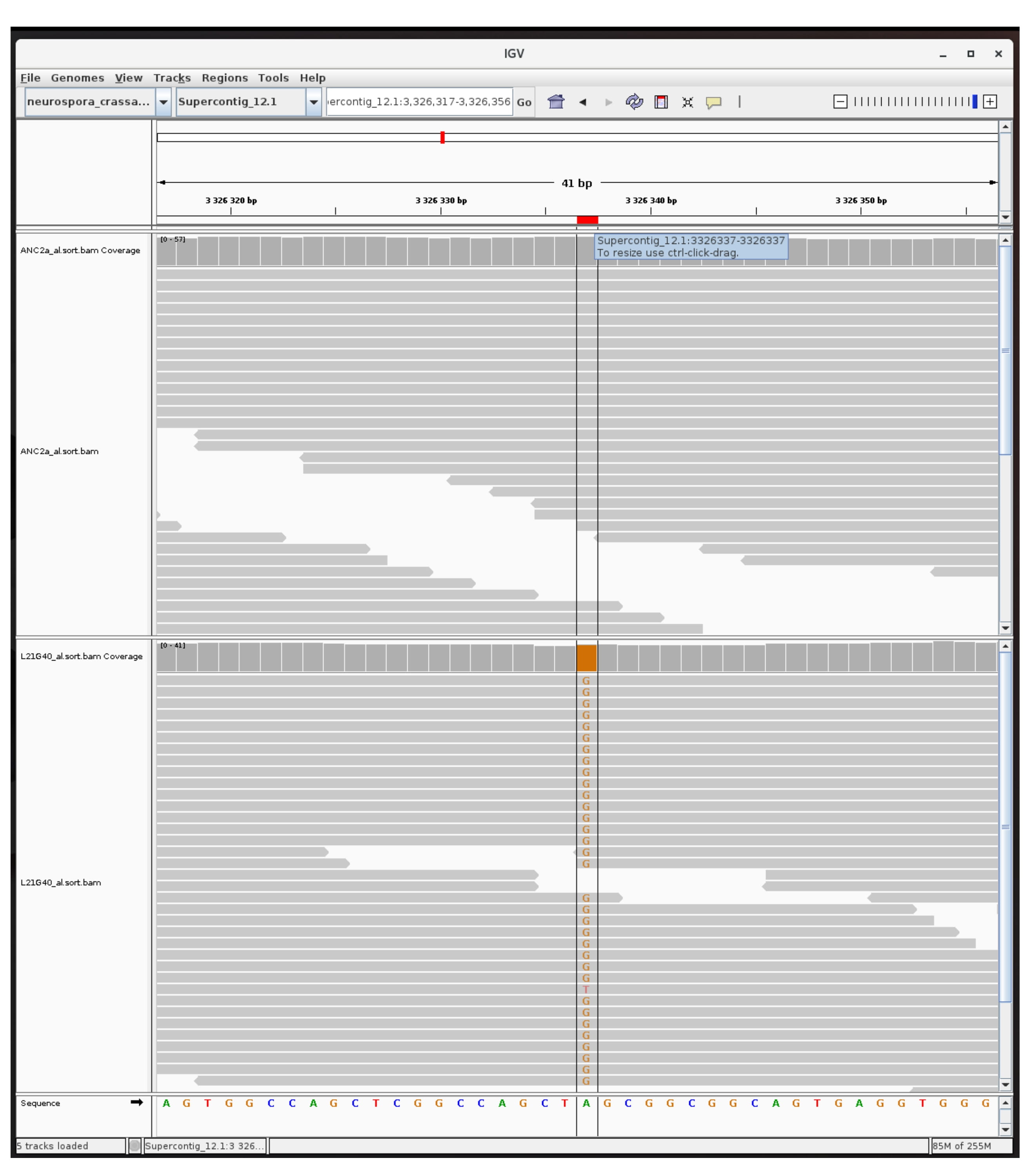

### mutation_H3K9_1.jpg

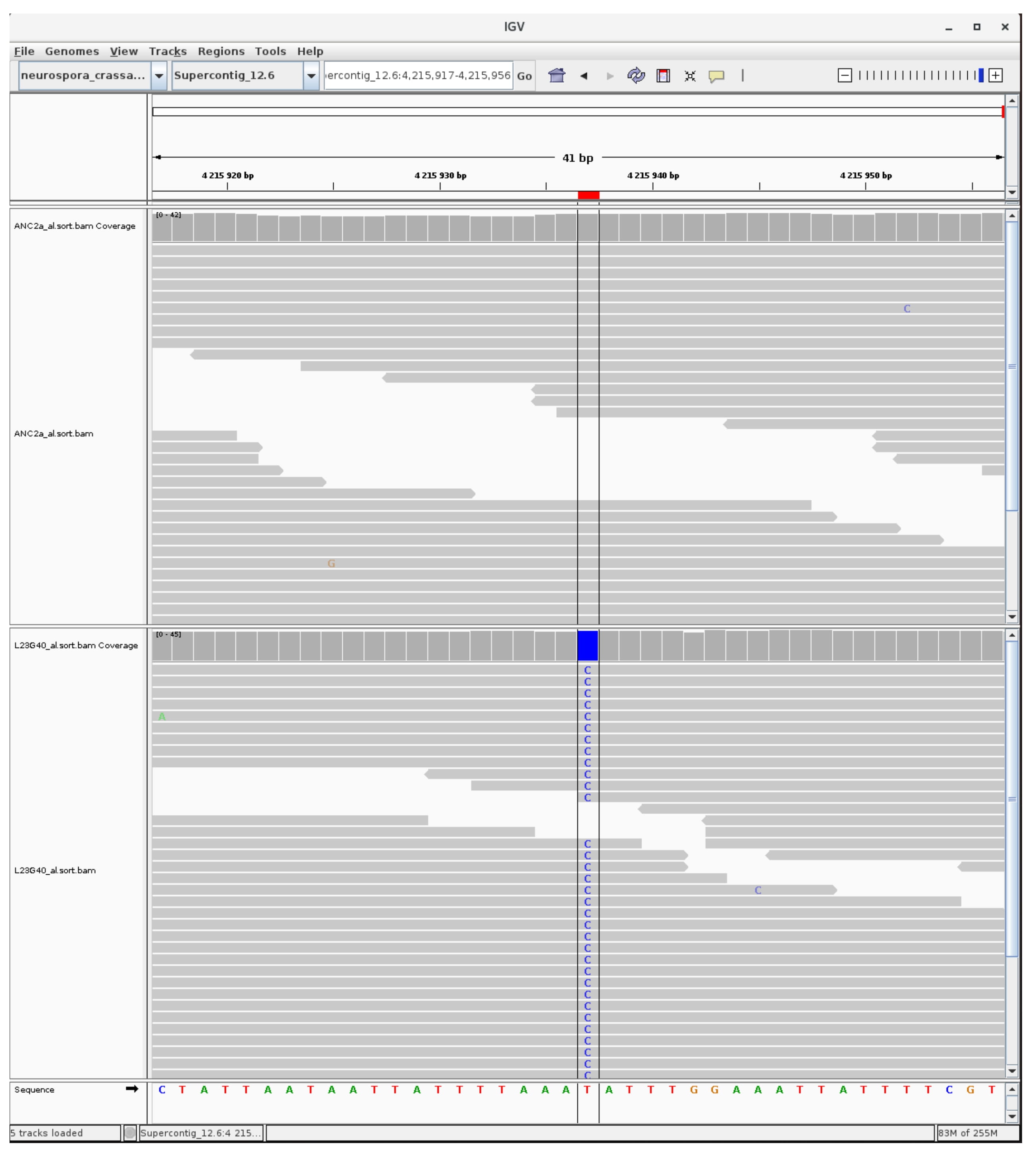

### mutation_H3K9_2.jpg

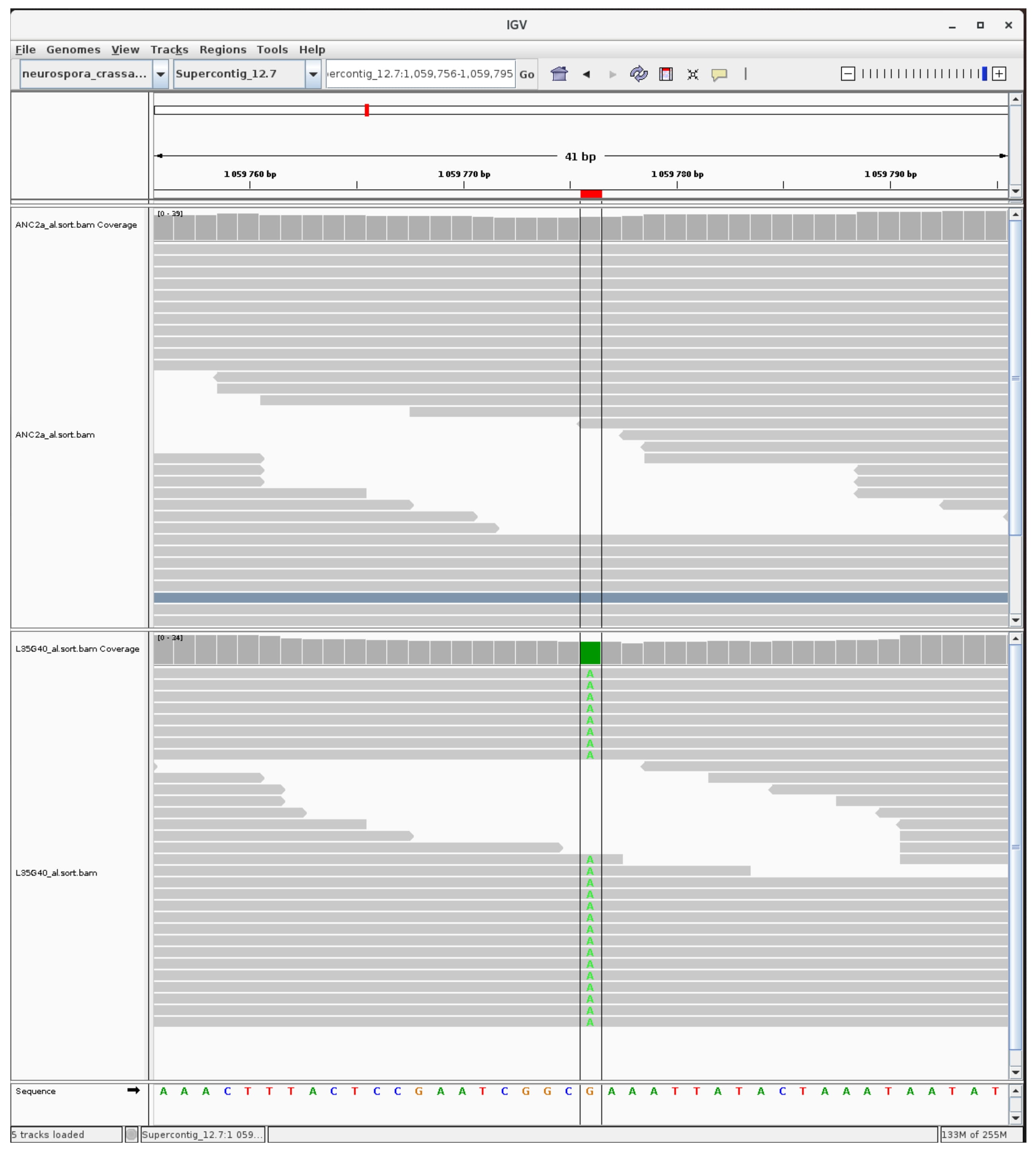

### mutation_H3K9_3.jpg

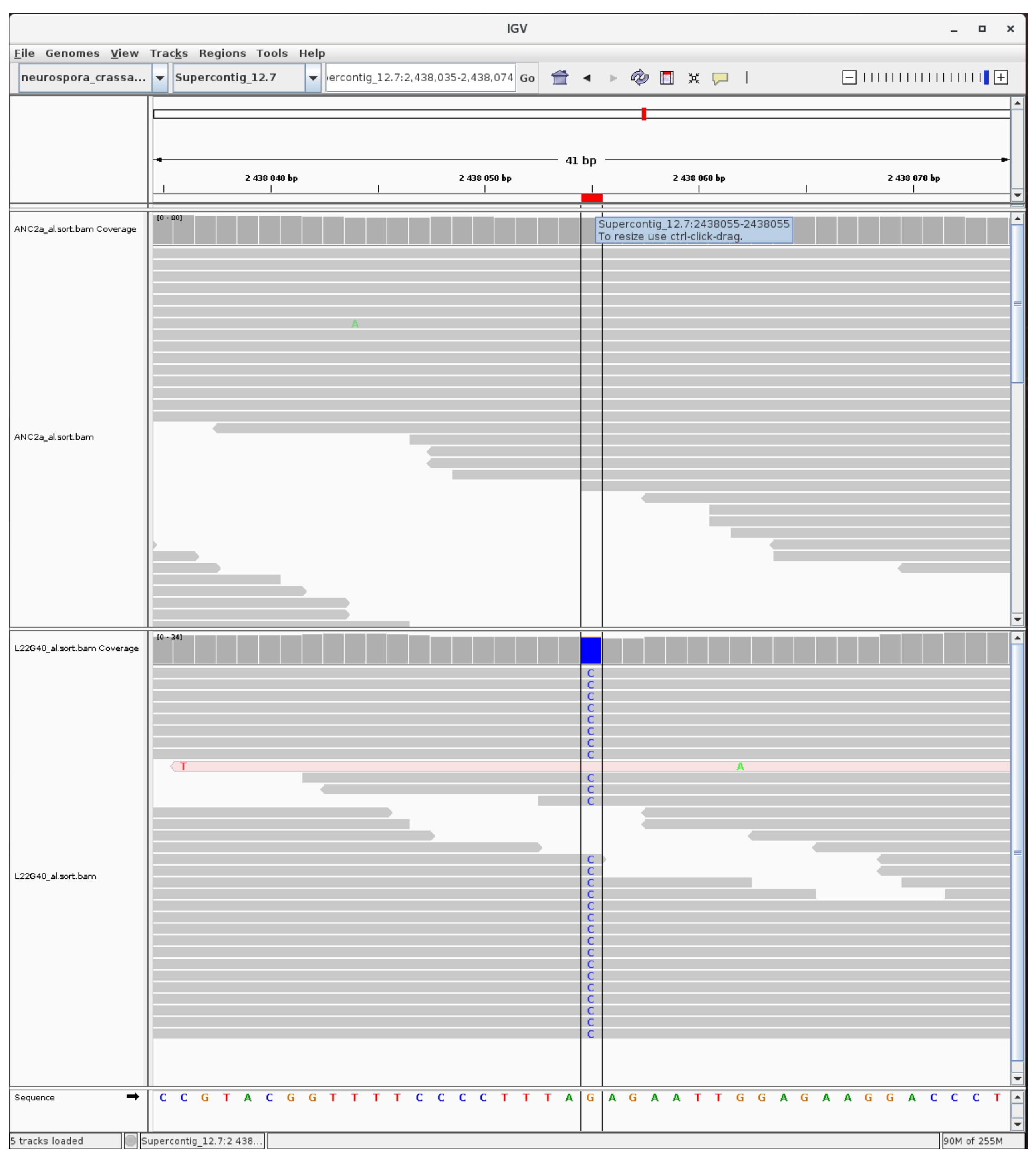

### mutation_H3K9_4.jpg

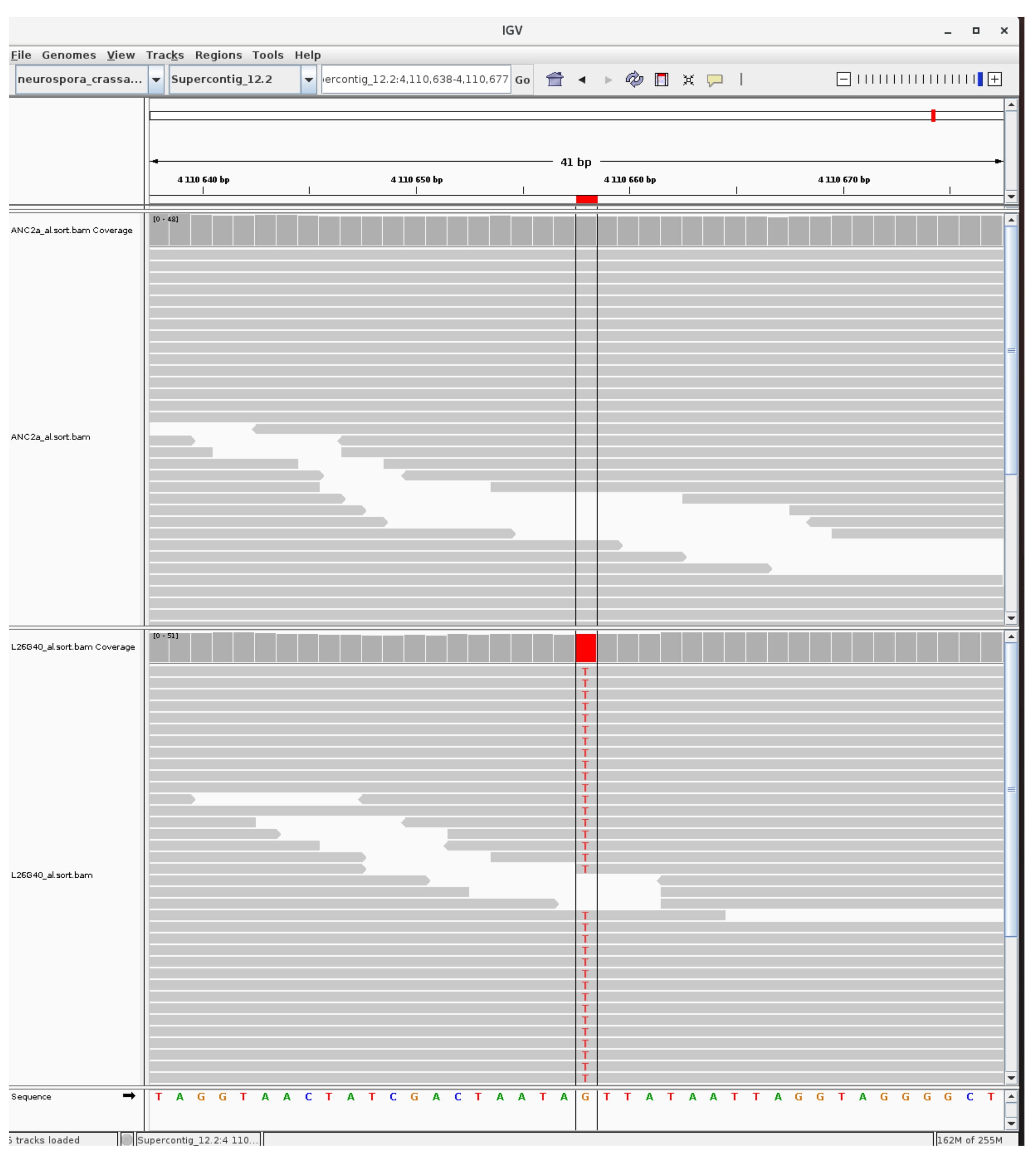

### mutation_H3K9_5.jpg

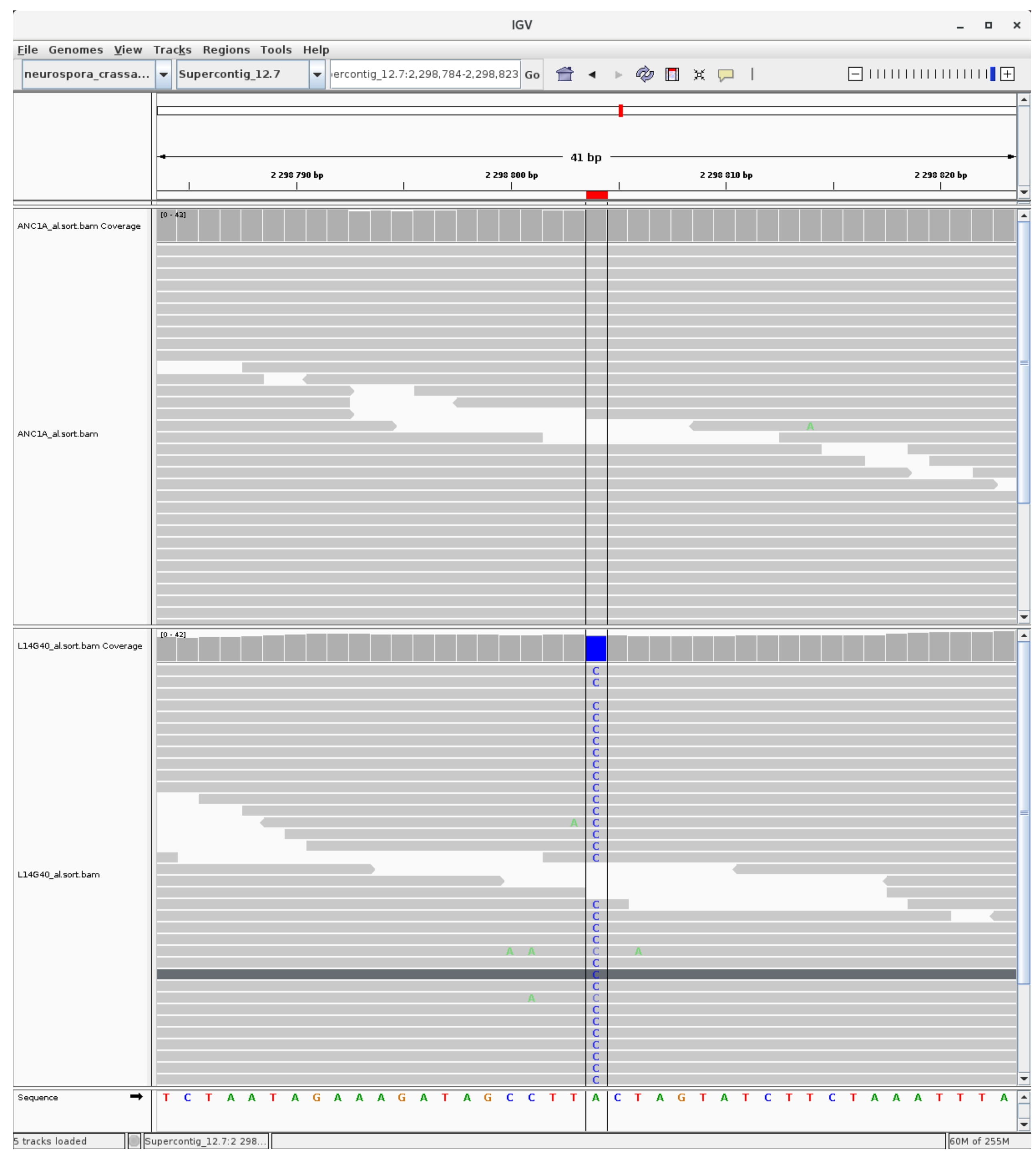

### mutation_H3K9_6.jpg

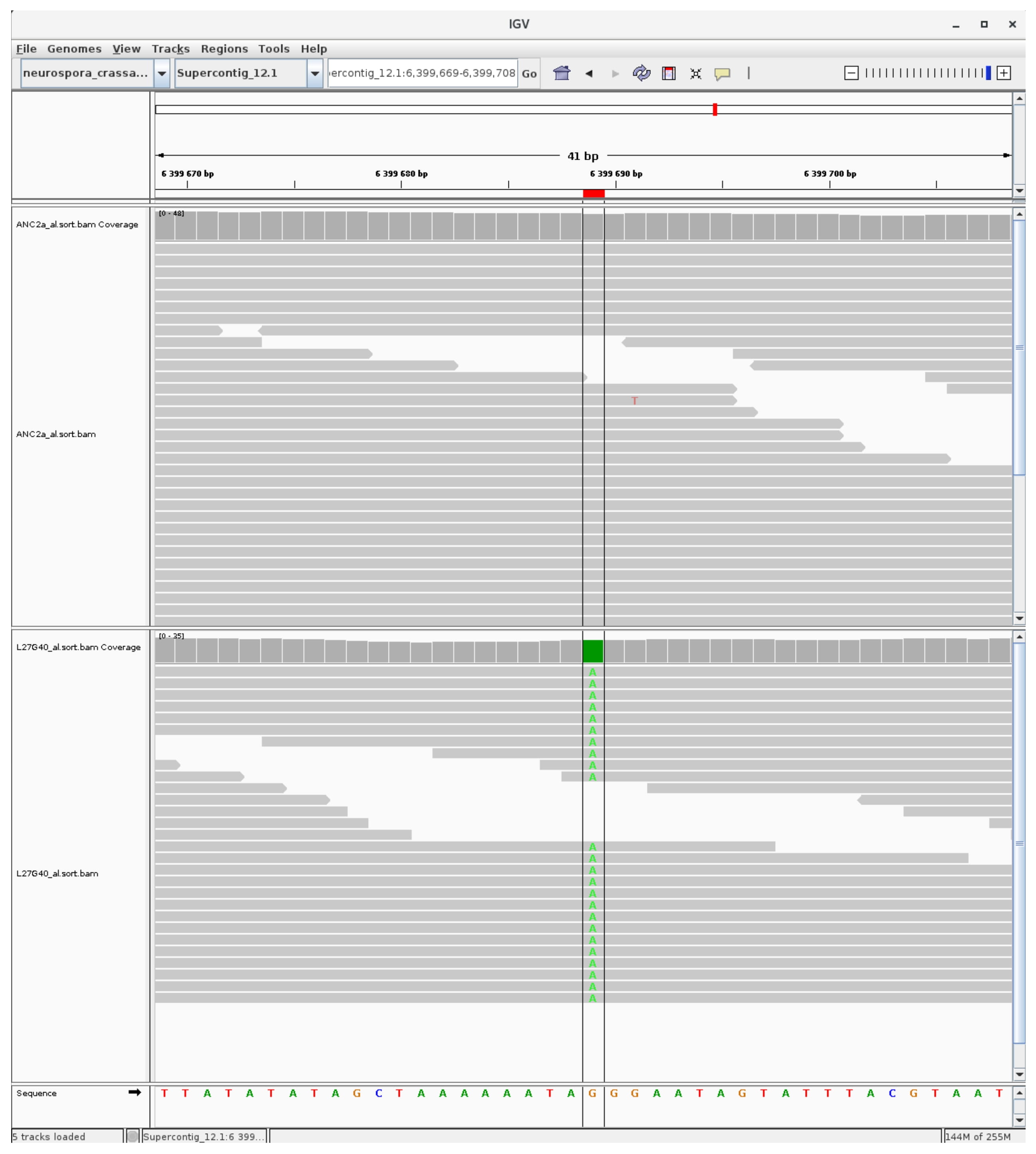

### mutation_H3K9_7.jpg

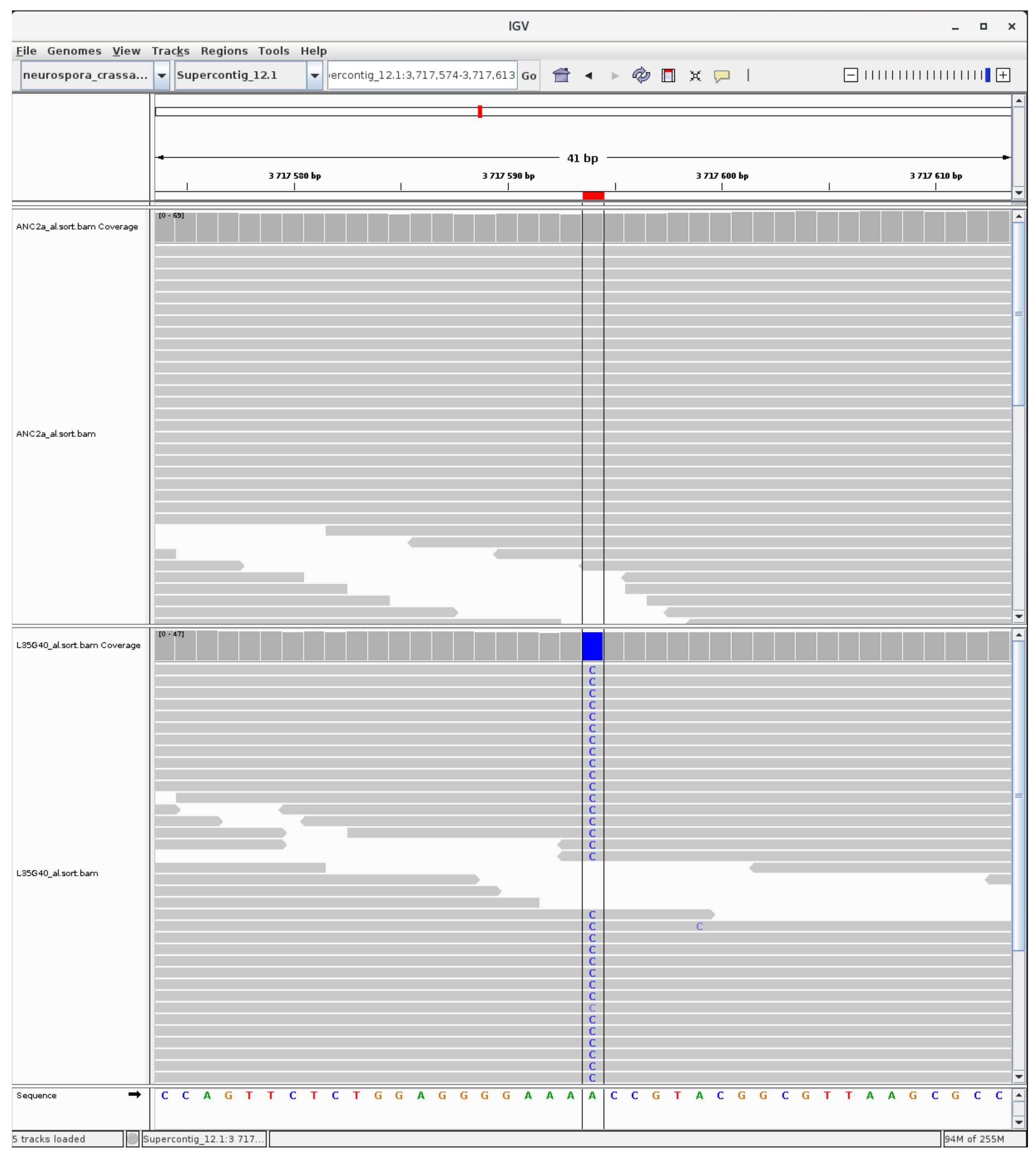

### mutation_H3K9_8.jpg

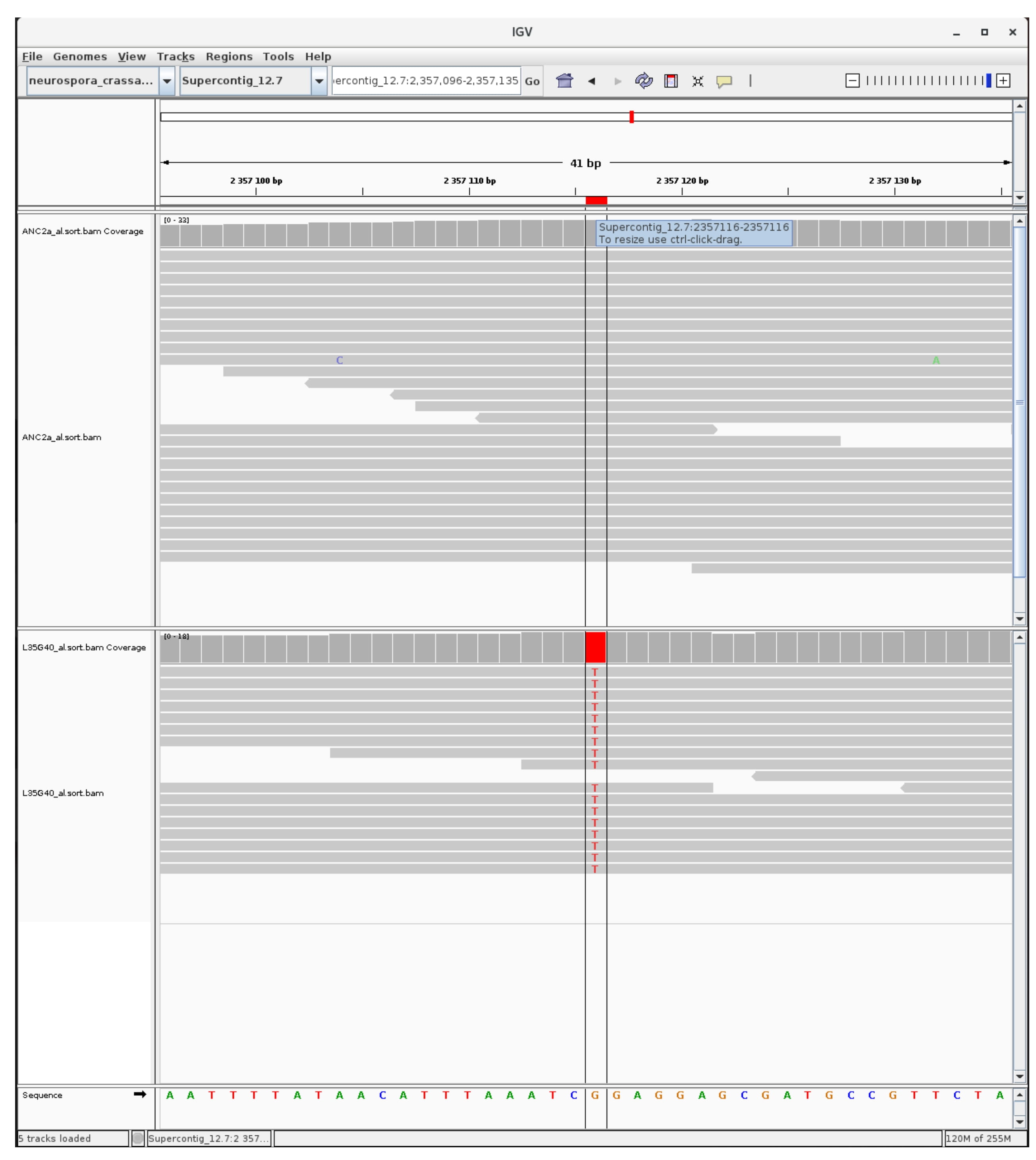

### mutation_H3K9_9.jpg

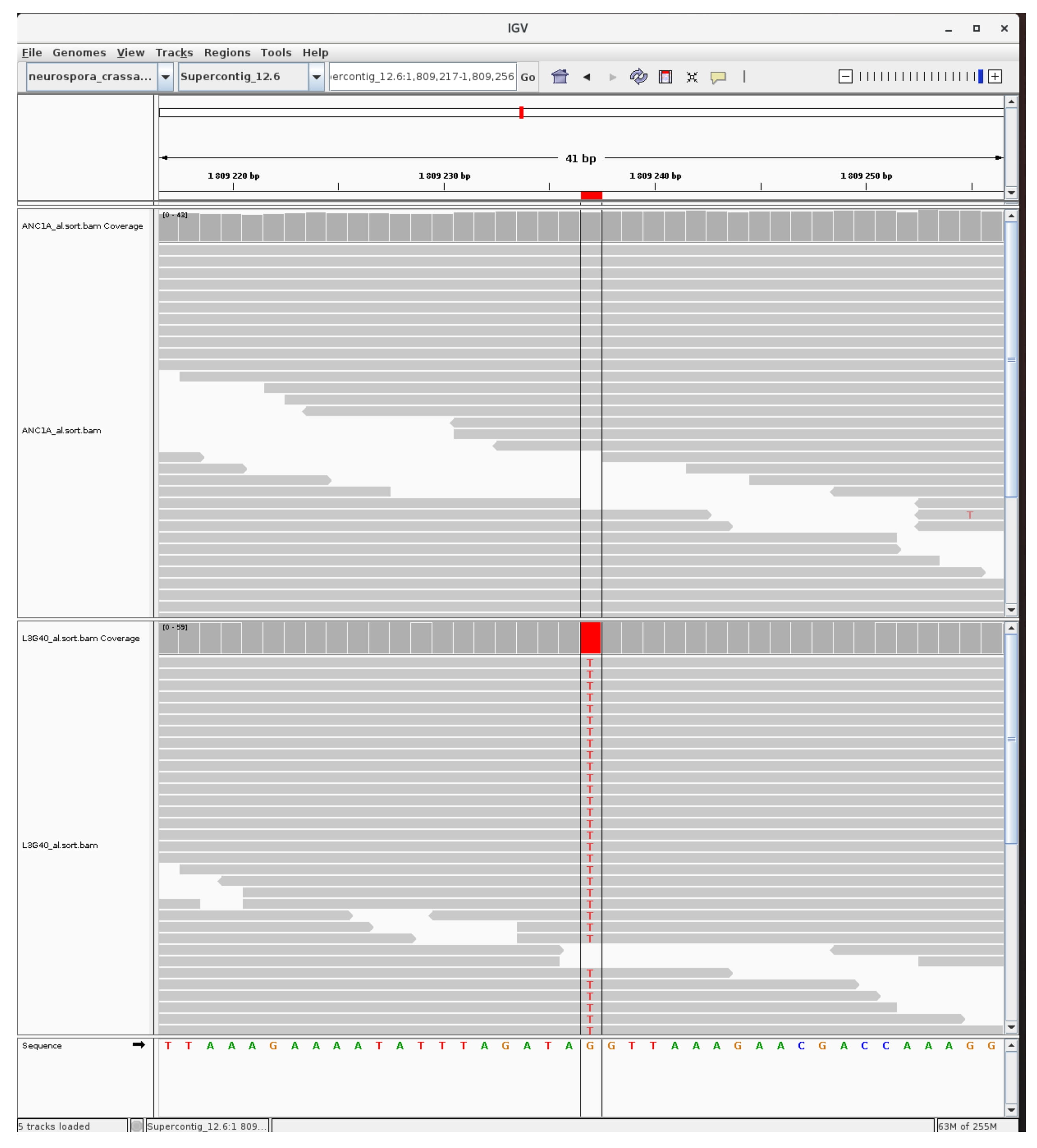

### mutation_H3K9_10.jpg

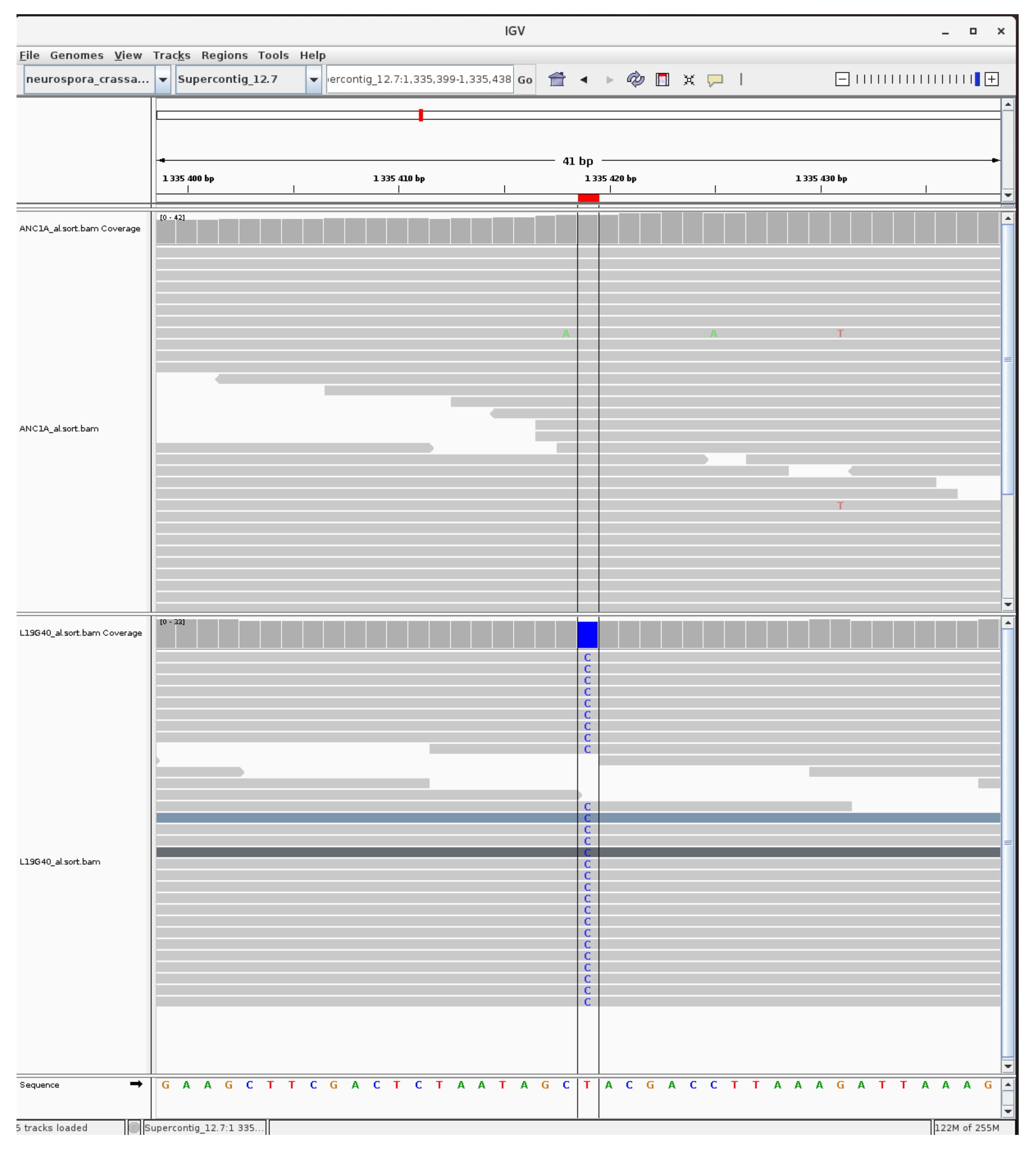

### mutation_H3K9_11.jpg

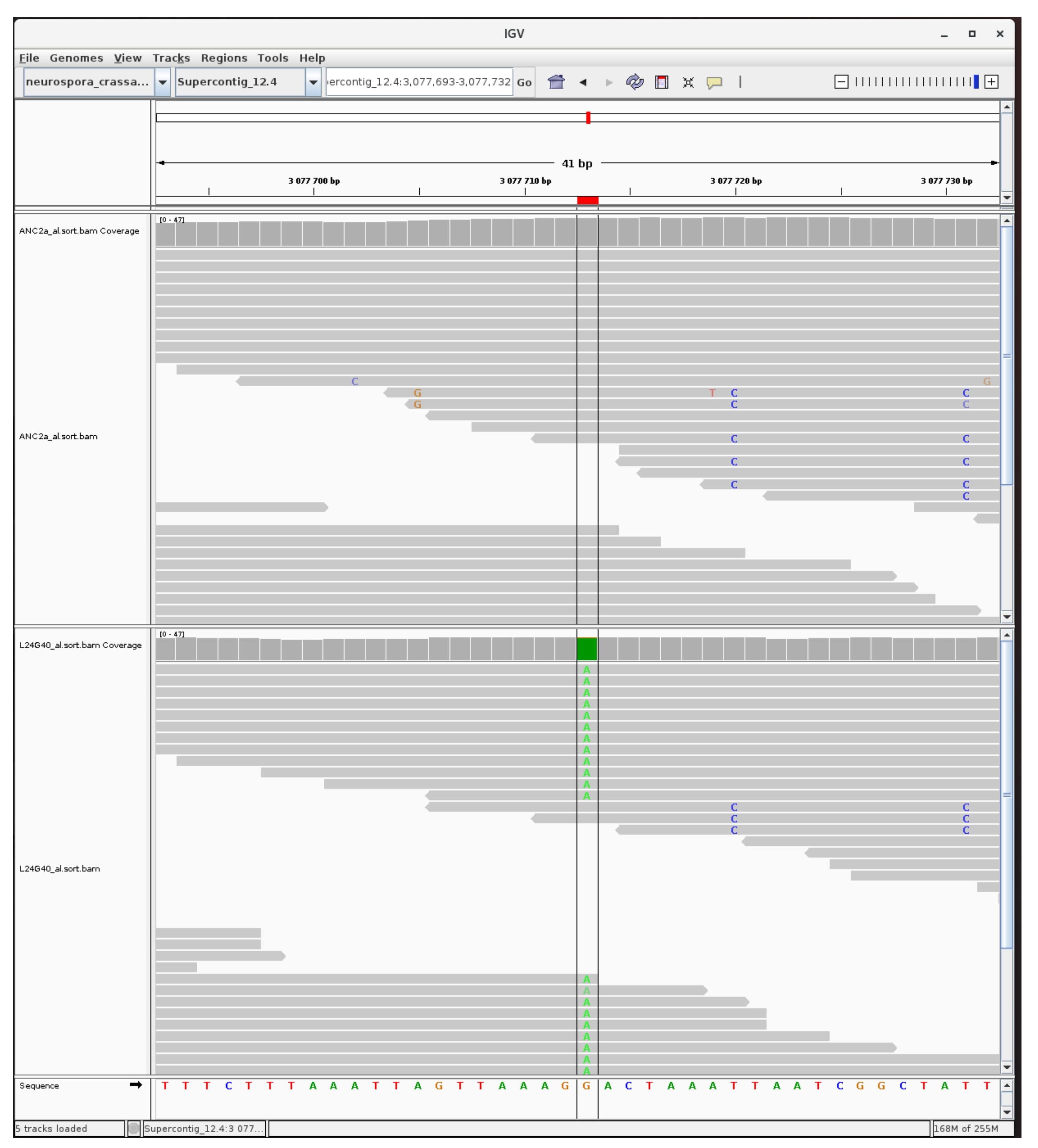

### mutation_H3K9_12.jpg

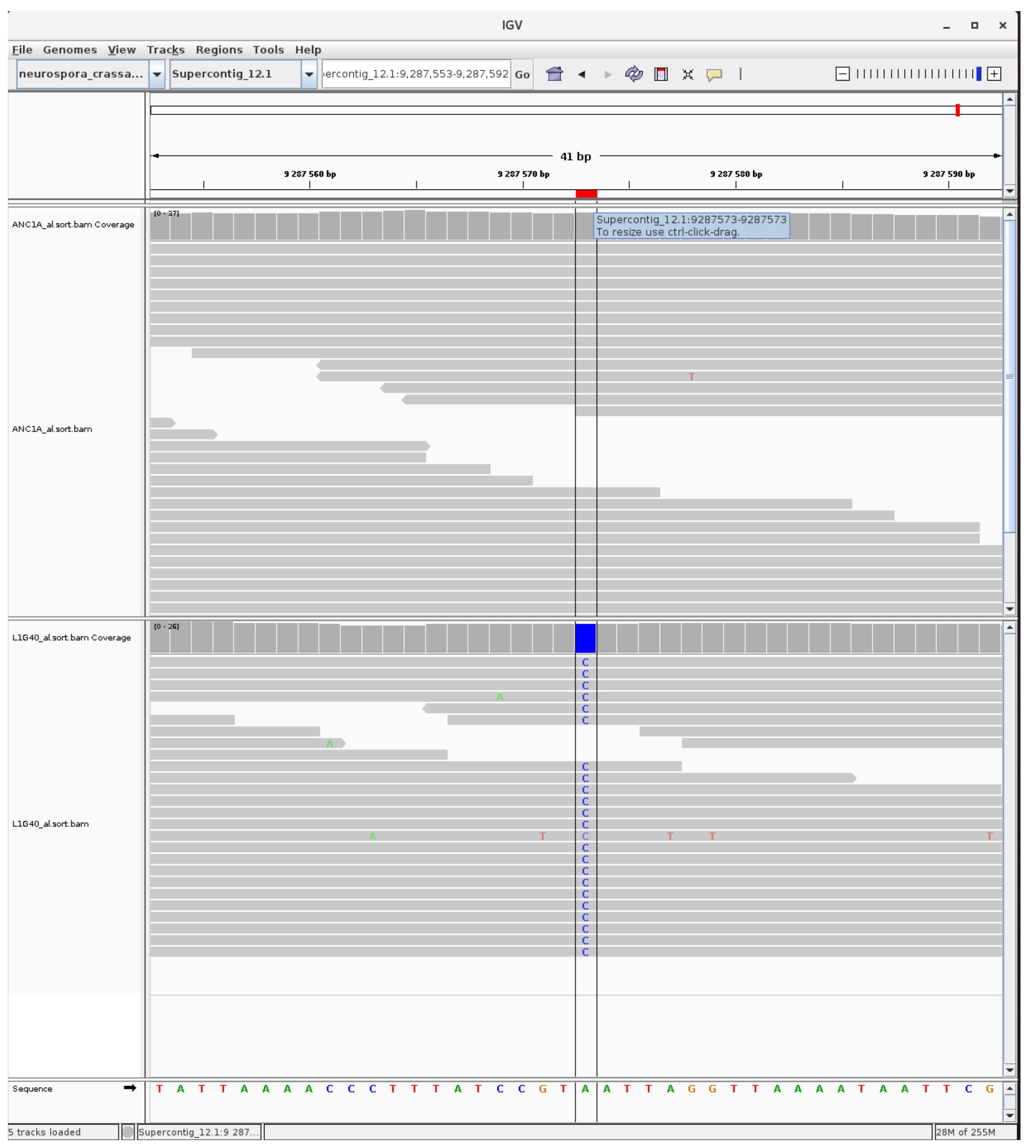

### mutation_H3K9_13.jpg

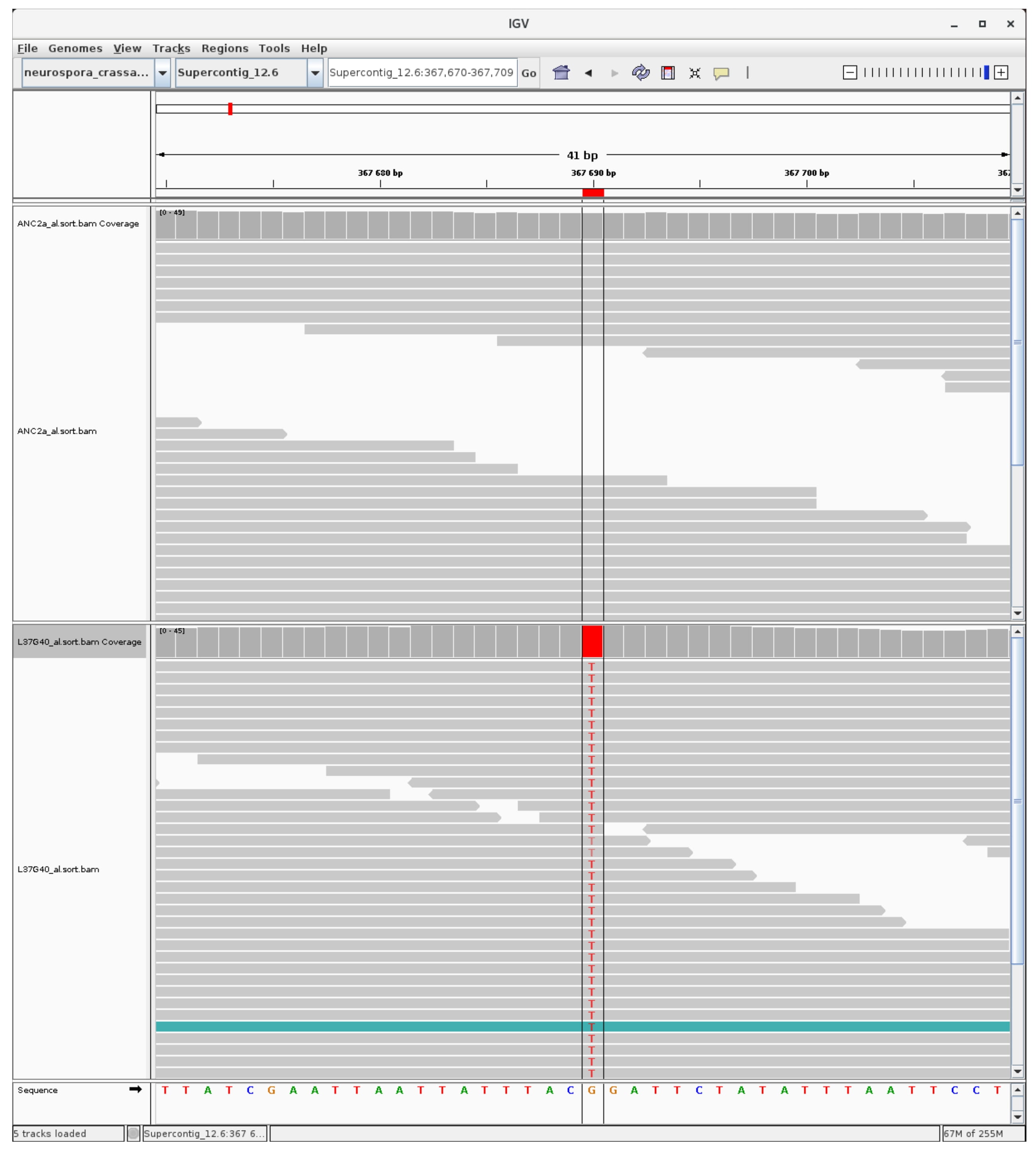

### mutation_H3K9_14.jpg

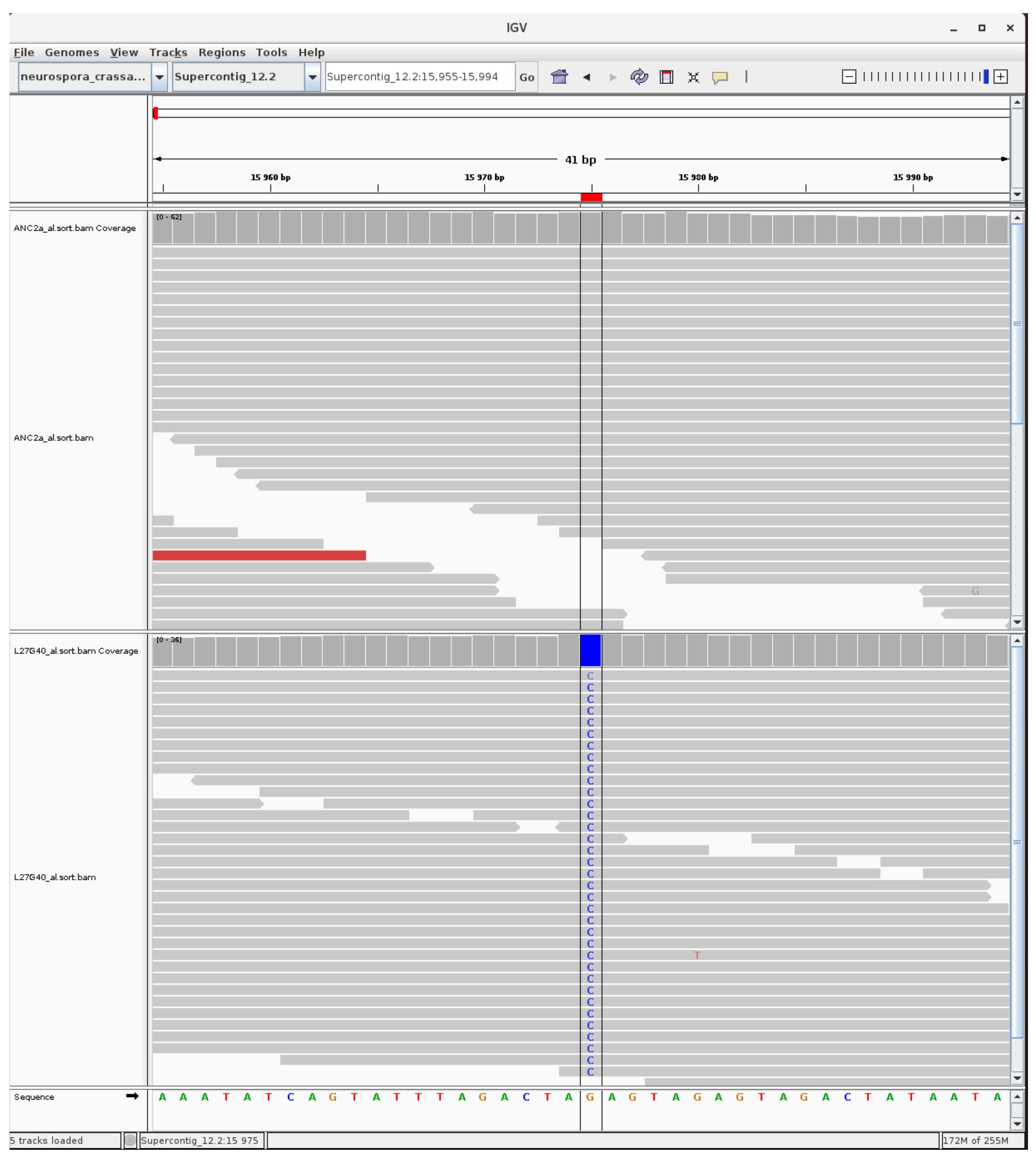

### mutation_H3K9_15.jpg

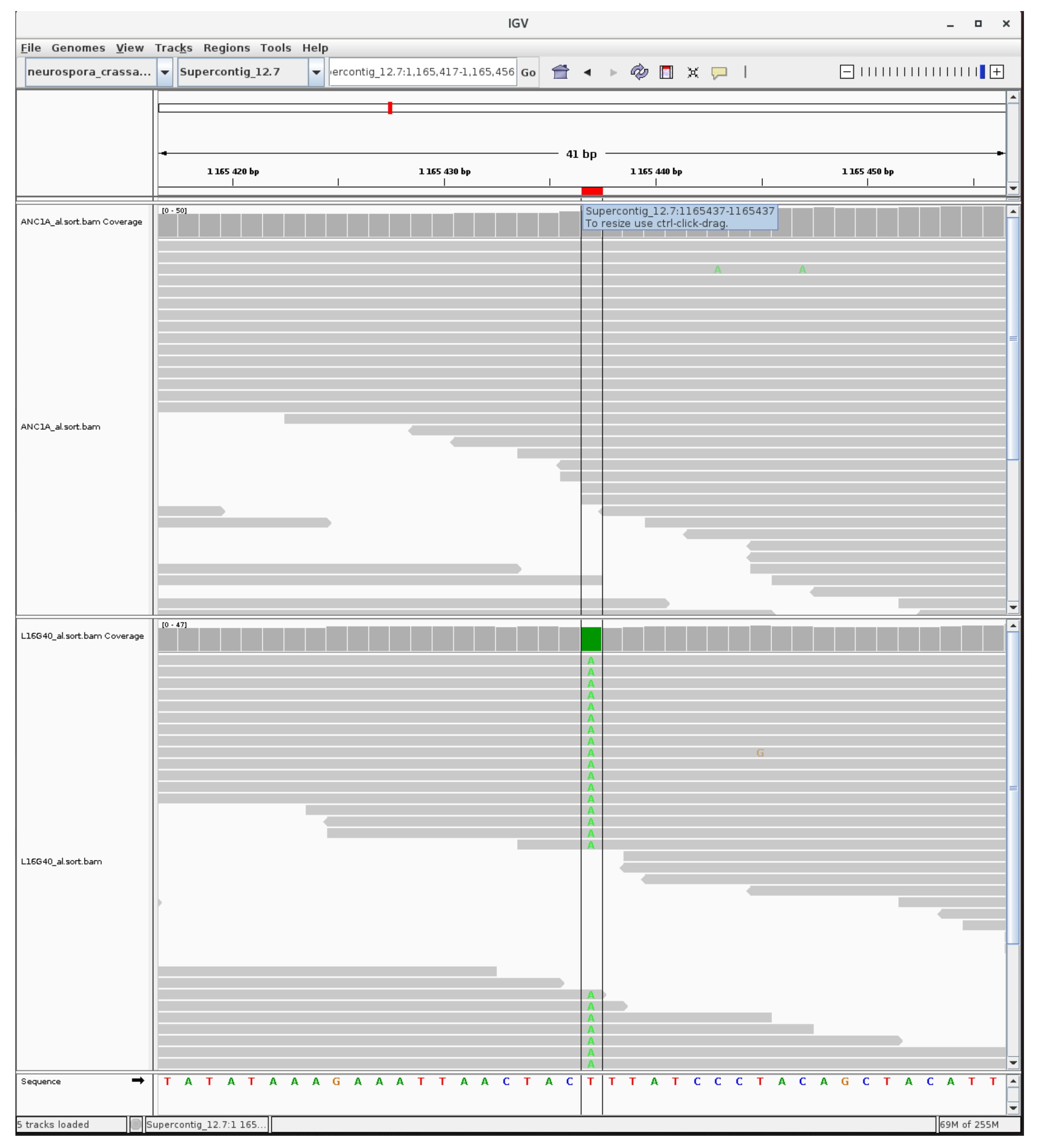

### mutation_H3K9_16.jpg

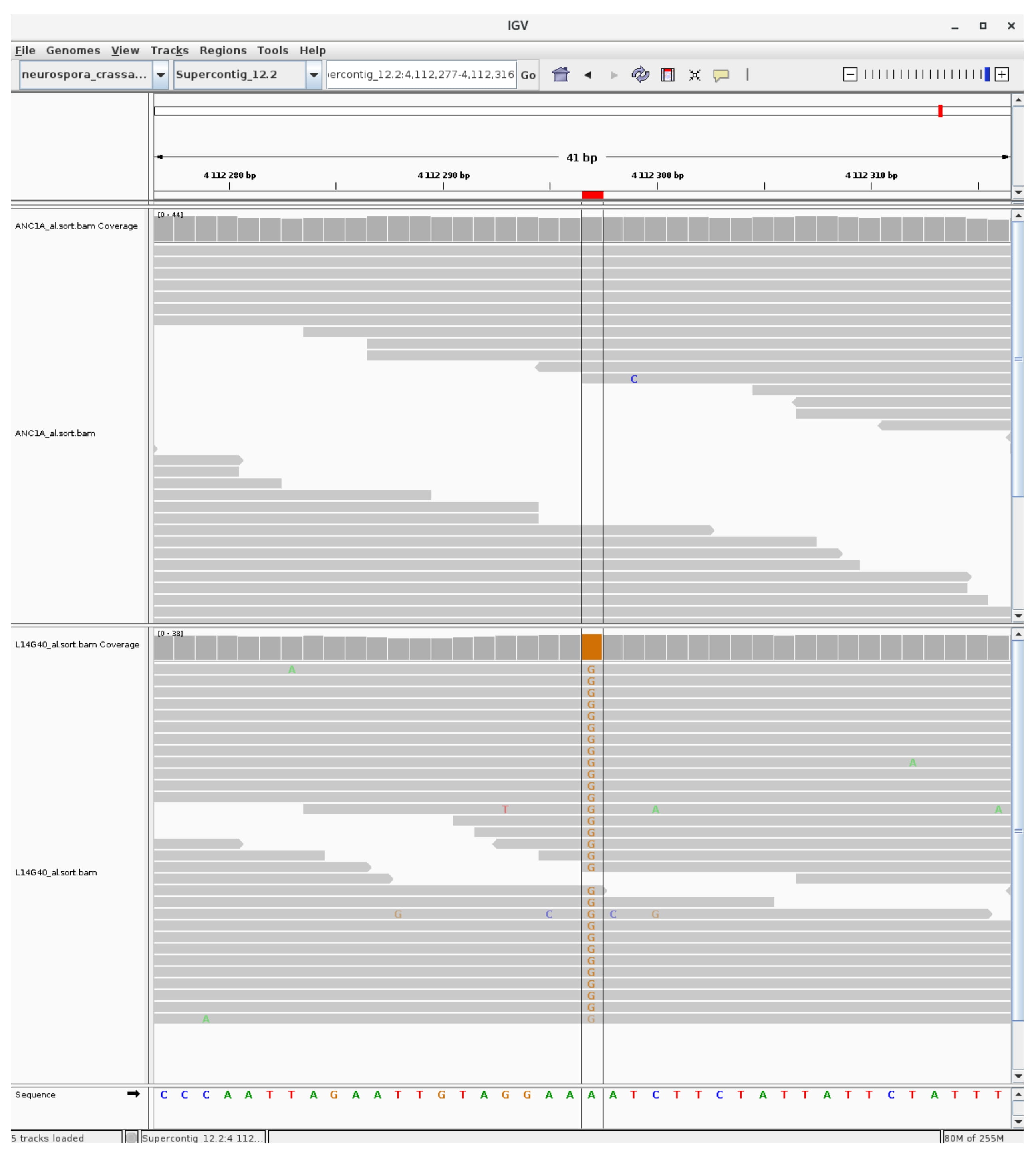

### mutation_H3K9_17.jpg

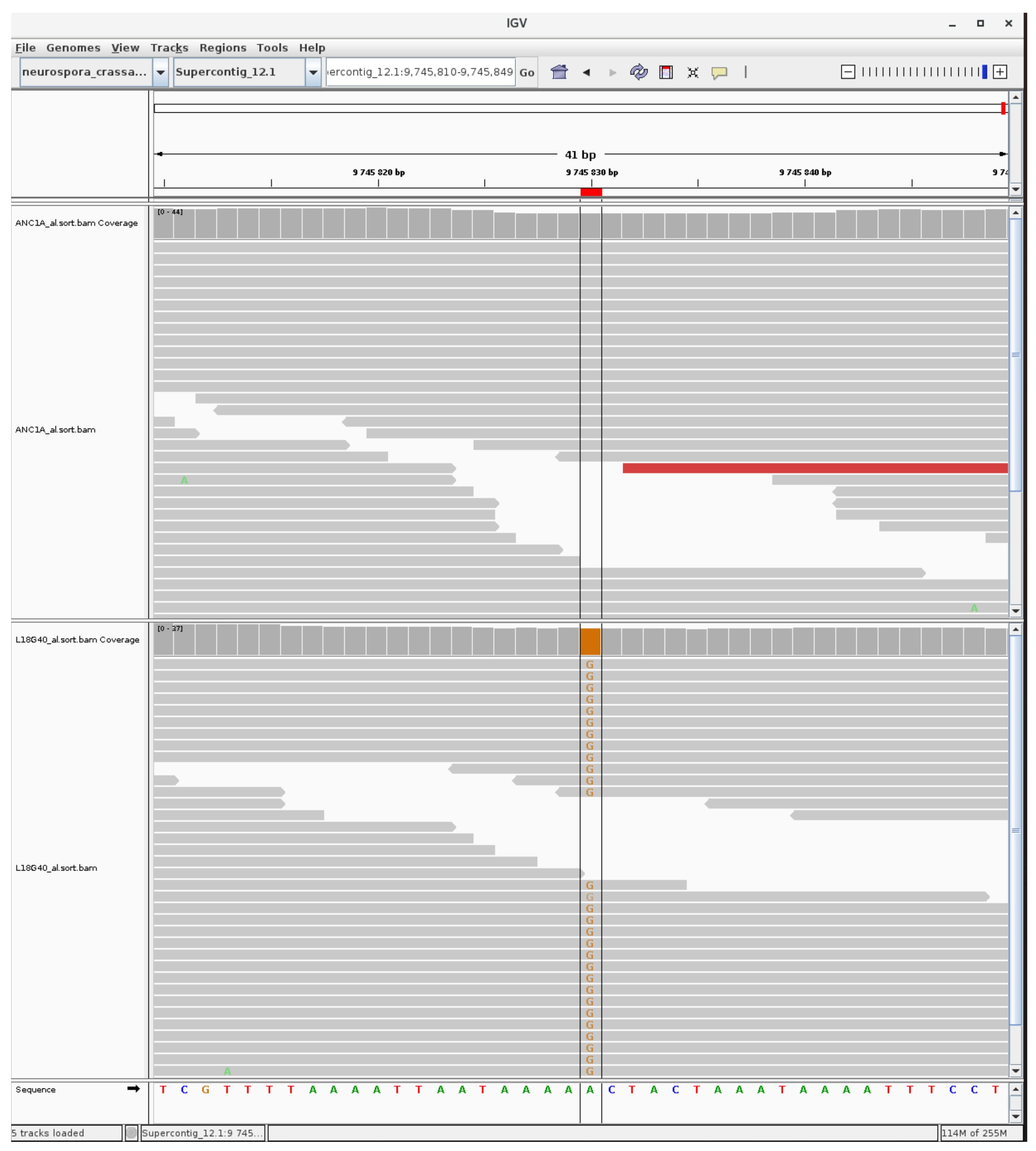

### mutation_H3K9_18.jpg

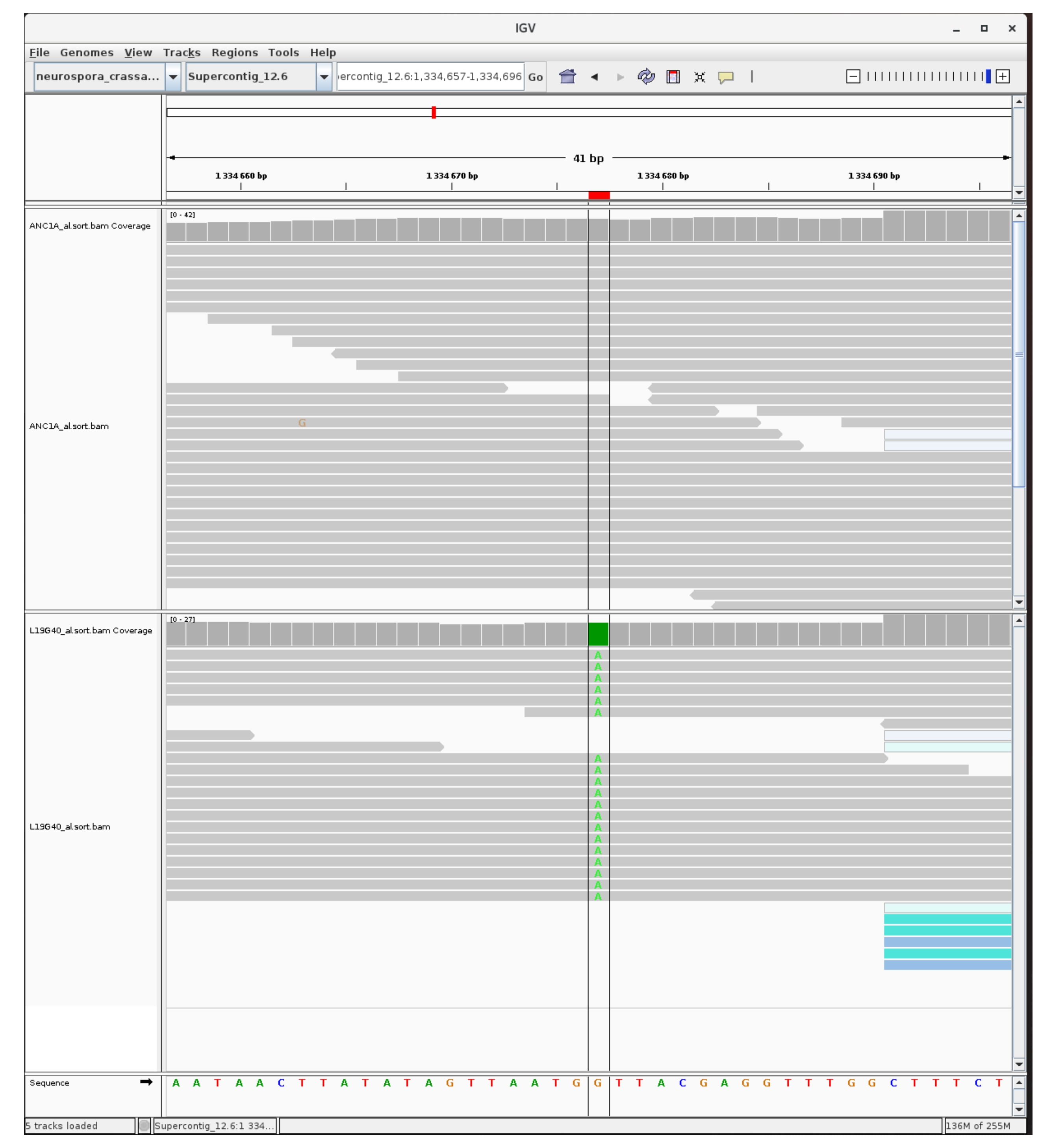

### mutation_H3K9_19.jpg

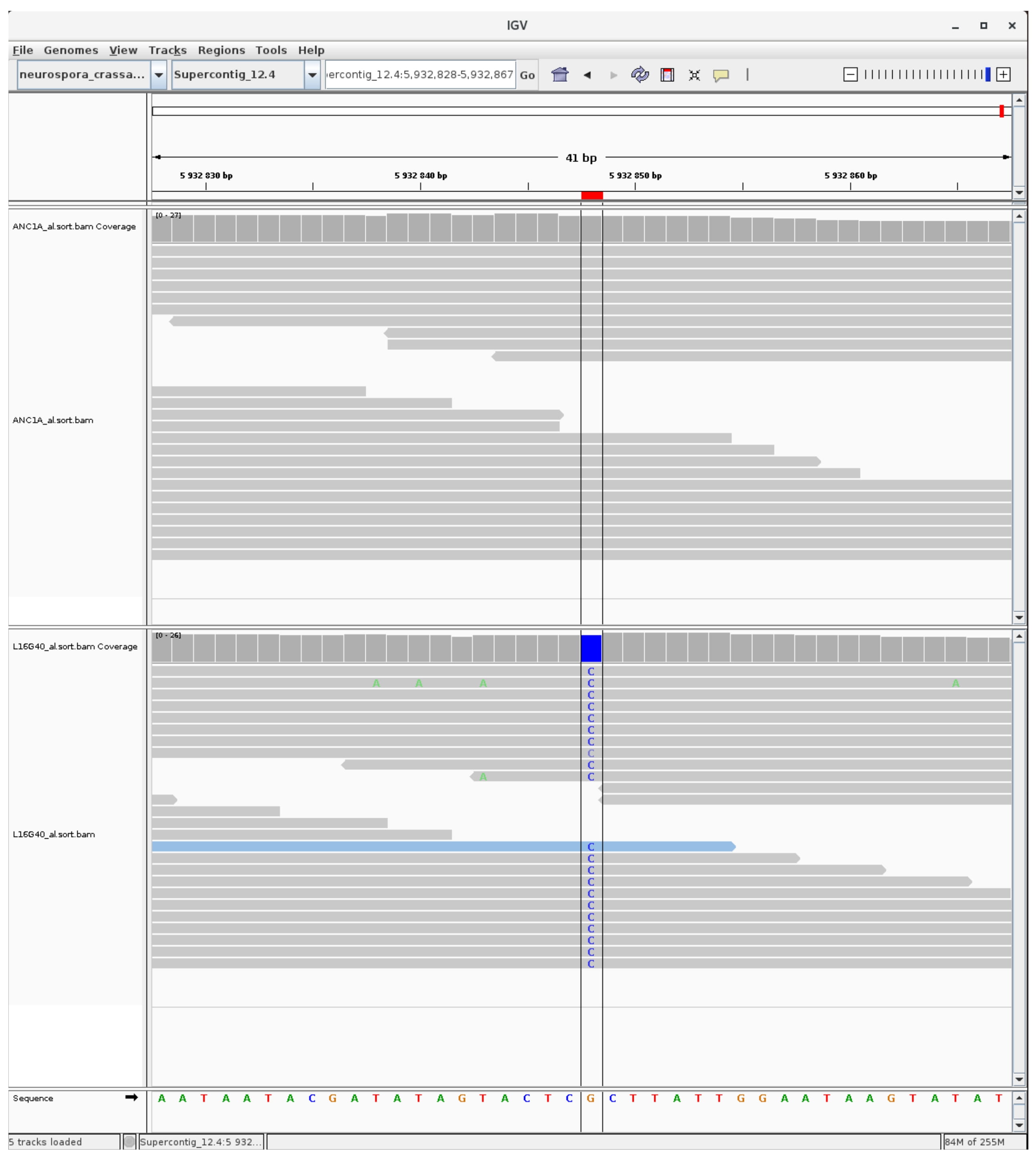

### mutation_H3K9_20.jpg

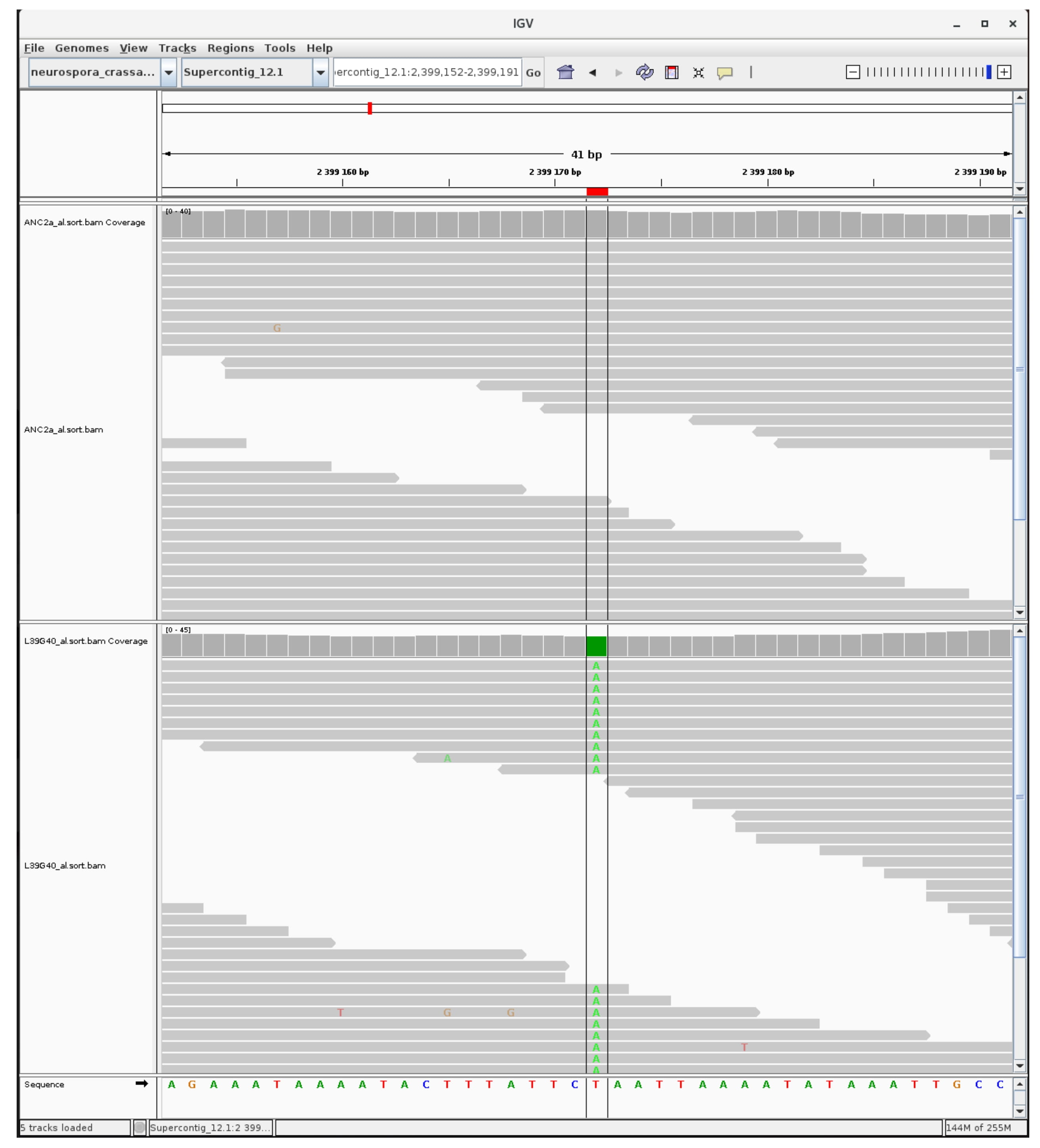

### mutation_H3K9_21.jpg

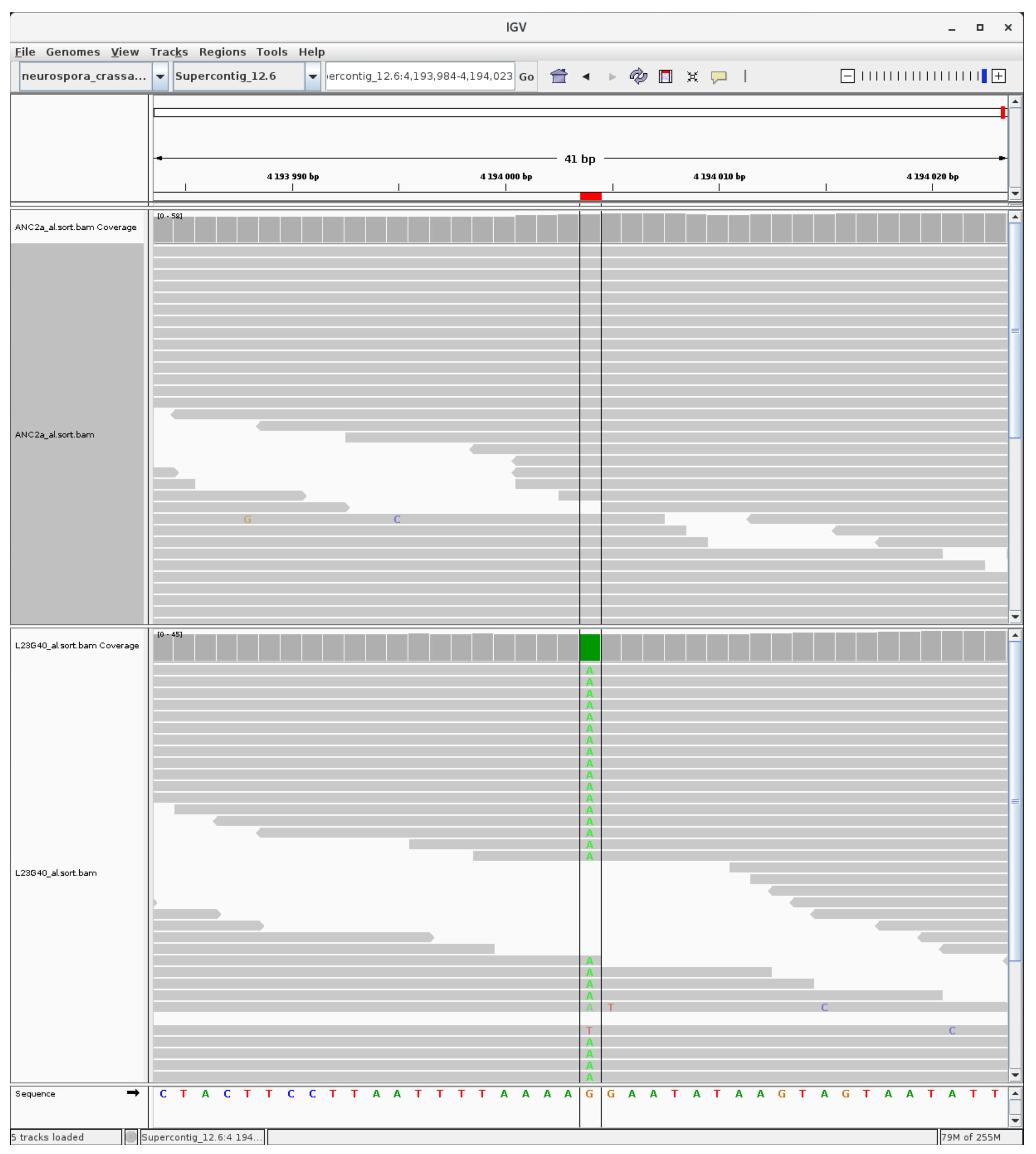

### mutation_H3K9_22.jpg

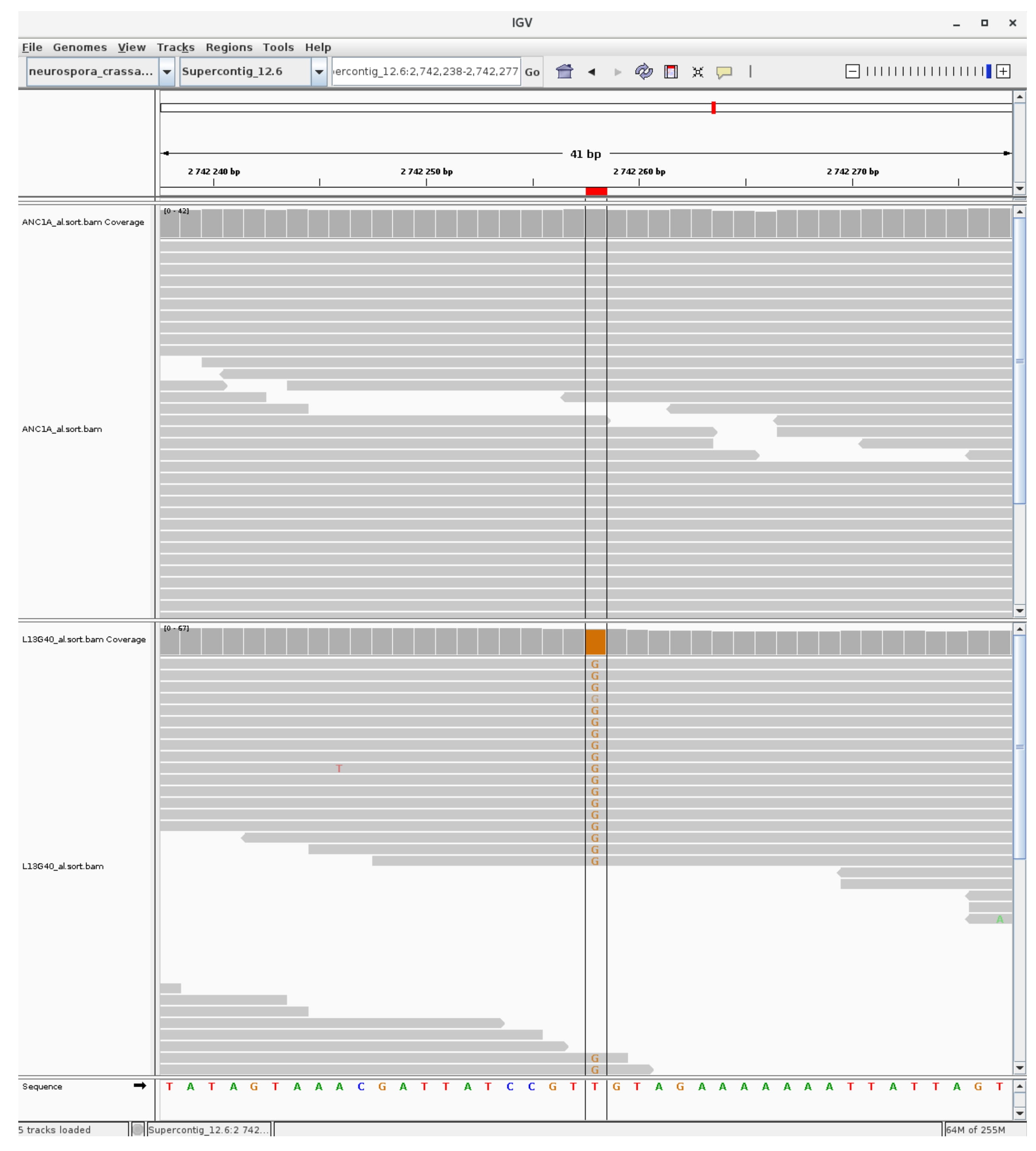

### mutation_H3K9_23.jpg

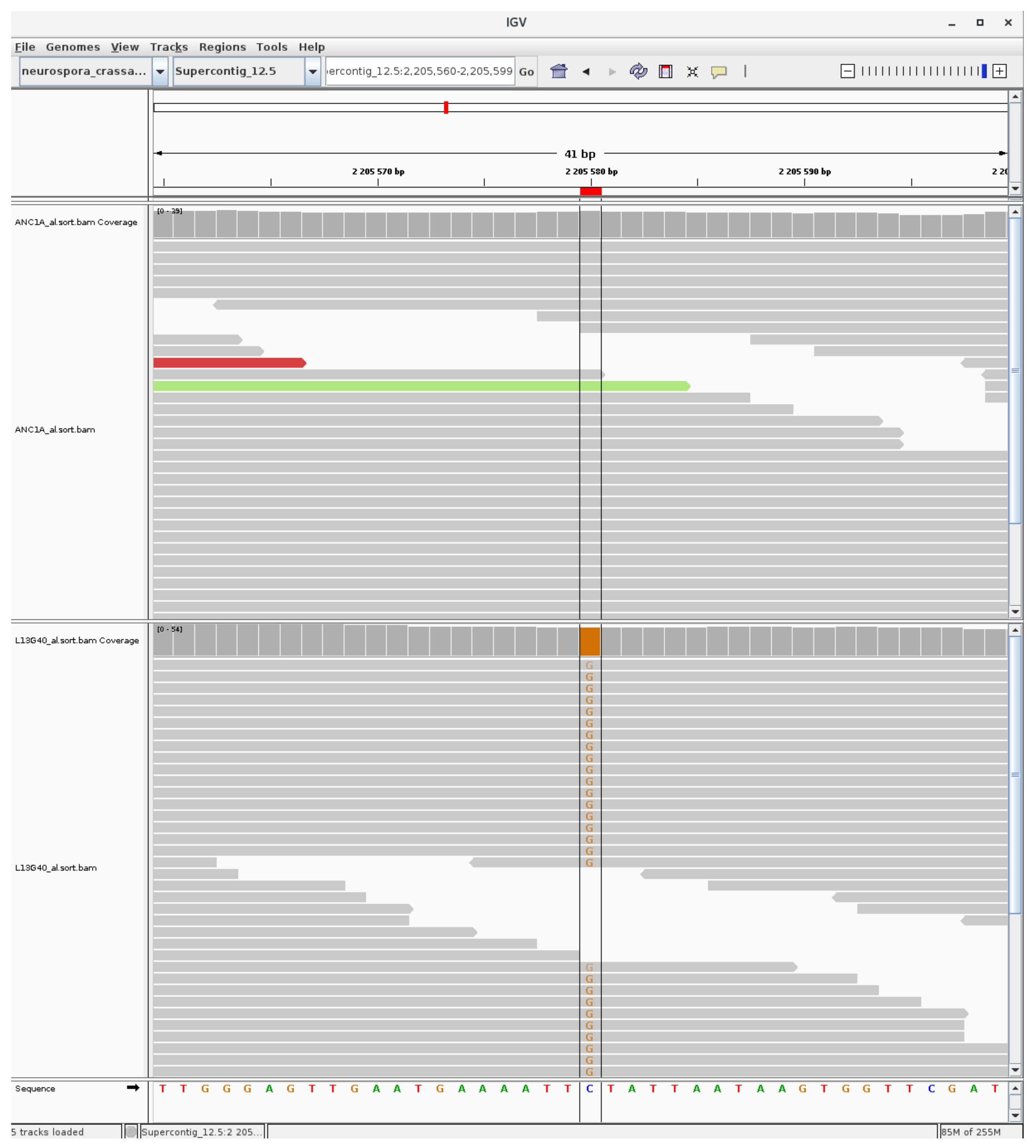

### mutation_H3K9_24.jpg

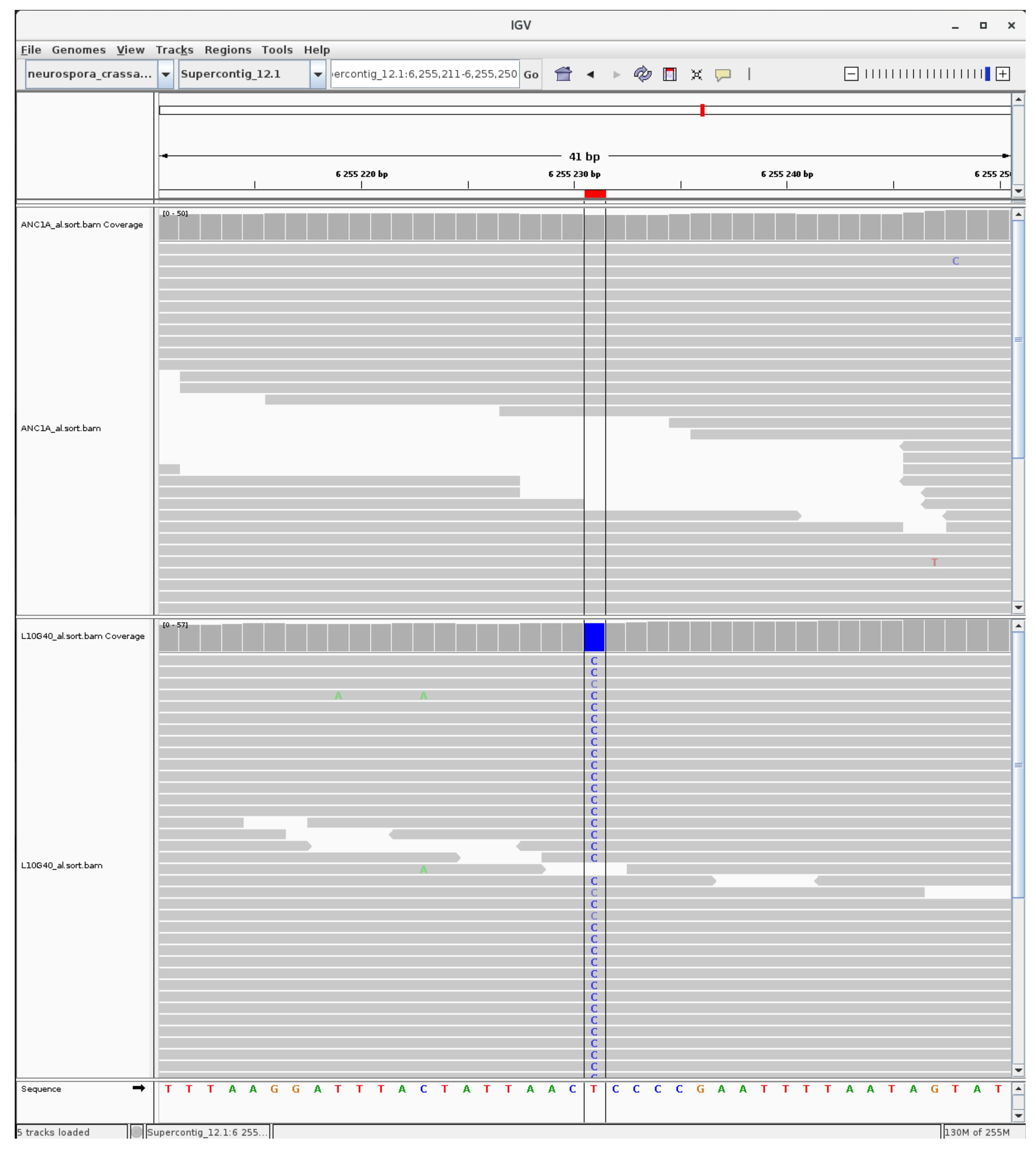

### mutation_H3K9_25.jpg

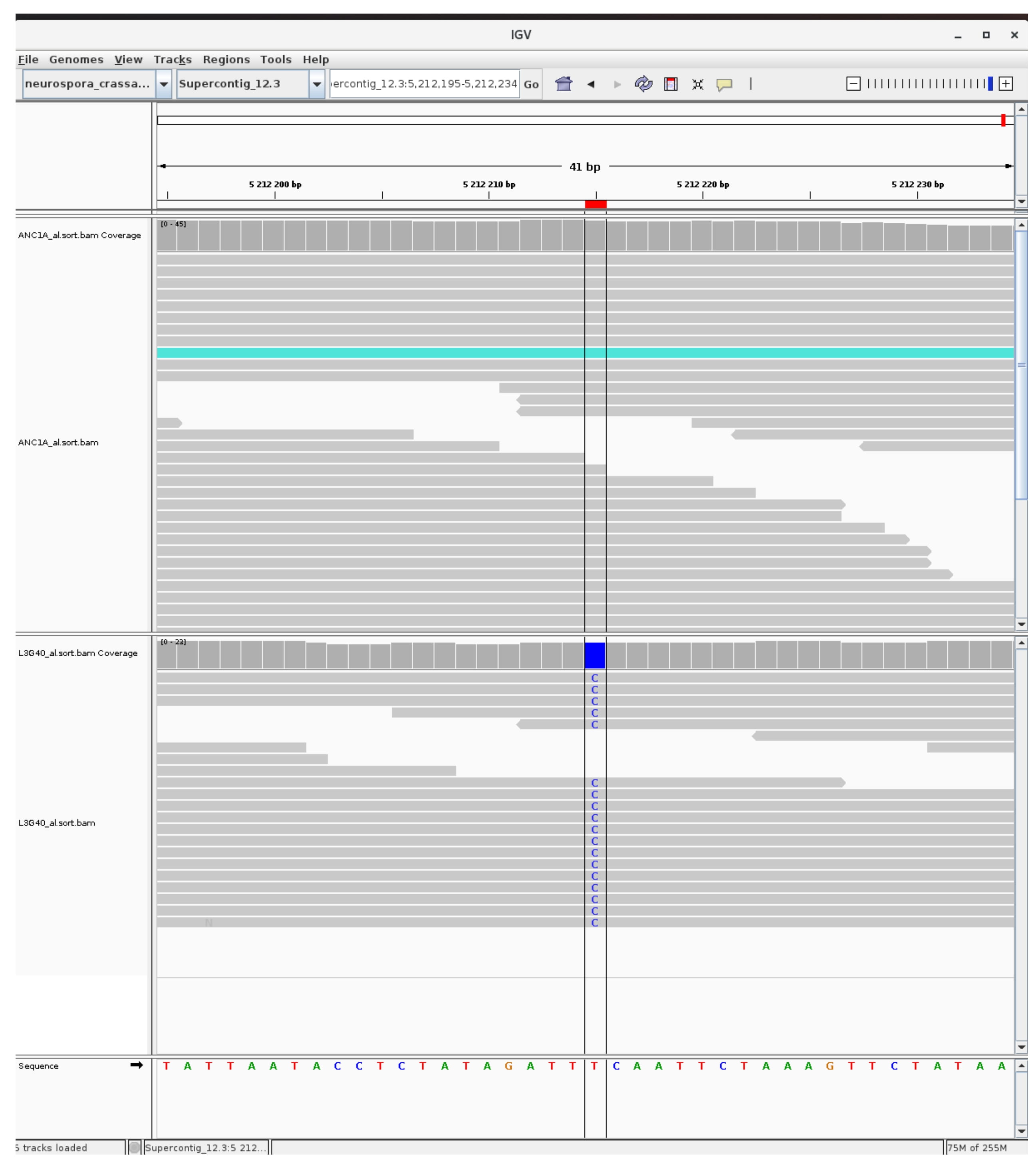

### mutation_H3K9_26.jpg

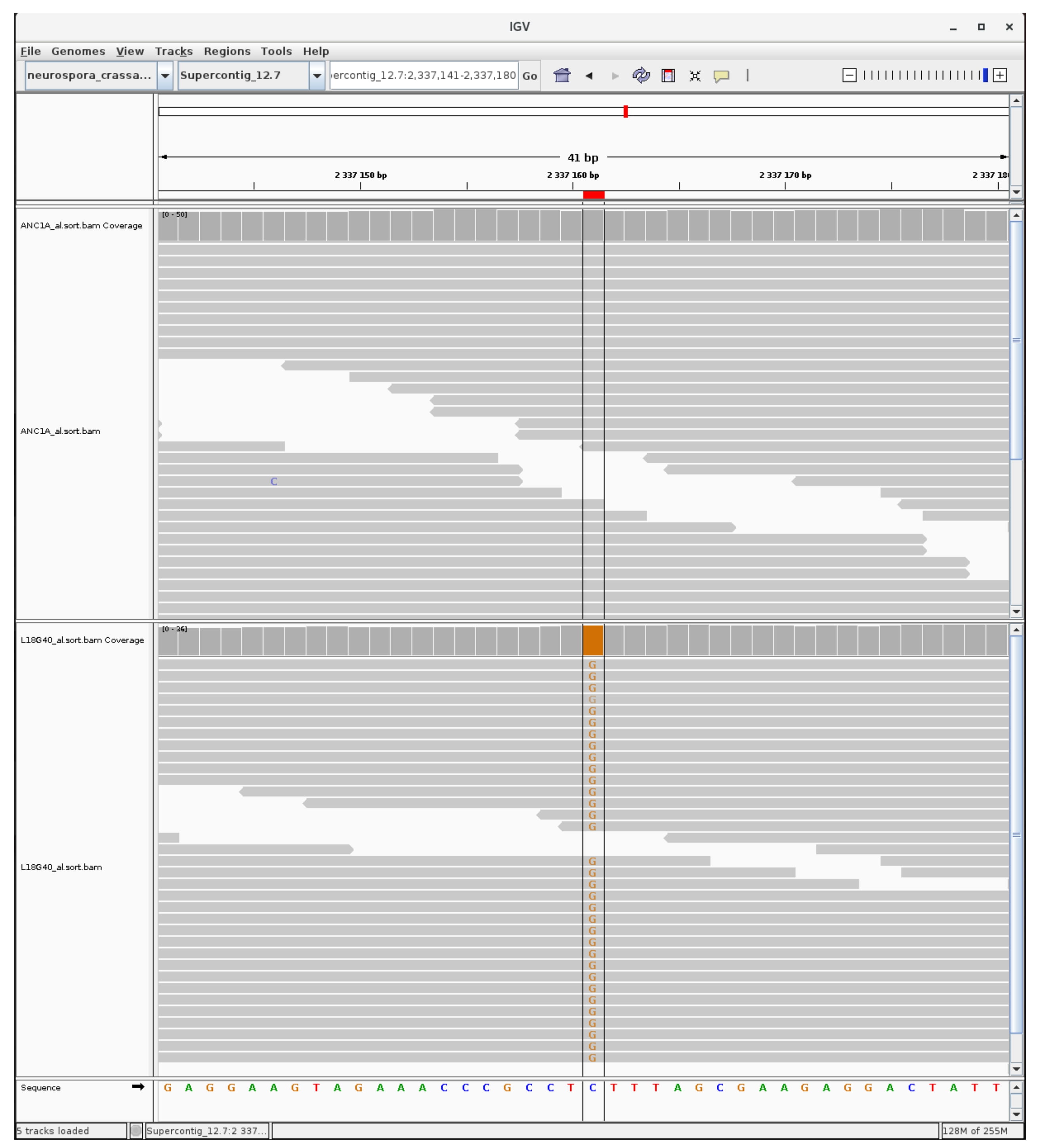

### mutation_H3K9_27.jpg

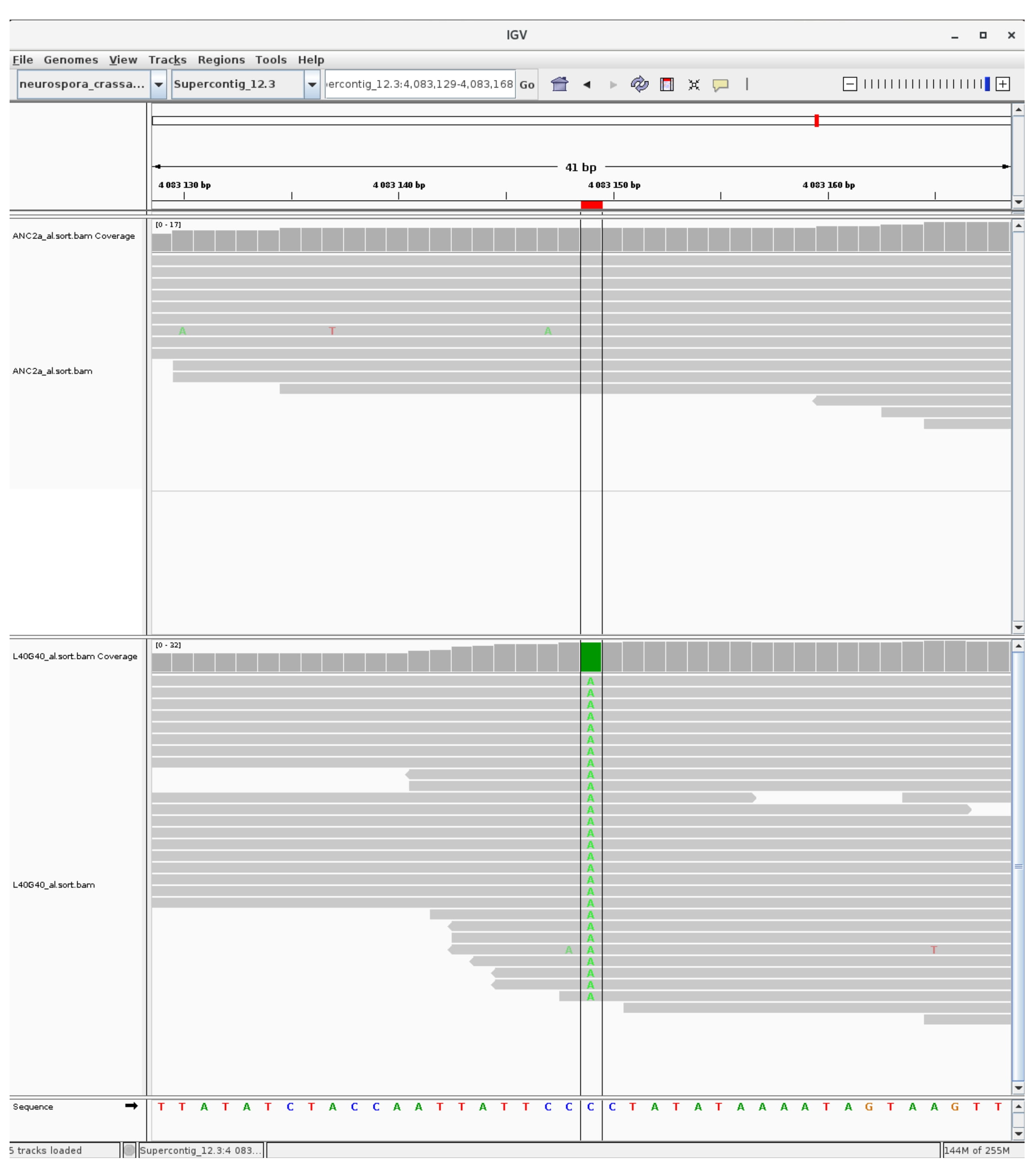

### mutation_H3K9_28.jpg

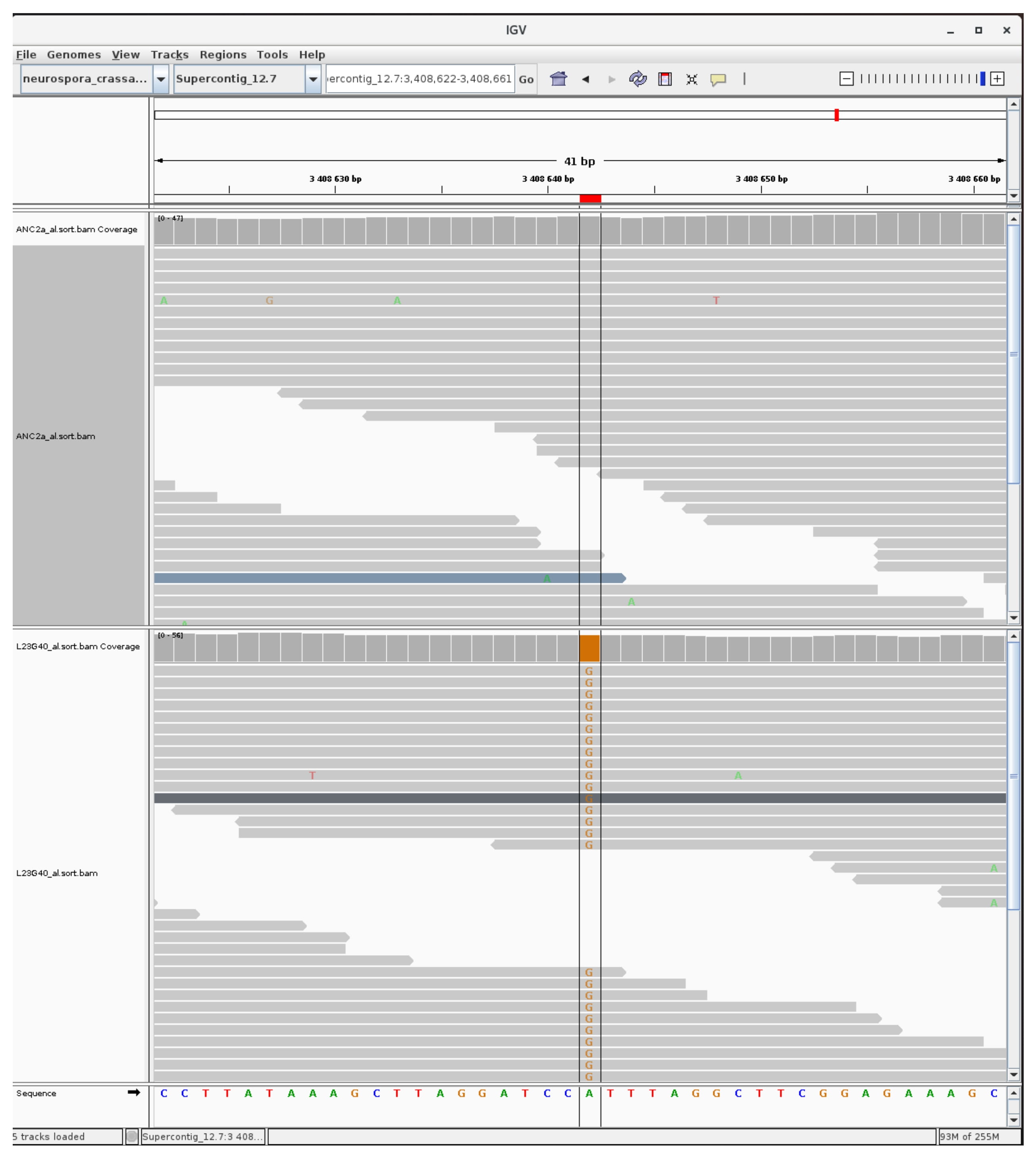

### mutation_H3K9_29.jpg

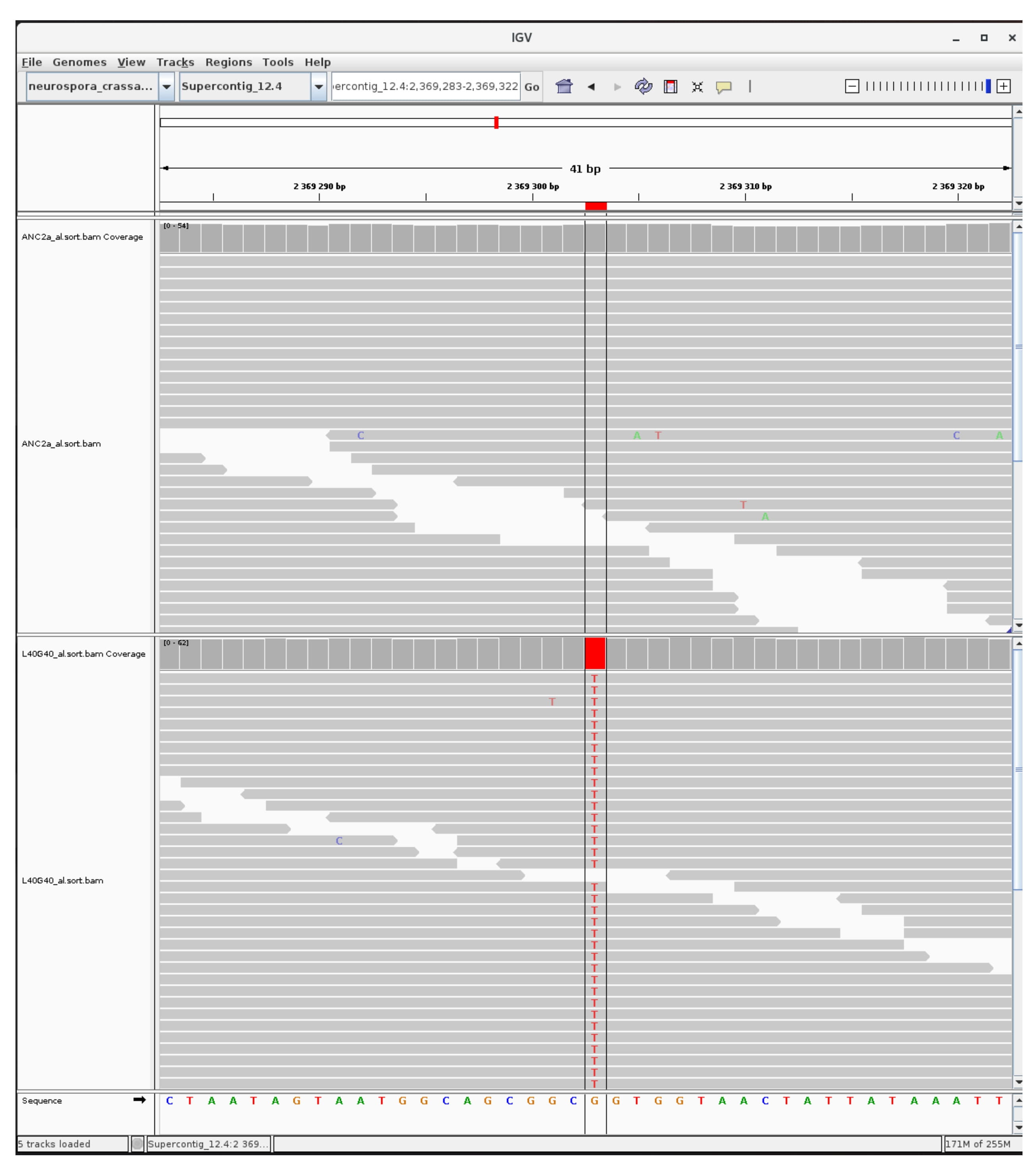

### mutation_H3K9_30.jpg

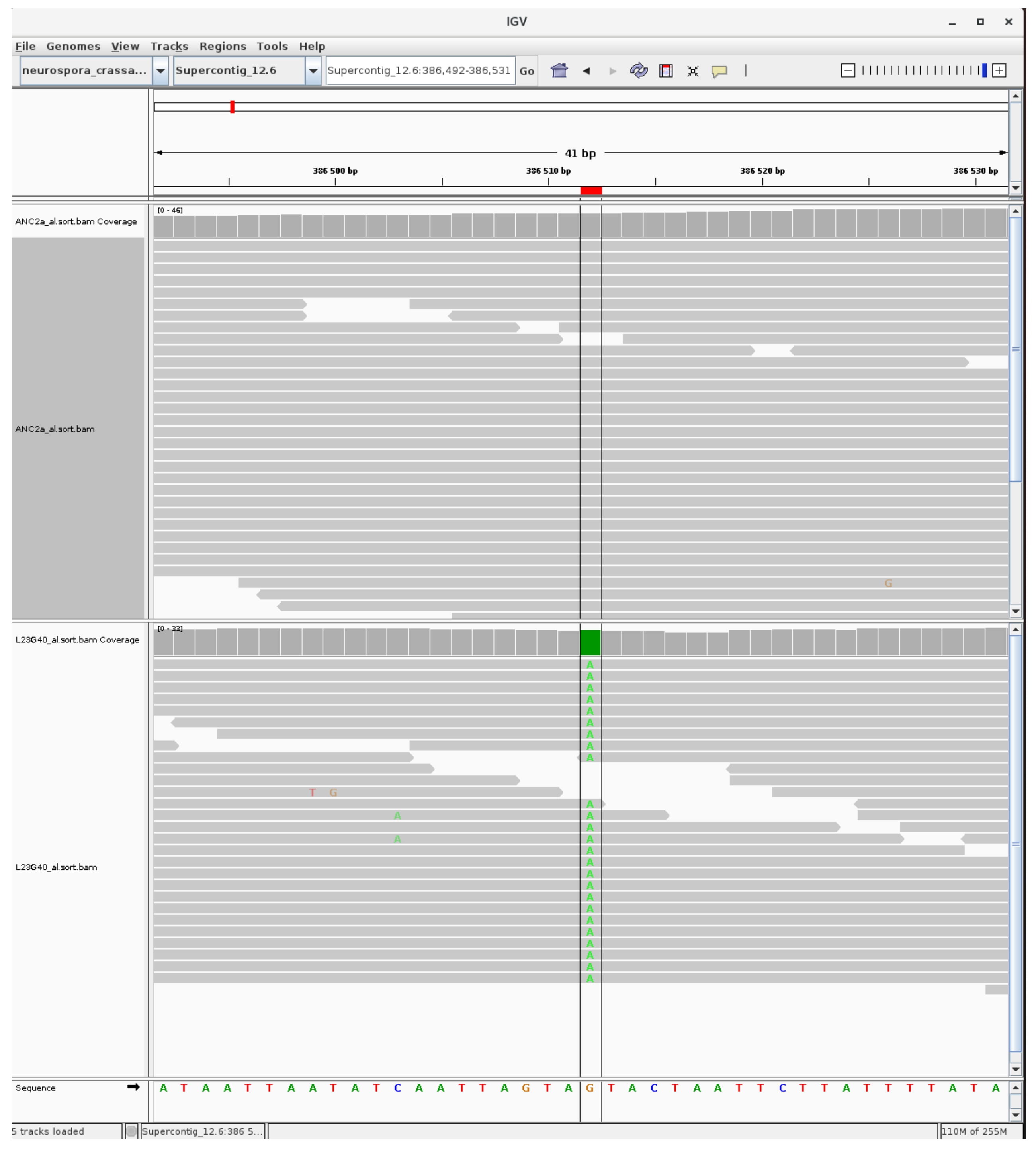
